# An AI-powered cloud biofoundry for autonomous biological research

**DOI:** 10.64898/2026.09.30.750244

**Authors:** Junyu Chen, Michael Volk, Saman Shafaei, Shih-I Tan, Vikas Upadhyay, Nilmani Singh, Xianrui Zhong, Katherine Arneson, Wenqi He, Chen Wang, Jingxia Lu, Aurosish Sharma, John McLaughlin, Matthew J. Berry, Steven Jarsen Harmon, Sophia Witola Reyes, George Heintz, Huimin Zhao

## Abstract

Autonomous experimentation, where an automated system designs, performs, analyzes, and iteratively improves experiments with minimal human intervention represents a transformative scientific research paradigm. However, it remains hampered by the disconnect between computational design and experimental execution. Here we present iCloudBiofoundry, a cloud-accessible, scalable, and self-evolving platform that bridges this divide. A scientific multi-agent system, iBioGenie, plans research workflows by converting natural language research objectives into testable hypotheses and experimentally grounded protocols using literature, databases, and computational tools. A physical AI layer, BiofoundryAI, translates them into executable workflows that run autonomously on distributed robotic biofoundries, and collects the data for iterative design. We demonstrate the end-to-end capabilities of iCloudBiofoundry across three representative case studies spanning enzyme retrieval, protein engineering, and metabolic engineering. By integrating agentic AI with physical AI, iCloudBiofoundry provides a cloud accessible platform that unifies computational design and robotic execution, enabling autonomous biological research.

The iCloudBiofoundry platform is available at: https://icloudbiofoundry.illinois.edu

## Main Text

By integrating artificial intelligence (AI), robotics, and real-time data analysis to enable closed-loop experimental workflows with minimal human intervention (*1–4*), autonomous experimentation or AI-powered autonomous laboratories can navigate vast, complex parameter spaces impractical to explore manually and is rapidly emerging as a new scientific discovery paradigm for biological, chemical, and materials research (*5–10*). In this paradigm, large language models (LLMs) and AI agents are used to plan, retrieve information, use computational tools, and carry out multi-step scientific tasks (*11–13*), while automated laboratories or biofoundries enable robotic experimental execution (*14, 15*). Recently, several agentic AI systems such as SciAgents (*16*), Robin (*17*), and CoScientists (*18*) have been developed to connect these capabilities, demonstrating that AI-driven scientific reasoning can be coupled to computational workflows and, in some cases, laboratory automation such as Biomni (*19*), ProtoPilot (*20*), and a GPT-5-driven autonomous lab (*21*). However, these efforts generally remain limited to specific tasks, narrow experimental workflows, or loosely connected components rather than providing a unified end-to-end environment. An autonomous experimentation system requires these capabilities to be integrated within a single platform that coordinates project context across computational and robotic experimental stages, generalizes across diverse scientific problems, and continuously incorporates new tools and capabilities as the field evolves (*22–26*). Moreover, while advanced computational systems are becoming increasingly accessible, access to automated biofoundries remains limited, making it difficult for many researchers to experimentally test and iterate on computationally generated designs at scale via robotic execution.

To address these limitations, here we present iCloudBiofoundry, an AI-powered cloud biofoundry platform that carries synthetic biology research from end-to-end, turning a natural language goal into experiments run on a set of remote biofoundries, and returning the measurements that inform the next design. We demonstrate iCloudBiofoundry across three distinct case studies: re-engineering of *Yersinia mollaretii* phytase (*Ym*phytase), discovery and directed evolution of aspartate decarboxylase (PAND) for improved 3-hydroxypropionic acid (3HP) production, and CRISPR-based metabolic engineering of *Saccharomyces cerevisiae* for β-carotene production. Across these studies, iCloudBiofoundry improved *Ym*Phytase activity by 5.28-fold over the previous best-engineered variant and by 70.17-fold over the wild type. Moreover, it discovered and further evolved novel PAND enzymes, achieving a 2.82-fold improvement in 3HP production. Lastly, it identified a double-gene deletion strategy that increased β-carotene production by 1.98-fold relative to the parental strain. These results demonstrate the effectiveness and versatility of iCloudBiofoundry for autonomous synthetic biology and contribute to the development of autonomous experimentation as an emerging paradigm for scientific discovery.

## Results

### iCloudBiofoundry as a scalable cloud platform for autonomous synthetic biology

iCloudBiofoundry connects agentic scientific reasoning with automated experimental execution (**Fig. 1 and fig. S1**). The system comprises four integrated components: (1) a unified web interface through which researchers initiate projects using natural language, review generated designs and experimental plans, monitor execution, and analyze returned results within a continuous session; (2) iBioGenie, a multi-agent system that interprets biological objectives, decomposes them into structured scientific actions, and runs workflows from relevant literature, tools, databases, models, and protocol-generation modules; (3) BiofoundryAI, a physical AI layer that interprets wet-lab protocols and translates them into validated workflows executable on a biofoundry; and (4) a set of automated biofoundries, including iBioFAB (*27*) and iBioFoundry (*15*), that run these workflows and return measurements to inform subsequent design. Collectively, these components couple agentic AI, physical AI, and distributed biofoundries into a single end-to-end loop for autonomous synthetic biology experimentation.

**Fig. 1.**
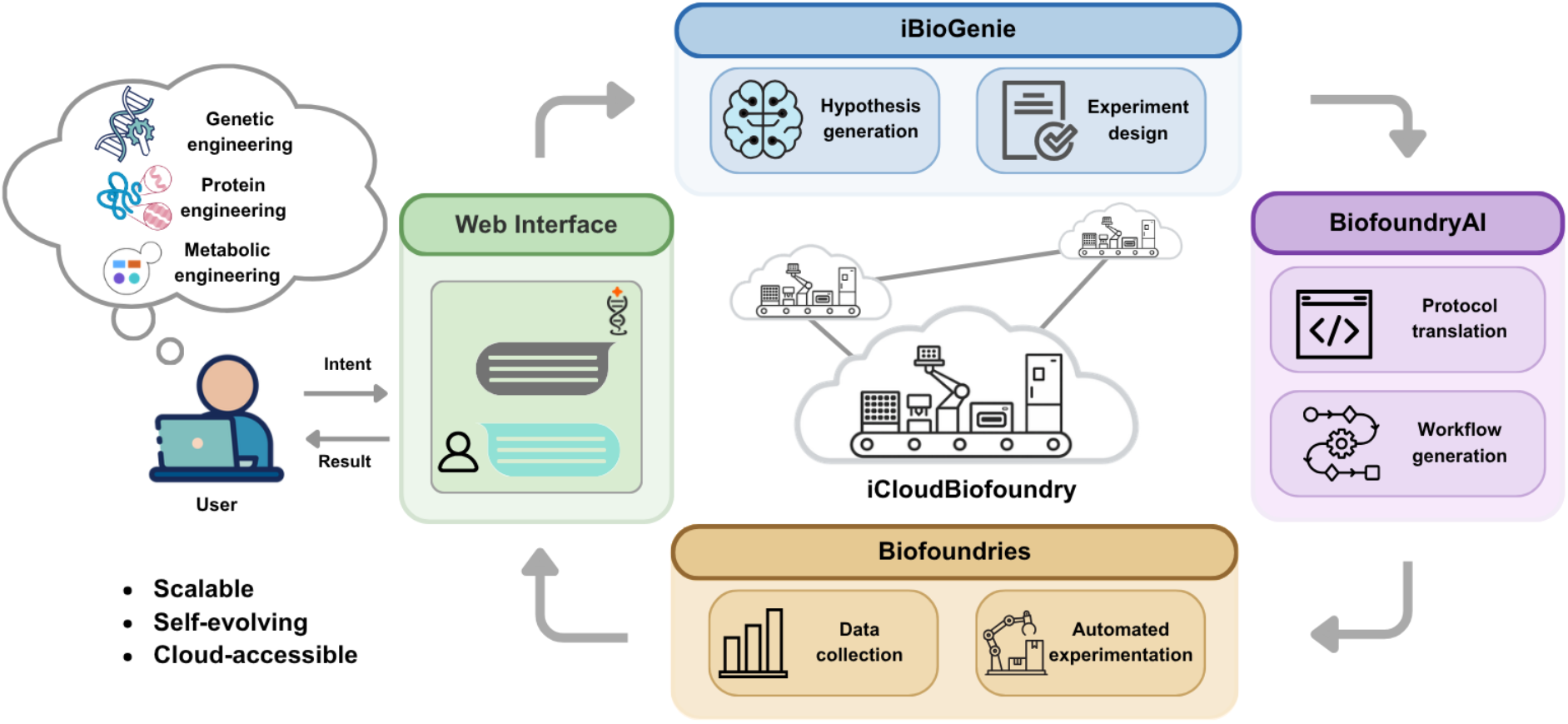
Architecture of the iCloudBiofoundry for autonomous biological research. iCloudBiofoundry connects scientific reasoning, computational design, protocol generation, and robotic execution into a single closed loop. This system comprises four components: (i) a unified **web interface**, through which researchers specify objectives in natural language and monitor jobs and results across biofoundry sites; (ii) **iBioGenie**, a multi-agent scientific intelligence layer that decomposes objectives into research tasks, invokes computational tools and databases, and generates experimentally grounded protocols; (iii) **BiofoundryAI**, a physical AI layer that compiles protocol-level plans into validated, simulation-tested automation workflows; (iv) **Biofoundries**, a network of distributed robotic infrastructure that executes the workflows and generates experimental measurements.

iCloudBiofoundry is designed to be a scalable, self-evolving platform rather than a fixed pipeline. iBioGenie can continuously expand and improve its capabilities through automated tool addition, incorporating validated new computational tools, databases, and specialized modules into its action space. In parallel, BiofoundryAI supports protocol translation across instruments, labware, and operational constraints, allowing workflows to be adapted across different compatible biofoundry environments. The cloud architecture connects multiple biofoundry sites within a single operational framework, allowing the platform to scale across its computational and physical layers. Additionally, the system uses experimental outcomes and user feedback to support automated continuous improvement for both iBioGenie and BiofoundryAI.

### Enabling computational and experimental design with iBioGenie

As the scientific intelligence layer of iCloudBiofoundry, iBioGenie is designed around the principle that biological research requires both a broad action space and a system capable of organizing that space into coherent workflows (**Fig. 2A**). The action space, defined as the capabilities available to the system, integrates computational tools, biological databases, literature, skills, code execution, and experimental procedures across major synthetic biology applications (**supplementary text 3.1 and 3.2**). Rather than relying on rigid workflows or loading every tool into the agent context, iBioGenie dynamically discovers, inspects, and invokes the capabilities needed for each research objective (**supplementary text 3.3**). This allows the system to remain flexible and extensible as new tools, databases, and procedures are added.

**Fig. 2.**
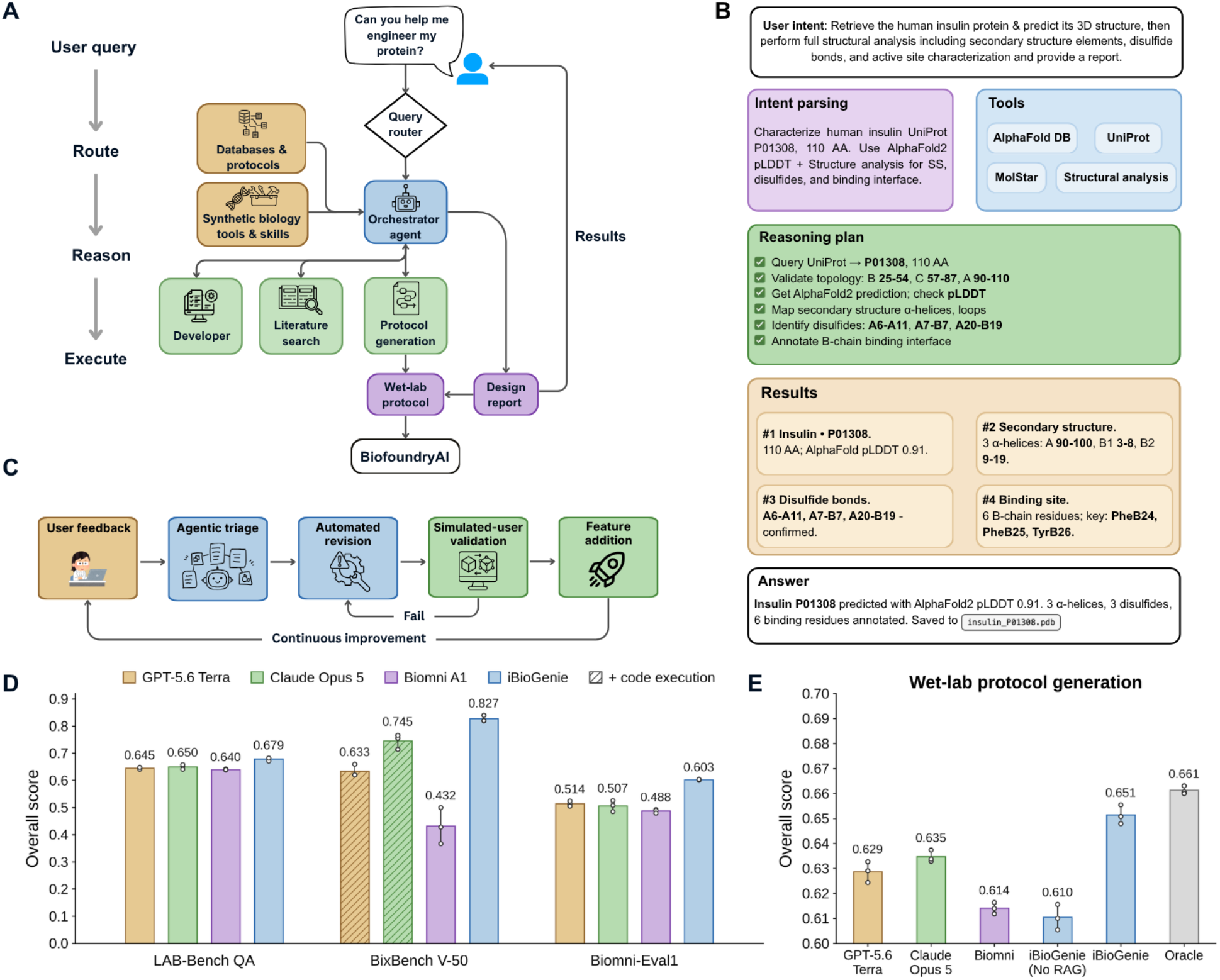
iBioGenie. **(A)** Overview of the iBioGenie multi-agentic AI architecture. User queries are routed to an orchestrator agent, which coordinates specialized agents, biological databases, curated protocols, and computational tools to generate design reports, and wet-lab protocols. (**B**) An example showing how iBioGenie parses user intent, selects tools, builds a reasoning plan, executes analyses, and returns a structured answer. (**C**) Continuous integration and self-improvement in iBioGenie via feedback from real-world use. (**D**) Benchmarking results comparing iBioGenie against baseline systems across, LAB-Bench QA subset with 1,096 questions, BixBench verified-50, and BiomniEval1 with 433 questions. iBioGenie and Biomni A1 are given GPT-5.6 Terra as their base model. For LAB-Bench and BiomniEval1, Claude Opus 5 and GPT-5.6 Terra use the base model without any tools; for BixBench, which requires code execution, they are run as coding agents using the benchmark’s baseline scaffold. Bars show means of three replicates, and error bars show the standard deviation. (**E**) Benchmarking results on the internal wet-lab protocol generation benchmark (139 questions) comparing iBioGenie protocol generation against baseline systems an Oracle reference that achieves the maximum possible score. Bars show means between 3 replicates, and error bars show the standard deviation.

On top of this action space, iBioGenie uses a multi-agent architecture to convert biological intent into executable research plans. A main orchestrator agent interprets the user’s objective, decomposes it into actionable scientific tasks, selects relevant tools and databases, and executes computational workflows (**fig. S10, supplementary text 3.4**). Other specialized agents and tools support this process by handling focused operations such as code execution, literature retrieval, protocol generation, and reporting (**figs. S11 and S12, supplementary text 3.5 and 3.6**). This architecture enables iBioGenie to connect high-level biological reasoning with reliable execution of targeted computational design tasks. Beyond executing the current action space, iBioGenie is designed to improve through continued use. Feedback submitted through the chat interface is linked to the corresponding session state and is automatically triaged into evidence-grounded engineering issues, which trigger revisions to tool implementations and behavioral features (**Fig. 2C, figs. S13 to S15, supplementary text 3.7 to 3.9**).

A representative interaction shows how iBioGenie works in practice (**Fig. 2B**). In this example, the user requests retrieval of human insulin, prediction of its 3D structure, and its structural characterization. iBioGenie parses the request into specific tasks, identifies the target sequence and executes a plan to assess topology, model confidence, secondary structure, disulfide bonds, and binding-site features. It returns a concise answer summarizing the main findings and provides the saved output.

We systematically evaluated iBioGenie across four complementary benchmark settings spanning biological reasoning, computational workflow execution, and wet-lab protocol generation. A subset of 1,096 LAB-Bench QA questions was used to examine performance on practical biology research tasks, including database-grounded question answering (DbQA), sequence–function inference (SeqQA), literature reasoning (LitQA2), protocol interpretation (ProtocolQA), and cloning scenarios (CloningScenarios) (*28, 29*) (**tables S1 and S2, supplementary text 4.2**). BixBench Verified-50 was included to test longer, multi-step computational biology workflows involving dataset exploration, analytical execution, and interpretation of complex outputs (*19, 30*) (**table S3, supplementary text 4.3**). Because BixBench requires code execution, the GPT-5.6 Terra and Claude Opus 5 baselines were run on it as coding agents using the benchmark’s own baseline scaffold, with their model-only runs retained as controls in the SI. BiomniEval1 provided a complementary assessment of tool-enabled biological problem solving across diverse biological tasks (*19*) (**table S4, supplementary text 4.4**). Finally, we developed an internal wet-lab protocol-generation benchmark to determine whether iBioGenie could convert biological objectives into experimental wet-lab procedures suitable for downstream execution (*31*) (**tables S5 and S6, supplementary text 4.5 and 4.6**). Across LAB-Bench, BixBench Verified-50, Biomni-Eval1, and the internal wet-lab protocol generation benchmark, iBioGenie achieved scores of 0.679, 0.827, 0.603, and 0.651, surpassing the strongest baseline by 2.9, 8.2, 8.9, and 1.6 percentage points, respectively (**Figs. 2D and 2E**). By supporting these time-intensive stages of research, iBioGenie can accelerate scientific decision-making and help researchers move more efficiently to actionable workflows.

### Connecting iBioGenie and robotic experimental execution with BiofoundryAI

iBioGenie generates a design and corresponding wet-lab protocol that a researcher could, in principle, follow at the bench. However, converting a wet-lab protocol into an executable automation workflow remains challenging, as each experimental step must be converted into precise, instrument-compatible operations and scripts. To address this challenge, we designed BiofoundryAI, a multi-agent physical AI system in which four specialized agents convert protocols designed by iBioGenie into executable workflows (**Fig. 3A, supplementary text 5.1**). A digital twin of the biofoundry, powered by the Momentum scheduler, encoding its instruments, labware, and status, is then used to validate and simulate workflows before execution (**supplementary text 5.2 and 5.3**). The BiofoundryAI orchestrator coordinates four specialized agents and deploys validated workflows to a distributed biofoundry platform (**Fig. 3B, figs. S8 and S9, supplementary text 2.1 to 2.4**). The planner agent translates wet-lab protocols into executable biofoundry workflows, specifying the required devices and labware. The script agent compiles these plans into platform-specific workflow scripts and instrument methods. The inspection agent identifies errors in the compiled workflows and issues targeted repair instructions. Finally, the biofoundry agent runs simulations and manages interactions with the physical platform (**supplementary text 5.1 to 5.4**).

**Fig. 3.**
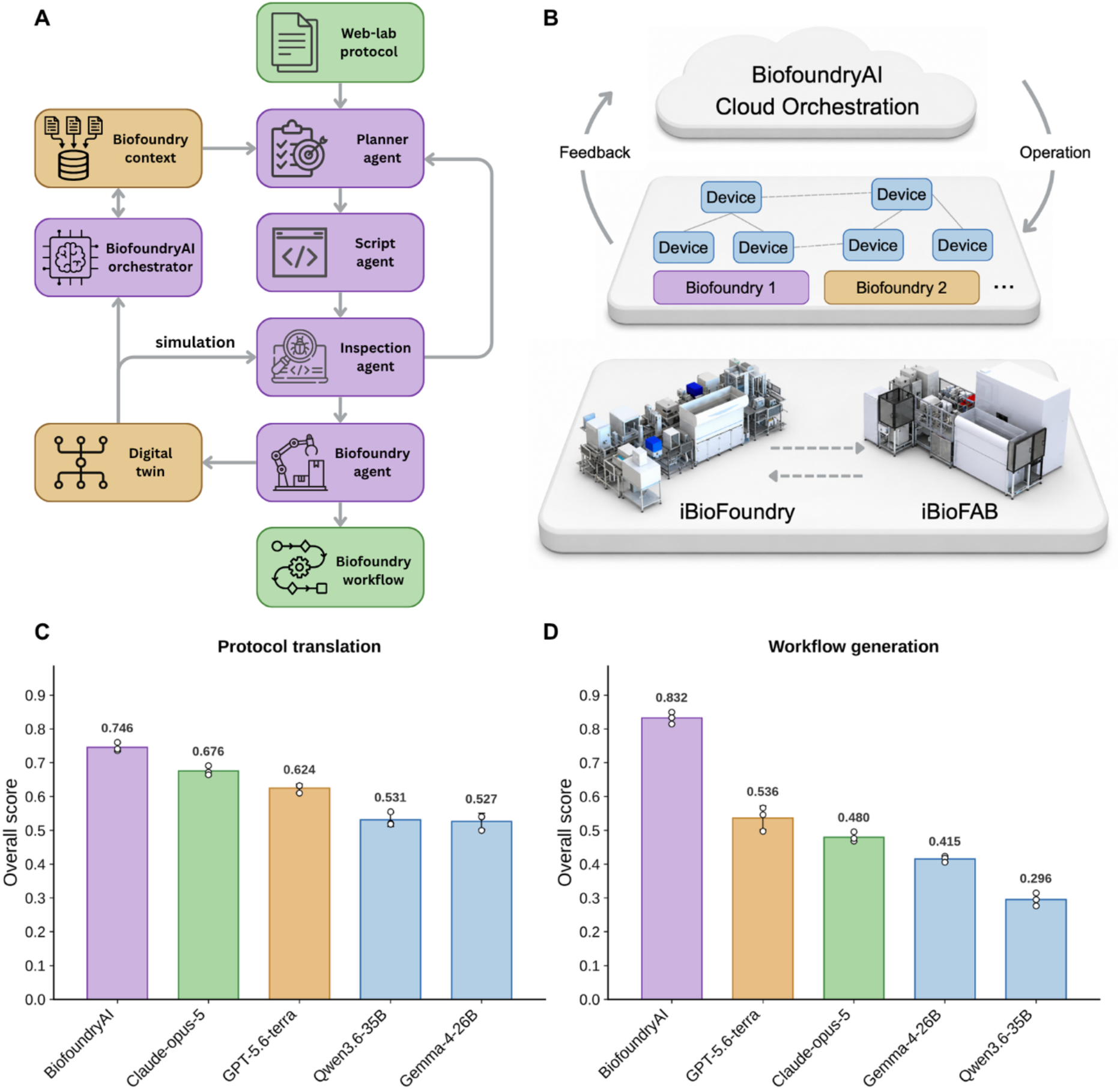
BiofoundryAI. **(A)** Overview of the BiofoundryAI physical AI architecture. Four specialized agents (Planner, Script, Inspection, and Biofoundry) coordinated by an orchestrator convert a natural language protocol into a validated, simulation-tested workflow. (**B**) Cloud connectivity across two biofoundries (iBioFAB, iBioFoundry) through the shared iCloudBiofoundry platform, with each site represented by its own instruments and capabilities. This also enables experiments to be distributed across biofoundries, with different stages executed at the facility best equipped for each task. (**C**) Level-1 benchmark results for protocol translation. Scores reflect device, operation, step sequence, labware, and parameter agreement with 40 hand-curated automation protocols. Bars show means between 3 replicates, and error bars show standard deviation. (**D**) Level-2 benchmark results for workflow generation. The overall score combines the digital twin’s assessment of workflow validity with the similarity between the automation script and the ground-truth script. Bars show means between 3 replicates, and error bars show standard deviation.

BiofoundryAI has three defining features: it is scalable across compatible facilities, accessible from the cloud, and self-evolving with use. First, agent behavior is configured through a structured context bundle that describes the target facility, including its device capabilities, plate templates, and unit-operation descriptions. Therefore, onboarding a new biofoundry means generating a context bundle rather than reprogramming the agents, which makes the platform easy to scale across compatible facilities (**supplementary text 5.5**). Second, the agents reach the physical instruments through the digital twin, so distributed biofoundries are accessible from the cloud through one common interface, and a protocol can be routed to whichever node carries the required capability (**Fig. 3B**). Third, every validated workflow is retained with its repair history and digital twin simulation outcome, allowing the agents that generate and check new workflows to learn from previous runs. In this way, BiofoundryAI improves as it is used (**supplementary text 5.6**).

To evaluate BiofoundryAI quantitatively, we constructed a two-level benchmark of 40 synthetic biology protocols (**table S8, supplementary text 6.1**). Level-1 tests translation of a platform-agnostic wet-lab protocol into a biofoundry automation protocol with explicit device and operation assignments; Level-2 tests the harder task of compiling that automation protocol into an executable workflow script, evaluated against the ground truth that was verified by human experts and the digital twin (**tables S9 to S11**). BiofoundryAI achieved a 0.746 overall score on protocol translation outperforming the best baseline model by 7% (**Fig. 3C, supplementary text 6.3**). When generating the workflow from the translated protocol, BiofoundryAI achieved 0.832 overall score, outperforming the best baseline model by 30% (**Fig. 3D, tables S13 to S16**).

With BiofoundryAI enabling the translation of experimental protocols into executable automation workflows, we sought to demonstrate the utility and versatility for autonomous synthetic biology. Specifically, we selected three case studies with different levels of complexity in computational design and experimental execution, including the re-engineering of *Ym*Phytase, discovery and directed evolution of PAND for improved 3HP production, and CRISPR-based metabolic engineering of *S. cerevisiae* for β-carotene production.

### Designing improved phytase enzyme variants using existing experimental data

Phytase is an industrially important feed enzyme that hydrolyzes phytic acid, improving phosphate availability, and reducing phosphorus waste in animal production (*32*). However, many phytases exhibit sharply reduced activity near neutral pH, limiting their performance across the variable pH environments encountered during feed processing and gastrointestinal digestion (*33, 34*). *Ym*Phytase had previously been engineered through four autonomous DBTL cycles, generating a multi-round dataset of sequence variants and measuring near-neutral pH activity (*7*) (**Fig. 4A**). The improvement from the final round of the previous study began to plateau, suggesting that the campaign was approaching saturation and making further optimization a challenging test case for data-driven re-engineering.

**Fig. 4.**
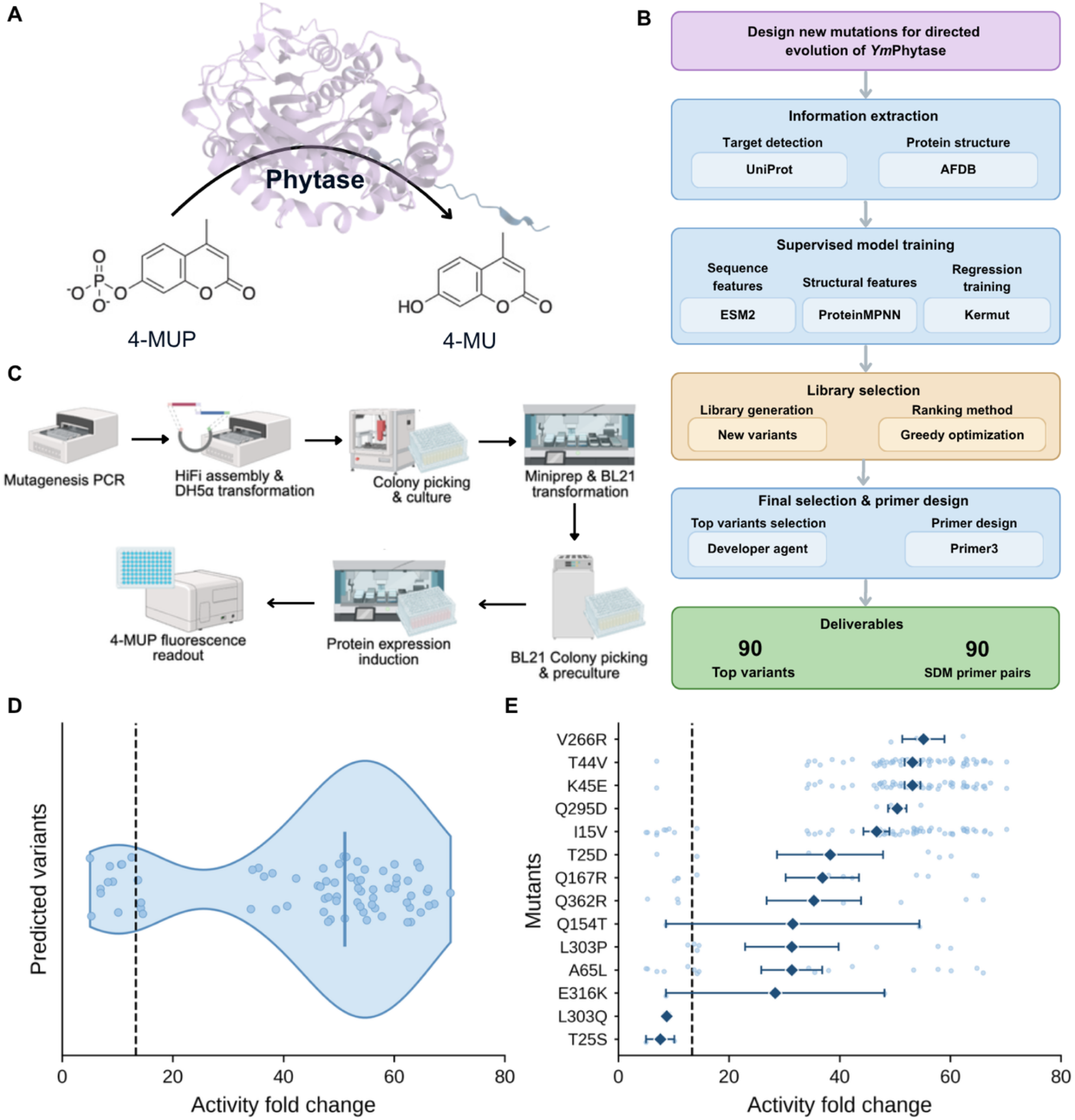
Automated, agent-driven enzyme engineering to improve phytase activity. **(A)** A previously engineered high-activity *Ym*Phytase variant, catalyzing phytate hydrolysis to release inorganic phosphate, served as the starting point for further optimization. **(B)** iBioGenie computational pipeline for designing new mutations for directed evolution of *Ym*Phytase. Tools used in this workflow include Uniprot (*40*), AlphaFoldDB (*41*), ESM2 (*42*), ProteinMPNN (*43*), Kermut (*44*) and Primer3 (*45*). **(C)** Automated experimental workflow generated by BiofoundryAI for SDM, from variant construction to activity measurement. **(D)** Per-variant activity of the round-5 *Ym*Phytase library relative to the wild-type enzyme. The vertical line marks the median and the dashed line marks the previous best. The best variant reached 70.17-fold over the wild-type enzyme. **(E)** Activity of designed variants containing each mutation. Individual points show the activity fold change of variants carrying the indicated mutation, diamonds show the mean, and error bars indicate the standard error of the mean. Mutations are ordered by mean activity, and the dashed line marks the activity of the previous best variant.

We therefore used the existing experimental dataset as the starting point for further optimization. iBioGenie was provided with the complete activity data from the previous campaign and tasked with collecting information about the enzyme and training a supervised model to design a new library of higher-order mutants. It was then tasked with designing corresponding site-directed mutagenesis (SDM) primers and a wet-lab protocol for library construction and screening (**Fig. 4B, table S17, supplementary text 7.1**). BiofoundryAI then translated the designs into an automation workflow, integrating primer handling, mutagenesis PCR, *Dpn*I digestion, HiFi DNA assembly, transformation, colony picking, plasmid preparation, protein expression, crude lysate preparation, and fluorescence-based phytase activity screening using 4-MUP hydrolysis (**Fig. 4C, supplementary text 7.2**). The new library designed by iBioGenie showed a high hit rate, with 64 of 75 tested variants outperforming the best variant from the previous study. The top variant achieved a 5.28-fold improvement over the previous best variant and a 70.17-fold improvement relative to wild-type *Ym*Phytase (**Figs. 4D and 4E, supplementary text 7.2**).

### Enzyme discovery and engineering for improved 3HP production

3HP is an industrially important platform chemical used in the production of acrylic acid, adhesives, coatings, and biodegradable plastics (*35, 36*). We previously engineered an acid-tolerant yeast *Pichia kudriavzevii* SD108 for cost-effective 3HP production through the β-alanine pathway and identified aspartate decarboxylase as a major rate-limiting enzyme (*37*) (**Fig. 5A**). To further improve 3HP production, we sought to identify a PAND enzyme with higher activity than the previously characterized enzyme and subsequently optimize it through an additional round of enzyme engineering.

**Fig. 5.**
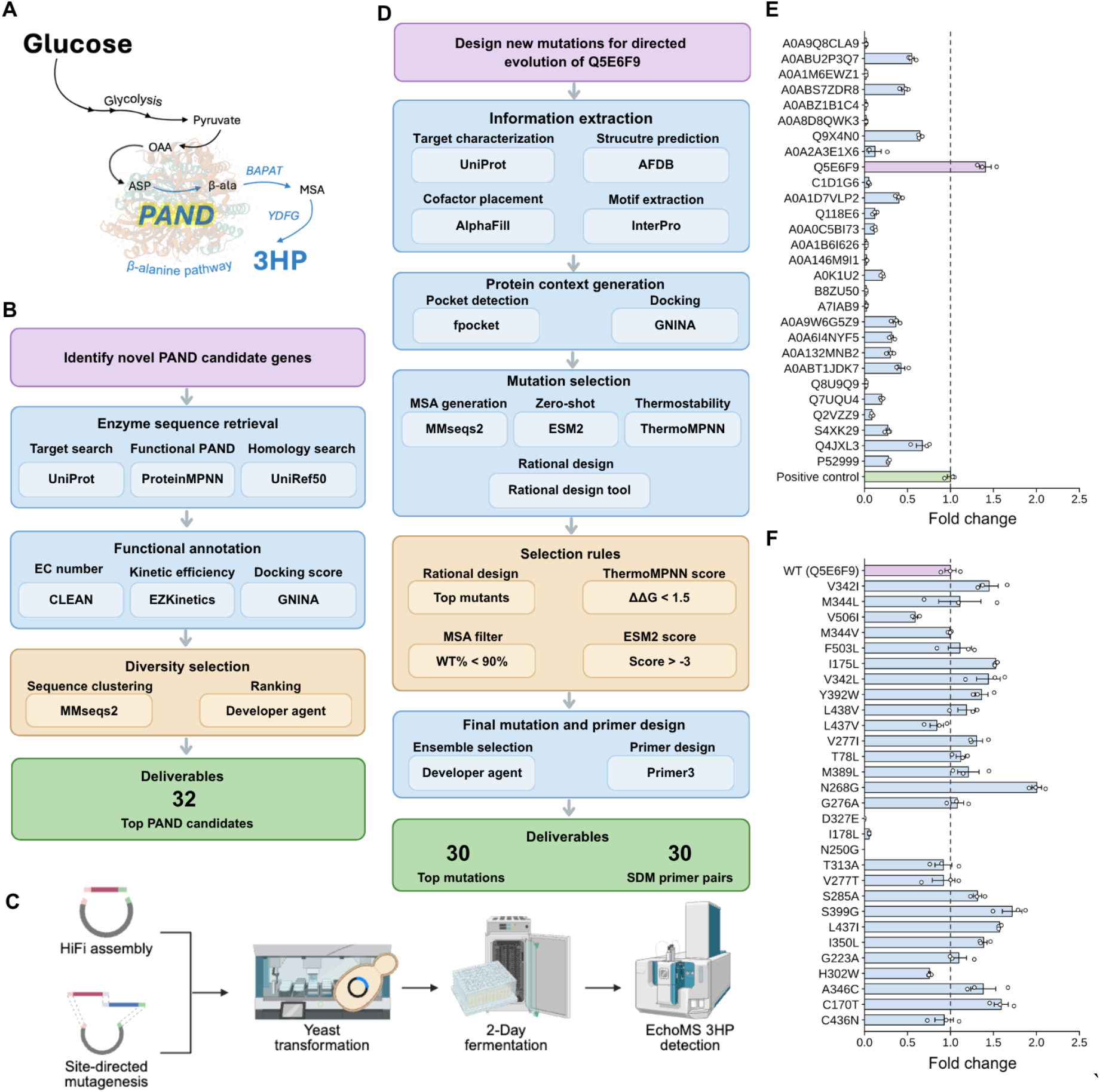
Closed-loop automated, agent-driven PAND enzyme discovery and iterative engineering. **(A)** Overview of the β-alanine pathway for 3HP biosynthesis, in which PAND catalyzes the conversion of aspartate to β-alanine. **(B)** First-iteration workflow designed by iBioGenie for retrieving novel PAND enzymes, yielding 32 candidates for experimental evaluation. Tools used in this workflow include Uniprot (*40*), BRENDA (*46*), UniRef50 (*47*), CLEAN (*48*), EZKinetics, GNINA (*49*), and MMseq2 (*50*). **(C)** Automated experimental workflow generated by BiofoundryAI from construction to EchoMS-based 3HP detection. **(D)** Second-iteration workflow for directed evolution of the top retrieved PAND (Q5E6F9) enzyme, producing 30 candidate mutations and corresponding SDM primers. Tools used in this workflow include Uniprot (*40*), AlphaFoldDB (*41*), AlphaFill (*51*), InterPro (*52*), fpocket (*53*), GNINA (*49*), MMseq2 (*50*), ESM2 (*42*), ThremoMPNN (*54*), iBioGenie’s internal rational design tool, and Primer3 (*45*). **(E)** Experimental screening of the retrieved PAND candidates, with 3HP production reported as fold change and benchmarked against the positive-control enzyme. Bars show the means of 3 biological replicates, and error bars show the standard deviation. **(F)** Per-variant activity of engineered Q5E6F9 variants relative to the retrieved enzyme. Bars show the means of 3 biological replicates, and error bars show the standard deviation.

We first tasked iBioGenie to identify novel PAND enzymes. iBioGenie then generated a computational discovery workflow integrating enzyme sequence retrieval from multiple databases, homology screening, functional annotation, catalytic-efficiency prediction, docking-based ranking, and diversity-aware clustering, ultimately nominating 32 candidates for experimental evaluation and a corresponding wet-lab protocol (**Fig. 5B, table S18, supplementary text 8.1**). BiofoundryAI then translated this protocol into an automated screening workflow incorporating HiFi DNA assembly, transformation, colony picking, miniprep, yeast transformation, fermentation and EchoMS-based 3HP detection (**Fig. 5C, supplementary text 8.2**). Of the 32 predicted enzymes, 28 were successfully transformed into *P. kudriavzevii*, and 18 produced detectable 3HP, corresponding to a 64% hit rate among successfully tested candidates. The best retrieved candidate increased 3HP production by 1.41-fold relative to the previously best known PAND enzyme (**Fig. 5E, supplementary text 8.2**). Building on this result, iBioGenie performed a second design iteration focused on engineering the best retrieved PAND enzyme, integrating target enzyme active site characterization, protein structure retrieval, motif extraction, substrate docking, rational design, multiple-sequence-alignment (MSA) based filtering, and two machine-learning-based zero-shot prediction models to nominate 30 single-site mutations and their corresponding primers for site-directed mutagenesis (**Fig. 5D, table S19, supplementary text 8.3**). BiofoundryAI’s generated automation workflow was then executed, and screening identified 19 variants with further improvements in 3-HP production relative to the retrieved parental enzyme (**supplementary text 8.4**). The best-performing variant achieved a 2.00-fold increase over the retrieved enzyme in the previous round, corresponding to a 2.82-fold overall improvement relative to the original starting enzyme (**Fig. 5F, supplementary text 8.4**).

### Metabolic engineering of *S. cerevisiae* for β-carotene production through combinatorial gene knockout

β-carotene is a high-value terpenoid and one of the most extensively engineered carotenoids in yeast. Over the past two decades, heterologous expression of *crtE, crtYB*, and *crtI*, alongside the native mevalonate pathway, in *Saccharomyces cerevisiae* has increased titers from trace levels to several hundred mg/L (*38, 39*). However, most of these gains came from manipulating a small set of genes at a time, yet the number of host modifications that could further redirect flux toward β-carotene is far larger than any single laboratory can construct and screen. We therefore used iCloudBiofoundry to nominate, build, and evaluate combinations of double-gene deletions that increase β-carotene production.

Starting from a base strain CEN.PK113-7D with the β-carotene pathway integrated into the genome and with additional Δ*CIT2*, we instructed iBioGenie to find additional gene deletions for improving β-carotene production using a multipronged literature and modeling approach (**Fig. 6A**). The final gene list contained 14 unique gene knockouts applied to the Δ*CIT2* strain. iBioGenie then designed *Sa*Cas9 gRNAs and sent the design to BiofoundryAI to generate the automation workflow, which included gRNA plasmid assembly, *E. coli* transformation, plasmid extraction, and yeast transformation for gRNA efficiency screening (**Fig. 6B, tables S20 to S22, supplementary text 9.1**). Of the 74 gRNAs tested across two rounds of screening, 21 were efficient and collectively targeted all 14 genes (**Fig. 6D**). Finally, iBioGenie was used to design primers for full open reading frame (ORF) deletions and then the BiofoundryAI workflow for yeast gene knockout was executed on the biofoundries (**Fig. 6C, supplementary text 9.2**). Eleven of the 14 targeted knockout mutants were successfully constructed and screened after fermentation by measuring absorbance at 454 nm normalized to OD600. The best mutant achieved a 1.69-fold increase in β-carotene production from the single knockout strain, and 1.98-fold increase compared to CEN.PK113-7D with the β-carotene pathway (**Fig. 6E, supplementary text 9.2**).

**Fig. 6.**
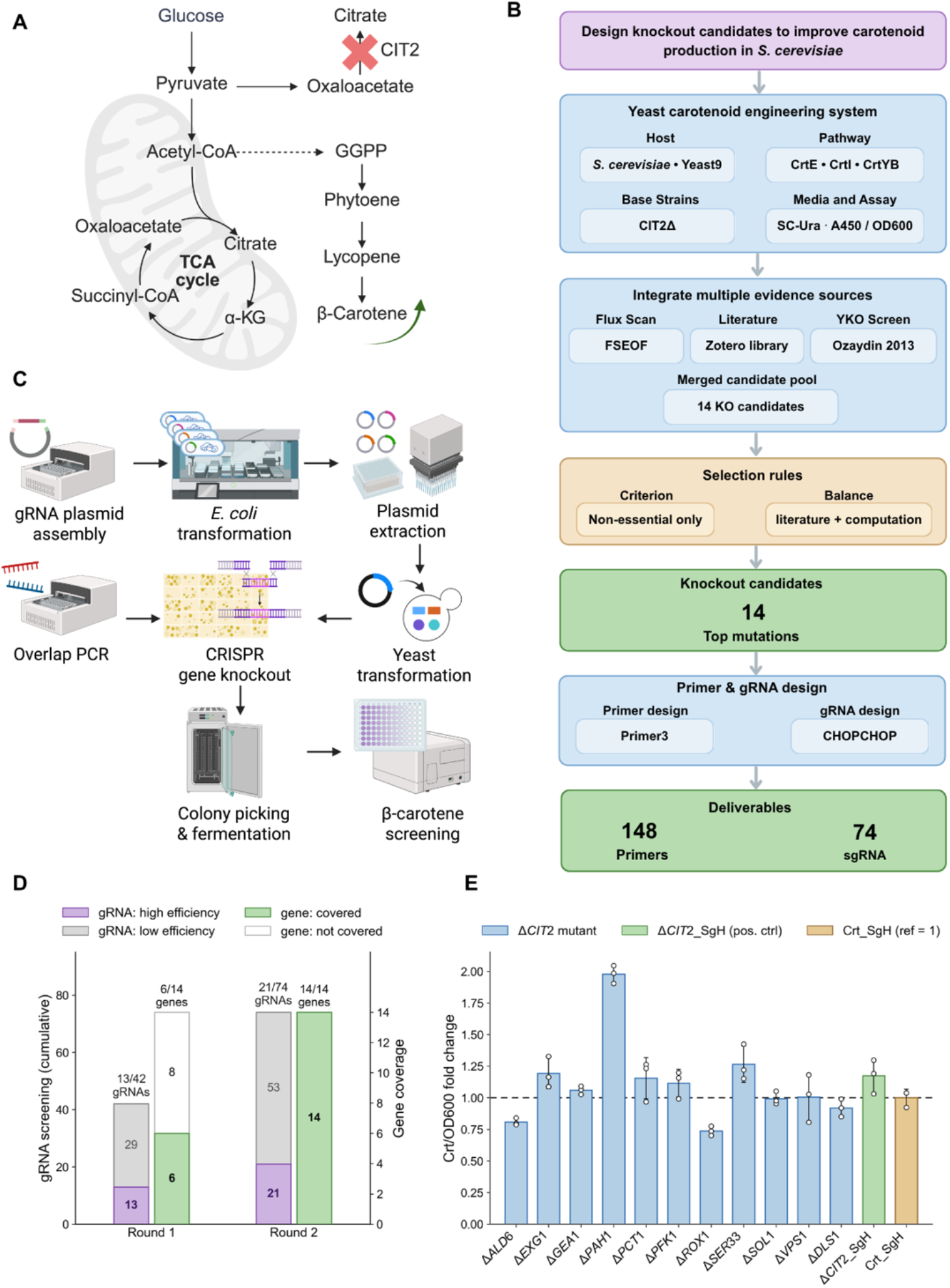
Agent driven metabolic engineering of *S. cerevisiae* through combinatorial gene knockout. **(A)** Overview of the β-carotene synthesis pathway and the Δ*CIT2* strain. (**B**) iBioGenie pipeline for target gene selection and CRISPR gRNA design. Tools used in this workflow include Yeast9 (*55*), COBRApy (*56*), FSEOF (*57*), the Saccharomyces Genome Database (*58*), the yeast deletion-collection carotenoid screen of Özaydin et al. (*59*), iBioGenie’s internal gene-candidate aggregation and homology-donor design tools, CHOPCHOP (*60*), and Primer3 (*45*). (**C**) Overview of the BiofoundryAI generated workflow for combinatorial yeast gene knockout. (**D**) CRISPR gRNA screening in two rounds. Cumulative data across two rounds are shown for gRNA efficiency and target genes covered by efficient gRNA. (**E**) β-carotene production of the final constructed 11 mutants. Production was measured by absorbance at 454 nm, normalized to cell density (OD600), and reported as a fold change relative to the β-carotene-producing strain (Crt_SgH). Bars show the means of 3 biological replicates, and error bars show the standard deviation.

## Discussion

Through systematic computational benchmarking and three experimentally validated case studies involving complex computational and experimental workflows, we demonstrate that iCloudBiofoundry provides a versatile framework for autonomous synthetic biology research. iCloudBiofoundry outperforms comparable agentic systems in synthetic-biology-relevant tasks and closes the key gap between computational design and robotic execution. Across the three campaigns, the agent used approximately 30 computational tools and databases during design, and around 20 instruments and 1000 scheduled operations during execution (**supplementary table S25**). These results establish a practical foundation for diverse autonomous synthetic biology experimentation by connecting AI-driven computational design with experimentally validated robotic execution in a unified research workflow.

Despite the advances of the iCloudBiofoundry platform, several limitations remain that can be further improved. The current capabilities of the system are still focused on synthetic biology, and future versions should expand to support broader scientific workflows, additional computational design modules, and integration with more biofoundry sites. Scaling the system will also require better large-scale planning, improved token efficiency, and more efficient access to GPU and cloud compute, especially for expensive ML-driven design campaigns that require repeated model inference, simulation, or optimization. Broader deployment will also require stronger safeguards for prompt injection, data privacy, biosafety, and provenance tracking, particularly when sensitive biological designs, proprietary datasets, or external model providers are involved. At the experimental layer, troubleshooting biofoundry execution remains challenging because many failures are difficult to predict before physical runs. Future systems may benefit from richer execution logs and vision-based monitoring of instruments. Finally, while the current digital twin supports pre-execution validation, future work should develop a digital twin that models the physical environment, derives empirical failure modes, and responds to real-time device feedback.

## Acknowledgments

This work used Delta at University of Illinois Urbana-Champaign through allocation BIO240329 from the Advanced Cyberinfrastructure Coordination Ecosystem: Services & Support (ACCESS) program, which is supported by U.S. National Science Foundation grants #2138259, #2138286, #2138307, #2137603, and #2138296. We thank the user testers who evaluated the iCloudBiofoundry system and provided feedback that guided its refinement. We thank the Momentum software team at Thermo Fisher Scientific for developing the developmental application programming interface (API) that enabled programmatic control of biofoundry scheduling in this work.

## Author contributions

H.Z. conceived the iCloudBiofoundry platform and supervised the project. M.V., S.S., and V.U. developed the iBioGenie multi-agent systems. J.C., X.Z., and J.M. developed the BiofoundryAI. W.H., C.W., K.A., M.V., and M.B. developed the web platform, cyberinfrastructure, and user interface. V.U., J.C., X.Z., M.V., S.S. designed and ran the benchmarks and evaluation framework. N.S., S.S., and S.H. carried out the phytase case study. S.I.T., S.S., J.L., J.C., N.S., S.H., carried out the PanD case study. J.C., M.V., and A.S. carried out the β-carotene metabolic engineering case study. All authors discussed the results and contributed to the final manuscript.

## Code availability

The source code and associated resources will be made publicly available upon acceptance of this manuscript for publication. All platform dependencies are open-source with the exception of the Momentum automation software, which is a commercial product of Thermo Fisher Scientific.

## Data and materials availability

All data are available in the main text or the supplementary materials.

## Supplementary Materials

Materials and Methods

Supplementary figures, text, and references

