## Supplementary Materials for "An AI-powered cloud biofoundry for autonomous biological research"

**The PDF file includes:**

Materials and Methods
Supplementary Text
Figs. S1 to S44
Tables S1 to S25
References

**Other Supplementary Materials for this manuscript include the following:**

**Table of Content**

1. General information: iCloudBiofoundry application 4
    1.1. System overview 4
    1.2. System architecture 4
    1.3. Federated identity and access management 5
    1.4. Deployment and operations 6
    1.5. Cloud access and scalability 6
    1.6. Data management and privacy 7
    1.7. Web user interface 8

2. Biofoundry design 11
    2.1. Overview of the biofoundry design 11
    2.2. Scheduler-driven biofoundry coordinated by Momentum 12
    2.3. Plate movement and standardized labware connect the instruments into a cell 12
    2.4. iBiofoundry and iBioFAB

3. iBioGenie 14
    3.1. iBioGenie synthetic biology action space 14
    3.2. iBioGenie agentic architecture 14
    3.3. Agentic coding framework and execution environment 15
    3.4. Semantic capability retrieval enables scalable access to heterogeneous scientific tools 16
    3.5. iBioGenie performs literature review through an automated retrieval pipeline 17
    3.6. iBioGenie generates protocols through evidence-grounded retrieval and composition 19
    3.7. Simulated user testing provides reproducible evaluation of autonomous scientific Workflows 21
    3.8. Automated tool addition expands the iBioGenie action space 24
    3.9. iBioGenie facilitates continuous capability integration and self-improvement 26

4. Benchmarking for iBioGenie 27
    4.1. iBioGenie benchmarking system configuration and baselines 27
    4.2. Multiple-choice QA benchmark (LAB-Bench QA Subset) 30
    4.3. Analytical benchmark (BixBench) 30
    4.4. Biomedical task benchmark (BiomniEval1) 31
    4.5. Construction of the internal wet-lab protocol generation benchmark 32
    4.6. Internal wet-lab protocol generation benchmark scoring and performance comparison 34

5. BiofoundryAI
    5.1. BiofoundryAI architecture and the biofoundry automation action space 37
    5.2. BiofoundryAI ensures reliable execution through layered validation and structured repair 37
    5.3. BiofoundryAI validates every workflow against a digital twin of the biofoundry 38
    5.4. BiofoundryAI generates instrument methods through deterministic and reasoning-based composition 40
    5.5. BiofoundryAI scales across compatible distributed biofoundries 43
    5.6. BiofoundryAI supports continuous improvement from operational experience 45

6. BiofoundryAI benchmarking 49
    6.1. Benchmark dataset construction 49
    6.2. Benchmark design 49
    6.3. Benchmark results 52

7. Case study I: Phytase engineering 55

7.1. Agentic AI pipeline for phytase engineering 55
7.2. Experimental and automation workflows for phytase engineering 62

8. Case study II: Integrated PAND discovery and engineering for enhanced 3-HP biosynthesis 65
8.1. Agentic AI pipeline for enzyme retrieval of aspartate decarboxylase 65
8.2. Experimental and automation workflows for new PAND discovery 73
8.3. Agentic AI pipeline for engineering the reterived aspartate decarboxylase 76
8.4. Experimental and automation workflows for PAND engineering 79

9. Case study III: Metabolic engineering of *Saccharomyces cerevisiae* for β-carotene production through combinatorial gene knockout 82
9.1. Agentic AI pipeline for gene knockout selection 82
9.2. Experimental and automation workflows for gene knockout selection 89

10. Conclusion on case studies 97
11. References 97

### 1. General information: iCloudBiofoundry application

#### 1.1. System overview

The iCloudBiofoundry application provides a unified interface connecting users, iBioGenie, and BiofoundryAI (**fig. S1**). Users submit research objectives through the chatbot interface, and the application routes these requests to iBioGenie, which interprets the user's objective, performs the required computational analyses, and generates experimental designs as well as wet-lab protocols. Intermediate results, explanations, and requests for clarification are returned to the application as agent responses, allowing users to review and refine the proposed strategy before experimental execution.

Once an experimental plan is approved, the computational design and protocol are transferred from iBioGenie to BiofoundryAI. BiofoundryAI converts the high-level protocol into a structured biofoundry workflow, including the required laboratory operations, instruments, parameters, and execution sequence. The application serves as the central orchestration and communication layer throughout this process. It manages interactions between the components, maintains the project state, presents execution progress, and collects the resulting experimental outputs. These outputs are organized into reports and returned to users through the same interface, creating a continuous workflow from a natural language research objective to computational design, laboratory execution, and interpretable experimental results.

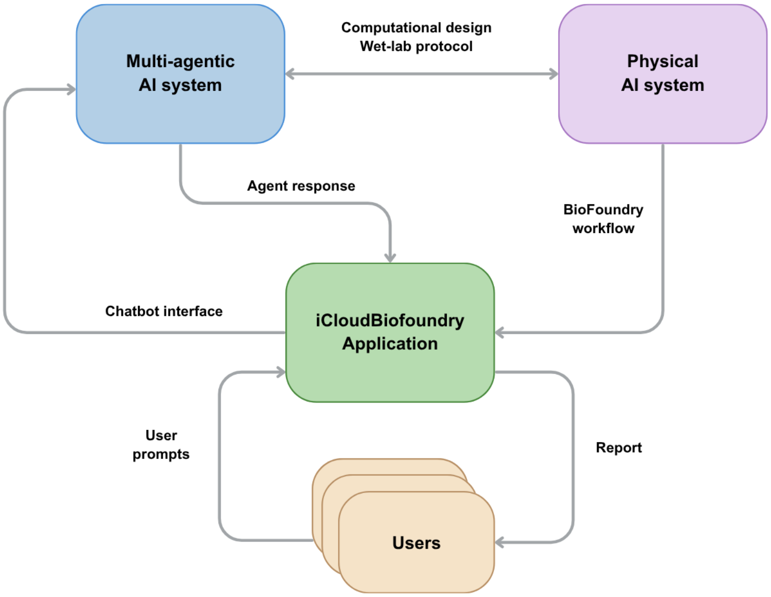


**Fig. S1. How information moves between the scientist, the reasoning system and the physical laboratory.** The platform has three parts, and the figure is the contract between them. The multi-agent AI system turns a natural language request into computational designs and wet-lab protocols. The physical AI system receives a protocol, translates it into an executable biofoundry workflow, and runs it. The iCloudBiofoundry application is the user's single point of contact with both systems. It relays user prompts to iBioGenie and streams the responses back through the chat interface; it receives the biofoundry workflow and the experimental results from BiofoundryAI and returns them to the user as a report.

#### 1.2. System architecture

iCloudBiofoundry is structured as a layered web application orchestrated by Kubernetes (**fig. S2**). The primary components are: (1) **Frontend**: The user interface is implemented as an Angular (https://angular.dev/) single-page application styled with Tailwind CSS (https://tailwindcss.com/) and PrimeNG (https://primeng.org/), communicating with the backend over Representational State Transfer (REST) and Server-Sent Events (SSE) for real-time updates. (2) **Backend:** A Python FastAPI (https://fastapi.tiangolo.com/) backend handles authentication, session management, job lifecycle operations, artifact storage, and proxied communication with iBioGenie and per-site Momentum instances. Real-time updates are delivered using FastAPI's built-in StreamingResponse, which streams SSE events over persistent per-user connections and pushes message, protocol, and job status updates to all active browser tabs. (3) **Data stores**: PostgreSQL (https://www.postgresql.org/) stores sessions, protocols, jobs, status history, and user records. MinIO (https://www.min.io/) provides Simple Storage Service (S3) compatible object storage for instrument output artifacts under artifacts/{session_id}/, with lazily refreshed 7-day presigned URLs and periodic backup and recovery. (4) **Message broker:** RabbitMQ (https://www.rabbitmq.com/) ensures real-time updates from all platform components are reliably delivered to the user interface, regardless of where in the system they originate. (5) **Ingress and TLS**: Traefik (https://traefik.io/) handles ingress routing and TLS termination for all incoming traffic. Traefik was chosen for its native Kubernetes integration and automatic TLS certificate management, ensuring all communication between clients and the platform is encrypted in transit without manual certificate configuration. (6) **Authentication**: All requests are authenticated before reaching the application. An OAuth2 Proxy (https://oauth2-proxy.github.io/oauth2-proxy/) intercepts unauthenticated traffic and redirects users through Keycloak (https://www.keycloak.org/) to CILogon (https://cilogon.org) for identity verification.

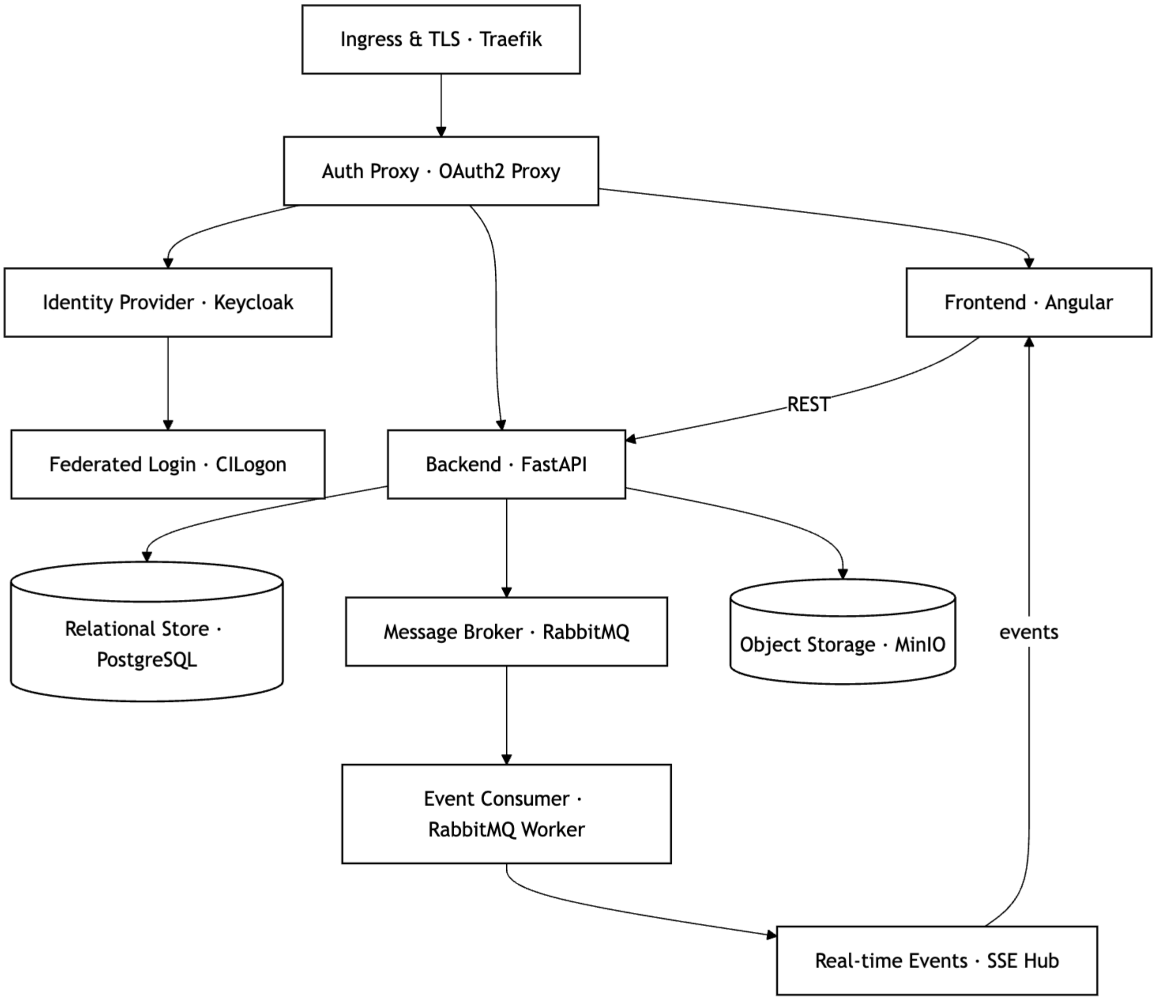


**Fig. S2. Architecture of the iCloudBiofoundry application.** Traefik terminates TLS and routes traffic to the OAuth2 Proxy, which authenticates users via Keycloak/CILogon before reaching the Angular frontend. FastAPI serves REST requests, persists to PostgreSQL and MinIO, and publishes job/status updates through RabbitMQ. A worker consumes these and streams them to the frontend via SSE.

#### 1.3. Federated identity and access management

Access is governed by CILogon, a federated identity broker supporting institutional single sign-on (SSO) via InCommon, Google, GitHub, and ORCID, allowing researchers to authenticate with home institution credentials rather than creating a separate account for iCloudBiofoundry use. Role-based access control on the application side separates researcher and administrator roles. Researchers manage sessions, submit jobs, and download artifacts. Administrators approve, reject, or cancel jobs and may attach notes or documents to any status update, ensuring that no instrument job reaches hardware without explicit human authorization.

#### 1.4. Deployment and operations

iCloudBiofoundry's components are containerized as separate Docker (https://www.docker.com/) images, ensuring environment consistency and reproducibility across development and production. These images are orchestrated with Kubernetes (https://kubernetes.io/) on an Ubuntu (https://ubuntu.com/) cluster hosted on National Center for Supercomputing Applications (NCSA)'s OpenStack on-premises cloud, providing automated scaling, self-healing, and rolling updates with minimal downtime. The full stack is packaged as a single Helm chart (https://helm.sh/) for one-command deployment and straightforward onboarding of new biofoundry sites. Component health is continuously monitored via Uptime Kuma (https://uptime.kuma.pet), an open-source monitoring dashboard that tracks service availability and delivers real-time alerts to a dedicated Slack channel for the development team. Operational data are backed up daily to NCSA's Taiga, a petabyte-scale NFS storage resource for research data.

Computational hardware is hosted on the Radiant OpenStack (https://www.openstack.org/) system at the NCSA (https://docs.ncsa.illinois.edu/systems/radiant/en/latest/user-guide/architecture.html). Compute is a two-node Slurm cluster running Ubuntu 24.04 LTS: the primary node (ibio-gpu-01) serves as the Slurm controller and hosts the containerized chat, agent, and simulation services on 3 NVIDIA Tesla V100 GPUs (32 GB each), 24 Intel Xeon virtual CPU cores, and 96 GB RAM, while a second node (ibio-gpu-02), configured via slurm, provides 1 NVIDIA A100 GPU (80 GB), 24 AMD EPYC virtual CPU cores, and 240 GB RAM. Persistent storage for the iCloudBiofoundry system is a 50 TB NFS export with a maximum throughput of 50 GB/s, hosted via the Taiga system (https://docs.ncsa.illinois.edu/systems/taiga/en/latest/index.html) at NCSA and mounted on both nodes for model weights, reference databases, session state, and experiment outputs.

#### 1.5. Cloud access and scalability

iCloudBiofoundry provides cloud-based access to iBioGenie through an interactive web interface, allowing researchers to use its scientific capabilities without directly interacting with the underlying computational tools. Within the same environment, users can define biological objectives, explore design strategies, perform computational analyses, and generate wet-lab protocols. Programmatic access is also available through an API, providing a common entry point for both interactive use and integration into external workflows.

For experiments requiring automated execution, iBioGenie can interact with BiofoundryAI to translate a proposed wet-lab protocol into an executable biofoundry workflow. Users can submit protocols for automation, inspect the resulting workflow plan, review and approve executable scripts, and monitor experimental execution remotely. Approved workflows can then be routed to a compatible biofoundry connected to the iCloudBiofoundry platform. The same cloud interface supports computational design, protocol development, automation planning, and physical experiment execution across distributed facilities, while allowing additional biofoundry sites and experimental capabilities to be incorporated as the platform expands.

The cloud architecture also allows computational and experimental resources to be shared across multiple users. Load testing with simulated users (**fig. S3**) showed that the current chat service sustained 3,200 concurrent users on two CPU cores, while the web backend supported 250 concurrent users without queuing. The main computational constraint is GPU-dependent iBioGenie design, for which four GPUs are available through the Slurm queue and therefore support up to four concurrent GPU-intensive design jobs. This capacity can be increased by adding computational resources or adjusting cluster scheduling.

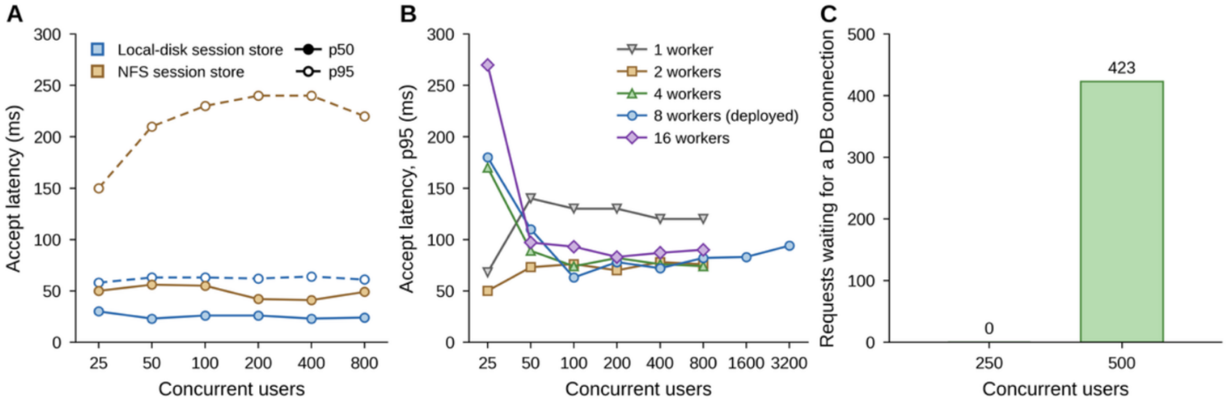


**Fig. S3. Concurrent user capacity of the iCloudBiofoundry platform.** Request-serving capacity was evaluated with Locust. For chat-service tests, model inference was replaced by a 27-s delay, matching the median duration of 2,317 production turns. The y-axis shows server acceptance delay as median (p50) and 95th percentile (p95) **(A)** iBioGenie on 40 CPU cores with 20 workers, latency remained flat from 25 to 800 concurrent users with no failures. Shared NFS storage added ~25 ms per turn relative to local disk. **(B)** The same sweep restricted to 2 CPU cores at 1, 2, 4, 8 and 16 worker processes. Every configuration holds p95 near or below 100 ms to 800 users, and the 8-worker production configuration holds it to 3,200 users. **(C)** The web backend and PostgreSQL database showed no connection waiting at 250 users, whereas 423 requests were queued at 500 users, placing the database-limited capacity between 250 and 500 concurrent users. Tests were performed in June–July 2026.

#### 1.6. Data management and privacy

Data handling follows four principles: everything a user generates is stored, every item is tied to a session, data is accessible only to authorized users, and all persisted data is backed up on a defined schedule. All user-generated content is persisted in the backend rather than maintained only as a transient session state. This includes designs, chat history, protocols, job definitions, execution status history, messages, and raw instrument outputs. When a client reconnects to an existing session, the complete prior state is restored from persistent storage. Structured data lives in PostgreSQL (https://www.postgresql.org/): sessions, protocols, jobs, status history, and user records. Binary instrument outputs are stored as objects in MinIO (https://www.min.io/) linked by session id and persist until the project is deleted. Clients never access MinIO directly; they receive 7-day presigned URLs that the backend re-issues on each access, so links expire but data does not.

Mirrored session data for iBioGenie accessibility in NFS file storage. This includes session logs not normally exposed to the user, and hidden LLM output files emitted during the generation process but are available from request in chat. Frontend forking of sessions creates symbolic links between session directories to keep context linked.

Access control is enforced at the session level. Every record and stored artifact is associated with the session in which it was created, and all authorization and data access are handled through the backend to ensure that users can access only the resources permitted within that session. A session's data is readable only by its owning user and by platform administrators (for support and maintenance). Since all traffic is authenticated before reaching the application, no unauthenticated client can read or download stored data.

As externally (non-locally) deployed LLMs are leveraged by iCloudBiofoundry agents, data sovereignty policies are specific to the models/providers selected by users (*e.g.* OpenAI). Internally managed user prompt data may be used to improve iCloudBiofoundry systems in the future. All scientific/research data included in prompts is kept confidential within iCloudBiofoundry systems to the largest extent possible, as is described above, however we cannot make guarantees due to the deployment and usage of third-party models.

An overview of the computational tools, specialized models, scientific databases, and computing infrastructure that can be accessed by the platform to support biological research workflows can be found at https://icloudbiofoundry.illinois.edu/resources.

**1.7. Web user interface**

The web user interface is organized into a public-facing informational site and an authenticated application. The public site (https://icloudbiofoundry.illinois.edu/) features resources related to the iCloudBiofoundry platform, such as tutorial content, an interactive listing of integrated resources, and related research publications (**fig. S4**).

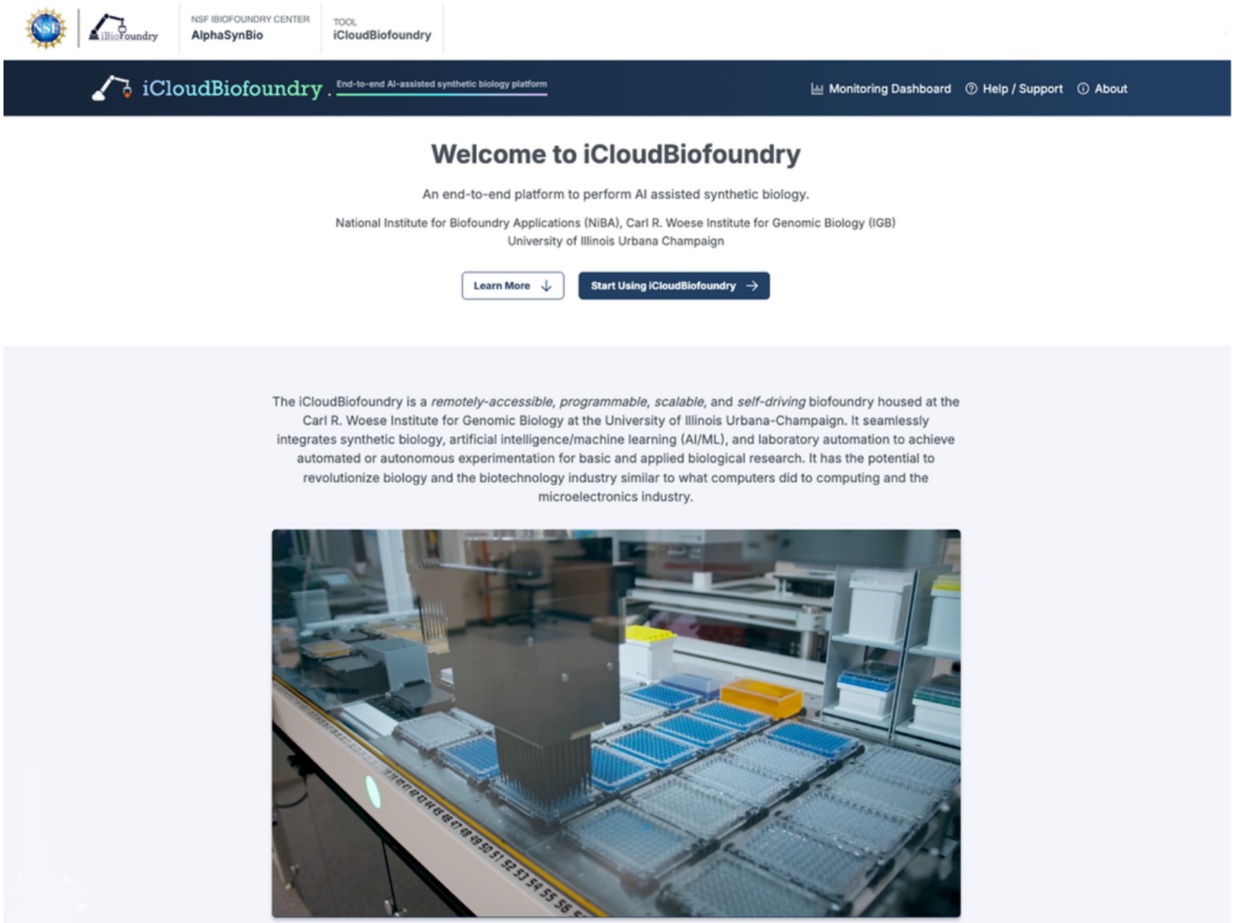


**Fig. S4. The iCloudBiofoundry landing page.** The pre-authentication entry point of the platform. The “Start Using iCloudBiofoundry” button launches the application.

**Design Projects:** After creating an account and logging into the authenticated application, the user begins by creating a new project. Each project corresponds to a chat and, optionally, a biofoundry experiment. Using natural language, the user can engage with the iBioGenie multi-agent system through the chat, entering prompts to complete a range of synthetic biology tasks, from hypothesis generation to in-silico design, to generating wetlab protocols. If the user chooses to generate a biofoundry protocol, which is compatible specifically with autonomous execution in one of the iCloudBiofoundry facilities, BiofoundryAI is called to generate and return a biofoundry protocol to the user (**fig. S5**).

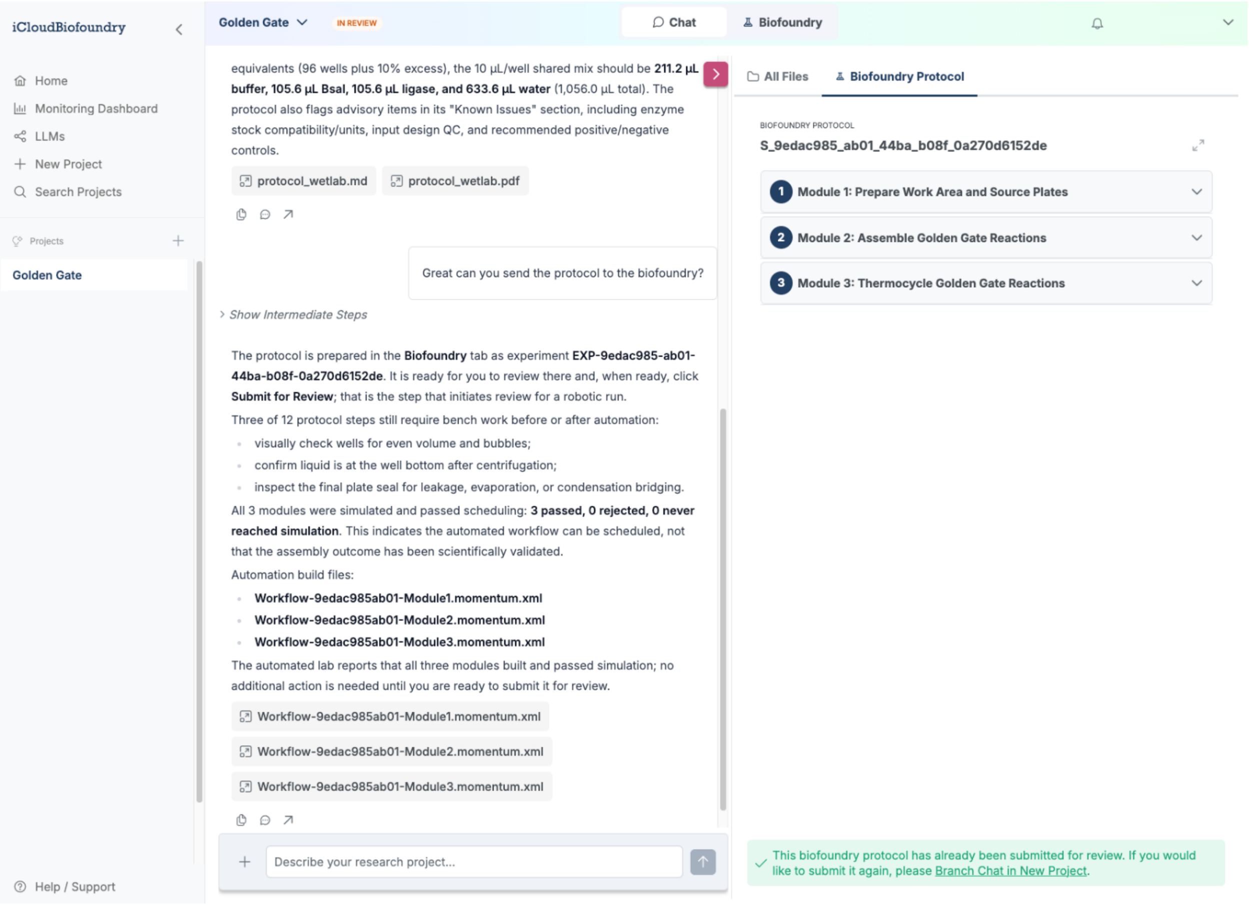

**Fig. S5. iCloudBiofoundry chat.** The chat interface combines chat with a collapsible files panel for viewing generated or uploaded files, including structures, protocols, and reference datasets. Biofoundry protocols can also be submitted for staff review through the Biofoundry protocol tab.

**Explore Results:** When experimental results become available for a given experiment, the corresponding biofoundry experiment page is updated to provide access to these results. Raw result data can be imported into chat for AI-assisted analysis with the multi-agent system, building on the existing project context. Alternatively, the user may download the results to use in their own downstream workflows. The web user interface also includes a feature for branching existing projects into new projects to support iterative experimentation (**fig. S6**).

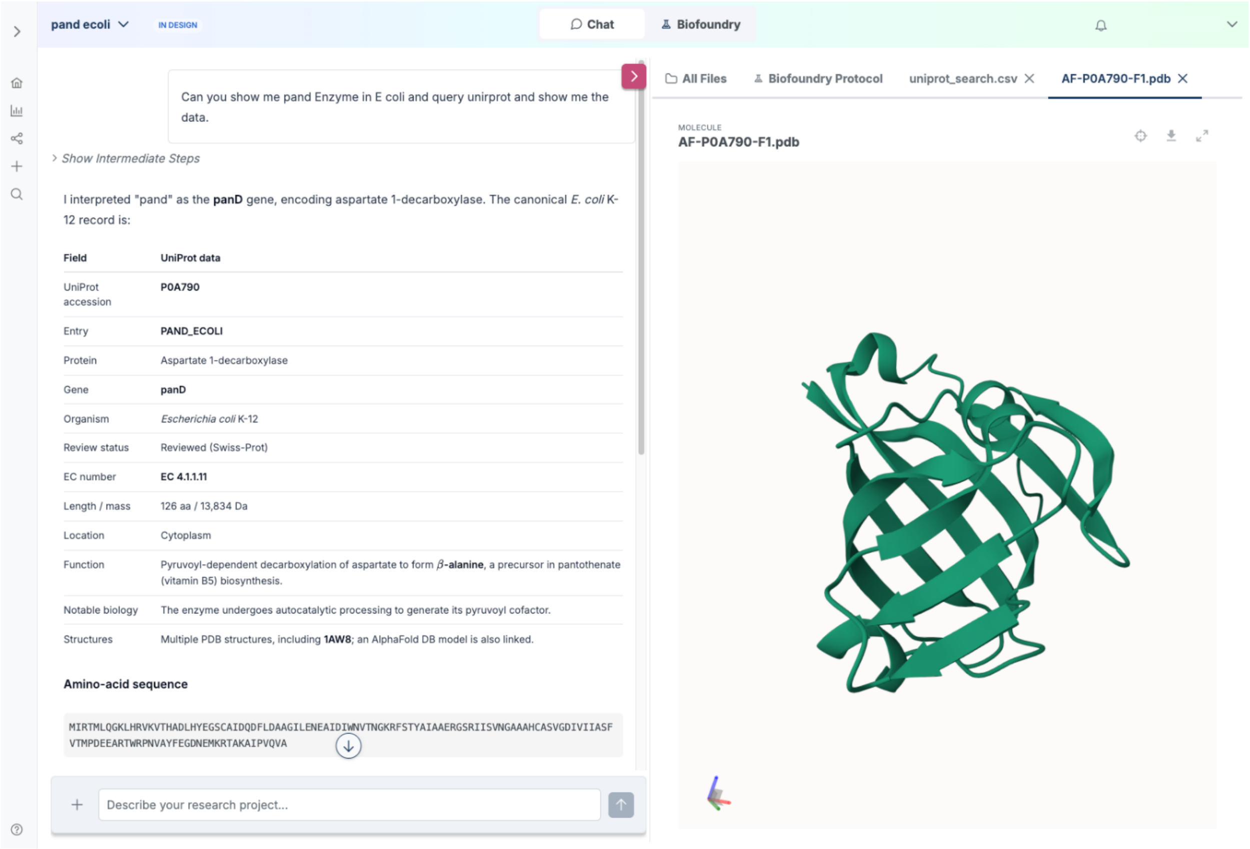


**Fig. S6. iCloudBiofoundry file viewer.** The iCloudBiofoundry platform features an in-app viewer with support for multiple file types, enabling users to explore results data, protein structures, and more within the project chat and biofoundry experiment page. The integrated viewing of a CSV file (left) and PDB file (right) are pictured above.

**Monitor Execution:** After generating a biofoundry protocol, the user can directly submit their protocol to iCloudBiofoundry staff for review. Once submitted, users can track the status of their protocol through review, approval, and execution (**fig. S7**).

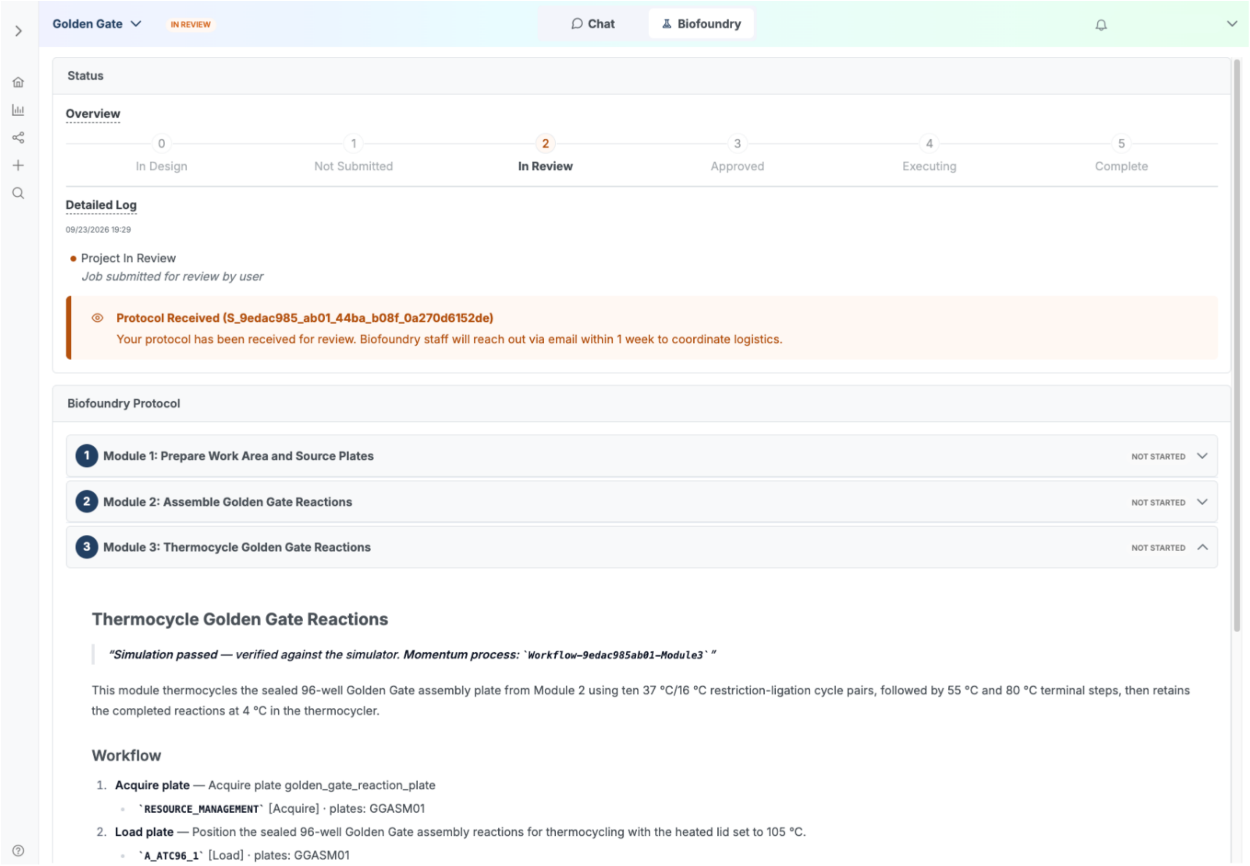


**Fig. S7. iCloudBiofoundry experiment page.** The biofoundry experiment page enables users to track the status of their biofoundry protocol and experiment, including a high-level stepper visualization and a detailed log of updates. Updates include both system alerts and messages or attachments from biofoundry staff.

### 2. Biofoundry design

#### 2.1. Overview of the biofoundry design

iCloudBiofoundry executes validated workflows on two integrated robotic biofoundries at the University of Illinois Urbana-Champaign (UIUC): iBioFoundry and iBioFAB. Each workcell combines specialized instruments for liquid handling, thermal cycling, plate reading, sealing and peeling, centrifugation, incubation, colony picking, and acoustic dispensing. The microplate is the unit of work, and a workflow is represented as an ordered sequence of plate operations. Scheduling software-controlled movement between devices preserves plate identity, timing, positional compatibility, and ensures safe transport.

Both facilities use the same scheduler Momentum (Thermo Fisher Scientific) coordinates instrument access and workflow execution (**Section 2.2**), while robotic arms, swap stations, plate hotels, and standardized labware connect the instruments physically (**Section 2.3**). The architecture is implemented differently at each site: iBioFoundry is a larger modular system with three transport zones, whereas iBioFAB is a compact workcell with one central arm and a different instrument complement (**Section 2.4**).

#### 2.2. Scheduler-driven biofoundry coordinated by Momentum

Momentum coordinates each biofoundry as an integrated workcell rather than a collection of independently automated instruments. A workflow specifies the required device operations, their dependencies, and associated labware. Momentum converts this workflow into a timed schedule that accounts for instrument availability and plate movement; the scientist defines what must occur, while the scheduler determines when each operation runs and directs transport between instruments.

Momentum performs four functions. It maintains the workcell inventory by registering each plate's container type and position; schedules instrument access based on workflow dependencies and device availability; directs robotic transport and enforces safety interlocks; and simulates workflows before hardware use. During simulation, Momentum advances the work queue, device methods, plate movements, and timing against a software representation of the workcell. This process reveals scheduling conflicts, incompatible transfers, and missing resources before samples are handled. Only workflows that complete the check proceed to hardware execution. The resulting simulated state serves as the operational digital twin in this study but does not model every physical or biological process.

#### 2.3. Plate movement and standardized labware connect the instruments into a cell

Robotic arms connect the device zones. Each arm services part of the workcell, and swap stations transfer plates between zones when a destination lies beyond one arm's reach. Plate hotels hold labware between operations. Refrigerated or temperature-controlled units are used when required, and lid parks temporarily hold covers while a plate is processed.

Shared robotic arm movement does not imply a single plate design. The supported labware includes flat-bottom 96- and 384-well assay plates, skirted plates for thermal cycling, deep-well plates for culture and storage, low dead volume 384-well source plates for acoustic dispensing, reagent reservoirs, and tip racks. Lids and seals are used when an operation requires a covered plate.

Each container template records the plate dimensions and device compatibility. Momentum accepts a container at a position only when the template is registered for that location. Workflow definitions therefore carry the container type and intended positions through the workcell and check those assignments before execution. Barcodes preserve plate identity as labware moves through the cell.

#### 2.4. iBioFoundry and iBioFAB

iBioFoundry is organized into three connected zones, each served by its own robotic arm, with the scheduler moving plates between zones through swap stations. It contains 60 devices, 47 physical instruments and 13 digital utilities. A subset are the physical instruments described below, and the remainder are storage hotels, robotic arms, swap and lid-handling stations, and control and data-management utilities. Selected representative devices in the iBioFoundry instrument set are labeled in **fig. S8**. Liquid handling is provided by a Tecan Fluent, Thermo Fisher Multidrop Combi dispensers, a BioTek MultiFlo FX dispenser and washer, and a Labcyte Echo acoustic dispenser. A SCIEX Echo MS couples acoustic ejection to mass spectrometry. Thermal cycling uses an Applied Biosystems QuantStudio 7 Pro real-time PCR system and automated 96- and 384-well cyclers. Detection uses a Tecan Spark reader, an Unchained Labs Lunatic spectrophotometer, and a fragment analyzer. The workcell also includes a Singer Instruments PIXL colony picker, a magnetic-bead processor, plate sealers and peelers, a microplate centrifuge, shakers, Cytomat incubators and storage, plate hotels, and a Scinomix SciPrint MP2 labeler. Spinnaker robots move plates between these devices through the zone swap stations.

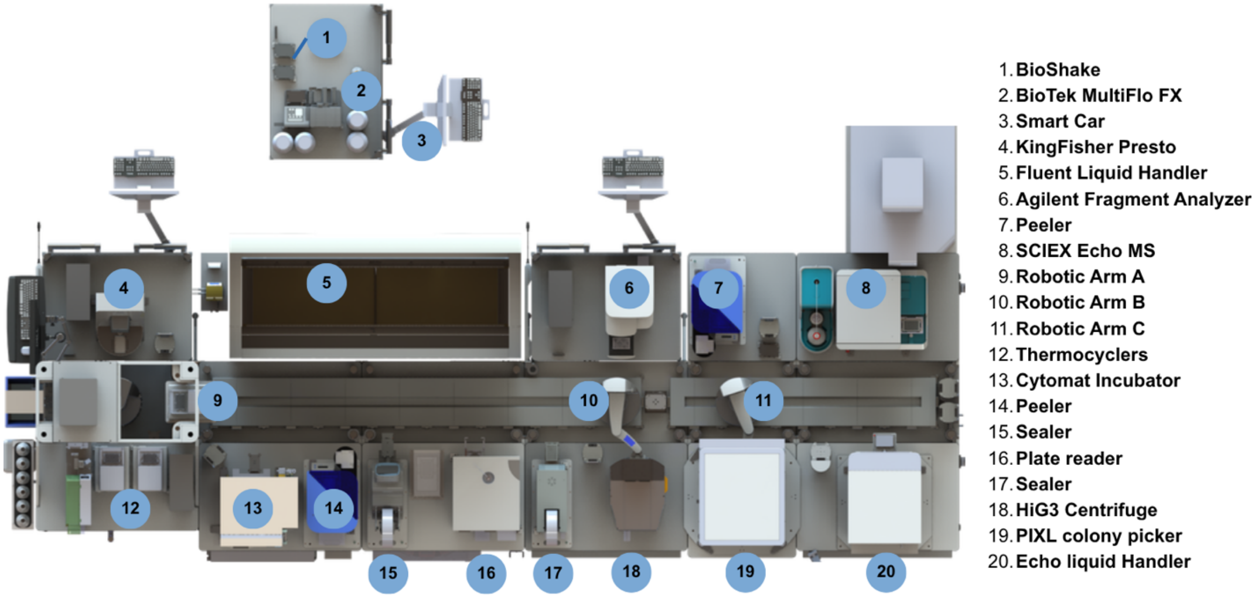


**Fig. S8. Overview of the iBiofoundry automation platform.** Top-down view with selected devices labeled.

The iBioFAB is an integrated workcell with one central arm. There are a total of 48 devices at iBioFAB, 33 physical and 15 digital. Selected devices from this instrument set are labeled in **fig. S9**. The instrument set includes a Tecan Fluent and four Thermo Fisher Multidrop Combi dispensers for liquid handling; a Labcyte Echo for nanoliter transfer; two Analytik Jena Biometra TRobot thermocyclers; BioTek Cytation and Tecan Infinite readers; a fragment analyzer; and an Agilent microplate centrifuge. Plate handling is supported by ALPS3000 and 4titude sealers, an XPeel, Cytomat incubators and storage units, chilled blocks, plate hotels, labeling equipment, regrip devices, and a central swap station. Momentum coordinates these devices through the operator layer.

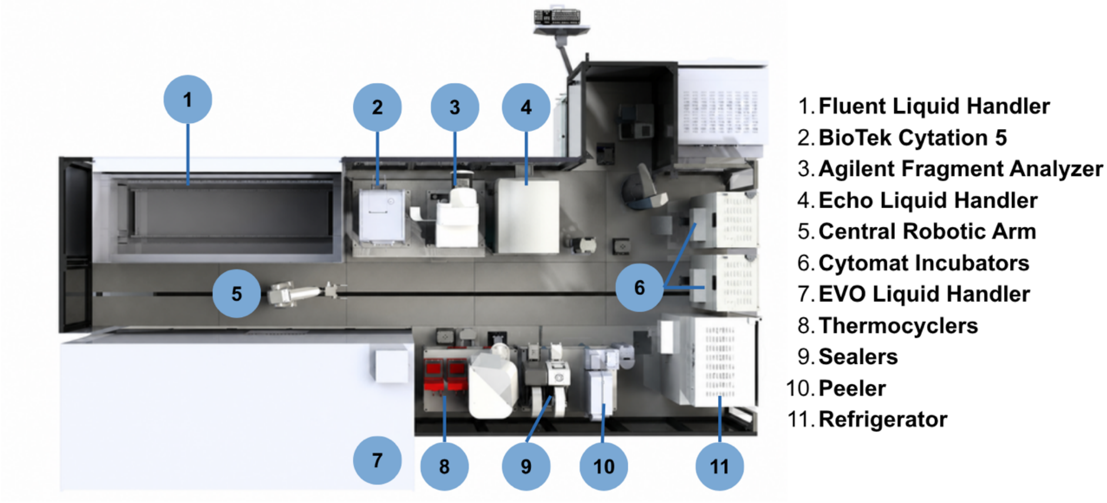


**Fig. S9. Overview of the iBioFAB automation platform.** The rendering labels the principal instruments and the central transport system.

Together, the two facilities provide the hardware and control context used by BiofoundryAI. A generated workflow is bound to site-specific devices and labware and run in the Momentum simulation before deployment. This sequence supports closed-loop experiments while keeping device scheduling and plate transport under deterministic control.

### 3. iBioGenie

#### 3.1. iBioGenie synthetic biology action space

iBioGenie is the scientific intelligence layer of iCloudBiofoundry, designed to convert a user's natural-language biological objective into a structured research process spanning computational design, biological data retrieval, code execution, literature synthesis, wet-lab protocol generation, and downstream handoff to robotic execution. Because synthetic biology research rarely consists of a single isolated task, iBioGenie is built around a broad action space that provides the capabilities required to address diverse, multi-stage research objectives. The current action space includes 293 integrated tools and databases, a Python sandbox with access to more than 193 pre-installed scientific packages, 30 CLI binaries, and a library of 627 procedural skills. The skill library combines skills developed in-house to support common laboratory work, experimental design, and biological analysis tasks with selected skills relevant to synthetic biology and related life-science workflows from publicly available collections, including bioSkills (https://github.com/GPTomics/bioSkills), SciAgent-Skills (https://github.com/jaechang-hits/SciAgent-Skills), and K-Dense scientific-agent-skills (https://github.com/K-Dense-AI/scientific-agent-skills). These components represent complementary forms of capability. Tools and databases provide executable access to specialized computational methods and biological resources. The Python environment supports flexible analysis, data processing, visualization, and custom computation. Skills provide reusable procedural knowledge that can be incorporated directly into the agent's reasoning when a task is better represented through scientific instructions than through a dedicated executable tool.

Integrated tools and databases are used for capabilities that require explicit execution logic, structured inputs and outputs, access to external software or biological resources, specialized computing environments, or deterministic control over how an operation is performed. Each integration is represented through a standardized interface that includes a functional description, supported inputs and outputs, usage constraints, and instructions governing when the capability is appropriate for a given scientific task. By abstracting implementation-specific details behind structured interfaces, iBioGenie can incorporate diverse computational resources into a common action space while preserving the operational requirements and scientific context needed for reliable execution. These integrations provide the executable backbone of the platform, supporting tasks ranging from biological information retrieval and sequence or structural analysis to model-based prediction, data processing, and more complex computational workflows.

Skills form a separate component of the action space. A skill is a self-contained natural-language specification of a scientific procedure, methodology, analytical convention, or best practice that can be retrieved and added directly to the agent's working context. Relevant skills are retrieved according to the current scientific objective and incorporated into the agent's context only when needed. This allows iBioGenie to access a large body of procedural knowledge without continuously loading the complete library and reduces reliance on the language model's parametric knowledge for specialized scientific procedures. The curated skill library spans genomics, transcriptomics, structural biology, cheminformatics, systems biology, statistical analysis, scientific reporting, and general research practices from multiple resources, with additional in-house skills developed specifically for synthetic biology workflows.

#### 3.2. iBioGenie agentic architecture

While the action space defines what iBioGenie can do, the agentic architecture determines how these capabilities are selected, combined, executed, and interpreted to accomplish a scientific objective. Rather than following a predefined workflow, iBioGenie dynamically assembles the appropriate sequence of actions based on the biological problem and intermediate results produced during execution.

iBioGenie's orchestrator agent coordinates the overall workflow. It interprets the user's intent, identifies the biological context, decomposes the objective into research tasks, selects relevant capabilities from the action space, coordinates their execution, tracks intermediate results, and synthesizes them into scientific conclusions or experimental plans. This enables iBioGenie to operate at the level of research strategy, determining which capabilities are required, the order in which they should be applied, how their outputs should be interpreted, and whether additional evidence or computation is needed before proceeding.

The orchestrator is supported by three agentic modules and a set of single-purpose LLM roles. A code-execution module handles custom Python analyses, specialized libraries, data processing, visualization, and computational routines not covered by predefined tools (**Section 2.3**); a literature-search module collects evidence from scientific publications (**Section 2.6**); and a protocol-generation module translates scientific conclusions and experimental objectives into experimentally grounded procedures (**Section 2.7**). Each module combines deterministic code with one or more narrowly scoped LLM calls, and only the code-execution module runs an autonomous tool-use loop of its own. A separate single-pass reporter consolidates results, generated artifacts, and citations into a final research report.

Scientific projects often unfold over many stages, with files, computational results, and experimental decisions accumulating along the way. To maintain continuity across these stages, iBioGenie maintains a persistent session state that tracks generated files, tool outputs, key findings, completed and pending tasks, and long-running jobs. The architecture also supports asynchronous, high-performance computation. Computationally intensive tasks are submitted as independent SLURM jobs while the agent continues coordinating the broader workflow. Once completed, their outputs are registered back into the session state and incorporated into subsequent reasoning, enabling iBioGenie to coordinate analyses that require specialized software environments, GPU resources, or execution times that extend beyond a single interactive exchange.

Overall, the architecture connects computational reasoning with experimental action. Depending on the task, iBioGenie can produce biological designs, analysis scripts, figures, experimental recommendations, reports, and wet-lab protocols, rather than being limited to natural-language responses. Within the full iCloudBiofoundry system, these protocols serve as the interface between iBioGenie and BiofoundryAI, with iBioGenie organizing the scientific reasoning and experimental intent, while BiofoundryAI translates the resulting protocol-level plan into validated workflows for robotic execution.

#### 3.3. Agentic coding framework and execution environment

iBioGenie includes a dedicated coding agentic framework (developer module) for tasks that require computation, data analysis, or programmatic manipulation of the user's data and provided information. Rather than having the orchestrator agent write and execute code directly, computational tasks are delegated to a separate coding workflow. Before this happens, a routing layer, a cascade of deterministic checks backed by a lightweight LLM classifier, determines whether code execution is needed, whether an existing scientific tool is a better fit, or whether the orchestrator should answer based on its own reasoning. Once a task is assigned to the coding framework, the original user question is preserved and passed to the developer agents so that quantitative requirements, thresholds, comparison operators, and other details are not lost through intermediate paraphrasing (**fig. S10A**).

The coding workflow separates planning from execution. A planning agent first defines the computational objective, identifies the required inputs and analysis method, and breaks the task into a sequence of verifiable steps. It also considers alternative approaches if the primary analysis fails. This plan is maintained throughout execution and repeatedly provided to the coding agent, which writes and runs the code needed to complete each step. The developer agent can revise its implementation when an execution fails, update the plan as work progresses, and stop once the requested result has been successfully produced. This separation allows iBioGenie to approach computational tasks as structured analyses rather than relying on a single code-generation response.

Generated code is executed in an isolated coding environment rather than inside the main iBioGenie process. Each execution attempt runs in a fresh subprocess with a new Python namespace, while the agent's conversation history, computational plan, saved files, and checkpoints can persist across attempts. The environment provides common scientific Python libraries and dedicated locations for input data and generated outputs. For analyses that require established scientific software rather than Python implementations, the coding agent can also call a controlled set of command-line programs through the same environment. These include commonly used tools for sequence analysis, alignment, phylogenetics, genomics, and statistical analysis, as well as R and Rscript. The execution environment applies wall-clock limits, restricts accessible environment variables, controls in-process network access, and limits external program invocation to an explicit binary allowlist.

Execution is iterative and includes explicit recovery from common failures. When code raises an exception, produces no useful output, exceeds its execution time, or refers to missing files, the failure is returned to the coding agent with information to revise the next attempt. Successful intermediate results can be saved as files or checkpoints so that later attempts do not need to repeat completed work. The system also checks for cases in which an apparently successful computation is actually a substitute for the requested analysis, such as returning a simplified proxy after the intended scientific method failed. Such outputs are rejected before they are recorded as successful results.

Finally, a completed computational result is checked against the user's original question before it is returned to the main iBioGenie workflow. Deterministic checks are applied first, followed, when appropriate, by a separate fidelity critic agent that evaluates whether the result answers the requested quantity, comparison, and scope. If a problem is identified, the coding agent is given one opportunity to revise the analysis. Outputs and files that pass these checks are returned to the orchestrator and registered as part of the project state, allowing results produced through code execution to be used by subsequent iBioGenie reasoning, analysis, and experimental-design steps.

#### 3.4. Semantic capability retrieval enables scalable access to heterogeneous scientific tools

One of the main challenges for autonomous scientific agents is the scale and diversity of the available action space. iBioGenie integrates a large and growing collection of tools and skills, but exposing every capability and its full input requirements to the agent at every turn would be inefficient and unreliable. It would consume substantial context, increase computational cost, and distract the agent with many irrelevant options. To address this, iBioGenie uses an on-demand capability-retrieval layer that allows the agent to discover, inspect, and invoke tools only when they are relevant to the current scientific task. Each capability is represented by a structured description containing both semantic information such as what it does, the inputs it accepts, the outputs it produces, and when it should or should not be used, and operational metadata, such as availability, computational requirements, data dependencies, and readiness for execution.

This retrieval layer acts as a compact interface between the reasoning model and the broader capability ecosystem. Rather than exposing the complete action space, iBioGenie retrieves a ranked set of candidate tools and skills based on the scientific intent of the current task. The agent can then inspect the detailed requirements of the selected capability before execution, reducing context burden while preserving access to a broad and continuously extensible tool library. Retrieval is supported by expert-curated descriptions of use cases, inputs, outputs, limitations, and negative usage guidance, which are used to build a semantic index over the available capabilities. This allows conceptually related requests to retrieve appropriate tools even when the user's terminology differs from the implementation name, while also helping distinguish between closely related but scientifically different tasks.

The action space is also dynamically controllable and independently evaluated. Capabilities can remain implemented and testable while being hidden from the active agent if they are experimental, unavailable, superseded, or temporarily unreliable, allowing new tools to be staged, replaced, or disabled without redesigning the overall system. This enables iBioGenie to evolve as new biological databases, predictive models, computational methods, and laboratory interfaces become available. Because retrieval quality directly determines whether these capabilities are practically accessible, iBioGenie evaluates capability discovery separately from downstream execution by testing whether the correct tool or skill can be retrieved from natural-language task descriptions without requiring exact capability names (**fig. S10B-D**).

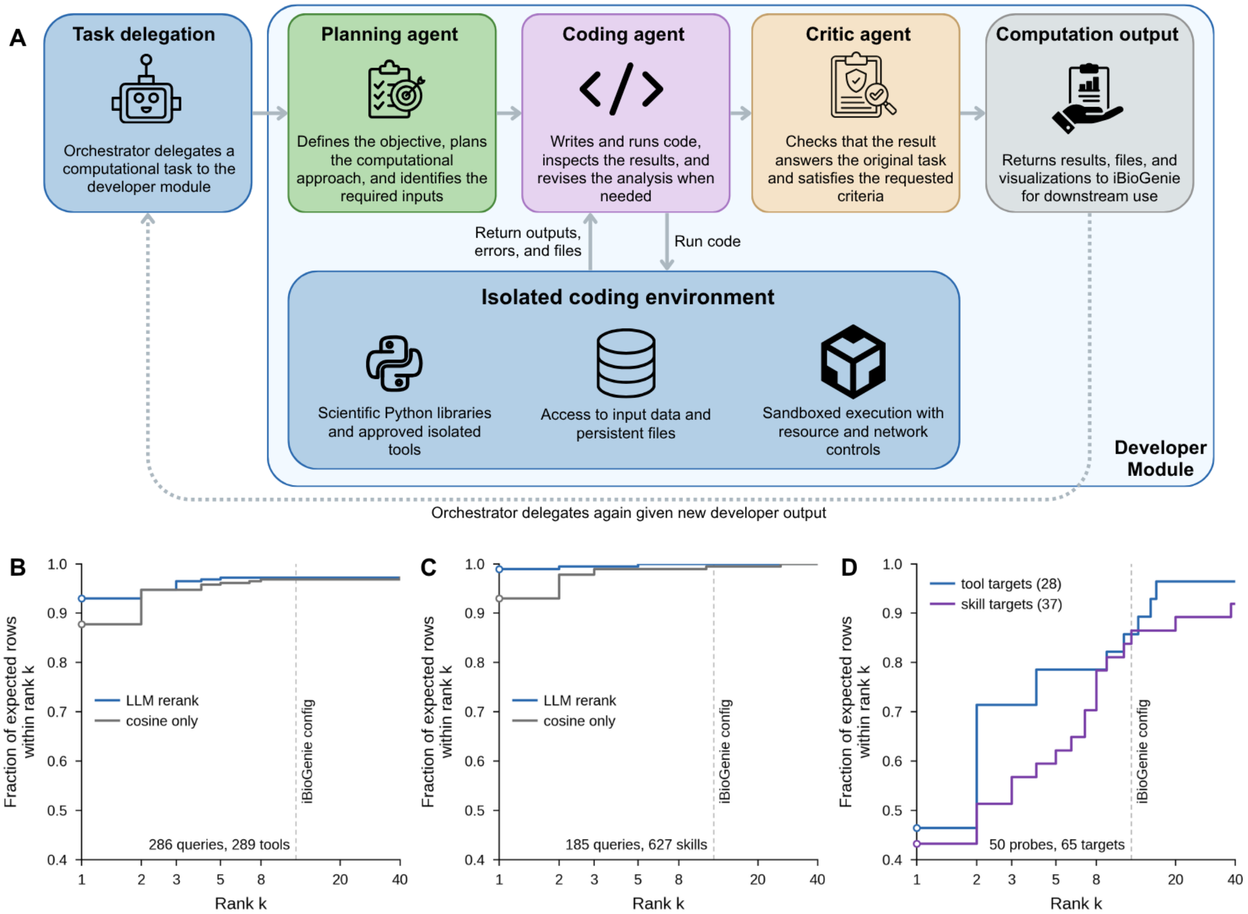


**Fig. S10. Developer module and semantic capability retrieval. (A)** Delegation architecture. The orchestrator hands a task to the developer module, where a planning agent sets the approach, a coding agent writes and runs code in a sandboxed scientific Python environment, and a critic agent checks the result against the task. Failures return to the coding agent for revision; finished results return to the orchestrator. **(B to D)** Retrieval recall as the fraction of expected items retrieved within rank k; the dashed line marks the 12-row window shown to the model. **(B)** Tool retrieval (286 queries, 289 tools) and **(C)** skill retrieval (185 queries, 627 skill cards) place the expected item within the window for nearly every query, with reranking lifting rank-1 accuracy to 93.0% and 98.9%. **(D)** When tools and skills are ranked together (50 probes, 65 targets), rank-1 accuracy drops below 50%, but roughly 86% of targets still land within the window, showing similarity search plus reranking creates a usable catalog.

#### 3.5. iBioGenie performs literature review through an automated retrieval pipeline

iBioGenie includes an automated literature review module to incorporate relevant scientific evidence from published work that can be activated whenever additional evidence is needed to support reasoning, workflow design, or experimental planning. Given a literature search task, the system first expands the original request into related academic search terms, allowing the literature agent to capture alternative terminology, synonyms, and closely related concepts that may appear across different subfields. The original and expanded queries are then searched across multiple scholarly sources, including Europe PMC (*1*), Semantic Scholar (*2*), and arXiv, enabling the retrieval of peer-reviewed articles and literature relevant to the tasked objective. Retrieved records are parsed to extract titles, abstracts, authors, identifiers, publication information, and open-access status, after which duplicate papers are removed using DOI and title-based matching. The remaining papers are semantically reranked against the original query, allowing iBioGenie to prioritize studies that are most relevant to the user's scientific question rather than relying only on keyword overlap.

After identifying the most relevant papers, iBioGenie attempts to acquire full-text articles when open-access versions are available. The system uses open-access discovery services and fallback strategies to locate downloadable PDFs. Downloaded articles are stored in a persistent library, together with structured metadata and citation keys, allowing the literature base to be reused across future sessions and projects. To make full papers searchable by the agent, PDFs are converted into machine-readable text through an Optical Character Recognition (OCR) and document-conversion pipeline (*3*). The extracted text is then segmented and embedded into a semantic index, enabling iBioGenie to search across the full content of papers and retrieve specific passages relevant to the current task. This allows the agent to move beyond abstract-level retrieval and use detailed evidence from full-text articles during scientific reasoning.

The literature review system was designed to be robust, and reusable across projects. Long-running operations such as PDF download, OCR, and semantic index construction are executed asynchronously on the computing infrastructure, while metadata, download results, OCR status, and index state are persisted to maintain continuity across sessions. The system also includes safeguards against redundant searches, duplicate library entries, failed downloads, and incomplete OCR outputs. In addition to structured literature retrieval, iBioGenie can access real-time web search tools when information is not yet captured in the scientific literature or when broader contextual information is required.

iBioGenie also allows users to bring their own scientific context into the workflow through connected literature libraries and direct file uploads. By uploading their files or connecting a Zotero library, the system can use a researcher's curated papers, downloaded PDFs, and metadata to further provide more project-specific context and support tasks such as protocol generation (**fig. S11**).

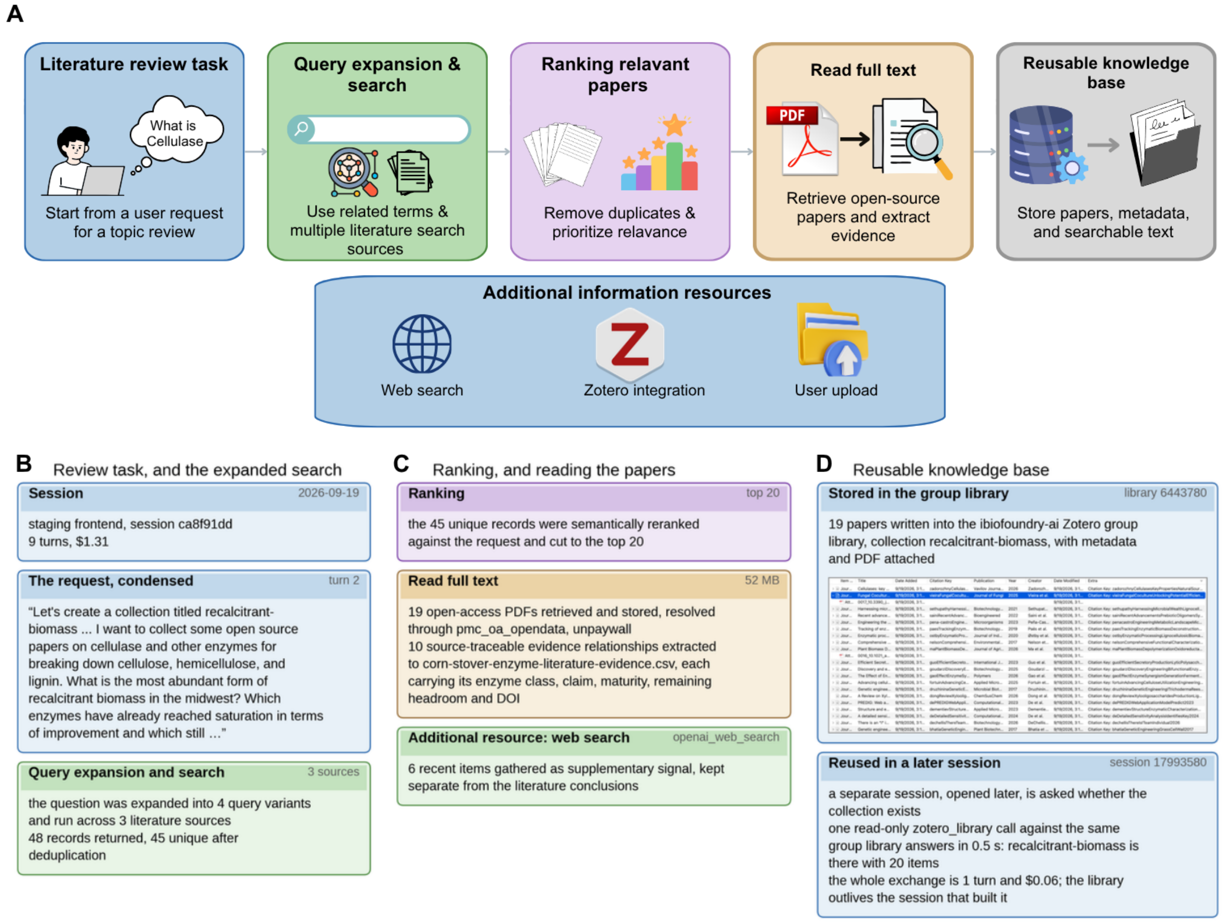


**Fig. S11. Literature review with an example session on cellulase. (A)** Literature review pipeline, including query expansion, multi-source search, deduplication, ranking, full-text retrieval, evidence extraction, and storage in a reusable knowledge base. Web search, Zotero connection, and user uploads are all available for the system. **(B)** The review task and expanded search. One request became 4 query variants across 3 sources, returning 48 records, 45 after deduplication. **(C)** Ranking and reading. The 45 records were reranked against the request and cut to the top 20; 19 open-access PDFs were retrieved via PubMed Central and Unpaywall, and 10 source-traceable evidence relationships were extracted to a table with enzyme class, claim, engineering maturity, remaining headroom, and DOI. A separate web scan contributed 6 recent items. **(D)** The reusable knowledge base. The 19 papers were written to the Zotero group library with metadata and PDFs. A later, unrelated session confirmed the collection in one read-only call.

#### 3.6. iBioGenie generates protocols through evidence-grounded retrieval and composition

iBioGenie generates wet-lab protocols through a multi-stage workflow that combines evidence retrieval, protocol generation, and post-generation verification. The process begins by breaking the requested experiment into individual procedural modules and arranging them in the appropriate experimental sequence. A separate search query is generated for each module, allowing evidence to be retrieved independently for each stage of the workflow. This stage-specific retrieval helps prevent well-documented procedures from dominating the available context and makes it easier to distinguish between protocol sections that are well supported and those for which evidence is limited.

Evidence is retrieved via a retrieval-augmented generation (RAG) workflow centered on iBioGenie's internal protocol corpus. The corpus comprises 42,541 protocol documents from 11 sources, including protocols.io (*4*), Nature Protocols, STAR Protocols, Methods in Enzymology, MethodsX, Bio-protocol,

Springer Open Access, F1000Research, Zenodo, bioRxiv, and the Morimoto Lab. Semantic retrieval is followed by LLM-based reranking to identify the procedural information most relevant to each experimental module. When sufficient evidence is not found in the protocol corpus, retrieval can be expanded to the user's literature collection, full-text methods from the literature review system, and relevant web sources. Candidate sources are evaluated for relevance before being assigned to individual modules. The retrieved information is then assembled into a structured context that records the supporting evidence available for each stage, as well as stages for which suitable evidence could not be identified.

Using this evidence, iBioGenie produces a structured wet-lab protocol that can include materials, reagent preparation, detailed procedures, quality-control checkpoints, anticipated results, troubleshooting, and other relevant sections. Each experimental module remains a distinct phase of the final procedure and is written using evidence retrieved specifically for that stage. The system also tracks citations for experimental parameters throughout the protocol. Values supported directly by retrieved evidence are linked to their corresponding sources. When reliable evidence for a particular experimental detail or parameter cannot be identified, iBioGenie does not present the value as evidence-backed. Instead, the lack of support is made explicit so that the parameter can be verified against the appropriate laboratory standard operating procedures (SOPs).

The initial protocol then undergoes several verification steps. iBioGenie checks that the procedure includes operational parameters, verifies user-provided values and citation provenance, and derives structured elements such as plate layouts and reagent totals from the underlying experimental declarations. A feasibility critic agent then reviews the complete protocol for issues such as missing intermediate steps, incompatible experimental conditions, and missing controls. Critical design problems are repaired within the affected sections, after which the revised protocol is re-evaluated. This process can continue for up to three rounds. Any issues that remain unresolved are explicitly reported in the final protocol rather than hidden from the user.

Protocols generated by iBioGenie can be used directly by researchers or passed downstream to BiofoundryAI for translation into executable biofoundry workflows. For automated execution, the wet-lab protocol can be converted into an automation-ready representation that incorporates the relevant biofoundry rules while preserving the original protocol's structure and citations. The adapted protocol is then schematized and submitted to BiofoundryAI for execution. At submission, iBioGenie preserves the exact protocol sent to BiofoundryAI, along with the schema version, BiofoundryAI commit, protocol size, module counts, and a hash of the source protocol, to provide a traceable record linking the original generated protocol to the version submitted for automated execution (**fig. S12**).

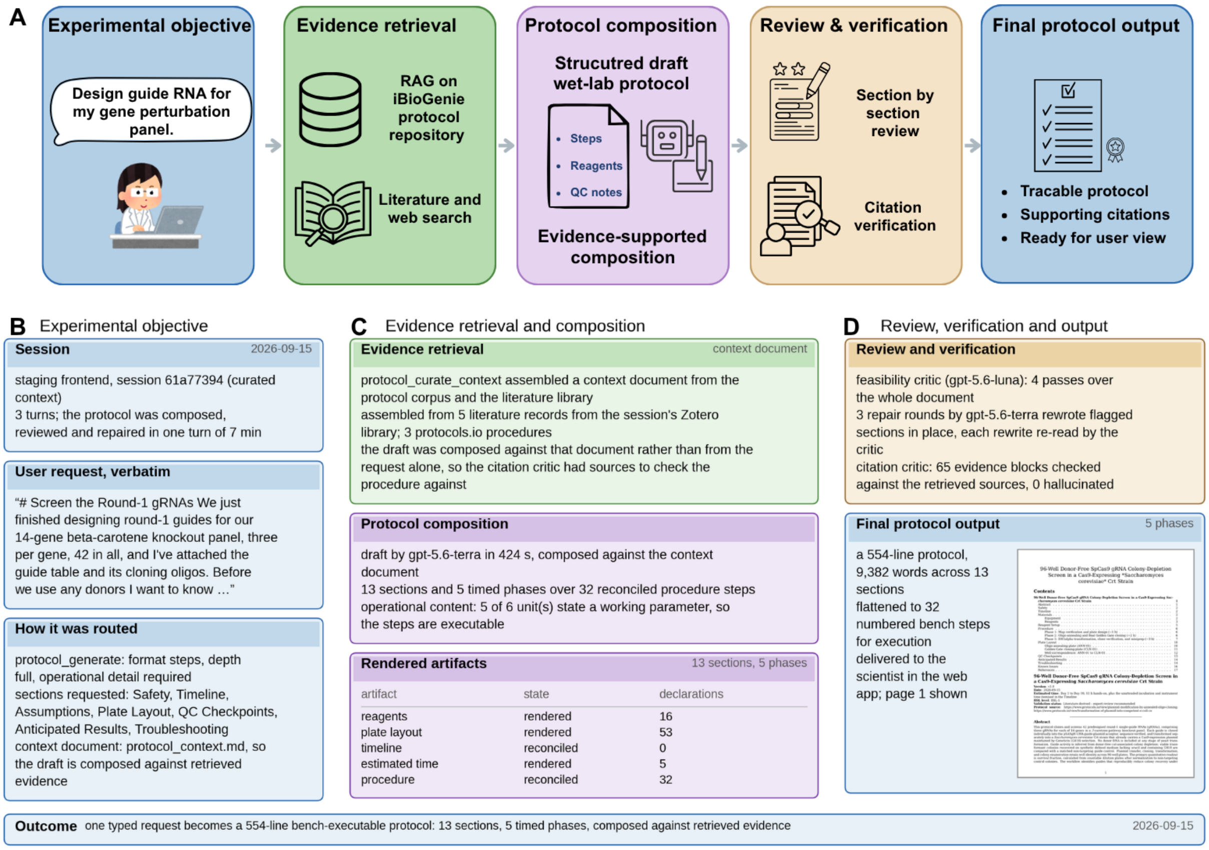


**Fig. S12. iBioGenie wetlab protocol generation with a genetic engineering example. (A)** The generation pipeline. A chat-stated objective drives evidence retrieval from the iBioGenie protocol repository, literature, and web. A structured protocol is then generated, reviewed section by section, citation-checked, and delivered as a traceable output. **(B)** The scientist's objective and routing card define the full protocol request, including safety, timeline, assumptions, plate layout, QC, expected results, and troubleshooting. The protocol was generated, reviewed, and repaired in a single 7-minute turn. **(C)** Evidence retrieval used eight sources. The resulting draft contained 13 sections, 5 timed phases, 32 reconciled steps, 16 reagent declarations, and a 53-well plate layout. **(D)** A feasibility critic completed four review passes across three repair rounds, while a citation critic checked 65 evidence blocks with no hallucinated citations identified. The final 9,382-word protocol was delivered as 32 numbered bench steps in the web application.

#### 3.7. Simulated user testing provides reproducible evaluation of autonomous scientific workflows

To help evaluate whether iBioGenie can reliably complete realistic biological research tasks, we developed a simulated researcher framework that tests the full agent system rather than isolated components. iBioGenie interacts with an independent LLM-based simulated user designed to mimic real scientists and assume different research roles, including principal investigators, graduate students, and research scientists. These simulated researchers submit plausible scientific requests, interact with the system over multiple turns, review generated outputs, and request refinements when necessary. This setup approximates real-world use, requiring iBioGenie to interpret open-ended requests, manage files, select appropriate capabilities, and produce usable scientific deliverables.

The framework supports both focused tool-level evaluations and broader workflow-level experiments. Tool-level tests assess whether a specific capability can be discovered, invoked, and used correctly within a constrained interaction budget. Workflow-level tests evaluate longer research tasks that require combinations of capabilities, planning, computation, and iterative refinement. Each task is specified in natural language to reflect how a researcher would interact with the deployed system, rather than in structured code-level instructions. Required input files are provided via the same file-upload pathway used in production, ensuring that evaluation also tests real session and file-handling behavior. The simulated user continues the interaction until the required deliverables are produced or the interaction limit is reached.

Scoring combines deterministic checks with rubric-based evaluation. For focused capability tests, the framework verifies that the intended capability was used. An independent LLM judge then evaluates the completed session using task-specific criteria derived from the conversation, tool-use trace, and generated files. Scores are computed from criterion-level judgments rather than from an overall score generated directly by the judge.

Finally, to support reproducibility and longitudinal comparison, each run records the task specification, rubric, agent and judge prompts, model and code versions, container version, simulated-user persona, interaction limits, output status, and final score. Results are stored in persistent run registries, enabling performance changes to be traced to differences in tasks, prompts, models, code, or evaluation settings across development cycles (**fig. S13**). The simulated researcher also enables the production of reproducibility trajectories for case studies by providing a single detailed prompt and continuing a multi-turn conversation until research deliverables are satisfied (**Section 9.1**).

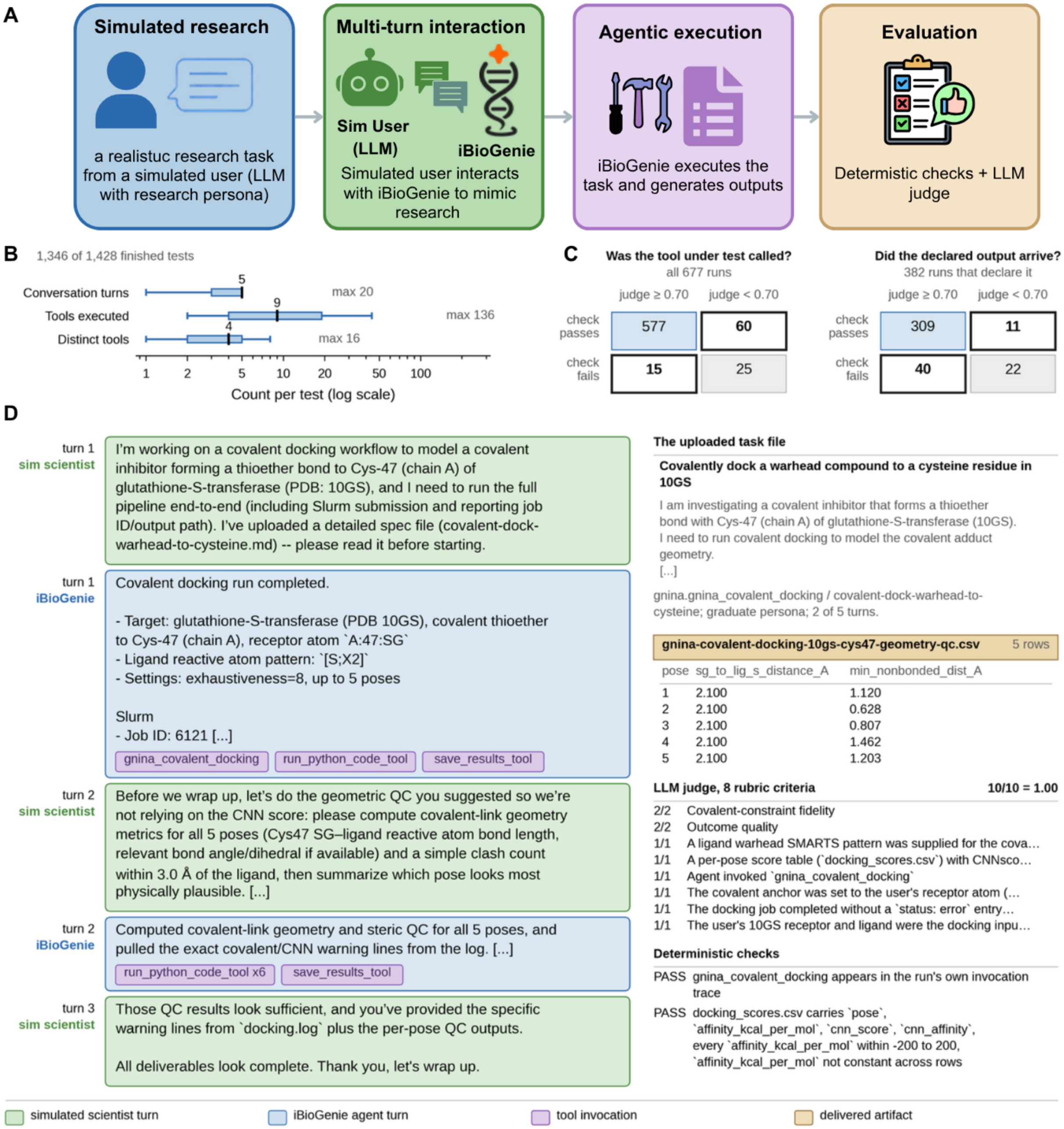


**Fig. S13. Simulated-user testing framework for testing tools and running 1-shot design trajectories.** **(A)** The evaluation loop. An LLM simulated user with a research persona works through a realistic task with iBioGenie over multiple turns; the run is then graded by deterministic checks, which validate the outputs, and by an LLM judge, which grades the work. **(B)** Scale of a single test, over 1,346 of 1,428 finished tests with a tool-chain log. Boxes span the quartiles, whiskers the 5th–95th percentiles, and the black line is the median. **(C)** Deterministic checks versus the LLM judge at a 0.70 pass threshold across 677 tests, including 382 with explicit output checks. The two methods identified complementary failures, with 51 tests failing checks but passing the judge and 44 showing the reverse. **(D)** an end-to-end example of covalent docking onto Cys-47 of glutathione S-transferase (PDB 10GS). The agent flags that the network score is uncalibrated for covalent docking and computes covalent-link geometry and steric metrics for all five poses instead. Both legs pass on different grounds: the judge grades the chemistry and reporting against eight rubric criteria; the deterministic check confirms the tool's own per-pose score table has the expected columns with in-range, non-constant affinities.

#### 3.8. Automated tool addition expands the iBioGenie action space

To make iBioGenie extensible beyond a fixed set of predefined tools and skills, we developed an independent automated tool-addition system that converts external scientific software into validated actions within the iBioGenie action space. This tool-addition platform operates as a separate system specialized for tool ingestion, implementation, testing, and evaluation. Given a candidate tool, it consumes available sources of technical information, including tool descriptions, documentation, API references, GitHub repositories, source code, examples, and associated publications. The system first extracts a structured specification that describes the tool's purpose, required and optional inputs, expected outputs, execution environment, dependencies, hardware requirements, runtime behavior, and the biological tasks for which the tool is appropriate. This specification is then used to generate tool wrappers, input schemas, execution logic, integration metadata, a detailed description, and example usage, enabling iBioGenie to discover, select, and invoke the tool during orchestration (**fig. S14**).

The tool addition process is designed to prevent shallow or unreliable integrations. Each candidate tool must pass multiple validation layers before entering the production action space. These layers include unit tests for individual functions, smoke tests on representative inputs, input–output schema validation, and runtime checks to confirm that returned artifacts match the expected format. For tools requiring high-performance computing (HPC), the system also performs SLURM-based execution tests to verify that jobs can be submitted, monitored, and completed correctly in the target computing environment.

Beyond execution-based testing, the system uses an LLM-based assessment layer to generate tool-specific specifications and evaluation rubrics. These define the tool's intended scientific capabilities, valid inputs and outputs, and the functionality that the generated integration should expose. This is especially important for biological tools that support multiple operating modes rather than a single function. The evaluation therefore assesses coverage of relevant parameters, file formats, batch behavior, execution modes, and biologically meaningful outputs, preventing integrations from being reduced to thin wrappers around a single example command. The rubric also evaluates whether iBioGenie can retrieve the tool from natural-language task descriptions and select it appropriately based on biological relevance.

After implementation and evaluation, each integration is scored against its specification and rubric. Tools that do not meet the required execution, capability-coverage, or retrieval criteria are returned for revision rather than added to the action space. This validation framework enables iBioGenie to incorporate new databases, predictive models, command-line tools, and web APIs while maintaining a reliable, reproducible scientific capability base.

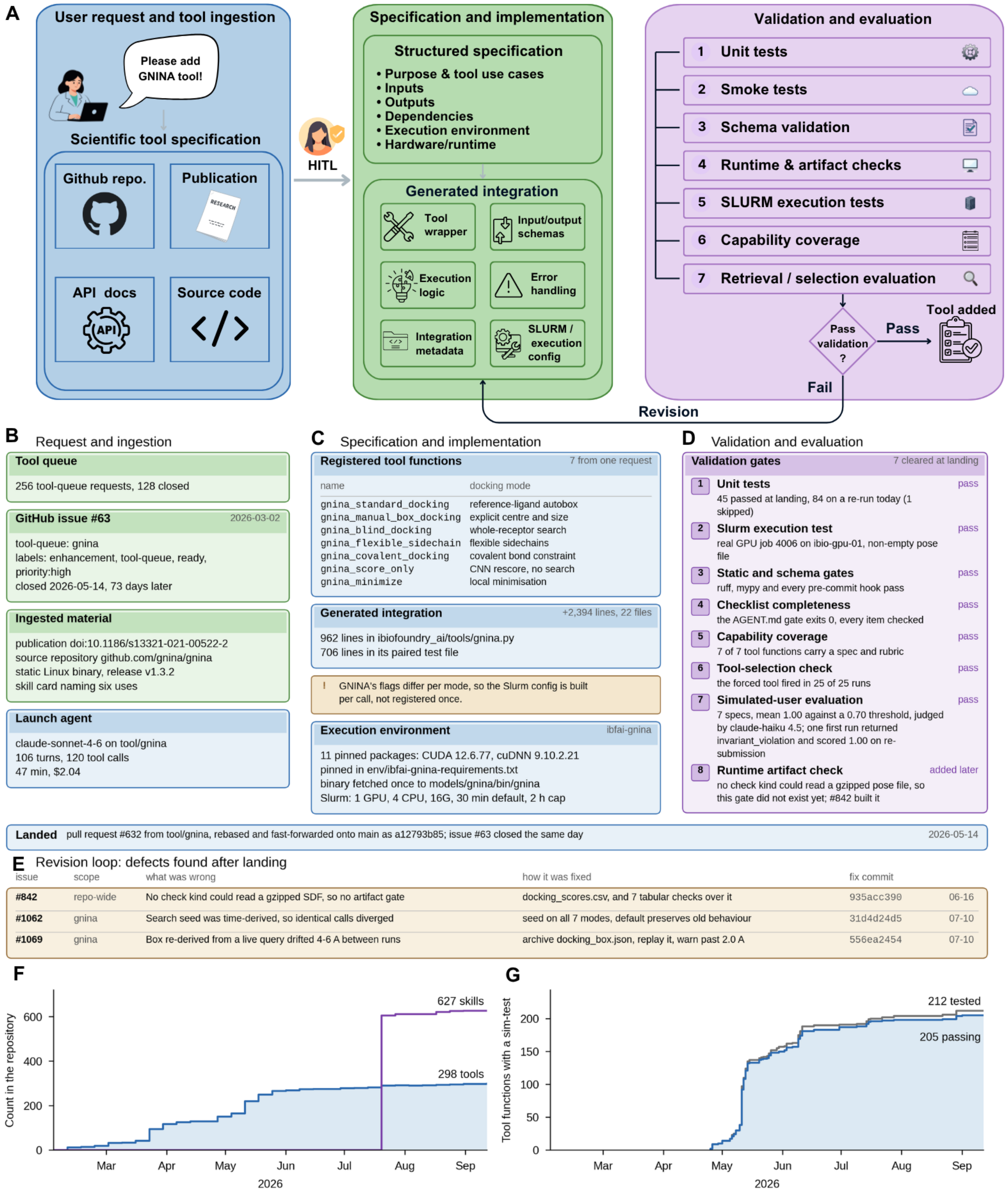


**Fig. S14. Automated tool addition with GNINA tool as an example. (A)** The addition pipeline. A user request names a tool; its repository, publication, and documentation are ingested into a structured specification; the agent generates the wrapper, schemas, execution logic, and Slurm configuration; and seven validation gates run in order. A failure returns to implementation; only a pass adds the tool. **(B)** The request. The recorded launch-agent session ran 106 turns and 120 tool calls in 47 minutes. **(C)** Implementation. One request yielded seven tool functions, one per docking mode, across 2,394 lines in 22 files, running as a one-GPU Slurm job in a pinned conda environment. **(D)** Validation. All seven gates passed before landing, including a real GPU execution test, a tool-selection check that fired in 25 of 25 runs, and seven simulated-user tests scoring a mean 1.00 against a 0.70 threshold. **(E)** Landing and follow-up. The branch landed and the issue closed the same day; three defects found between 16 June and 10 July were each closed by a named commit with its own regression test. **(F)** Cumulative tools and skill cards from commit history: 298 tools and 627 skills. **(G)** Cumulative simulated-user coverage: 212 tool functions have been tested and 205 have passed at least once.

#### 3.9. iBioGenie facilitates continuous capability integration and self-improvement

Maintaining iBioGenie as a growing scientific platform requires more than adding new capabilities. Tools evolve, users expose failure modes that are difficult to anticipate during development, and changes to either tool implementations or their use by the agent can affect system performance. To address this, iBioGenie uses a continuous self-improvement framework that integrates feedback from real users, allowing capabilities to be updated and revalidated as the platform evolves while keeping the underlying action space reliable for the orchestrator and other agent systems that may use it.

Tools and skills remain updatable even after they enter the action space. Real-world use may reveal bugs on specific inputs, missing functionality, limitations, or cases where the agent selects an inappropriate mode or parameterization. To capture these issues directly from users, the iCloudBiofoundry application includes a feedback option at each interaction turn, allowing researchers to comment on a specific response, the broader conversation, or a particular use of the system. This feedback provides direct evidence of how capabilities behave in practice and can trigger revisions to a tool or skill, its interface, or the metadata that determines when and how it is selected. iBioGenie handles these revisions through a specification-driven pipeline. When testing or user feedback identifies a problem, a revision system receives the existing integration and the relevant evidence, modifies the implementation or associated metadata, and submits the change through the same validation gates required for new tools. (**fig. S15**).

Every new or revised capability or feature is evaluated through an automated simulated-user testing pipeline before deployment. Each test begins with a biological objective written as a researcher might enter it through the interface and is paired with a task-specific evaluation rubric. An LLM-based simulated researcher interacts with iBioGenie over multiple turns, while the orchestrator runs the same containerized service used in production. After each interaction, an independent LLM judge scores the conversation and generates files against the task-specific rubric. Each run also records the evaluation setup and execution trace, so performance can be compared over time. Deterministic checks complement this scoring by confirming that the intended capability was used and that it can be retrieved from a realistic scientific request without being explicitly named.

Real-user feedback is the other major source of improvement. Because iBioGenie is designed for researchers who may not be software developers, feedback can be submitted directly alongside an agent response rather than through a technical issue tracker. Each report is sent to the maintenance pipeline with a link to the corresponding session. A triage agent evaluates the feedback using the full session context, including the conversation, tool-use trace, logs, and generated files, rather than relying solely on the short user comment. It then assigns a category and priority, identifies supporting evidence, and either creates a new engineering issue or adds information to an existing one if the same problem has already been reported. Repeated reports can increase an issue's priority, while feedback that reveals a broader coverage gap can be converted into a new simulated-user test (**fig. S15**).

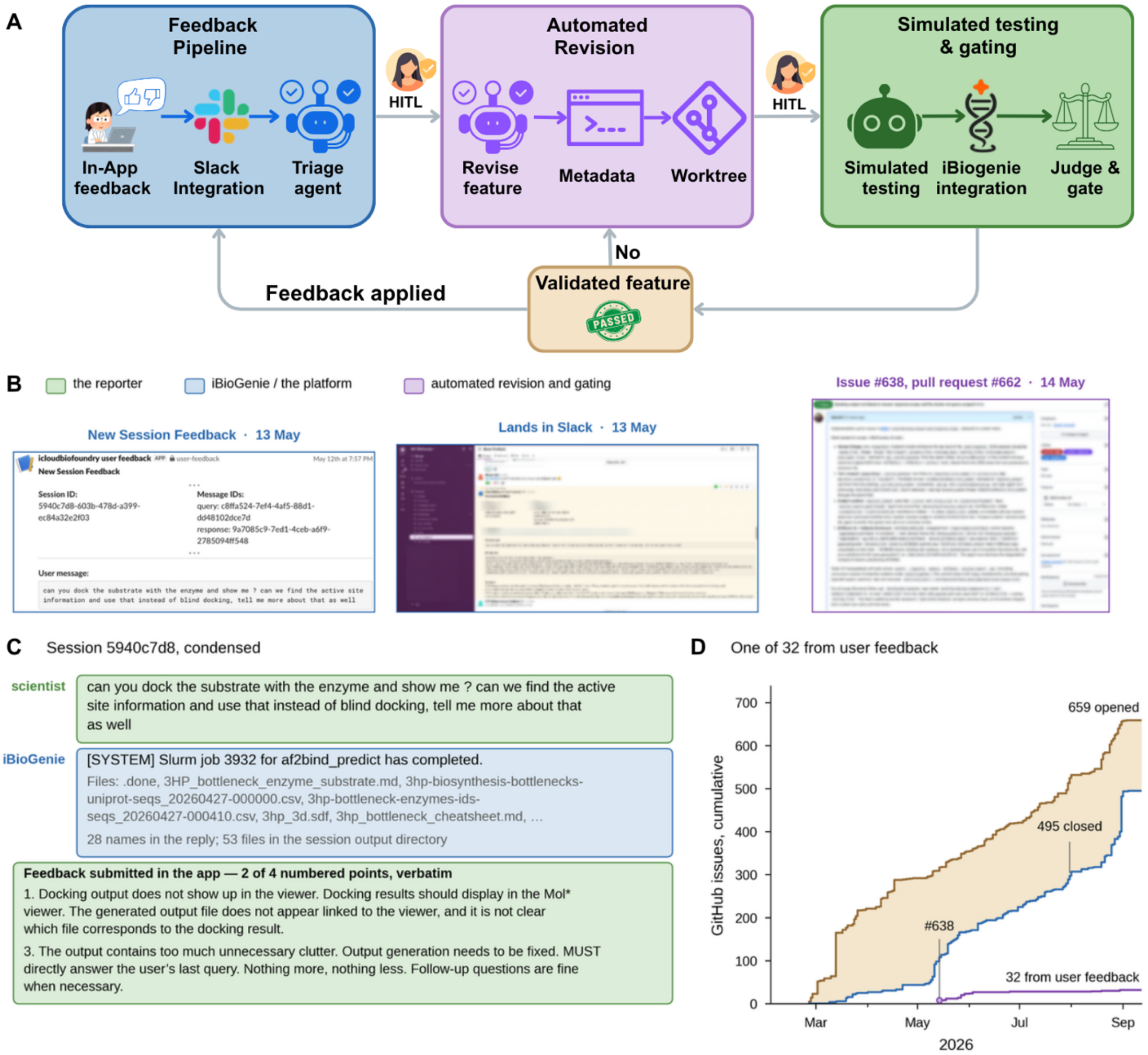


**Fig. S15. Feedback-driven continuous improvement with an example. (A)** The improvement loop. In-app feedback is persisted and posted to Slack; a triage agent classifies it into an issue; a revision agent turns the issue into a change; and the change is validated by simulated testing and gated by an LLM judge before it lands. A human researcher from iBioGenie team approves after triage and before the gate; a change that fails validation returns to revision. **(B)** One report traced end to end. Feedback on 13 May opened issue #638 on 14 May; pull request #662 followed on 15 May with 67 new or updated unit tests and a new simulated-user test; the change landed on 17 May and the issue closed on 18 May, 5 days after the report, with an automatic message back to the reporter. **(C)** The session that provoked the report, condensed verbatim to the flagged exchange and two of the reporter's four points. **(D)** Cumulative GitHub issues: 659 opened, 495 closed, and 32 user feedbacks, of which 27 are closed.

### 4. Benchmarking for iBioGenie

#### 4.1. iBioGenie benchmarking system configuration and baselines

To evaluate iBioGenie alongside alternative systems under identical conditions, all systems were run through a single harness that exposes one standardized input-output interface. For each benchmark item, the harness renders the prompt, fixes the answer-option order where applicable, submits the item to the system under test, extracts the answer from the raw response, and applies the scoring rule; only the component that produces the response differs between systems. Each evaluated item is stored as a per-question record containing the rendered prompt (or its hash), the option order, the complete raw response, the extracted answer, the scoring outcome and its class, latency, retry count, token usage, estimated model cost, and the identity of the software build that produced the response. Every quantity reported in this work is recomputed directly from these records by scripts released with the data.

Four systems were evaluated. iBioGenie coordinates multiple specialized agents, whereas Biomni A1 (v0.0.8, commit 400c1f3) processes each task through a general-purpose agent workflow with code execution. Before any reported Biomni run, one correction was made to its agent loop. Biomni's controller assumes that each model message contains at most one action tag and relies on stop sequences to enforce this behavior. However, reasoning models such as gpt-5.6-terra do not reliably honor these stop sequences, allowing a single response to contain both an <execute> code block and a premature <solution> answer. In the unmodified loop, the presence of the solution caused execution to terminate before the accompanying code was run, meaning the agent could answer from an unexecuted plan. In a diagnostic run of the analytical benchmark performed before the correction, 46 of 50 tasks terminated after the first model response without executing any code. The corrected loop instead executes any pending code before accepting a solution and terminates only when a solution is returned without an accompanying code block. This modification favors the baseline, was applied consistently across all reported Biomni runs, and is included in the released code.

The remaining two systems serve as model-only references. Each submits the benchmark prompt as a single completion call, without tools, planning, or retrieval: one uses gpt-5.6-terra at low reasoning effort, and the other uses claude-opus-5 with extended thinking disabled, the corresponding setting for that model. iBioGenie, Biomni A1, and the gpt-5.6-terra reference arm therefore share the same underlying language model, allowing differences among these three systems to be attributed to their surrounding agent architectures rather than to the base model.

For reproducibility, the model identity of every iBioGenie response was verified from per-turn server-side usage records rather than inferred from configuration. All iBioGenie results were generated using a single frozen build (2026.08.31-ab2da0971), with the build identifier recorded in the manifest of every run. In addition, the scored input files used for the question-answering replicates were preserved as write-protected snapshots, preventing later activity on shared storage from modifying the data used for scoring.

Each benchmark was executed three times per system as fully independent replicate runs, and results are reported as the mean ± sample standard deviation over the three replicates, with all individual run values released. Within a replicate, all systems receive byte-identical prompts with identical option order, so item-level outcomes are paired across systems. Accuracy is computed over answered questions: a request that fails at the infrastructure level (a transport failure, or a response that never reached the system) is excluded from the denominator and reported as a separate outcome class, and infrastructure-failure rates were at most 2.3% in every reported run. Acceptance of a replicate required completeness, a single recorded build across all records, and a failure rate below 5%. One earlier iBioGenie question-answering run failed this gate during a service outage, with 63.4% of its requests failing at the infrastructure level; it was excluded under this rule and replaced by a complete run, and every reported replicate passed the gate, except as noted for the coding-agent reference arm in Section 4.3. Wrong answers remain in the denominator, including responses whose content could not be interpreted as an answer. Per-question token usage and estimated model cost are recorded for every run and released with the data; estimated cost covers model tokens only and excludes non-model compute, storage, and external-tool charges. The evaluation scores final answers only: complete response text, and for the agentic systems the execution trajectory and step counts, are recorded in the released files, but no step-level or process-level judgment contributes to any reported number.

Answer extraction and grading were audited rather than assumed correct. Because the systems format answers differently (answer tags, solution blocks, reasoning preambles, markdown emphasis), the shared extraction layer recognizes each of these envelopes for every system. Extraction rules were derived from the response formats in general, never from any individual item, and each rule change was verified to leave the other systems' scored outcomes byte-identical. Two grading corrections are applied uniformly on top of the benchmarks' own scoring. First, answers whose gold standard is numerically decidable, an exact value, a stated interval, or a leading numeric magnitude, are decided arithmetically before any language-model judge is consulted: interval bounds are sorted, and a value is accepted when it matches the gold at its written precision or within 1% relative error. This confines the judge to genuinely categorical prose, where its verdict is otherwise not reproducible across runs. Second, dictionary-shaped answers whose values are emitted as unquoted tokens are restored to valid literals before comparison; this repair is purely syntactic and never consults the gold answer. In addition, a deliberately over-permissive independent re-grade of every scored item served as an audit: every disagreement between the audit and the reported grades was inspected individually, and the audit is released with the data. The released per-question records contain all corrections, the items they affect, and both the benchmark-native and corrected scores (**fig. S16**).

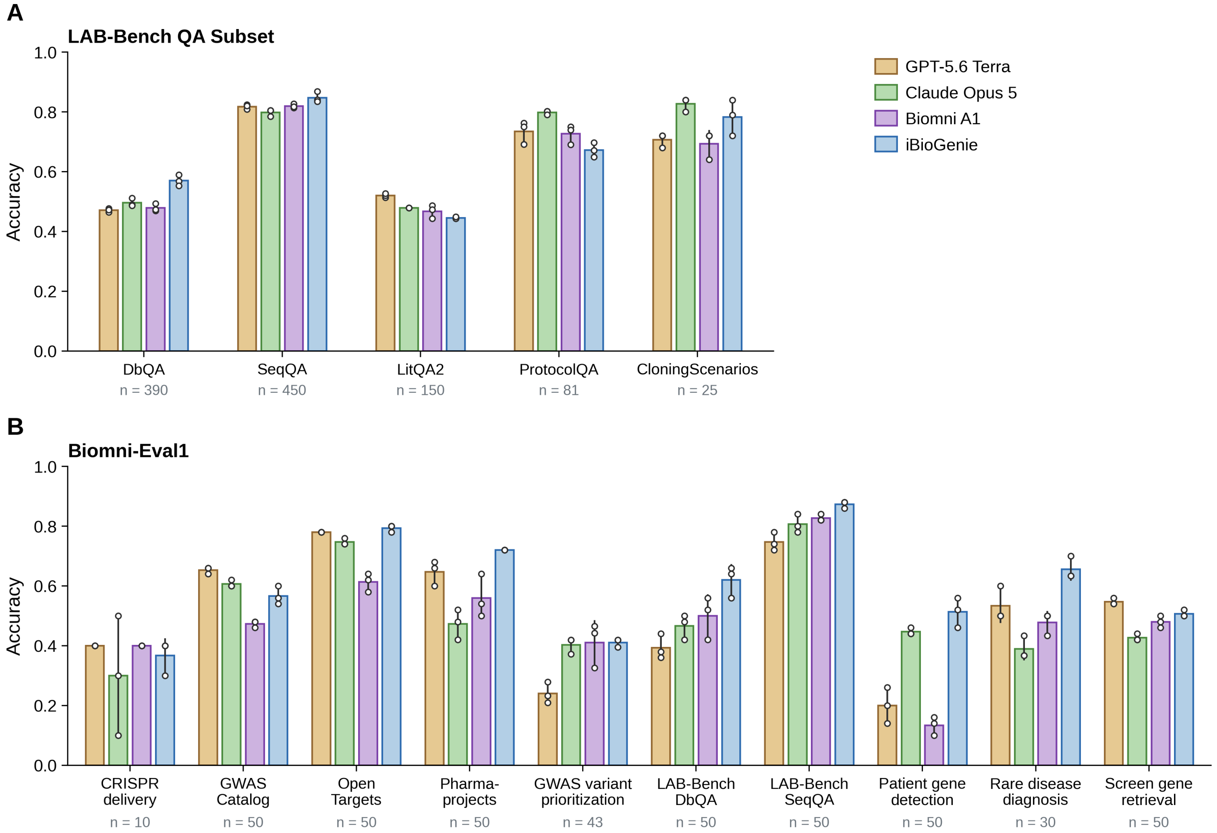


**Fig. S16. Per-class benchmarking results comparing iBioGenie against baseline systems on LAB-Bench Subset and BiomniEval1.** iBioGenie and Biomni A1 are given GPT-5.6 Terra as their base model; Claude Opus 5 and GPT-5.6 Terra use the base model without any tools. All systems were run on the same items with the same grader. Bars show means of three replicates, error bars show the standard deviation, and open circles show individual replicates. **(A)** LAB-Bench QA subset by dataset (1,096 questions): DbQA (n = 390), SeqQA (450), LitQA2 (150), ProtocolQA (81), and CloningScenarios (25). **(B)** BiomniEval1 by task family (433 queries): CRISPR delivery (10), three GWAS causal-gene families (50 each), GWAS variant prioritization (43), LAB-Bench DbQA and SeqQA subsets (50 each), patient gene detection (50), rare disease diagnosis (30), and screen gene retrieval (50). Per-dataset LAB-Bench means are tabulated in table S2.

#### 4.2. Multiple-choice QA benchmark (LAB-Bench QA Subset)

The LAB-Bench evaluation covers the five text-based components, DbQA, SeqQA, LitQA2, ProtocolQA, and CloningScenarios, which assess database interpretation, nucleotide- and protein-sequence reasoning, literature synthesis, protocol troubleshooting, and multistep molecular-cloning reasoning, respectively (*5*). FigQA, TableQA, and SuppQA require figure, table, or supplementary-file inputs and were outside the scope of this evaluation. Questions of each component were partitioned into four near-equal subsets with a fixed seed; three of these subsets form the evaluated pool of 1,096 questions, and the fourth was not used in this work (table S1). Every replicate run evaluates all 1,096 questions. Answer choices are permuted deterministically from the question identifier, so the order is identical for every system and reproducible across executions, and no system can benefit from a fixed position of the correct answer. A response is correct only when the parsed answer letter matches the designated answer (**table S2**).

**Table S1. Dataset sizes.** Full public split and one 25% subsample.

| Dataset | Full | Per subsample |
|---|---|---|
| DbQA | 520 | 130 / 130 / 130 |
| SeqQA | 600 | 150 / 150 / 150 |
| LitQA2 | 199 | 50 / 50 /50 |
| ProtocolQA | 108 | 27 /27 /27 |
| CloningScenarios | 33 | 9 / 8 / 8 |
| Total | 1,460 | 1,096 over three subsets |

**Table S2. LAB-Bench QA Subset accuracy.** Values are the mean ± SD over three independent replicate runs (pooled over the three subsamples within each run; accuracy over answered questions)**.** The highest accuracy in each row is in bold.

| Dataset | Biomni A1 | OpenAI (no tools) | Claude (no tools) | iBioGenie |
|---|---|---|---|---|
| DbQA | 47.84 ± 1.32 | 47.09 ± 0.65 | 49.60 ± 1.37 | **57.01 ± 1.85** |
| SeqQA | 81.93 ± 0.68 | 81.78 ± 0.80 | 79.78 ± 1.15 | **84.75 ± 1.80** |
| LitQA2 | 46.77 ± 2.24 | **52.00 ± 0.67** | 47.86 ± 0.00 | 44.50 ± 0.35 |
| ProtocolQA | 72.66 ± 3.20 | 73.46 ± 3.80 | **79.84 ± 0.71** | 67.24 ± 2.44 |
| CloningScenarios | 69.33 ± 4.62 | 70.67 ± 2.31 | **82.67 ± 2.31** | 78.32 ± 6.02 |
| Pooled | 64.01 ± 0.15 | 64.49 ± 0.35 | 65.02 ± 0.84 | **67.88 ± 0.65** |

#### 4.3. Analytical benchmark (BixBench)

The second evaluation track uses BixBench to test end-to-end analytical capability (*6*). Each task provides the raw data from a published study and asks the system to carry out a specified analysis and report the result. Tasks are distributed as capsules containing both the raw input data and a complete reference analysis with the expected answer. For evaluation, the harness exposes only the raw-data portion to the system and aborts the task if any part of the reference analysis becomes accessible. This ensures that systems work from the original study data without access to the answer or to the analysis steps used to obtain it. Each attempt is limited to 900 seconds of wall-clock time, and tasks that exceed this limit are recorded as timeouts within the infrastructure-failure class.

We use the expert-audited Verified-50 subset, which contains fifty questions drawn from thirty-three capsules. In this subset, domain experts removed flawed items and corrected both question wording and reference answers, providing more reliable ground truth than the unaudited benchmark. Of the fifty questions, seventeen are graded by exact string match, thirteen by numeric-range containment, and twenty by a language-model judge using gpt-4o in all reported runs (**table S3**). During evaluation, we found two reproducible problems in the benchmark-native grading. Some interval answers were stored with bounds in descending order, making the range check impossible to satisfy, and the language-model judge sometimes returned different verdicts for identical answer–gold pairs across runs, including purely numeric cases. We therefore use the corrected grading procedure (**Section 4.1**); interval bounds are sorted before comparison, and all numerically decidable answers are evaluated arithmetically before invoking the judge. These corrections are applied identically to every system, while the original benchmark-native grade is preserved in the released records.

Because BixBench requires code execution on the staged data, the two model-only reference arms cannot meaningfully complete the benchmark and are included only as behavioral and data leakage controls. gpt-5.6-terra nevertheless tends to produce confident numeric answers, with 45 of 50 items changing between replicates, and obtains a score of 10.00 ± 2.00. By contrast, claude-opus-5 usually attempts to invoke execution tools that are not available and scores 0.67 ± 1.15. When the comparison is restricted to genuine numeric attempts (predictions that parse as numbers, excluding tool-invocation output), the two models are statistically indistinguishable: 1 of 17 attempts was correct for claude-opus-5 and 8 of 122 for gpt-5.6-terra across the three replicates (Fisher's exact test, $p = 1.00$). The attempt counts are themselves the signal: claude-opus-5 produced 17 numeric attempts against 122 for gpt-5.6-terra, so the former declined most items while the latter answered anyway. Refusal here means the model states that it cannot answer without the information the analysis would supply, which is the calibrated response when no execution tool is available. These model-only rows therefore reflect behavior under an impossible task setting, namely the tendency to guess versus refuse, and are excluded from the headline comparisons.

To evaluate these models under conditions that permit code execution, both were additionally assessed as coding agents using the benchmark's own baseline scaffold, in which the model drives a Python Jupyter notebook opened in the task's staged data directory through three tools: editing a cell, executing it, and submitting an answer. No browsing or retrieval tools are provided to either baseline model in this scaffold, and the recorded trajectories contain no network access, so every analysis draws only on the staged data. Each task allows 20 agent steps within the same 900-second budget; one to three tasks per replicate exceeded it on the largest capsules and are recorded as timeouts, the only replicates in this work above the 5% infrastructure-failure gate (**Section 4.1**). Under the identical harness, grading, and three-replicate protocol, claude-opus-5 reaches 74.51 ± 2.72 and gpt-5.6-terra 63.33 ± 2.31. Both fall between Biomni A1 and iBioGenie, indicating that the remaining gap to iBioGenie reflects its agent architecture and access to relevant actions for the benchmark.

**Table S3. Verified-50 composition and accuracy.** Values are percentages under corrected grading, mean ± SD over three replicate runs, with individual runs in parentheses. The highest accuracy in each row is in bold.

| | n | Biomni A1 | OpenAI agent | Claude agent | iBioGenie |
|---|---|---|---|---|---|
| String match | 17 | 38.11 ± 7.64 | 60.78 ± 6.79 | 68.01 ± 3.01 | **78.43 ± 3.40** |
| Numeric range | 13 | 38.46 ± 13.32 | 66.67 ± 4.44 | **83.76 ± 0.74** | 76.92 ± 0.00 |
| LLM-judged category (arithmetic pre-decision applied) | 20 | 50.61 ± 14.09 | 63.33 ± 7.64 | 74.21 ± 4.50 | **90.00 ± 5.00** |
| Overall | 50 | 43.20 ± 6.64 (50.00 / 36.73 / 42.86) | 63.33 ± 2.31 | 74.51 ± 2.72 | **82.67 ± 1.15 (82.00 / 84.00 / 82.00)** |

#### 4.4. Biomedical task benchmark (BiomniEval1)

BiomniEval1 contains 433 open-answer questions spanning 10 task families (*7*). These include CRISPR delivery selection (10 questions per run); GWAS causal-gene identification using GWAS Catalog, Open

Targets, and Pharmaprojects (50 questions each); GWAS variant prioritization (43); database and sequence question answering (50 each); patient-gene detection (50); rare-disease diagnosis (30); and screening-hit gene retrieval (50). Following the Biomni report, we score each task family separately using its corresponding exact-match rule and report the macro average across the ten families. This prevents the larger task families from contributing disproportionately to the overall score. We also retain per-question results, allowing the micro average to be recomputed; for the systems evaluated here, it is roughly three percentage points higher than the macro average.

One detail is worth noting for comparison with the original Biomni results. Screening-hit gene retrieval was evaluated there using the average post-perturbation effect rather than exact-match accuracy, so our score for this family is not directly comparable. In practice, however, removing this family changes the overall macro averages by less than one percentage point. All four systems were evaluated using the same extraction and grading procedure described in Section 4.1. This includes a small syntactic literal-repair step, which did not change any result for the three baseline systems but recovered thirteen iBioGenie responses that contained the correct answer in a structurally malformed format (**table S4**).

**Table S4. BiomniEval1 macro-averaged accuracy.** Mean ± SD over three replicate runs is shown**.** The highest accuracy is in bold.

| System | Macro accuracy | Individual runs |
|---|---|---|
| iBioGenie | **60.26 ± 0.16** | 60.32 / 60.09 / 60.39 |
| OpenAI (no tools) | 51.40 ± 0.94 | 51.29 / 52.39 / 50.53 |
| Claude (no tools) | 50.65 ± 2.04 | 50.85 / 48.52 / 52.59 |
| Biomni A1 | 48.75 ± 0.67 | 49.25 / 49.02 / 47.99 |

#### 4.5. Construction of the internal wet-lab protocol generation benchmark

We assembled 3,307 protocol PDFs from eleven repositories (Bio-protocol, bioRxiv, F1000Research, Methods in Enzymology, MethodsX, the Morimoto Lab, Nature Protocols, protocols.io, Springer Open Access, STAR Protocols, and Zenodo) and converted them to structured Markdown using gemma-4-26B-A4B-it. The converted protocols were classified into 17 technique classes, and those in Synthetic Biology and Bioengineering were reviewed to form the 139-item evaluation set (**fig. S17**). Each item pairs a self-contained natural-language objective (median 665 words) with an expert-reviewed reference protocol; the question does not name its source publication, and the reference is never shown to the system under test.

To ensure consistent scoring across expert-annotated and model-generated protocols, both are expressed in a single JSON template with five sections: biological context (organism or chassis and genetic materials), overview (workflow string, pause points, controls), safety and compliance (hazard, material, protective equipment, disposal), advance preparation, and major steps. Each major step includes its purpose, duration, inputs, instruments, reagents, numeric conditions with tolerances, quality checks, an ordered list of actions individually flagged for criticality, an expected outcome, troubleshooting entries, and a block that names which later step consumes each output. Reagents, instruments, and genetic materials carry short identifiers that step actions reference, and step outputs point forward using step tokens; both conventions are read by the structural scoring (**Section 4.6**).

The reference protocols contain 812 major steps (median of 5 per protocol) and 3,584 actions, of which 116 (3.2%) are annotated as critical (**table S5)**. The template is populated unevenly: biological context, overview, and major steps appear in all 139 references; advance preparation appears in 85%; and safety and compliance in 53%. At the step level, actions appear in 99% of steps and declared outputs in 90%, but quality checks in only 29% and troubleshooting in 13%.

**Table S5. Composition of the internal wet-lab protocol generation benchmark (n = 139).**

| Property | Value |
|---|---|
| Source repositories | protocols.io 95 (68%), STAR Protocols 15 (11%), Nature Protocols 13 (9%), Bio-protocol 12 (9%), Springer Open Access 2 (1%), Morimoto Lab 2 (1%) |
| Publication year | 2006–2026; 101 of 136 records with a parseable date are 2019 or later |
| Question length | median 665 words (range 405–903) |
| Major steps per protocol | median 5 (range 1–21); 812 steps in total |
| Actions per major step | median 3; 3,584 in total, of which 116 (3.2%) annotated critical |
| Distinct reagents, instruments per protocol | median 9 (range 0–42), median 4 (range 0–18) |
| Section populated in the reference | biological context 100%, overview 100%, major steps 100%, advance preparation 85%, safety and compliance 53% |
| Step field populated (% of 812 steps) | actions 99%, output 90%, input 88%, materials and reagents 79%, conditions 70%, instruments 63%, duration 44%, expected outcome 40%, quality checks 29%, troubleshooting 13% |
| Steps whose output names its consuming step | 590 of 812 (73%) |

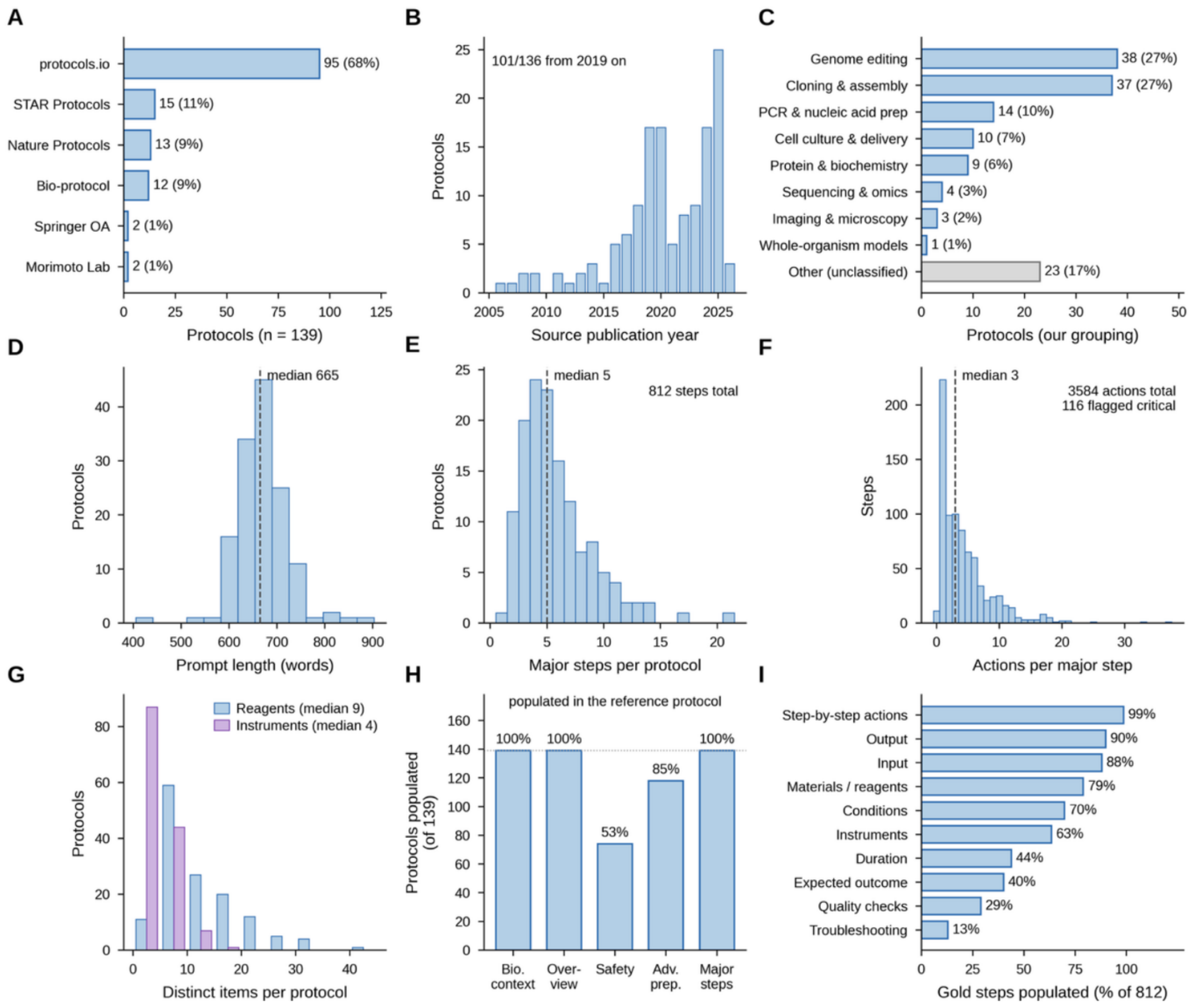

**Fig. S17. Composition of the internal wet-lab protocol generation benchmark.** All panels describe the 139-protocol corpus: the top row the corpus, the middle row cover tasks posed to the model, and the bottom row the structured reference protocols. **(A)** Source repository. **(B)** Publication year; 101 of 136 dated records are from 2019 or later. **(C)** Technique theme, assigned by us from subcategory tags and keywords; the unclassified residual is shown in gray. **(D)** Prompt length in words; median 665, range 405–903. **(E)** Major steps per reference protocol; median 5, range 1–21, 812 in total. **(F)** Actions per major step; median 3, 3,584 in total, 116 (3.2%) flagged critical. **(G)** Distinct reagents and instruments per protocol; medians 9 and 4. **(H)** Fraction of references populating each judged section; safety and compliance is present in only 53%. **(I)** Fraction of the 812 reference steps carrying each per-step field; actions, inputs, and outputs are nearly always present, while duration, quality checks, and troubleshooting rarely are.

#### 4.6. Internal wet-lab protocol generation benchmark scoring and performance comparison

Seven systems were scored on all 139 items, with three independent generations each: the four systems of Section 3.1, two additional iBioGenie configurations differing only in retrieval, and an Oracle arm in which the exact source protocol is injected into the context. The Oracle measures what perfect retrieval buys on this corpus rather than setting an upper bound, which is 1.0 by construction; its distance from 1.0 reflects how much of the residual gap is attributable to the structured protocol output and grading rather than the generating system.

Each generated protocol is scored on three axes. A deterministic structural score, D = 0.5 s_schema + 0.3 s_ref + 0.2 s_wf, measures schema validity, the fraction of internal identifier and step references that resolve, and agreement between the step count declared in the overview and the number of major-step objects. An LLM judge (gpt-5.4, medium reasoning effort) scores each of the five template sections on [0, 1] against fixed anchors, from 1.00 for semantic equivalence, with synonyms and reordering permitted, to 0.00 for missing or contradictory content, aligning candidate to reference major steps by purpose. Reference-based BLEU, METEOR and ROUGE-L were computed only as a negative control: two unrelated expert protocols score a higher corpus BLEU against each other (1.94) than four of the seven systems score against their own correct reference, so these metrics are not used to rank systems.

The reported score is the unweighted mean of seven equally weighted terms: the five judged sections plus two terms computed directly from the generated JSON. Step chaining is 1 only when every declared step output resolves to a later step or to the terminal sink; a single unresolved edge fails the term. 590 of 812 reference steps declare such an output. Numeric fidelity is the fraction of reference quantities reproduced within tolerance (5% relative, 1 °C or 0.1 pH) after label-matched one-to-one alignment; a protocol cannot match by emitting many quantities. Both terms were admitted on a gate requiring the ratio of between-system to within-system standard deviation to exceed approximately 2, which they clear at 11.5 and 9.0; a third candidate, severity-weighted omission of critical actions, failed at 1.4 and is not reported. The deterministic score is reported as a structural floor rather than a scored term, because it spans only 0.979 to 0.999 across all systems and runs. Scored on the judged sections alone, reweighted on reproducibility grounds and without the two computed terms, the iBioGenie margin over Claude (no tools) reverses from +0.016 to −0.001.

Results are the mean ± SD over three generations. Unparseable predictions are scored as zero on every term and are never excluded from any denominator; they were stable across replicates at 2 of 139 per run for Claude (no tools) and Biomni A1, 1 for OpenAI (no tools), and 0 for iBioGenie. Regarding identical frozen predictions, the judge, not the model, dominates the error bars, accounting for 45 to 100% of each system's replicate variance, so differences below approximately 0.008 are not resolvable on this benchmark. iBioGenie reached 0.651, above the strongest baseline, Claude (no tools) at 0.635, and approaching the Oracle at 0.661 (**table S6**). The per-term breakdown (**fig. S18)** shows the two leading systems reaching similar totals by different routes: Claude (no tools) leads on safety and compliance and on numeric fidelity, while iBioGenie leads on biological context, overview, and step chaining, the one term on which Claude (no tools) falls below 0.97 while every other system exceeds it. Claude (no tools) also generates 9,323 words per protocol against 1,347 for the expert references and 2,056 to 2,989 for every other system, with no overlap in length distribution. Because no other system produces protocols of comparable length, there is no common length range over which to fit a correction, so any length-adjusted comparison against Claude (no tools) would have to extrapolate beyond the observed data. Assigning each protocol to the term accounting for the largest difference between the two systems, step chaining decides 45 protocols and favors iBioGenie 44 to 1, while safety and compliance decides 21 and favors Claude (no tools) 18 to 3.

**Table S6. Per-term and overall scores on the internal wet-lab protocol generation benchmark.** Mean ± SD over three replicate runs**.** Each term is in [0, 1]. The first five are scored by the LLM judge against the expert reference; step chaining and numeric fidelity are computed from the generated JSON. The overall score is the unweighted mean of the seven terms. The highest score in each column, excluding the Oracle, is in bold.

| System | Biological context | Overview | Safety / compliance | Advance preparation | Major steps | Step chaining | Numeric fidelity | Overall |
|---|---|---|---|---|---|---|---|---|
| Biomni A1 | 0.774 ± 0.006 | 0.729 ± 0.008 | 0.760 ± 0.005 | 0.482 ± 0.004 | 0.514 ± 0.001 | 0.974 ± 0.004 | 0.067 ± 0.003 | 0.614 ± 0.002 |
| OpenAI (no tools) | 0.785 ± 0.007 | 0.739 ± 0.013 | 0.782 ± 0.006 | 0.488 ± 0.009 | 0.518 ± 0.004 | 0.981 ± 0.004 | 0.108 ± 0.002 | 0.629 ± 0.004 |
| Claude (no tools) | 0.728 ± 0.003 | 0.737 ± 0.007 | **0.831 ± 0.017** | 0.575 ± 0.004 | 0.561 ± 0.002 | 0.801 ± 0.015 | **0.211 ± 0.005** | 0.635 ± 0.002 |
| iBioGenie(no retrieval) | 0.759 ± 0.012 | 0.710 ± 0.006 | 0.748 ± 0.013 | 0.475 ± 0.012 | 0.496 ± 0.006 | 0.981 ± 0.004 | 0.104 ± 0.006 | 0.610 ± 0.005 |
| iBioGenie(retrieval only) | 0.764 ± 0.008 | 0.731 ± 0.003 | 0.742 ± 0.018 | 0.483 ± 0.004 | 0.511 ± 0.003 | 0.974 ± 0.004 | 0.108 ± 0.005 | 0.616 ± 0.004 |
| iBioGenie | 0.787 ± 0.011 | 0.752 ± 0.009 | 0.767 ± 0.015 | 0.563 ± 0.010 | 0.556 ± 0.005 | **0.988 ± 0.004** | 0.146 ± 0.008 | 0.651 ± 0.004 |
| Oracle | **0.790 ± 0.012** | **0.755 ± 0.011** | 0.776 ± 0.003 | **0.584 ± 0.008** | **0.578 ± 0.002** | 0.986 ± 0.007 | 0.160 ± 0.009 | **0.661 ± 0.002** |

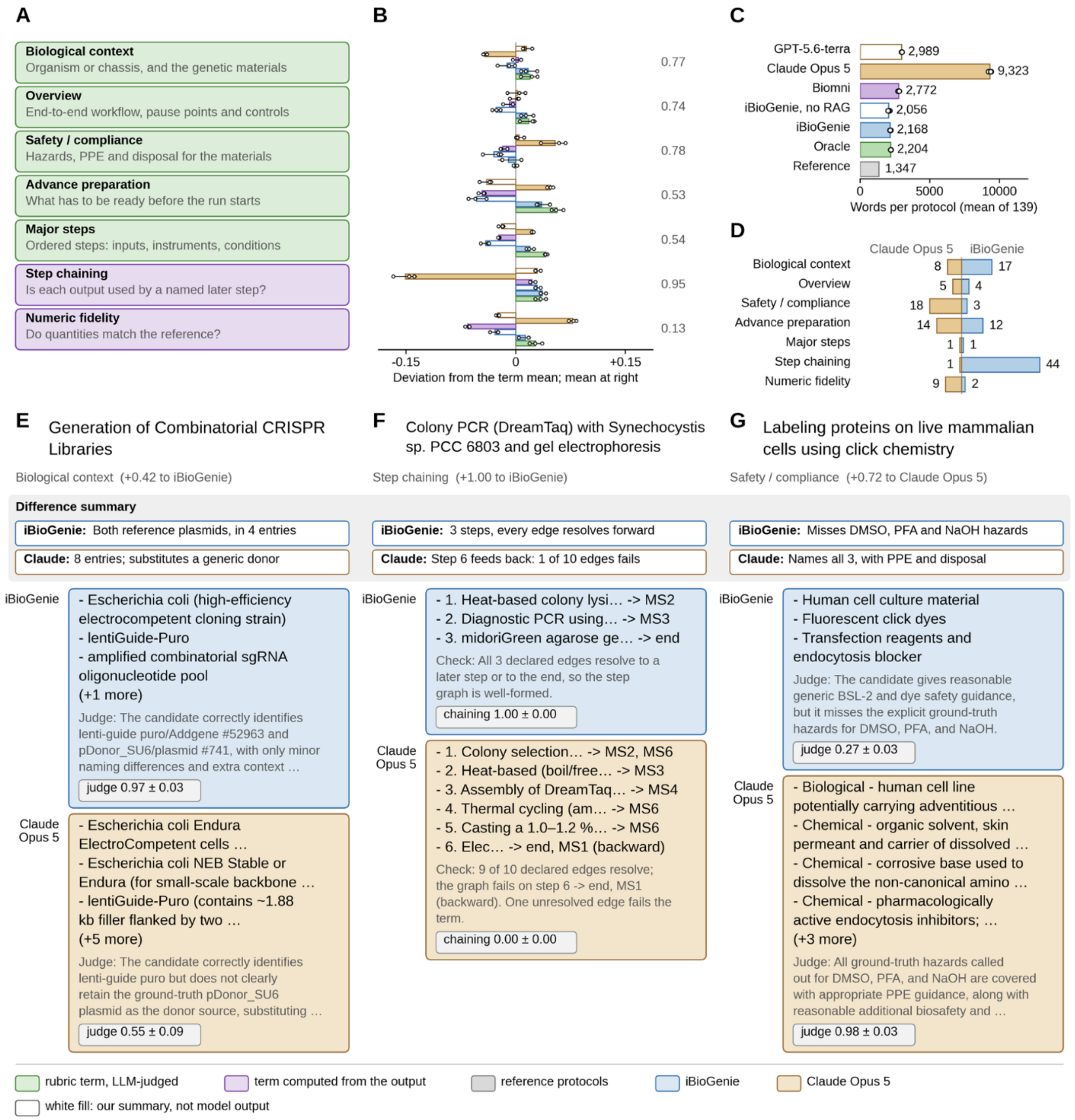


**Fig. S18. Per-term performance breakdown on the internal wet-lab protocol generation benchmark.** Six systems over all 139 protocols, three generations each; green marks judge-rated terms, purple computed terms, blue iBioGenie and brown Claude Opus 5. **(A)** The score is the mean of seven equal terms: five rubric sections rated by an LLM judge (gpt-5.4) and two computed from the generated JSON. **(B)** Each system's deviation from the term mean (printed at right); bars are the mean of three generations, error bars one SD. The two leading systems reach similar totals by different routes. **(C)** Output length: Claude Opus 5 writes 9,323 words per protocol, 6.9 times the 1,347-word reference mean and three to four times any other system. **(D)** The term that decides each protocol between iBioGenie and Claude Opus 5: step chaining decides 45 and favors iBioGenie 44 to 1, safety and compliance decides 21 and favors Claude 18 to 3, and advance preparation splits 12 to 14. **(E to G)** One cleanly separated protocol per block in D, with each system's output, the judge's rationale, and the term score (mean ± SD). **(E)** Biological context, 0.97 vs 0.55: iBioGenie names both reference plasmids; Claude substitutes a generic donor. **(F)** Step chaining, 1.00 vs

0.00: one backward edge among ten fails Claude. **(G)** Safety and compliance, 0.27 vs 0.98: Claude names the DMSO, PFA and NaOH hazards with PPE and disposal; iBioGenie gives generic BSL-2 guidance.

### 5. BiofoundryAI

#### 5.1. BiofoundryAI architecture and the biofoundry automation action space

BiofoundryAI is the physical AI layer of iCloudBiofoundry. It converts wet-lab protocols produced by iBioGenie into validated, simulation-tested workflows for execution on a biofoundry platform. Protocol translation and workflow generation are distributed among four specialized agents in a multi-agent architecture called FORGE (Foundry Orchestration Generation Engine). FORGE follows a single operating principle: a language model proposes a structured, typed description of an experiment, and a deterministic compiler verifies that description and renders it into platform-specific workflows, separating scientific judgment from the mechanical generation of device code. The inspection and biofoundry agents interact with the digital twin (**Section 5.3**) to close the loop, returning machine-generated diagnostics whenever a workflow fails to import, validate, or simulate (**fig. S19**). On the iCloudBiofoundry platform, general users use only the compiler to render workflows before submitting them for review. Administrators then use script agents to develop device methods (**Section 5.4**) and interact more extensively with the biofoundry through the full set of 77 CLI tools supported by the biofoundry agent (**Section 5.5**).

The planner agent operates on a site-specific context compiled from the biofoundry platform. This context includes exact device and operation contracts, plate templates, method registries, shared hotel positions, and nest inventory. These site-specific contexts constrain the plan and define what the compiler accepts. As a result, adding a contracted operation, plate type, or registered method is primarily a change to the site context rather than the agent (**Section 5.5**).

The planner agent writes one automation plan, which is the master contract for downstream generation. The plan contains an ordered set of typed Momentum steps, with the device, operation, plate identifiers, method parameters, and explicit units required for each step. It also records the plate inventory and the derived labware plan, including each plate's template, container type, barcode, lid state, and assigned shared-hotel position. For a structured plan, the script agent invokes a deterministic compiler. The compiler validates each planned step against the selected site's device-operation contract, binds labware and method parameters, assembles the Momentum process, and validates the rendered workflow before import. On this contract path, a language model does not write or patch raw Momentum workflow text. If a validation or review issue remains, the planner revises the typed plan, and the script agent recompiles it.

Some protocols cover discontinuous blocks of automation, such as an overnight incubation followed by a transformation the next morning. Such protocols are split into modules, each representing one contiguous block. If the protocol declares its own module, phase, or stage, that split is used as written; otherwise, the planner decides the split. Plates that pass from one module to the next are represented as explicit handoff objects that carry the container type, template, barcode, lid state, and the source and target modules and positions, and inconsistent handoffs are reported as warnings. A protocol that forms a single contiguous block runs as a one-module experiment. When there are several modules, per-module planning, script assembly, file writing, and review run concurrently for parallel generation.

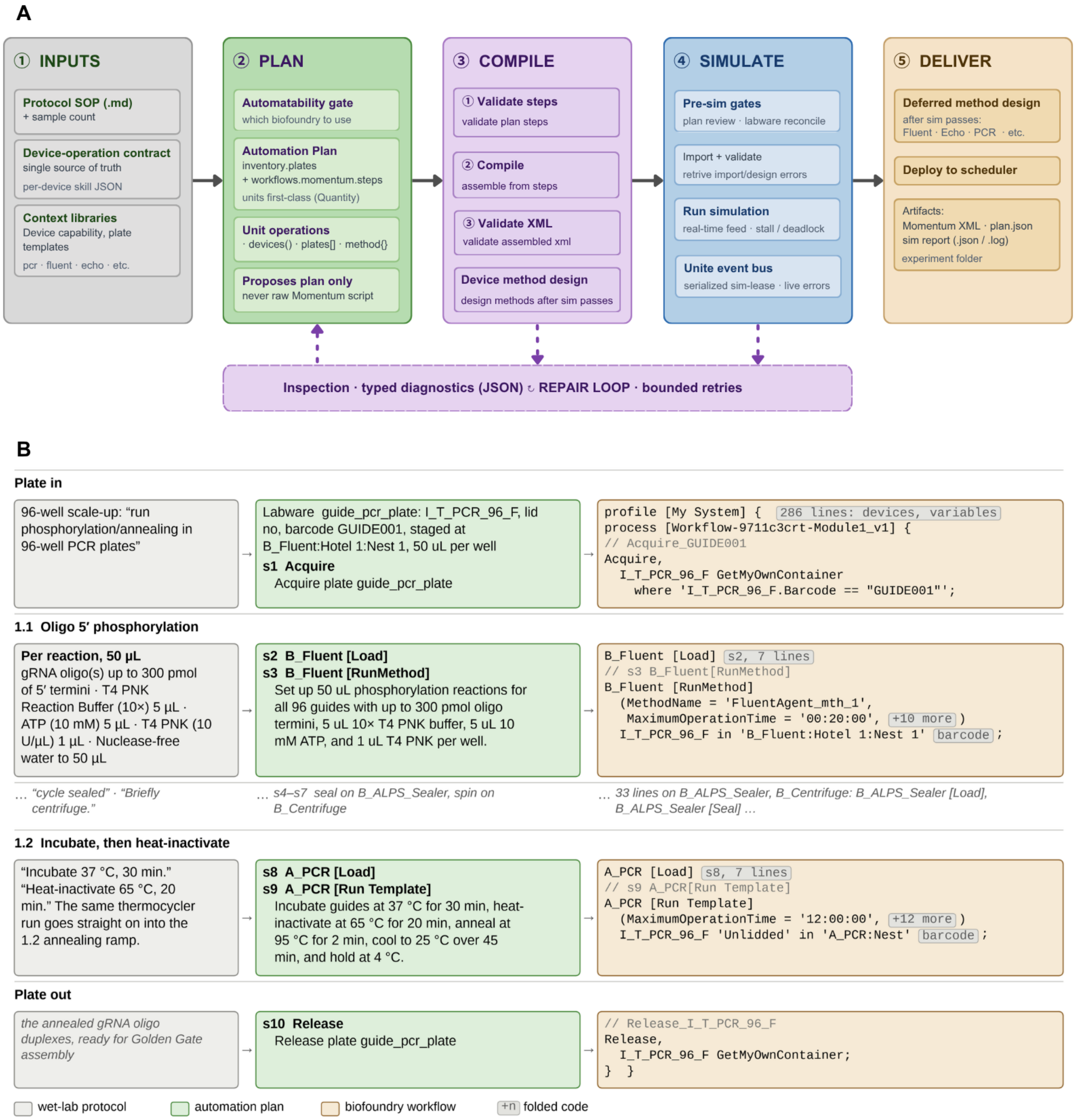


**Fig. S19. FORGE workflow overview. (A)** The FORGE workflow runs in five stages. (1) Inputs: the protocol SOP, the device-operation contract and context libraries. (2) Plan: the planner agent applies the automatability gate and proposes a typed automation plan of plates and ordered steps. (3) Compile: The script agent assembles and validates the workflow using a compiler, and further designs the device methods after simulation. (4) Simulate: the biofoundry agent runs pre-simulation gates and simulates the workflow, streaming live errors through the Unite event bus. (5) Deliver: instrument methods are designed, the workflow is deployed to the scheduler and the artifacts are saved. The inspection agent returns typed diagnostics from stages 3-4 to a repair loop with bounded retries. **(B)** Worked example of 96-well gRNA oligo 5′-phosphorylation and annealing, traced from the wet-lab protocol (grey) through the automation plan (green) to the Momentum workflow (brown).

#### 5.2. BiofoundryAI ensures reliable execution through layered validation and structured repair

Every generated workflow passes through three layers of checking before it can reach physical hardware: deterministic plan and script validation, review by the Inspection Agent, and import, validation, and simulation in Momentum. A failure at any gate is returned with evidence to the structured repair loop, which revises the design and renders the module again, up to a budget of five attempts. If the attempt keeps failing for the same reason, the repair loop will stop earlier.

The first layer is purely deterministic and uses no language model. It confirms that every device named in the workflow exists in the biofoundry, that each operation is one the named device can perform, that every container type matches a known plate template, that every method name appears in the instrument library, and that all numeric parameters fall within the device's physical limits, so that a centrifuge spin is never emitted outside the velocity the contract allows. These checks catch the failures language models produce, hard errors send the attempt back for repair without contacting the platform.

The second layer is a review by the inspection agent, which reads the complete workflow alongside the labware plan and the device-contract rules and looks for inconsistencies that pattern matching cannot identify, such as a centrifugation step without the counterweight its contract requires, or a container type that disagrees with what a downstream operation expects. When it finds a problem, it does not rewrite the whole workflow; it emits a precise instruction to rewrite the plan. The cost of repair is therefore proportional to the number of errors rather than to the size of the workflow.

The third layer is the automation platform itself. The workflow is imported into the digital twin of the platform, which checks device availability, inventory position conflicts, and the logic of the workflow as a whole. Anything it rejects is routed back to the inspection agent for another round of targeted repair. Once a workflow passes import and validation, the system registers each plate from the labware plan at its assigned position in the platform's inventory. The simulation exercises the entire workflow with simulated timing, moving plates between devices, and processing the work queue as it would during a physical run. This surfaces problems that static checking cannot, such as step conflicts between devices or race conditions in parallel branches. A simulation failure re-enters the same repair loop, and any failed unit of work is aborted automatically. If errors remain after every retry, the run reports failure and returns the last set of errors together with the last workflow it produced, rather than deploying anything. This end-to-end simulation is the simplified digital twin of the biofoundry, and **Section 3.4** describes what it exercises and how it reports.

The repair loop's effect is visible across the runs. Of the 1,710 modules run between 14 August and 5 September 2026, 0.905 passed simulation on the first attempt and 0.950 by the fifth, so 78 modules reached a simulated pass that the first attempt had not (**fig. S20A**). Those 78 are 0.48 of the 163 modules whose first attempt failed; 85 never passed, and each further attempt returns a smaller share of the modules still failing (**fig. S20B**). Two rounds show what one pass through the loop looks like: a Momentum import rejection whose next attempt imported (**fig. S20C**), and a simulation stall whose next attempt passed (**fig. S20D**). Because every retry re-plans and re-renders the whole module, these figures measure what the loop recovers, not the effect of any single instruction.

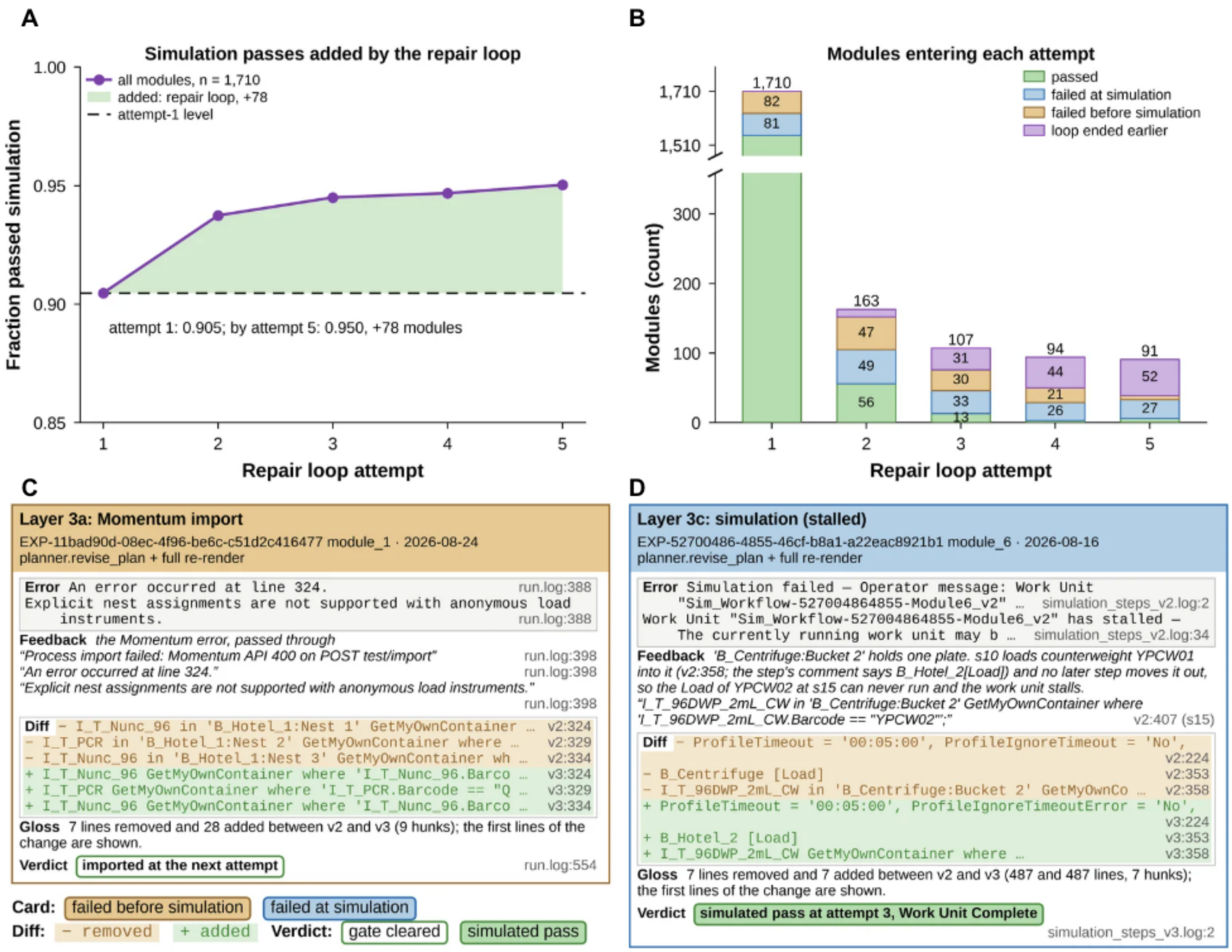


**Fig. S20. Repair-loop convergence of Momentum simulation. (A)** Cumulative fraction of the 1,710 modules run between 14 August and 5 September 2026 that had passed simulation by repair-loop attempt n. **(B)** The same modules as a sequence. Attempt 1 takes all 1,710; the modules still failing after attempt n enter attempt n + 1, and each bar shows what became of them there: passed (green), failed during simulation (blue), failed before simulation (brown), or loop ended earlier (purple). **(C)** A Momentum import rejection occurred due to an incorrect nest assignment, but the error was fixed in the next attempt and successfully imported. **(D)** A simulation stalled due to the occupancy of the device nest, but the next attempt fixed the error and the simulation passed. Each card quotes the gate's error, the instruction that went back, the first changed lines and the verdict, with the file and line of each.

#### 5.3. BiofoundryAI validates every workflow against a digital twin of the biofoundry

The third validation layer uses Momentum's Simulate mode as a digital twin of the biofoundry. The twin reproduces the site's configured scheduler, devices and operations, labware locations, inventory, and shared work queue without moving physical plates (**fig. S21**). BiofoundryAI can therefore test dynamic execution, not only whether a workflow is syntactically valid. In particular, the simulation asks whether Momentum can acquire the planned labware, reserve the required nests, route plates between devices, and complete the operation sequence without a scheduling or resource conflict. These behaviors cannot be established by static inspection alone.

BiofoundryAI validates workflows through three sequential gates (**fig. S21**). The inspection agent first imports the exact workflow into the simulation endpoint (Gate 1) and records whether Momentum accepts it or returns an import fault. It then queries Momentum for design errors associated with that process (Gate

2). Only an accepted process with no reported design error advances to dynamic simulation (Gate 3). This ordering separates workflow-format or profile errors from failures that arise only when the scheduler and devices execute the process.

Before each run, the biofoundry agent clears the twin's inventory, verifies that it is empty, and registers the planned labware at each plate's assigned nest. The agent submits the workflow as a single work unit of one iteration and monitors it, draining the unit's events and polling the work queue. As the twin executes, its events are rendered as a live step feed in which each entry records, where the platform reports it, the operation, the device, the plate or barcode, the nests involved, and the resulting state, so that an operator can watch the run unfold. The feed is kept as an ordered record together with a short report of the outcome, the number of recorded steps, and the labware-loading events, and it is written to the experiment folder alongside the run log.

The digital twin detects failure the moment it appears, from several signals: an operation that enters an error state or a work unit that enters an error, aborted, or paused state, a blocked queue, and event categories for a missing plate, a scheduler fault, or a nest conflict. Acquisition failures are recognized specifically, and the missing plate's container type and requested barcode are read out of the message, so the repair stages exactly the plate that is absent. A workflow that stops making progress is caught as a deadlock rather than being left to resume on its own. On any failure the digital twin's work unit is aborted and its plates are released from every nest, returning the deck to a clean state, and the corrected workflow re-enters the same targeted repair loop used by the other layers.

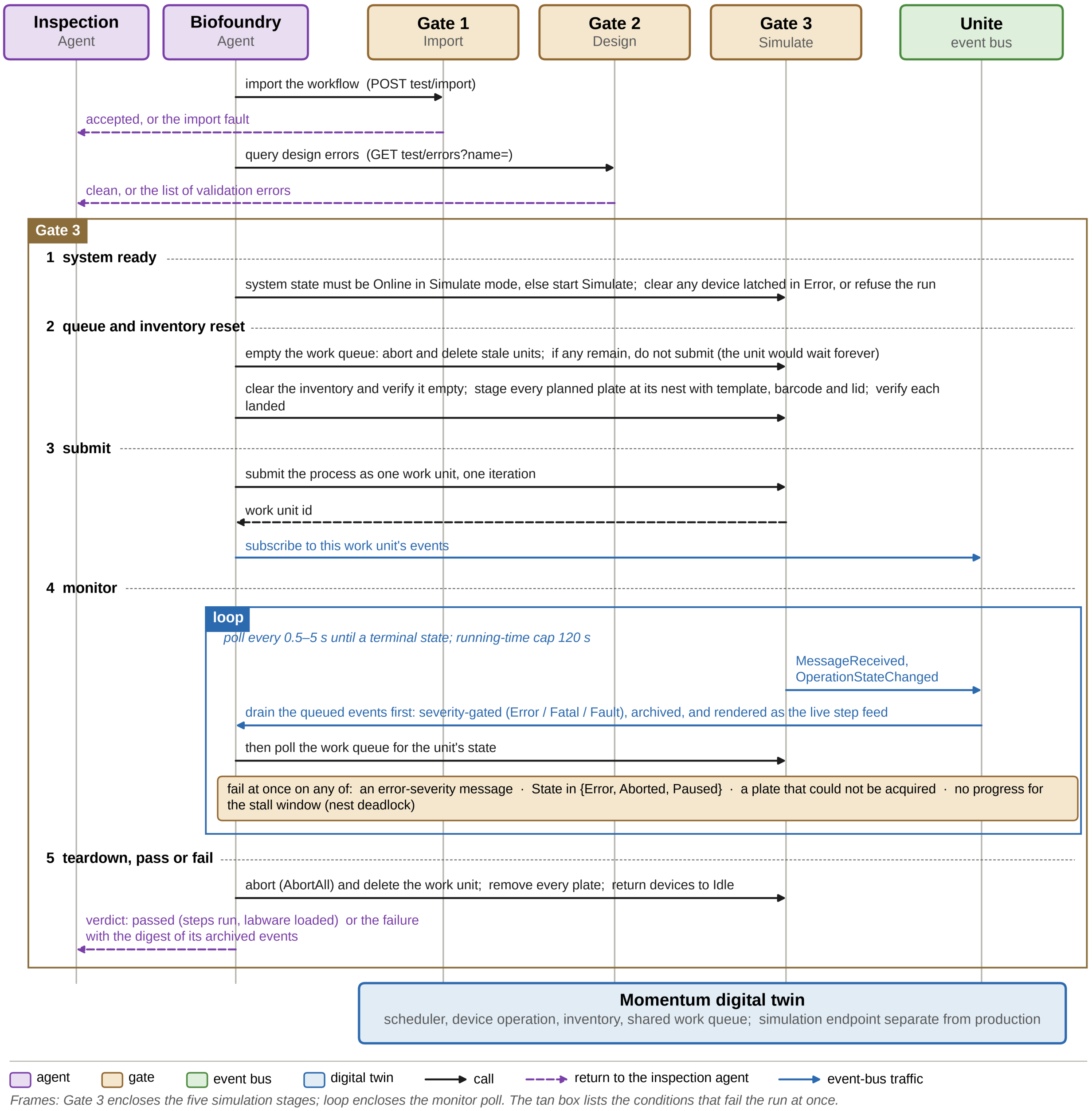


**Fig. S21. Momentum digital-twin validation and event flow.** The biofoundry agent runs three gates against the digital twin, which is Momentum in Simulate mode. Gate 1 (Import): the workflow is imported, and acceptance or the import fault is returned to the inspection agent. Gate 2 (Design): design errors are queried, and a clean result or the list of validation errors is returned. Gate 3 (Simulate) has five stages. (1) System ready: the system must be Online in Simulate mode and any device latched in Error is cleared, or the run is refused. (2) Queue and inventory reset: stale work units are aborted and deleted, and nothing is submitted if any remain; the inventory is cleared and verified empty, and every planned plate is staged at its nest with its template, barcode and lid and verified in place. (3) Submit: the process is submitted as one work unit of one iteration, and the agent subscribes to that unit's events on the Unite event bus. (4) Monitor: every 0.5-5s, the agent drains queued events and polls the unit's state until the unit terminates or reaches the 120s cap. (5) Teardown: whatever the outcome, the unit is aborted and deleted, all plates are removed and devices return to Idle. The verdict (passed, with the steps run and labware loaded, or the failure with a digest of its archived events) is returned to the inspection agent. Solid black arrows, calls; dashed purple arrows, returns to the inspection agent; blue arrows, event-bus traffic.

#### 5.4. BiofoundryAI generates instrument methods through deterministic and reasoning-based composition

Beyond the platform workflow, many experiments require method files in the native format of a particular instrument. These files are typically generated by the default software supplied with each device. BiofoundryAI not only uses the existing methods in the library but also uses the script agents to generate these files (**fig. S22A)**. The generated workflow contains the placeholder for all the methods that need to be designed. Once the workflow has been validated, the script agent will work on the design section for each instrument the experiment uses, and the files are rendered from those sections. Worklists for the Tecan Fluent liquid handler require reasoning about the deck and are produced by a dedicated multi-agent system (**fig. S22B)**; the acoustic picklist, the thermocycler program, and the plate-reader method are rendered deterministically.

The Fluent worklist is produced by FluentAgent. A multi-channel liquid handler is programmed against its deck: the generator must place labware on the worktable, pick up and discard tips, and keep the arms clear of one another. FluentAgent runs as an ordered sequence of specialized agents. A planner converts the standard operating procedure into a structured protocol of six sections that fix the deck layout, the arm assigned to each operation, the liquid classes, the ordered operations themselves, the constraints on transitions of the arm state machine, and the safety checks. A collision validator then examines the planned placements. The multi-channel head can collide with tall labware placed in an adjacent deck position. The validator detects this case by computing the physical clearance directly in code, and when a collision would occur, it relocates the labware to a non-adjacent position and updates the protocol accordingly. A Coder writes a Python program against the site's pyFluent library (*8*), and an Executor runs the program in a bounded subprocess to produce a GWL file without controlling the robot. Deterministic preflight checks and a Reviewer validate the program and GWL; categorized feedback is returned to the Coder for another bounded attempt when needed.

The Labcyte Echo picklist is rendered from the experiment design's acoustic-dispensing section. Each CSV row maps a source plate and well to a destination plate and well and records the transfer volume in nanoliters. The builder validates plate indices, well syntax, transfer volumes, and duplicate destinations before writing the file.

Thermocycler instructions can be written as a platform-independent PCR program table or, for Analytik Jena Biometra instruments, as a native TSuite text method. The PCR builder records run-wide settings such as lid temperature and block type, followed by one record per thermal step. Temperatures and ramp rates are encoded as integers scaled by 100, and a loop containing N total passes stores N -1 repeats in the native format.

The plate-reader builder writes a Tecan MDFX measurement file for the Infinite M1000 Pro. It supports absorbance, fluorescence, luminescence, emission scans, and excitation scans, together with optional temperature control and kinetic blocks. Wavelength, gain, plate range, and timing settings are taken from the instrument section of the experiment design.

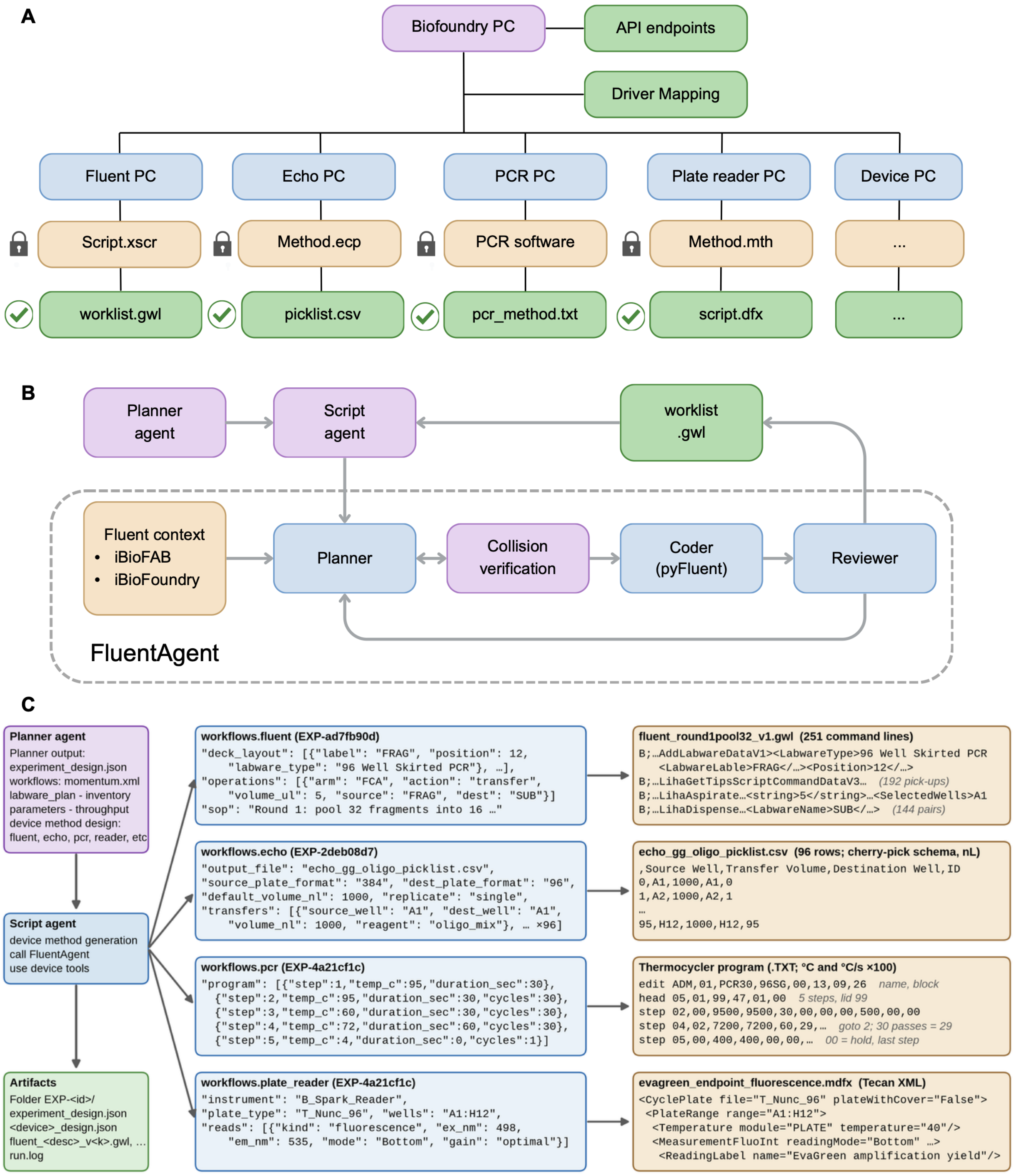


**Fig. S22. Instrument methods generation for biofoundry. (A)** Example layout for method generation across the biofoundry. The Biofoundry PC coordinates the workcell through its API endpoints and a driver mapping that links each scheduled operation to the PC controlling its instrument (Fluent, Echo, PCR, plate reader etc.). On each instrument PC, the native method format (Script.xscr, Method.ecp, the PCR vendor software, Method.mth) is locked by the vender. BiofoundryAI instead generates an open, importable file that the instrument software executes (check mark): a Fluent worklist (.gwl), an Echo pick list (.csv), a thermocycler program (.txt) and a plate-reader script (.dfx). **(B)** Architecture of the FluentAgent. The planner agent's structured intent passes through the script agent to the Fluent agent. There, a planner draws on Fluent context from iBioFAB and iBioFoundry, its deck plan is checked by collision verification, a coder renders it with pyFluent, and a reviewer either sends it back to the planner for revision or releases the worklist (.gwl) to the script agent. **(C)** End-to-end example for different method generation. The planner agent writes experiment design. The script agent turns each device block into a native method by calling FluentAgent or device-specific tools. Left to right: the structured device specification and an excerpt of the method it produces.

All outputs are collected in a versioned experiment folder. The folder contains the master experiment design, the wet-lab to automation translation, the validated Momentum workflow, per-instrument design records, the generated instrument files required by the experiment, a run log, and the simulation event record. Multi-module experiments also include a module manifest and per-module outputs.

Deployment is an explicit operation after validation and method design. On sites with separate simulation and production endpoints, the biofoundry agent imports the validated workflow into production Momentum but does not start it or stage production labware. The deployment backend then uploads the generated method files and transfers them to the configured device computers, where file type and upload order map them to the numbered placeholder slots referenced by the workflow (**section 5.5**). The operator receives the required labware placements and starts physical execution only after completing the physical setup.

#### 5.5. BiofoundryAI scales across compatible distributed biofoundries

BiofoundryAI separates shared workflow logic from site-specific context. The FORGE framework and its agents are reused across sites. Each site supplies a compiled context of its Momentum devices, operations, plate templates, and inventory, together with the credentials and endpoints needed to reach the platform.

The site context is built by a context-engineering workflow (**fig. S23A**). Devices, plate templates, and nest inventory are read from the site's Momentum database through the Momentum web-services API. The operations each device supports, and their descriptions, are extracted from the Momentum driver documentation and from the site's existing process scripts, then supplemented with curated device capabilities, operation contracts, and plate definitions. The driver documentation (from Momentum 7.3.2) contains 177 device drivers and 761 operations across 31 instrument categories supported by Momentum (**fig. S23B**), demonstrating the scale that BiofoundryAI can support and expand upon.

Contexts are currently provided for iBioFoundry (Momentum 7.3.2) and iBioFAB (Momentum 7.3.3), which differ in scale and instrument complement. iBioFoundry has 60 devices supporting 199 operations (**fig. S23C**), and iBioFAB has 48 devices supporting 204 operations (**fig. S23D**). The two sites also carry different device type, such as centrifuge and plate reader, yet the same agents plan, generate, validate, and deploy on both, because site selection, simulation, and deployment are resolved from the compiled context and the endpoint configuration.

All interactions with the physical biofoundry go through Momentum that handles device scheduling, plate tracking, and workflow execution. The biofoundry agent reaches Momentum through a direct client for its web-services API, and instrument method files are delivered to the device computers through a separate BiofoundryAI backend service. BiofoundryAI defines 77 platform tools (**table S7**): 67 Momentum tools, seven tools for the deployment backend, and three for session files.

The cost of adding a platform is therefore bounded mainly by describing its context and configuring its endpoints. A site is compatible if it is scheduled by Momentum and its instruments, plate templates, and nest positions can be described in this form. The site must expose the Momentum web-services API.

Connecting its Unite event bus adds real-time feedback during simulation, and installing the deployment backend allows method files to be delivered to individual instruments (**table S7**).

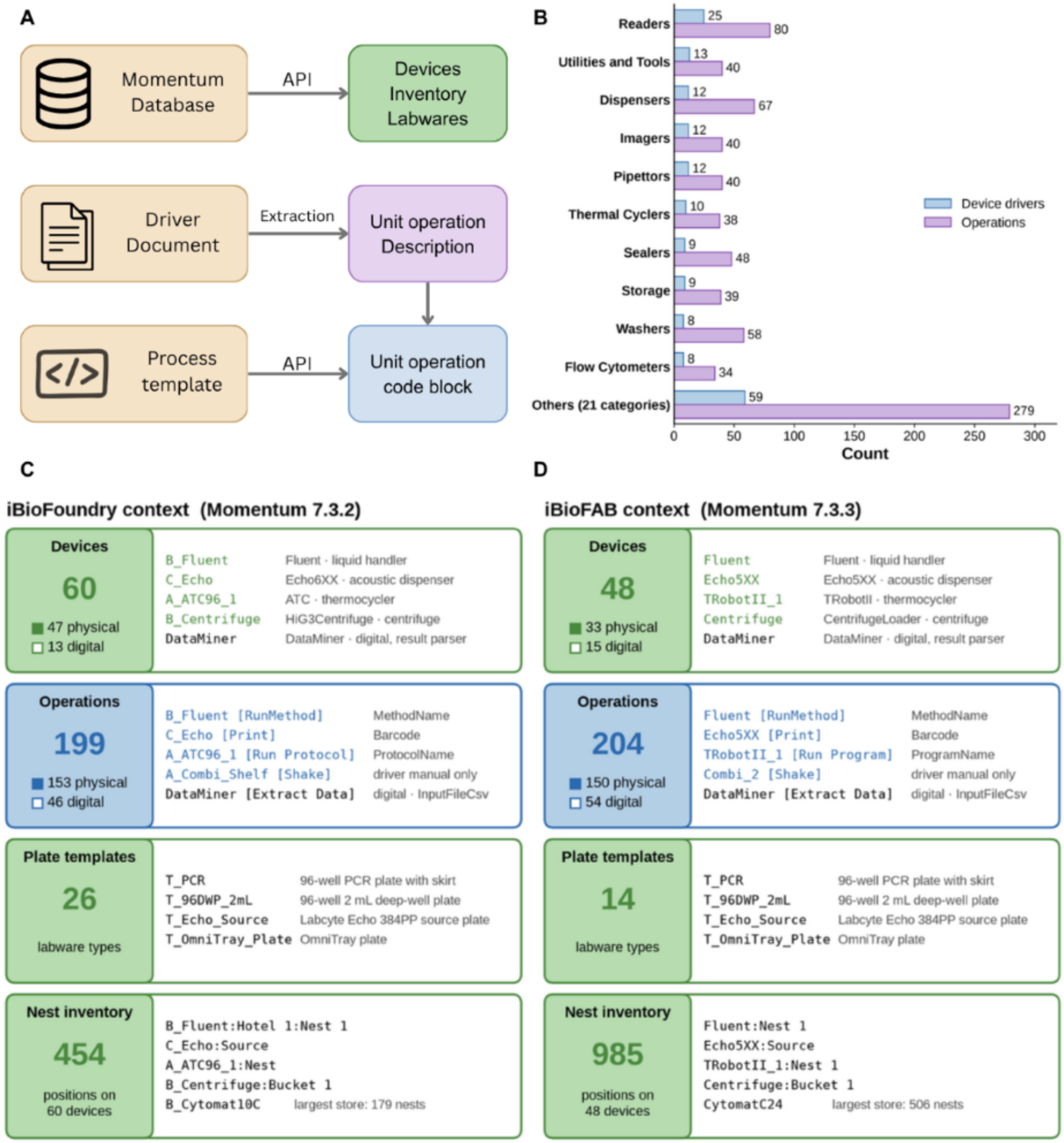


**Fig. S23. Biofoundry context engineering overview. (A)** Biofoundry context engineering workflow. **(B)** Devices and operations that Momentum (version 7.3.2) supports. Top 10 device categories were shown in the figure. **(C)** The iBioFoundry context and **(D)** the iBioFAB context: the number of devices, operations, plate templates, and nests, with examples of each. A physical device (filled square) owns at least one nest and holds labware; a digital device (hollow square) owns none.

**Table S7. Biofoundry Agent CLI tools for biofoundry interaction and cloud deployment**

| Group | CLI tools number | What it covers |
|---|---|---|
| System control | 5 | status, version, start/stop, attended mode |
| Device discovery and control | 13 | devices, nests, device tasks, device state/availability/presence, waste |
| Inventory management | 15 | containers, inventory items, templates, add/remove plates, snapshots |
| Process development | 4 | import, export, delete a process, read its validation errors |
| Process execution | 6 | list processes, variables, run a process or worklist, experiments, runtime variables |
| Work queue control | 15 | pause/resume/abort the queue; inspect, load, unload, abort, delete work units and batches |
| Operator messages | 3 | simple message, confirmation dialog, message XML |
| Simulation | 6 | full simulate, instrument state, simulation queue, reset instrument, import and validate, run evidence |
| Session files | 3 | read, list, clean up saved large responses |
| Backend deployment | 7 | create/status/delete experiment, upload files, transfer to devices, deploy folder, health |
| Total | 77 | 67 Momentum platform tools + 3 session-file tools + 7 backend tools |

#### 5.6. BiofoundryAI supports continuous improvement from operational experience

BiofoundryAI includes a human-gated learning subsystem that turns operational experience into improvements on request. It works in four stages: capture, diagnose and propose, evaluate, and approve and promote (**fig. S24A**). Capture is automatic. Every run on BiofoundryAI is recorded to a local store with its plan, each script version, its validation and simulation results, the number of repair attempts it required and its outcome. An edit to a plan or a script, and the difference between two successive repair attempts, are stored as corrections. Each run is labelled a success, a partial success or a failure. A failure opens a case to investigate, and a clean success becomes a hold-out case that later changes must not break.

On request, the subsystem clusters the chosen failures by root cause and names the kind of fix each cluster needs. It then drafts a candidate fix aimed only at that weakness: a lesson, a reusable module, or a revised prompt or a tool fix. It regenerates the failed cases and the hold-out cases twice, without and with the candidate fix, in a sandbox whose changes never persist, and scores both by deterministic structural checks; the measured effect is the change in the rate of valid scripts. A candidate that turns any previously passing case into a failure is flagged as a regression. An administrator reviews each candidate fix with its measured effect and its regression flag, and approves, rejects or later rolls it back. An approved prompt becomes a new version with earlier versions kept, an approved lesson is appended to the site's lessons file as a block that can be removed on its own, and an approved module is added to the site's module library. The agents read the change on the next run, which is captured in turn, and this closes the loop.

The system improved over the same period through a simpler path, in which the software detects and records its failures, while people apply the fixes. Across 3,046 modules run between 1 June and 5 September 2026, the fraction passing all three gates rose from 0.18 in the first batch of 50 experiments to 0.81 in the last (**fig. S24B**). On 24 and 25 June 2026 the benchmark harness ran the same 24 cases through import, the design check, and simulation three times, and 2, 10 and then 23 of them passed (**fig. S24C**). After each run it wrote a lesson for every error that recurred in at least two cases, in total there are six lessons in three blocks (L1-L3), on import time-outs, rejected imports, unregistered acoustic-dispenser protocol names, missing picklists and out-of-range temperatures. Two examples trace the progression from an error to the change that resolved it (**fig. S24D and E**). For L2, a script post-processor left a dangling comma that caused Momentum to reject the import; this was corrected in the orchestrator. For L3, a 37 °C step was sent to an incubator with a limited temperature range of 4-25 °C, a planner routing rule was added, and the validator's ranges were corrected.

In practice, BiofoundryAI improves with a human in the loop. The system captures every run and records the errors that recur, and a developer decides what changes. Each run adds to the collected data, so the system improves as it is used.

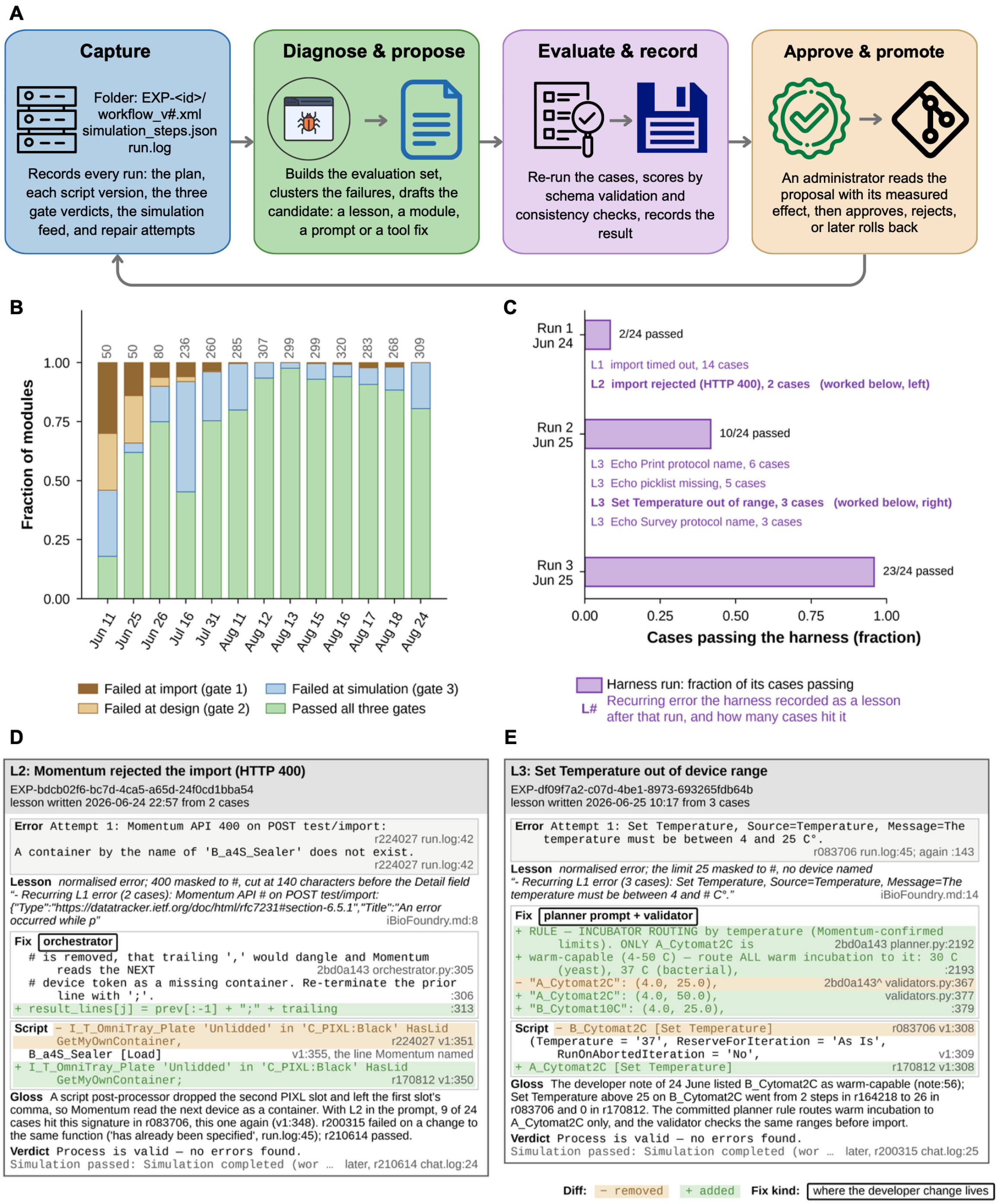


**Fig. S24. BiofoundryAI improves with operational experience.** Operational simulation outcomes across sequential experiment batches. **(A)** The four stages of the learning subsystem. **(B)** All 3,046 modules run between 1 June and 5 September 2026, in sequential batches of 50 experiments labelled by the first date in the batch (modules per batch above each bar). **(C)** Three successive benchmark harness runs on the same 24 cases, 24-25 June 2026, as the fraction of cases passing. Under each bar, in purple, are the recurring errors the harness wrote to the lessons file after that run (L1-L3). **(D, E)** Two of those lessons traced from the error to the change that resolved it. L2: Momentum rejected the import (HTTP 400). L3: a Set Temperature step outside the incubator's range. Every quoted line carries its source file and line, or its commit.

### 6. BiofoundryAI benchmarking

#### 6.1. Benchmark dataset construction

To compare BiofoundryAI with general-purpose large language models (LLMs), we constructed a benchmark of 40 cases on the iBioFoundry platform, spanning four categories of increasing complexity (**table S8**).

Operations comprise 8 single-instrument tasks, including an OD600 read on the plate reader, a plate move between storage positions, a thermocycler run and an acoustic transfer on the Echo. The 12 modules cover curated platform workflows such as HiFi assembly, transformation, colony picking, and protein-expression induction. The 8 automation SOPs are multi-step standard operating procedures, including Illumina DNA library preparation, Echo-based 384-well qPCR setup and a dose-response series. The 12 wet-lab protocols represent the complete experiments, including PCR variants, ligation, Golden Gate and Loop assembly, and CRISPR/Cas9 genome editing.

**Table S8. BiofoundryAI benchmark dataset statistics.**

| Category | Cases | Avg steps | Min | Max | Sum |
|---|---|---|---|---|---|
| Operations | 8 | 9.4 | 3 | 16 | 75 |
| Modules | 12 | 31.3 | 11 | 67 | 376 |
| Automation-SOP | 8 | 35 | 19 | 50 | 280 |
| Wet-lab protocols | 12 | 36.8 | 8 | 87 | 442 |
| Total | 40 | 29.3 | 3 | 87 | 1173 |

Each case pairs a wet-lab protocol with a Level-1 automation plan that translates the protocol into instrument-level steps and a Level-2 Momentum workflow script that implements the plan. Automation experts wrote and reviewed both references, checking every device, operation, container type and biological parameter against the platform. All 40 reference scripts were validated on the Momentum simulator. Simulation defects, including missing lid qualifiers, nest collisions and missing plate declarations, were corrected. The full set then ran to completion in three consecutive simulator passes.

#### 6.2. Benchmark design

**Level-1: Protocol Translation Task**

In the protocol translation task, each system converts a wet-lab protocol into an automation plan. Every LLM receives one system prompt and one user message. The system prompt begins with a fixed task instruction: "You are a laboratory-automation engineer for the iBioFoundry biofoundry. Translate the wet-lab protocol below into an instrument-level automation plan for this platform, using only the devices, operations, labware and methods documented in the reference material that follows."

The prompt then provides a frozen platform context. Its seven blocks cover devices and capabilities, the device-operation contract, labware and plate templates, the liquid-handler worktable, shared hotel positions, registered methods and protocols, and plan-level flow control. The user message contains the benchmark protocol as input. The model returns a Markdown plan with labware assignments, workflow, and parameters sections.

**Level-2: Workflow Generation Task**
In the workflow generation task, each system converts an automation plan into a Momentum process. The system prompt begins with a fixed task instruction. "You are a laboratory-automation engineer for the iBioFoundry biofoundry. Implement the instrument-level automation plan below as Momentum process statements, using only the devices, operations, parameters, nests and container types documented in the reference material that follows."

The prompt then provides a frozen context covering the Momentum language, the device catalog, device skill details and registered methods. The model returns the Momentum statements inside the process body, covering labware acquisition, device operations, and control flow.

To separate planning errors from script generation errors, Level-2 is run on two tracks: GT-plan track, the system receives the curated ground truth plan, isolating script generation from planning. Own-plan track, the system receives the plan it produced at Level-1 by the model it used. This is the realistic end-to-end result that from wet-lab protocol to automation plan then to the platform script, in which planning errors will also propagate.

We have evaluated the performance of BiofoundryAI against other four LLMs (**table S9**). All four LLMs receive the prompts described above through one shared harness. The prompts are built once from the frozen release and are identical for every model; only provider transport differs. Every baseline run recorded the same prompt digest (hashes 27027c91e7801ea1). The output limit is 16,000 tokens at Level-1 and 32,000 tokens at Level-2 for every model.

Both open-weight models were served with vLLM (0.22.0) on four NVIDIA RTX 6000 Ada Generation GPUs using tensor parallelism 4, bfloat16 weights, an 81,920-token context window and 0.92 GPU memory utilisation. Qwen used the qwen3 reasoning parser, while Gemma used the gemma4 parser with the Gemma-4 chat template. Thinking was enabled for Gemma-4-26B-A4B-it and disabled for Qwen3.6-35B-A3B. Sampling used temperature 1.0 and replicate seeds 1, 2 and 3. The hosted models ran with provider-default reasoning; their routes did not accept seeded sampling.

At Level-1, BiofoundryAI's planner agent (GPT-5.6 Terra) works from the same platform facts as the baselines. The shared reference is built from the context sources the planner loads. At Level-2, FORGE compiler converts the plan into structured intent and renders the script deterministically against the device-operation contract. We have run the BiofoundryAI with two set up. One is the one-shot design that only try for once, the other is the feedback design that allows the agent to fix the error through the repair loops with maximum five attempts.

**Table S9. Systems evaluated in the BiofoundryAI benchmark.**

| System | Type | Level-1 | Level-2 | Access and serving |
|---|---|---|---|---|
| BiofoundryAI | Proposed agent system | planner agent (GPT-5.6 Terra) | contract-driven compiler | Native pipeline |
| GPT-5.6 Terra (gpt-5.6-terra) | Closed-weight LLM | One call | One call | OpenAI API |
| Claude Opus 5 (claude-opus-5) | Closed-weight LLM | One call | One call | Anthropic API |
| Qwen3.6-35B-A3B | Open-weight mixture-of-experts LLM | One call | One call | vLLM, OpenAI-compatible |
| Gemma-4-26B-A4B-it | Open-weight mixture-of-experts LLM | One call | One call | vLLM, OpenAI-compatible |

**Evaluation Metrics**

To better compare the performance, we come up with a standard evaluation metrics for both Level-1 protocol translation (**table S10**) and Level-2 workflow generation (**table S11**). For protocol translation, the metrics covers the devices, operation, step order, labware and parameters. For workflow generation, similar metrics were applied to the script with more emphasis on the validity of the script that screen if it can pass the import, design, and simulation test via the digital-twin.

**Table S10. Level-1 protocol translation metrics.** The predicted plan is parsed into its workflow steps, as (device, operation) tuples, and its labware and parameter sections, and each is compared with the reference plan.

| Score | Definition |
|---|---|
| device | F1 between predicted and reference device sets. Similarity is 1.0 for identical IDs or the same model with another instance number in one functional category, 0.5 for another model in the same category, and 0 otherwise. |
| operation | F1 between operation sets. Similarity is 1.0 for identical operations, 0.5 for synonyms, and 0 otherwise. |
| step_lcs | Mean of two order measures over step tuples: a longest-common-subsequence ratio normalized by the longer sequence and F1 over adjacent step pairs. A step match uses the lower of its device and operation similarities; weights below 0.5 count as 0. |
| labware | Harmonic mean of container correctness and completeness. Correctness is F1 over container types, with 1.0 for identical types or documented aliases and 0.5 for the same physical plate format. Completeness credits a lid state and a valid container type. |
| parameters | F1 over biological parameters, extracted by class with normalized units. Recall uses reference parameters, while precision uses only predicted values in reference classes. |
| overall_score | Harmonic mean of the five scores, each floored at 0.05: $5 / \Sigma k\ 1 / \max(\text{score}, 0.05)$. |

The harmonic mean of the overall score reflects the conjunctive structure of a protocol: the devices, operation, order, labware and parameters must all be correct.

**Table S11. Level-2 workflow generation metrics.** Generated statements are parsed into ordered device-operation steps with parameters and compared with the reference script. Devices, operations, steps and parameters require exact matches.

| Score | Definition |
|---|---|
| device | F1 between the device set used by the script and the reference set. |
| operation | F1 between the operation sets. |
| gt_score | Mean of step F1, the longest-common-subsequence ratio over ordered tuples, and F1 over adjacent step pairs, all against the reference script and all using exact matches. |
| labware | F1 over script containers, with similarity 1.0 for identical containers or documented aliases and 0.5 for the same physical plate format. |
| parameter | Steps are aligned to the reference by longest common subsequence. The score is the fraction of reference parameters reproduced exactly on aligned steps; parameters on unaligned reference steps count as misses. |
| harm_5 | Harmonic mean of the five scores above, each floored at 0.05; this measures script fidelity before validity. |
| Gate 1 import | Momentum accepts the file and the import request returns HTTP 200. |
| Gate 2 design | The process imports and passes design validation without errors. |
| Gate 3 simulate | A submitted work unit reaches Work Unit Complete on the simulator. |
| validity | Per case, (gate 1 + gate 2 + gate 3) / 3. The gates are nested, so validity is 0, ⅓, ⅔ or 1. |
| overall_score | Per case, (validity + harm_5) / 2, averaged over cases. |

Validity averages the three nested gates rather than requiring all of them, so a script that imports but does not run can earn ⅓ or ⅔, since it will reduce the repair cost to let it pass the simulation test. The Level-2 overall_score averages validity and fidelity. It measures script quality together with progress through the validation gates.

For each system and replicate, the evaluation pipeline loads the per-case predictions, scores each case against the curated reference and averages the case scores. Level-1 uses the 40 cases common to all systems. Each system was generated and scored in three independent replicates. Tables report the mean across replicates ± sample standard deviation. Truncated, unparseable, refused or empty outputs remain in the mean and are scored as produced.

For Gate 3, each script was run on the iBioFoundry Momentum simulator with the same simulation tool used by the biofoundry agent. Before each validation run, the work queue was drained, inventory was cleared, devices were checked for error states, and a test import had to succeed and without design errors. Plate type, barcode and lid state came from the script's container declarations. Starting nests came from the Level-1 reference plan on the GT-plan track and the system's own Level-1 plan on the own-plan track.

#### 6.3. Benchmark results

All scores are reported as mean ± sample SD across three replicates (n = 3; **tables S12–S16**).

**Table S12. Level-1 protocol translation.**

| System | device | operation | step_lcs | labware | parameters | overall_score |
|---|---|---|---|---|---|---|
| BiofoundryAI (feedback) | 0.816 ± 0.013 | **0.855 ± 0.010** | 0.670 ± 0.020 | **0.820 ± 0.011** | 0.778 ± 0.016 | 0.746 ± 0.013 |
| BiofoundryAI (one-shot) | **0.820 ± 0.011** | 0.853 ± 0.011 | **0.681 ± 0.018** | 0.818 ± 0.015 | 0.771 ± 0.009 | **0.748 ± 0.009** |
| claude-opus-5 | 0.766 ± 0.015 | 0.792 ± 0.013 | 0.493 ± 0.008 | 0.773 ± 0.008 | 0.787 ± 0.018 | 0.676 ± 0.014 |
| gpt-5.6-terra | 0.772 ± 0.010 | 0.795 ± 0.016 | 0.441 ± 0.008 | 0.766 ± 0.020 | **0.801 ± 0.008** | 0.624 ± 0.012 |
| gemma-4-26B | 0.723 ± 0.019 | 0.741 ± 0.022 | 0.367 ± 0.008 | 0.756 ± 0.007 | 0.639 ± 0.020 | 0.527 ± 0.023 |
| Qwen3.6-35B | 0.726 ± 0.002 | 0.747 ± 0.015 | 0.307 ± 0.010 | 0.677 ± 0.011 | 0.756 ± 0.007 | 0.531 ± 0.020 |

BiofoundryAI produced the best automation plans in one-shot, with overall_score 0.748 compared with 0.676 for the strongest LLM, Claude Opus 5. While BiofoundryAI with feedback loops is enabled, it will be based on simulation feedback to adjust the automation plan. This will lead to a lower score compared to the reference, but it allows for a higher success rate in designing the script and passing the simulation. The largest difference is step order: BiofoundryAI's step_lcs is 0.681 compared with 0.307-0.493 for the LLMs. The systems are closer on the other four scores, differing by at most 0.143. The LLMs usually select appropriate instruments, operations, labware and quantities, but order the steps less faithfully.

**Table S13. Level-2 workflow generation GT-plan track script fidelity.**

| System | device | Operation | gt_score | labware | parameter | harm_5 |
|---|---|---|---|---|---|---|
| BiofoundryAI (feedback) | 0.931 ± 0.006 | 0.963 ± 0.004 | 0.886 ± 0.009 | 0.918 ± 0.008 | 0.871 ± 0.007 | 0.906 ± 0.006 |
| BiofoundryAI (one-shot) | 0.991 ± 0.000 | **0.976 ± 0.000** | **0.959 ± 0.000** | **0.990 ± 0.000** | **0.929 ± 0.000** | **0.967 ± 0.000** |
| claude-opus-5 | **0.999 ± 0.001** | 0.971 ± 0.001 | 0.926 ± 0.009 | 0.982 ± 0.008 | 0.917 ± 0.007 | 0.954 ± 0.007 |
| gpt-5.6-terra | 0.848 ± 0.045 | 0.823 ± 0.044 | 0.734 ± 0.037 | 0.855 ± 0.034 | 0.720 ± 0.046 | 0.784 ± 0.030 |

| System | device | Operation | gt_score | labware | parameter | harm_5 |
|---|---|---|---|---|---|---|
| gemma-4-26B | 0.974 ± 0.001 | 0.955 ± 0.001 | 0.917 ± 0.005 | 0.963 ± 0.003 | 0.907 ± 0.004 | 0.942 ± 0.002 |
| Qwen3.6-35B | 0.991 ± 0.005 | 0.973 ± 0.002 | 0.913 ± 0.011 | 0.863 ± 0.088 | 0.907 ± 0.007 | 0.854 ± 0.067 |

**Table S14. Level-2 workflow generation GT-plan track gate pass rates and overall scores.**

| System | n | gate 1 import | gate 2 design | gate 3 simulate | validity | overall_score |
|---|---|---|---|---|---|---|
| BiofoundryAI (feedback) | 40 | 40 / 40 / 40 | 39 / 39 / 39 | 38 / 38 / 37 | **0.972 ± 0.005** | **0.939 ± 0.005** |
| BiofoundryAI (one-shot) | 40 | 37 / 37 / 37 | 35 / 35 / 35 | 34 / 34 / 34 | 0.883 ± 0.000 | 0.925 ± 0.000 |
| claude-opus-5 | 40 | 38 / 38 / 36 | 7 / 10 / 8 | 4 / 4 / 2 | 0.408 ± 0.025 | 0.681 ± 0.009 |
| gpt-5.6-terra | 40 | 29 / 29 / 30 | 27 / 28 / 29 | 12 / 10 / 11 | 0.569 ± 0.013 | 0.677 ± 0.011 |
| gemma-4-26B | 40 | 29 / 27 / 24 | 24 / 21 / 20 | 10 / 9 / 10 | 0.483 ± 0.038 | 0.712 ± 0.018 |
| Qwen3.6-35B | 40 | 35 / 28 / 25 | 27 / 20 / 19 | 14 / 6 / 6 | 0.500 ± 0.117 | 0.678 ± 0.088 |

Level-2 GT-plan track. Given the same reference plan, BiofoundryAI, Claude Opus 5, Gemma-4-26B-A4B-it, and achieve relative high script fidelity, with harm_5 values of 0.967, 0.954, and 0.942. The different is whether the script can be run on the digital twin. With script template and grammar in the context, LLMs can write script with high fidelity, however, still would fail on the execution. While BiofoundryAI (one-shot) only use the deterministic compiler (SD = 0) can translate the script most similar to the ground truth but still 6 cases failed in validity test. With feedback repair loops enabled, the validity increase from 0.883 to 0.972, while the script fidelity drops from 0.967 to 0.906, since it would required to change the ground truth plan to make the script valid. The systems differ mainly in whether Momentum accepts and runs the scripts. BiofoundryAI's validity is 0.972 compared with 0.408-0.569 for the LLMs, producing overall_scores of 0.939 and 0.677-0.712.

**Table S15. Level-2 workflow generation own-plan track script fidelity.**

| System | device | operation | gt_score | labware | parameter | harm_5 |
|---|---|---|---|---|---|---|
| BiofoundryAI (feedback) | 0.738 ± 0.028 | **0.828 ± 0.020** | 0.673 ± 0.022 | **0.747 ± 0.026** | 0.688 ± 0.028 | 0.706 ± 0.024 |
| BiofoundryAI (one-shot) | **0.748 ± 0.028** | 0.826 ± 0.026 | **0.692 ± 0.021** | 0.740 ± 0.017 | **0.705 ± 0.020** | **0.714 ± 0.019** |
| claude-opus-5 | 0.673 ± 0.013 | 0.773 ± 0.013 | 0.557 ± 0.011 | 0.685 ± 0.011 | 0.678 ± 0.014 | 0.645 ± 0.008 |
| gpt-5.6-terra | 0.653 ± 0.016 | 0.728 ± 0.001 | 0.469 ± 0.008 | 0.644 ± 0.025 | 0.493 ± 0.013 | 0.548 ± 0.008 |
| gemma-4-26B | 0.690 ± 0.012 | 0.759 ± 0.017 | 0.467 ± 0.009 | 0.641 ± 0.012 | 0.480 ± 0.004 | 0.545 ± 0.009 |
| Qwen3.6-35B | 0.639 ± 0.005 | 0.752 ± 0.025 | 0.437 ± 0.020 | 0.388 ± 0.026 | 0.521 ± 0.020 | 0.424 ± 0.014 |

**Table S16. Level-2 workflow generation own-plan track gate pass rates and overall scores.**

| System | n | gate 1 import | gate 2 design | gate 3 simulate | validity | overall_score |
|---|---|---|---|---|---|---|
| BiofoundryAI (feedback) | 40 | 39 / 39 / 38 | 39 / 39 / 38 | 38 / 38 / 37 | **0.958 ± 0.014** | **0.832 ± 0.018** |
| BiofoundryAI (one-shot) | 40 | 39 / 39 / 38 | 30 / 33 / 30 | 30 / 30 / 29 | 0.828 ± 0.021 | 0.771 ± 0.018 |
| claude-opus-5 | 40 | 35 / 32 / 30 | 4 / 2 / 3 | 2 / 2 / 3 | 0.314 ± 0.024 | 0.480 ± 0.014 |
| gpt-5.6-terra | 40 | 24 / 30 / 29 | 20 / 27 / 27 | 10 / 9 / 13 | 0.525 ± 0.066 | 0.536 ± 0.035 |

| System | n | gate 1 import | gate 2 design | gate 3 simulate | validity | overall_score |
|---|---|---|---|---|---|---|
| gemma-4-26B | 40 | 17 / 17 / 16 | 12 / 13 / 12 | 6 / 5 / 5 | 0.286 ± 0.010 | 0.415 ± 0.009 |
| Qwen3.6-35B | 40 | 14 / 9 / 11 | 8 / 7 / 8 | 1 / 1 / 1 | 0.167 ± 0.025 | 0.296 ± 0.019 |

Level-2 own-plan track. When each system scripts its own plan, fidelity falls because Level-1 errors carry into the script. BiofoundryAI's overall_score is 0.832 compared with 0.296-0.536 for the LLMs. For one-shot workflow generation of the BiofoundryAI achieves 0.828 validity, while the feedback loops help to improve the validity to 0.958. Scores are reported by category for both tracks (**fig. S25**).

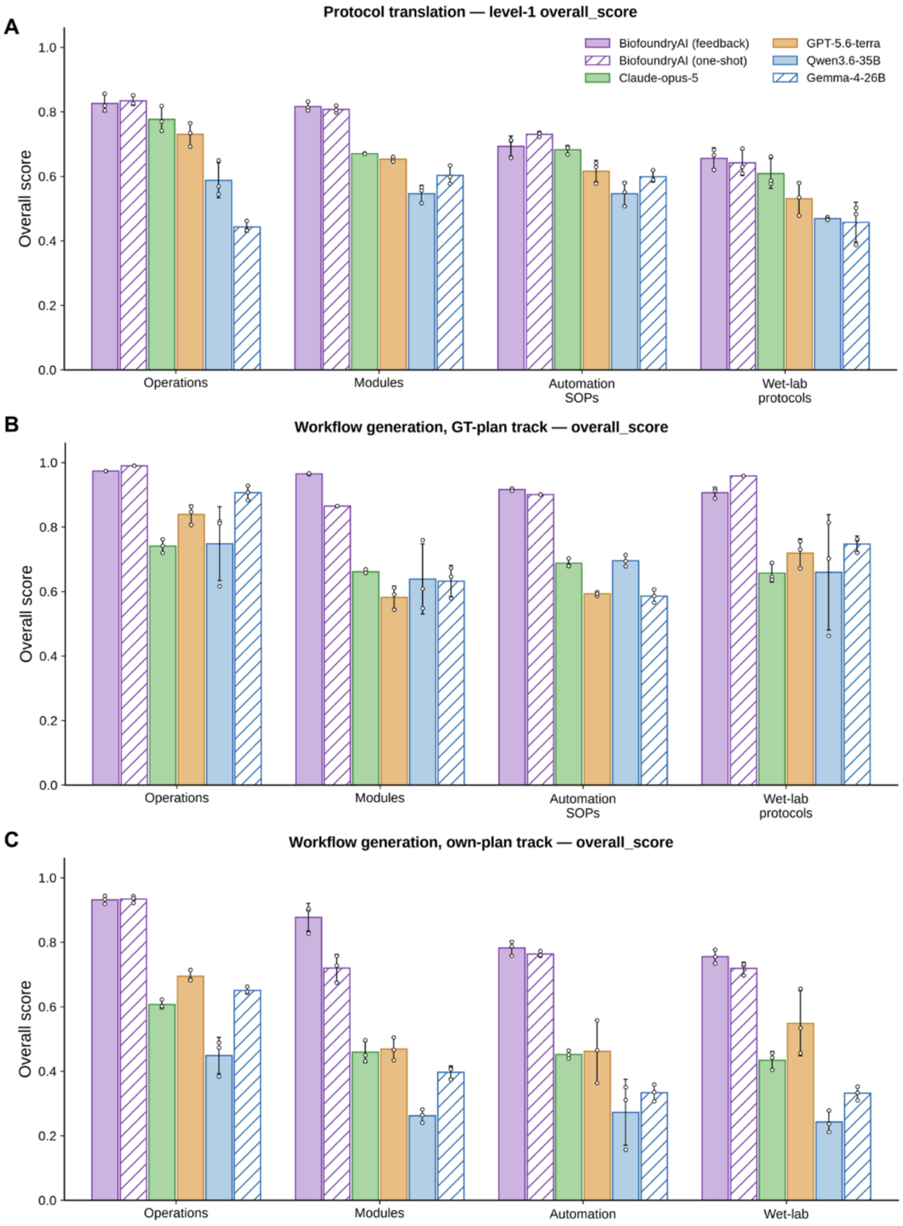


**Fig. S25. Detailed performance on the four categories of the protocols. (A)** overall score for Level-1 protocol translation in four categories. **(B)** overall score for Level-2 GT-plan track in four categories. **(C)** overall score for Level-2 own-plan track in four categories.

### 7. Case study I: Phytase engineering

#### 7.1. Agentic AI pipeline for phytase engineering

To further engineer phytase for improved activity, we used iBioGenie to extend the prior four-round campaign into a fifth round of directed evolution (*9*). The system was first prompted to design new mutations for *Ym*Phytase by using the experimentally measured data from previous campaign (4 rounds) rather than starting from sequence information alone. iBioGenie began by extracting target-specific information, including the *Ym*Phytase protein record from UniProt and the corresponding predicted structure from AlphaFold DB. This provided the target identity, sequence context, and structural framework needed to guide downstream model training and mutation prioritization.

Next, iBioGenie organized the supervised model-training workflow. The previous four rounds of *Ym*Phytase activity measurements were processed into a training dataset, allowing each variant to be represented by its mutation composition and measured functional output. To capture both sequence- and structure-level information, the workflow incorporated sequence features derived from ESM2 and structural features derived from ProteinMPNN. These features were then used to train Kermut, a supervised regression model designed to learn from the accumulated sequence–function data and predict the activity of new higher-order mutants. In this way, the fifth-round design process was not only based on general protein-sequence priors, but also on the specific experimental trajectory observed during the previous *Ym*Phytase engineering campaign.

After model training, iBioGenie generated a fifth-round candidate library containing possible 5$^{th}$ and 6$^{th}$ order mutant designs. These candidates represented new mutation combinations built from the existing engineering background and were intended to explore sequence space beyond the variants already tested in the first four rounds. The candidate library was then ranked using greedy optimization, prioritizing mutants predicted to provide the largest functional improvement while remaining experimentally actionable for site-directed mutagenesis. From this ranked library, iBioGenie selected 90 top candidate mutations for experimental validation (**fig. S26**).

Finally, iBioGenie generated the experimental design outputs needed to move from computational prediction to laboratory construction. For each selected mutation, the system produced site-directed mutagenesis primer designs using Primer3, yielding 90 SDM primer pairs corresponding to the selected fifth-round mutants. These deliverables converted the model-ranked mutation list into an experimentally executable library, enabling direct construction and testing of the predicted *Ym*Phytase variants (**table S17**). Of the 90 proposed variants, 73 were successfully constructed, tested, and yielded retrievable activity measurements.

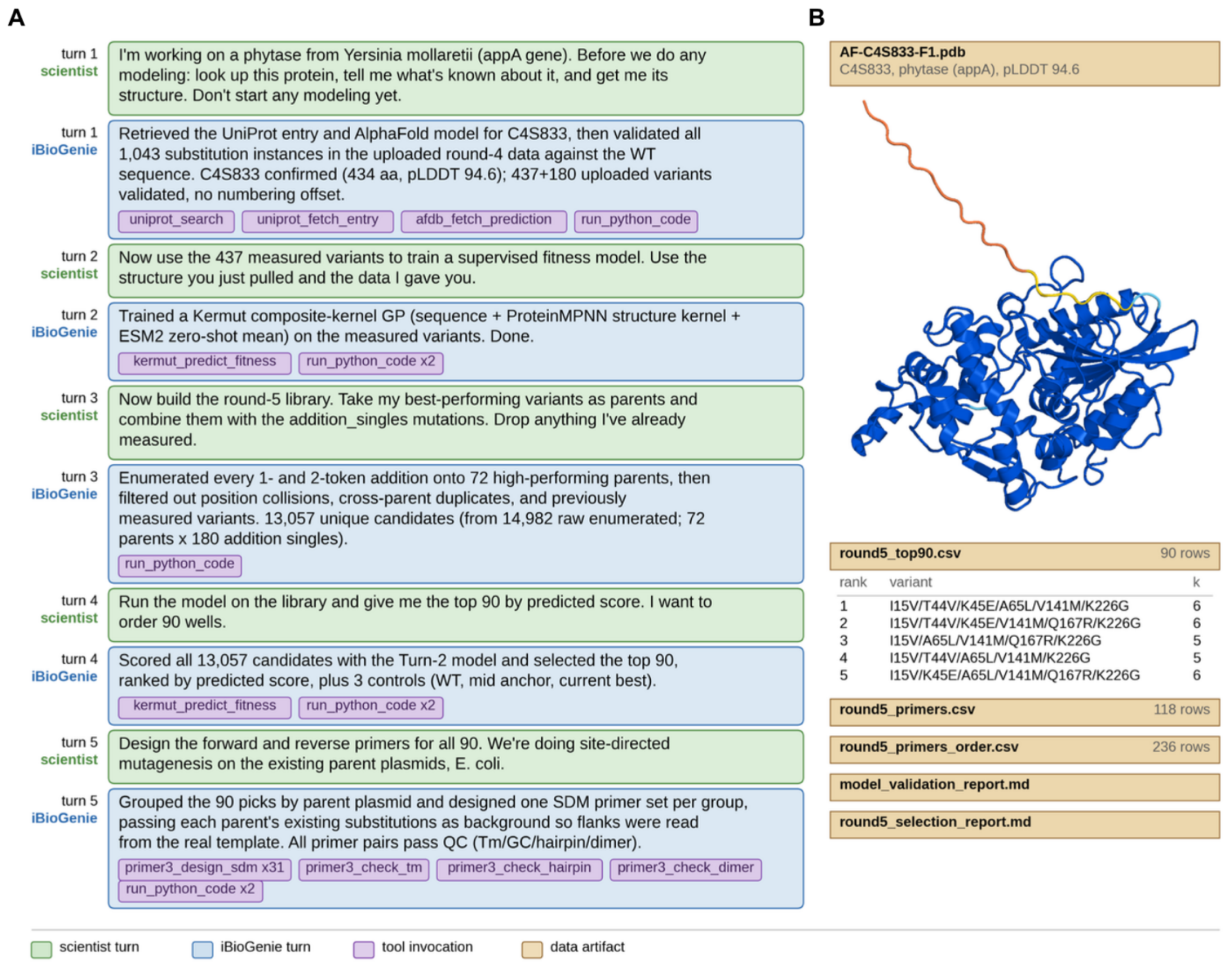


**Fig. S26. iBioGenie design for phytase engineering using prior experimental data. (A)** A summary of the scientist guided a data-driven design workflow starting from the *Ym*phytase sequence and structure. A supervised fitness model was trained on previously measured variants, then used to score higher-order candidates generated from high-performing parents and previously tested single mutations. The top 90 predictions were selected with three controls, followed by SDM primer design and quality checks. **(B)** Representative outputs include the AppA structure, the ranked 90-variant panel, primer designs and ordering files, model-validation results, and the final selection report.

**Table S17. SDM design and primers for *Ym*Phytase.**

| Well | Combined Mutant | Parent | Forward (5' to 3') | Reverse (5' to 3') |
|---|---|---|---|---|
| A1 | I15V/T44V/K45E/A65L/V141M/K226G | I15V/A65L/V141M/K226G | GTTCGTTCGCCGGTCGAACAAACAGAGTTAATGAATG | CATTCATTAACTCTGTTTGTTCGACCGGCGAACGAAC |
| A2 | I15V/T44V/K45E/V141M/Q167R/K226G | I15V/V141M/Q167R/K226G | GTTCGTTCGCCGGTCGAACAAACAGAGTTAATGAATG | CATTCATTAACTCTGTTTGTTCGACCGGCGAACGAAC |
| A3 | I15V/T25D/T44V/K45E/V141M/K226G | I15V/T25D/V141M/K226G | GTTCGTTCGCCGGTCGAACAAACAGAGTTAATGAATG | CATTCATTAACTCTGTTTGTTCGACCGGCGAACGAAC |

| Well | Combined Mutant | Parent | Forward (5' to 3') | Reverse (5' to 3') |
|---|---|---|---|---|
| A4 | I15V/T44V/K45E/V141M/K226G / Q362R | I15V/V141M/K226G/Q362R | GTTCGTTCGCCGGTCGAACAAACAGAGTTAATGAATG | CATTCATTAACTCTGTTTGTTCGACCGGCGAACGAAC |
| A5 | I15V/T44V/K45E/V141M/K226G/V266R | I15V/V141M/K226G/V266R | GTTCGTTCGCCGGTCGAACAAACAGAGTTAATGAATG | CATTCATTAACTCTGTTTGTTCGACCGGCGAACGAAC |
| A6 | I15V/T44V/K45E/V141M/K226G/L303P | I15V/V141M/K226G/L303P | GTTCGTTCGCCGGTCGAACAAACAGAGTTAATGAATG | CATTCATTAACTCTGTTTGTTCGACCGGCGAACGAAC |
| A7 | I15V/T44V/K45E/V141M/K226G /V298L | I15V/V141M/K226G/V298L | GTTCGTTCGCCGGTCGAACAAACAGAGTTAATGAATG | CATTCATTAACTCTGTTTGTTCGACCGGCGAACGAAC |
| A8 | I15V/T44V/K45E/A65L/V141M/Q362R | I15V/A65L/V141M/Q362R | GTTCGTTCGCCGGTCGAACAAACAGAGTTAATGAATG | CATTCATTAACTCTGTTTGTTCGACCGGCGAACGAAC |
| A9 | I15V/T44V/K45E/V141M/K226G/Q295D | I15V/V141M/K226G/Q295D | GTTCGTTCGCCGGTCGAACAAACAGAGTTAATGAATG | CATTCATTAACTCTGTTTGTTCGACCGGCGAACGAAC |
| A10 | I15V/T44V/K45E/V141M/H161Q/K226G | I15V/V141M/H161Q/K226G | GTTCGTTCGCCGGTCGAACAAACAGAGTTAATGAATG | CATTCATTAACTCTGTTTGTTCGACCGGCGAACGAAC |
| A11 | I15V/T44V/K45E/V141M/K226G/L303G | I15V/V141M/K226G/L303G | GTTCGTTCGCCGGTCGAACAAACAGAGTTAATGAATG | CATTCATTAACTCTGTTTGTTCGACCGGCGAACGAAC |
| A12 | I15V/T44V/K45E/E48Q/V141M/K226G | I15V/E48Q/V141M/K226G | GTTCGTTCGCCGGTCGAACAAACAGAGTTAATGAATG | CATTCATTAACTCTGTTTGTTCGACCGGCGAACGAAC |
| B1 | I15V/T44V/K45E/V141M/Q154D/K226G | I15V/V141M/Q154D/K226G | GTTCGTTCGCCGGTCGAACAAACAGAGTTAATGAATG | CATTCATTAACTCTGTTTGTTCGACCGGCGAACGAAC |
| B2 | I15V/T44V/K45E/L103V/V141M / K226G | I15V/L103V/V141M/K226G | GTTCGTTCGCCGGTCGAACAAACAGAGTTAATGAATG | CATTCATTAACTCTGTTTGTTCGACCGGCGAACGAAC |
| B3 | I15V/T44V/K45E/V141M/K226G/V298S | I15V/V141M/K226G/V298S | GTTCGTTCGCCGGTCGAACAAACAGAGTTAATGAATG | CATTCATTAACTCTGTTTGTTCGACCGGCGAACGAAC |
| B4 | I15V/P19A/T44V/K45E/V141M/K226G | I15V/P19A/V141M/K226G | GTTCGTTCGCCGGTCGAACAAACAGAGTTAATGAATG | CATTCATTAACTCTGTTTGTTCGACCGGCGAACGAAC |
| R4_A2 | V141M/K226G/I15V/A65L | — | | |

| Well | Combined Mutant | Parent | Forward (5' to 3') | Reverse (5' to 3') |
|---|---|---|---|---|
| B6 | I15V/T44V/K45E/V141M/K226G/G334A | I15V/V141M/K226G/G334A | GTTCGTTCGCCGGTCGAACAAACAGAGTTAATGAATG | CATTCATTAACTCTGTTTGTTCGACCGGCGAACGAAC |
| B7 | I15V/T25D/T44V/K45E/A65L/V141M | I15V/T25D/A65L/V141M | GTTCGTTCGCCGGTCGAACAAACAGAGTTAATGAATG | CATTCATTAACTCTGTTTGTTCGACCGGCGAACGAAC |
| R4_A3 | V141M/K226G/I15V/Q167R | — | | |
| B9 | I15V/T44V/K45E/Q106W/V141M /K226G | I15V/Q106W/V141M/K226G | GTTCGTTCGCCGGTCGAACAAACAGAGTTAATGAATG | CATTCATTAACTCTGTTTGTTCGACCGGCGAACGAAC |
| B10 | I15V/T44V/K45E/V141M/H161T/K226G | I15V/V141M/H161T/K226G | GTTCGTTCGCCGGTCGAACAAACAGAGTTAATGAATG | CATTCATTAACTCTGTTTGTTCGACCGGCGAACGAAC |
| B11 | I15V/T44V/K45E/V141M/T158E/K226G | I15V/V141M/T158E/K226G | GTTCGTTCGCCGGTCGAACAAACAGAGTTAATGAATG | CATTCATTAACTCTGTTTGTTCGACCGGCGAACGAAC |
| B12 | I15V/T44V/K45E/V141M/Q154S/K226G | I15V/V141M/Q154S/K226G | GTTCGTTCGCCGGTCGAACAAACAGAGTTAATGAATG | CATTCATTAACTCTGTTTGTTCGACCGGCGAACGAAC |
| C1 | I15V/V141M/K226G/Q295R/L303Q | I15V/V141M/K226G/Q295R | ACGGCGCTAGTGCAACAGCGTAAGGGG | CCCCTTACGCTGTTGCACTAGCGCCGT |
| C2 | I15V/T44V/K45E/V141M/K226G/R305Q | I15V/V141M/K226G/R305Q | GTTCGTTCGCCGGTCGAACAAACAGAGTTAATGAATG | CATTCATTAACTCTGTTTGTTCGACCGGCGAACGAAC |
| C3 | I15V/T44V/K45E/V141M/K226G/T318N | I15V/V141M/K226G/T318N | GTTCGTTCGCCGGTCGAACAAACAGAGTTAATGAATG | CATTCATTAACTCTGTTTGTTCGACCGGCGAACGAAC |
| C4 | I15V/T44V/K45E/V141M/K226G/G334S | I15V/V141M/K226G/G334S | GTTCGTTCGCCGGTCGAACAAACAGAGTTAATGAATG | CATTCATTAACTCTGTTTGTTCGACCGGCGAACGAAC |
| C5 | T44V/K45E/A65L/V141M /K226G/Q362R | A65L/V141M/K226G / Q362R | GTTCGTTCGCCGGTCGAACAAACAGAGTTAATGAATG | CATTCATTAACTCTGTTTGTTCGACCGGCGAACGAAC |
| C6 | I15V/T44V/K45E/V141M/K226G/K319R | I15V/V141M/K226G/K319R | GTTCGTTCGCCGGTCGAACAAACAGAGTTAATGAATG | CATTCATTAACTCTGTTTGTTCGACCGGCGAACGAAC |
| C7 | I15V/T44V/K45E/V141M/S157P/K226G | I15V/V141M/S157P/K226G | GTTCGTTCGCCGGTCGAACAAACAGAGTTAATGAATG | CATTCATTAACTCTGTTTGTTCGACCGGCGAACGAAC |
| C8 | I15V/T44V/K45E/V141M/ | I15V/V141M/S157K/K226G | GTTCGTTCGCCGGTCGAACAAACAGAGTTAATGAATG | CATTCATTAACTCTGTTTGTTCGACCGGCGAACGAAC |

| Well | Combined Mutant | Parent | Forward (5' to 3') | Reverse (5' to 3') |
|---|---|---|---|---|
| | S157K / K226G | | | |
| C9 | I15V/T44V/K45E/V141M/Q154A/K226G | I15V/V141M/Q154A/K226G | GTTCGTTCGCCGGTCGAACAAACAGAGTTAATGAATG | CATTCATTAACTCTGTTTGTTCGACCGGCGAACGAAC |
| C10 | T44V/K45E/V141M/Q167R/K226G/Q362R | V141M/Q167R/K226G/Q362R | GTTCGTTCGCCGGTCGAACAAACAGAGTTAATGAATG | CATTCATTAACTCTGTTTGTTCGACCGGCGAACGAAC |
| C11 | I15V/T44V/K45E/V141M/K226G/L303S | I15V/V141M/K226G/L303S | GTTCGTTCGCCGGTCGAACAAACAGAGTTAATGAATG | CATTCATTAACTCTGTTTGTTCGACCGGCGAACGAAC |
| R4_C2 | V141M/K226G/I15V/R305Q | — | | |
| D1 | I15V/T44V/K45E/V141M/K226G/V298D | I15V/V141M/K226G/V298D | GTTCGTTCGCCGGTCGAACAAACAGAGTTAATGAATG | CATTCATTAACTCTGTTTGTTCGACCGGCGAACGAAC |
| D2 | I15V/T44V/K45E/V141M/K226G/E316S | I15V/V141M/K226G/E316S | GTTCGTTCGCCGGTCGAACAAACAGAGTTAATGAATG | CATTCATTAACTCTGTTTGTTCGACCGGCGAACGAAC |
| D3 | I15V/T44V/K45E/E48R/V141M/K226G | I15V/E48R/V141M/K226G | GTTCGTTCGCCGGTCGAACAAACAGAGTTAATGAATG | CATTCATTAACTCTGTTTGTTCGACCGGCGAACGAAC |
| D4 | I15V/T44V/K45E/V141M/K226G/A339L | I15V/V141M/K226G/A339L | GTTCGTTCGCCGGTCGAACAAACAGAGTTAATGAATG | CATTCATTAACTCTGTTTGTTCGACCGGCGAACGAAC |
| D5 | I15V/T44V/K45E/V141M/S173A/K226G | I15V/V141M/S173A/K226G | GTTCGTTCGCCGGTCGAACAAACAGAGTTAATGAATG | CATTCATTAACTCTGTTTGTTCGACCGGCGAACGAAC |
| D6 | I15V/T44V/K45E/V141M/K226G/E316P | I15V/V141M/K226G/E316P | GTTCGTTCGCCGGTCGAACAAACAGAGTTAATGAATG | CATTCATTAACTCTGTTTGTTCGACCGGCGAACGAAC |
| R4_D1 | V141M/K226G/I15V/E316P | — | | |
| D8 | I15V/T44V/K45E/V141M/Q154T/K226G | I15V/V141M/Q154T/K226G | GTTCGTTCGCCGGTCGAACAAACAGAGTTAATGAATG | CATTCATTAACTCTGTTTGTTCGACCGGCGAACGAAC |
| D9 | T25D/T44V/K45E/V141M/Q167R/K226G | T25D/V141M/Q167R / K226G | GTTCGTTCGCCGGTCGAACAAACAGAGTTAATGAATG | CATTCATTAACTCTGTTTGTTCGACCGGCGAACGAAC |
| D10 | I15V/T44V/K45E/V141M/H161R / K226G | I15V/V141M/H161R/K226G | GTTCGTTCGCCGGTCGAACAAACAGAGTTAATGAATG | CATTCATTAACTCTGTTTGTTCGACCGGCGAACGAAC |

| Well | Combined Mutant | Parent | Forward (5' to 3') | Reverse (5' to 3') |
|---|---|---|---|---|
| D11 | T25D/T44V/K45E/A65L/V141M/K226G | T25D/A65L/V141M/K226G | GTTCGTTCGCCGGTCGAACAAACAGAGTTAATGAATG | CATTCATTAACTCTGTTTGTTCGACCGGCGAACGAAC |
| D12 | I15V/V141M/Q154T/K226G/E316K | I15V/V141M/K226G/E316K | GCCGGTGTGTGTACGCTAGATTCGACA | TGTCGAATCTAGCGTACACACACCGGC |
| E1 | A65L/V141M/K226G/V302G / L303P | A65L/V141M/K226G / L303P | GTGACGGCGCTAGGGCTTCAGCGTAAG | CTTACGCTGAAGCCCTAGCGCCGTCAC |
| E2 | I15V/A65L/V141M/Q167R / K226G | I15V/A65L/V141M/K226G | GCCATTGAGGCGCGACTGGGGGCGCCA | TGGCGCCCCCAGTCGCGCCTCAATGGC |
| E3 | T44V/K45E/A65L/V141M/K226G / L303P | A65L/V141M/K226G / L303P | GTTCGTTCGCCGGTCGAACAAACAGAGTTAATGAATG | CATTCATTAACTCTGTTTGTTCGACCGGCGAACGAAC |
| E4 | T44V/K45E/V141M/K226G/Q295D/L303P | V141M/K226G/Q295D/L303P | GTTCGTTCGCCGGTCGAACAAACAGAGTTAATGAATG | CATTCATTAACTCTGTTTGTTCGACCGGCGAACGAAC |
| E5 | I15V/T44V/K45E/E48D/V141M/K226G | I15V/E48D/V141M/K226G | GTTCGTTCGCCGGTCGAACAAACAGAGTTAATGAATG | CATTCATTAACTCTGTTTGTTCGACCGGCGAACGAAC |
| E6 | G7A/V141M/Q167R/K226G/Q362R | V141M/Q167R/K226G/Q362R | TTAACTGCACTGGCCTTAATGCTAGGT | ACCTAGCATTAAGGCCAGTGCAGTTAA |
| R4_G3 | V141M/K226G/Q167R/Q362R | — | | |
| E8 | I15V/T25P/T44V/K45E/V141M/K226G | I15V/T25P/V141M/K226G | GTTCGTTCGCCGGTCGAACAAACAGAGTTAATGAATG | CATTCATTAACTCTGTTTGTTCGACCGGCGAACGAAC |
| E9 | T44V/K45E/V141M/K226G/V266R/Q295D | V141M/K226G/V266R/Q295D | GTTCGTTCGCCGGTCGAACAAACAGAGTTAATGAATG | CATTCATTAACTCTGTTTGTTCGACCGGCGAACGAAC |
| E10 | T44V/K45E/V141M/K226G/Q295D/Q362R | V141M/K226G/Q295D/Q362R | GTTCGTTCGCCGGTCGAACAAACAGAGTTAATGAATG | CATTCATTAACTCTGTTTGTTCGACCGGCGAACGAAC |
| E11 | I15V/T44V/K45E/V141M/S173E/K226G | I15V/V141M/S173E/K226G | GTTCGTTCGCCGGTCGAACAAACAGAGTTAATGAATG | CATTCATTAACTCTGTTTGTTCGACCGGCGAACGAAC |
| E12 | I15V/T44V/K45E/V141/T158Q/K226G | I15V/V141M/T158Q/K226G | GTTCGTTCGCCGGTCGAACAAACAGAGTTAATGAATG | CATTCATTAACTCTGTTTGTTCGACCGGCGAACGAAC |

| Well | Combined Mutant | Parent | Forward (5' to 3') | Reverse (5' to 3') |
|---|---|---|---|---|
| F1 | T44V/K45E/A65L/V141M/Q167R/K226G | A65L/V141M/Q167R/ K226G | GTTCGTTCGCCGGTCGAACAAACAGAGTTAATGAATG | CATTCATTAACTCTGTTTGTTCGACCGGCGAACGAAC |
| F2 | I15V/T44V/K45E/V141M/K226G/S248A | I15V/V141M/K226G/S248A | GTTCGTTCGCCGGTCGAACAAACAGAGTTAATGAATG | CATTCATTAACTCTGTTTGTTCGACCGGCGAACGAAC |
| F3 | I15V/V141M/K226G/A262E/E316K | I15V/V141M/K226G/E316K | CGATTAAGTGGAGAGGAAAATTGGGTC | GACCCAATTTTCCTCTCCACTTAATCG |
| F4 | A65L/Q106H/V141M/K226G/L303P | A65L/V141M/K226G/ L303P | ACCCTCTATGCGCACGCCGATATCGAT | ATCGATATCGGCGTGCGCATAGAGGGT |
| R4_H3 | V141M/K226G/A65L/L303P | — | | |
| F6 | I15V/T44V/K45E/V141M/R162A/K226G | I15V/V141M/R162A/K226G | GTTCGTTCGCCGGTCGAACAAACAGAGTTAATGAATG | CATTCATTAACTCTGTTTGTTCGACCGGCGAACGAAC |
| M16 | T44V/K45E | — | GTTCGTTCGCCGGTCGAACAAACAGAGTTAATGAATG | CATTCATTAACTCTGTTTGTTCGACCGGCGAACGAAC |
| F9 | I15V/A65L/V141M/ K226G / L303Q | I15V/A65L/V141M/K226G | ACGGCGCTAGTGCAACAGCGTAAGGGG | CCCCTTACGCTGTTGCACTAGCGCCGT |
| F11 | I15V/T44V/K45E / V141M / E174T / K226G | I15V/V141M/E174T/K226G | GTTCGTTCGCCGGTCGAACAAACAGAGTTAATGAATG | CATTCATTAACTCTGTTTGTTCGACCGGCGAACGAAC |
| F12 | I15V/T44V/K45E/V141M/H161L/K226G | I15V/V141M/H161L/K226G | GTTCGTTCGCCGGTCGAACAAACAGAGTTAATGAATG | CATTCATTAACTCTGTTTGTTCGACCGGCGAACGAAC |
| G2 | I15V / T25S / V141M / Q167R / K226G | I15V / V141M / Q167R / K226G | GCTGCACCAGTGTCGGGTTACACTTTG | CAAAGTGTAACCCGACACTGGTGCAGC |
| G4 | I15V / T25D / A65L / V141M / K226G | I15V / A65L / V141M / K226G | GCTGCACCAGTGGACGGTTACACTTTG | CAAAGTGTAACCGTCCACTGGTGCAGC |
| G5 | I15V / V141M / Q167R / K226G / V266R | I15V / V141M / Q167R / K226G | GCGGAAAATTGGCGCTCATTATTATCG | CGATAATAATGAGCGCCAATTTTCCGC |
| WT | WT | — | | |
| G9 | I15V / T44V / K45E / V141M / R162K / K226G | I15V / V141M / R162K / K226G | GTTCGTTCGCCGGTCGAACAAACAGAGTTAATGAATG | CATTCATTAACTCTGTTTGTTCGACCGGCGAACGAAC |

| Well | Combined Mutant | Parent | Forward (5' to 3') | Reverse (5' to 3') |
|---|---|---|---|---|
| G10 | I15V / A65L / V141M / K226G / Q362R | I15V / A65L / V141M / K226G | TTTGAACTATGGCGGAATCCAGATAAC | GTTATCTGGATTCCGCCATAGTTCAAA |
| G11 | I15V / T25S / A65L / V141M / K226G | I15V / A65L / V141M / K226G | GCTGCACCAGTGTCGGGTTACACTTTG | CAAAGTGTAACCCGACACTGGTGCAGC |
| G12 | I15V / T25D / V141M / Q167R / K226G | I15V / V141M / Q167R / K226G | GCTGCACCAGTGGACGGTTACACTTTG | CAAAGTGTAACCGTCCACTGGTGCAGC |
| H3 | I15V / T44V / K45E / V141M / K226G / E316L | I15V / V141M / K226G / E316L | GTTCGTTCGCCGGTCGAACAAACAGAGTTAATGAATG | CATTCATTAACTCTGTTTGTTCGACCGGCGAACGAAC |
| H4 | A65L / V141M / K226G / G240S / L303P | A65L / V141M / K226G / L303P | TCATCCACATTGAGTGAAATCTTCTTG | CAAGAAGATTTCACTCAATGTGGATGA |
| H5 | V141M / Q167R / K226G / V302N / Q362R | V141M / Q167R / K226G / Q362R | GTGACGGCGCTAAACCTTCAGCGTAAG | CTTACGCTGAAGGTTTAGCGCCGTCAC |
| H6 | I15V / T44V / K45E / A65L / V141M / Q167R | I15V / A65L / V141M / Q167R | GTTCGTTCGCCGGTCGAACAAACAGAGTTAATGAATG | CATTCATTAACTCTGTTTGTTCGACCGGCGAACGAAC |
| H7 | M9L / A65L / V141M / K226G / L303P | A65L / V141M / K226G / L303P | GCACTGGGCTTACTGCTAGGTAGTTTT | AAAACTACCTAGCAGTAAGCCCAGTGC |

### 7.2. Experimental and automation workflows for Phytase engineering

The experimental workflow was organized engineering campaign could begin with the required biological materials, reagents, and plate layouts prepared. The starting *Ym*Phytase plasmid and control constructs were sequence verified, mutation-specific primers were supplied in plate format, and the plasmid templates and vector backbone were prepared and quantified before use. Competent *Escherichia coli* (*E. coli*) DH5α and BL21 cells, selective growth media, autoinduction medium, lysis reagents, 4-MUP assay reagents, and the required PCR, deep-well, omnitray, and assay plates were also prepared in advance. Variant identities, parent templates, primer positions, and control wells were mapped before execution so that each construct could be tracked across the full workflow.

iBioGenie first generated the detailed experimental workflow and organized it into seven modules (**fig. S27A-B).** BiofoundryAI then mapped each module to the available automation capabilities, generated and simulation-tested the executable workflows, and scheduled their execution on the biofoundry through Momentum and the central robotic arm (**fig. S27C).**

Module 1 generated the mutagenic DNA fragments. Parent plasmids and mutation-specific primers were distributed into 96-well PCR plates using computationally generated worklists, and forward and reverse mutagenesis reactions were carried out separately. The workflow used Q5 polymerase with a common 65 °C annealing temperature across the library to simplify automation. Amplification was first checked using EvaGreen fluorescence. For the YmPhytase workflow, the SDM PCR products were further verified by capillary electrophoresis using an Agilent Fragment Analyzer with the DNF-920 kit. The forward and reverse PCR products for each variant were analyzed alongside a DNA size ladder, and the observed fragment sizes were compared with their expected sizes to confirm successful amplification before the products were carried forward for DpnI treatment and HiFi assembly (**fig. S28).** Module 2 combined the mutation-containing PCR fragments with the prepared vector backbone by HiFi assembly to generate the mutant plasmids. The assemblies were incubated at 50 °C for 30 min, then transformed into competent DH5α cells by heat shock, recovered in SOCS medium, and spread onto kanamycin-containing 8-well omnitray plates for overnight selection (**fig. S27D).**

Module 3 recovered individual DH5α clones using the PIXL colony-picking system and inoculated them into Terrific Broth with kanamycin in 96-deepwell plates while preserving the identity of each variant. The cultures were grown overnight to provide sufficient material for plasmid isolation. Module 4 purified plasmid DNA from the DH5α cultures in 96-well format and used the resulting plasmids to transform competent BL21 expression cells. Selected constructs were sequenced as quality-control checks rather than requiring sequence verification of every variant during each engineering cycle. Module 5 established BL21 precultures from individual colonies in selective medium and carried the wild type, empty vector, positive control, and the best variant from the preceding round alongside the new variants as internal references.

Module 6 transferred the BL21 precultures into fresh autoinduction medium in 96-deepwell plates for parallel protein expression. YmPhytase expression was performed overnight for 16 h at 37 °C and 900 rpm in a Cytomat automated shaking incubator. Module 7 harvested the expression cultures, lysed the cells, clarified the crude lysates, and performed the functional screen without an intermediate protein-purification step. Cells were pelleted and resuspended in 200 µL of lysozyme-containing Tris-HCl lysis buffer, followed by incubation and centrifugation to remove cell debris. YmPhytase activity was measured directly from the clarified lysate using 4-MUP hydrolysis in a fluorescence-based plate assay. The screen combined 10 µL of crude lysate with 90 µL of 4-MUP substrate in Tris-maleate buffer at pH 6.6 and monitored fluorescence at 354 nm excitation and 465 nm emission.

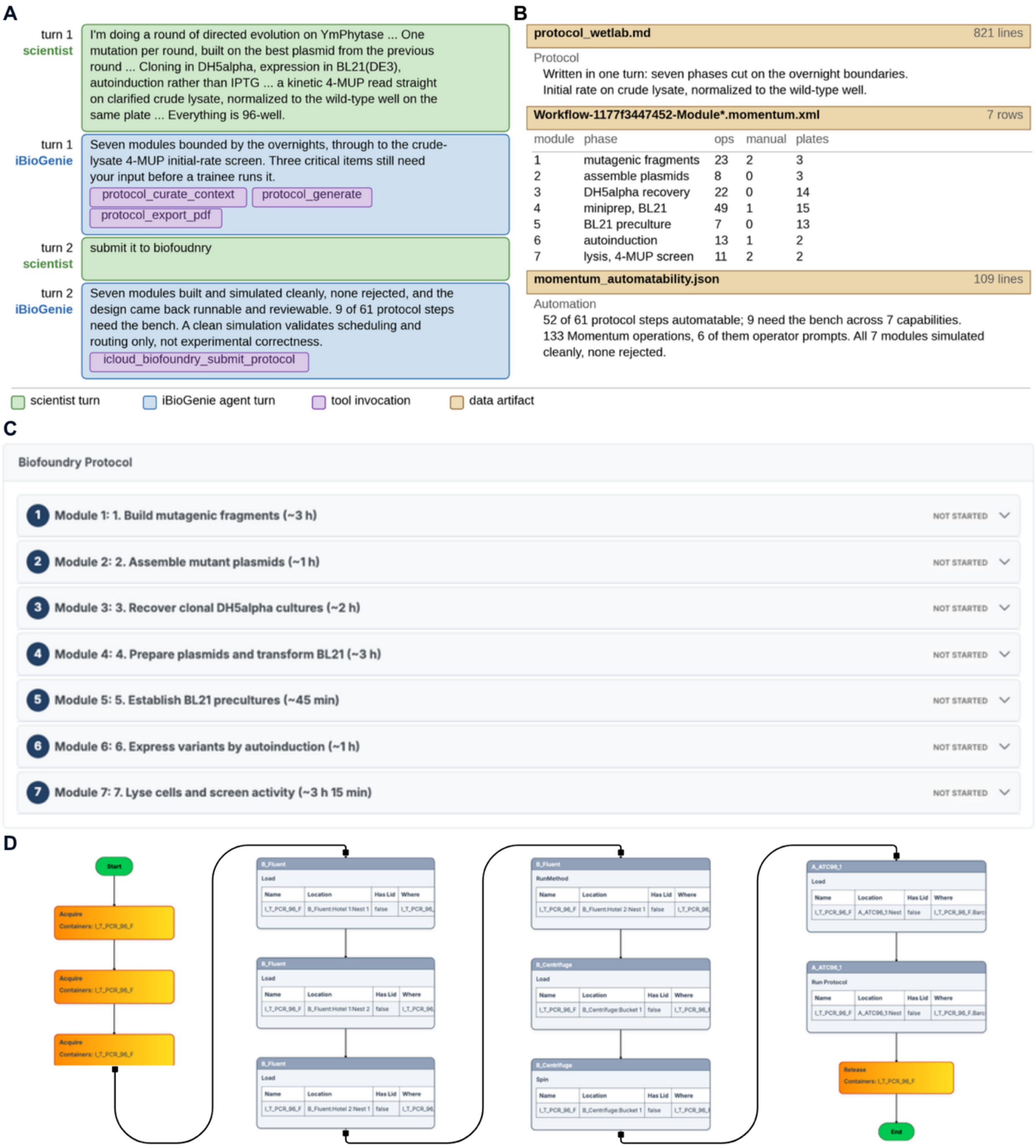


**Fig. S27. Directed evolution of YmPhytase, one round, written in chat and built by the automated laboratory.** iBioGenie writes and submits the bench protocol for one round of site-directed mutagenesis, cloned through *E. coli* DH5α, expressed in BL21(DE3) and read out as a crude-lysate 4-MUP kinetic screen. A clean simulation validates scheduling, labware and routing, not experimental correctness. **(A)** Both exchanges. The scientist card is the prompt as typed, elided with an ellipsis; the agent cards are condensed, and the tools each turn ran are lavender chips carrying their real invocation counts. **(B)** The delivered artifacts, read from the session's own output: the 821-line protocol and its scoring rule, the seven Momentum modules with the operations, operator prompts and containers each carries **(C)** The same protocol in the web application, as the seven modules the biofoundry built from it with their estimated durations. **(D)** The Momentum workflow graph for module 2, assembly of the mutant plasmids, showing container acquisition, liquid handler and centrifuge operations, the thermocycler run and container release.

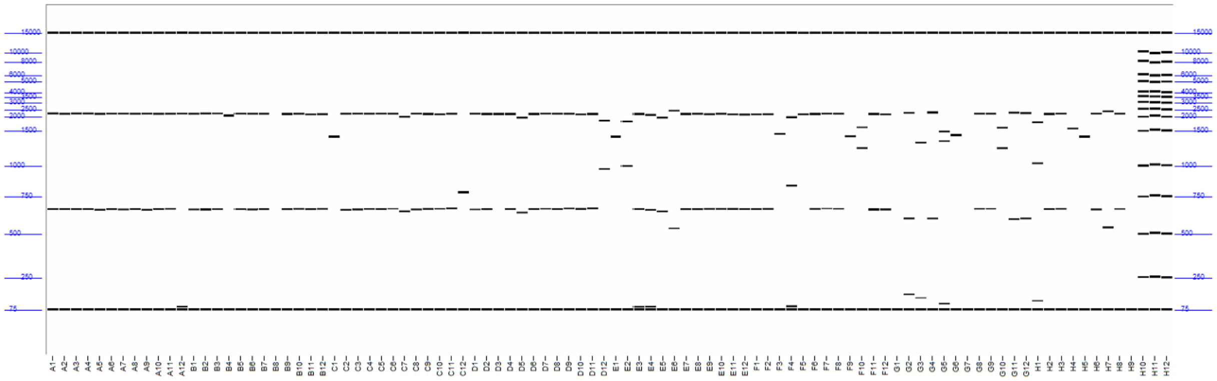

**Fig. S28. Quality check and verification of SDM PCR products by capillary electrophoresis (Agilent Fragment Analyzer, kit #DNF-920) for phytase case study.** Observed fragment sizes were compared to expected product sizes to confirm the identity of SDM PCR products. Each well contains the forward and reverse PCR reactions, run alongside a DNA ladder for size reference.

### 8. Case study II: Integrated PAND discovery and engineering for enhanced 3-HP biosynthesis

#### 8.1. Agentic AI pipeline for enzyme retrieval of aspartate decarboxylase

We first sought to identify novel aspartate 1-decarboxylase candidate enzymes that could outperform the current best enzyme before applying additional engineering. The goal was to move beyond direct mutagenesis of the known enzyme and instead search for alternative aspartate decarboxylase candidates that might provide a stronger starting point for 3HP pathway optimization in *P. kudriavzevii*. The previously known enzyme served as the functional benchmark as it was the best-performing enzyme from the previous screening campaign, producing approximately 1.64 g/L β-alanine and 0.63 g/L 3HP under the tested conditions (*10*). We therefore asked iBioGenie to identify new PAND candidate genes with predicted aspartate decarboxylase activity, potential compatibility with yeast expression, and predicted catalytic performance greater than the baseline.

iBioGenie began by constructing a broad enzyme-retrieval workflow centered on aspartate 1-decarboxylase activity, corresponding to EC 4.1.1.11. The sequence search combined three complementary sources of evidence. First, a targeted UniProt search was used to collect reviewed enzymes annotated with the relevant EC number and appropriate sequence length, producing a curated starting pool of candidate decarboxylases. After removing duplicate amino-acid sequences, this search yielded a nonredundant set of known PAND-like enzymes spanning bacterial, archaeal, and eukaryotic lineages. Second, functionally characterized PAND enzymes from BRENDA were included to capture candidates with reported kinetic evidence, including enzymes with high catalytic efficiency or known beneficial mutations. Third, a homology-expansion step was performed from 5 active selected anchor sequences, including the experimentally validated known enzyme and top 3 high-value curated candidates from BRENDA, to retrieve additional related sequences from large protein-sequence databases with the most phylogenetic similarity to the host.

After sequence retrieval, iBioGenie performed functional annotation to distinguish potential PAND enzymes from broader decarboxylase homologs. Because homology alone can retrieve proteins with related folds but different substrate preferences, each candidate was passed through a function-prediction filter. CLEAN was used to predict enzyme commission class and retain candidates whose predicted function remained within the 4.1.1.x decarboxylase family, thereby enriching for enzymes with PAND-like chemistry while removing candidates predicted to belong to unrelated enzyme classes. This step ensured that the final pool was not simply sequence-similar to panD, but also functionally consistent with aspartate decarboxylation or closely related PLP-dependent decarboxylase activity.

The remaining candidates were then prioritized using predicted catalytic performance. iBioGenie used EZKinetics to estimate kinetic efficiency for the candidate enzymes and compared each prediction. Only candidates predicted to have the highest catalytic efficiency were retained, focusing the search on enzymes with the potential to improve β-alanine formation rather than merely reproduce the existing pathway performance. To further incorporate substrate-level compatibility, candidate structures were evaluated by docking L-aspartate into the predicted enzyme active sites using GNINA. Candidates with favorable docking scores were prioritized, providing an additional structure-based filter for enzymes whose active sites could plausibly bind and position aspartate for decarboxylation.

Finally, iBioGenie applied diversity-aware selection to avoid returning a redundant set of closely related enzymes. The filtered candidates were compared by sequence similarity using MMseqs2, and sequences were chosen if their identity to the initial enzyme was less than 80%. All together the final list was selected based on integrated functional annotation, predicted kinetic efficiency, docking score, and sequence diversity. The ranked and clustered output was manually structured into a final set of 32 top PAND candidates for experimental validation. These candidates represented diverse, computationally prioritized enzymes predicted to retain PAND-like function, exceed the positive control benchmark in catalytic potential, and provide stronger starting points for downstream engineering (**fig. S29, table S18**). Of the 32 proposed enzymes, 28 were successfully constructed, tested, and yielded retrievable activity measurements.

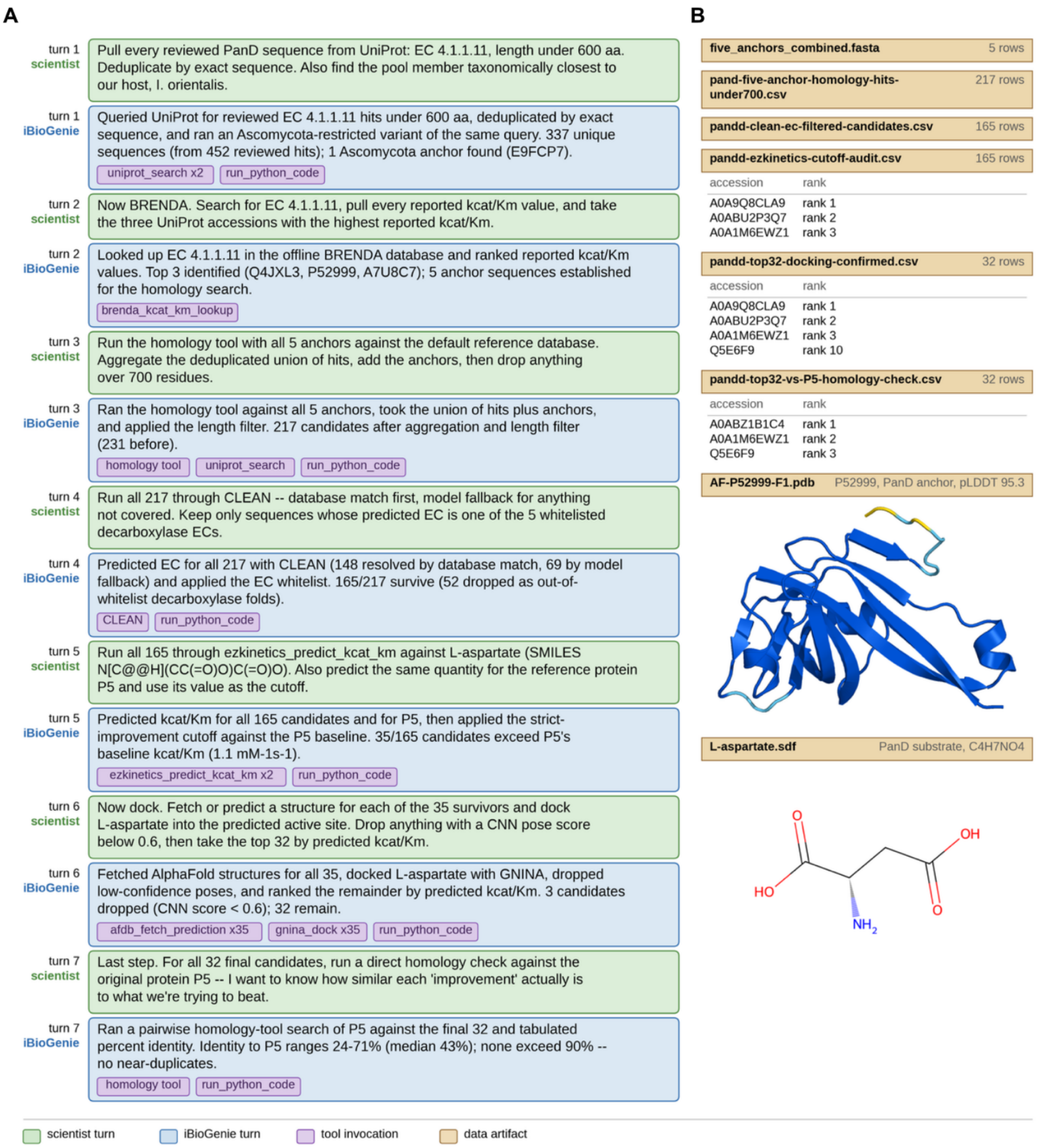


**Fig. S29. iBioGenie PAND retrieval case study. (A)** A summary of the scientist-guided, stepwise search for PAND enzymes and their homologs, beginning with UniProt and BRENDA retrieval, followed by homology expansion, EC filtering with CLEAN, predicted kcat/Km screening, structure-based representation of L-aspartate, and a final homology check against the reference PAND. **(B)** Representative outputs from the workflow, including the five anchor sequences, intermediate candidate tables, the final 32 ranked candidates, homology comparisons to the reference enzyme, and the protein structure and L-aspartate used for docking.

**Table S18. 32 PAND candidates**

| Uniprot ID | Sequence (5' to 3') |
|---|---|
| A0A9Q8CLA9 | AAAACACACAAACAAACACAATTACAAAAAATGATGCATGGCAAAATTCACAGAGCAAGAGTGACACAGG<br>CTGACTTAAACTACGTTGGTTCTATTACTATCGATACTGATATTCTCGAAGCTGTCGACATCCTTCCAAA<br>CGAAAAGGTCGCTATCGTTAATAATAATAACGGTGCAAGATTAGAGACCTATGTCATCGCAGGAGAGCGT<br>GGTTCCGGAGTTATCTGTTTGAATGGTGCGGCAGCCAGATTGGTCCAAGTGGGTGACATTGTTATCATTA<br>TGAACTACGTTGAATTGACGGATGAGGAAGCCAGACAACATAGGCCTAAGGTTGCCGTTATGGATGCGGA<br>TAATCGTATTGCAGAAATGATTGAAGAAAAGGAGGCTACAATCTTGATTTAGATCTAACTAATGCTTTTA<br>CTAAATATCTAG |
| A0ABU2P3Q7 | AAAACACACAAACAAACACAATTACAAAAAATGCAACGTACTTTGCTTAATGGTAAGATTCATAGGGCTA<br>CTGTTACCCAAGCAGATTTACATTACGTTGGTTCCTTGACGATCGACCAAGACCTCATGGATGCAGCTGA<br>TATTGCCGAAGGTGAGCAGGTCCATGTCGTCGATATTGACAACGGTGCTAGATTGACAACTTATGCTATC<br>ACAGGTGCAAGAGGTACCGGTGTGATTGGAGTTAATGGCGCAGCAGCCCACTTAGTGCACCCAGGCCACC<br>TTGTTATCATCATGTCGTTTGCTGCCTTGGACGAGGCCGAAAGAGCTGTCCATCGTCCAAGAGTCGTTCA<br>TGTTGATAAAGCGAACAGAATTGTTGCCTTAGGTGCAGATCCTGCAGAACCAGTTCCAGGTGCGCCTGAT<br>CAATACTCTGGAGCGTTCACACAACAGGAAGGAAATTAGATCTAACTAATGCTTTTACTAAATATCTAG |
| A0A1M6EWZ1 | AAAACACACAAACAAACACAATTACAAAAAATGGAATTGACTATGTTAAAGGGAAAAAATTCATAGAGTGA<br>CTGTTAAGGAAGCGAATATTAATTATGTTGGTTCTATCACGGTCGACAAAGACTTGCTCGACGCTTCCGG<br>AATCTTGGAGTACGAAATGGTCCAGGTGGTTGATATTGATAACGGTGCCAGATTGGAAACCTACACAATT<br>CCTGGCGAAAGGGGTTCAGGTATCATCTCCTTAAATGGTGCAGCTGCTCGTCTTGTTCAAGTCGGCGATA<br>AGGTTATTATTATGGCCTATGCAAAGATGTCGGAGGAAGAAGCGAAGAAGCATAAACCAAGAGTTGTCTT<br>AGCAGATGAGAACAACAAAGTTTCTAAGATCTTTGATTACGAGAAGCACGGTTTCATGTAGATCTAACTA<br>ATGCTTTTACTAAATATCTAG |
| A0ABS7ZDR8 | AAAACACACAAACAAACACAATTACAAAAAATGATGATTGGTAAGATTCATAGAGCAACTGTTACCCAAG<br>CCGATTTGCATTACGTTGGTTCGGTTACAATCGATGCTGACTTATTAACTGCCGCGGACCTTGTCCCTGG<br>TCAACAGGTCGATATCGTTGATGTTACGAATGGTGCTAGGTTGACTACCTATGTGATTCCTGGTGAACCA<br>GGCTCAGGTCAAATTTCCATCAACGGAGCGGCTGCCCACCTCGTTCACCCAGGTGATGTTGTCATTATTA<br>TCGCATATGGAACAATGTCAGATGCTGACGCTAGAACATTCGATCCAGCAGTGGTCTTTGTTGATTCTGA<br>TAATAGAATCGTCGAGTTGGACTCCGATCCAGGTCAGGTTCCAGACGGATACGGCTTACAAGCATCTGGT<br>TTGCCTTTTGCCGCAGTCCGTTAGATCTAACTAATGCTTTTACTAAATATCTAG |
| A0ABZ1B1C4 | AAAACACACAAACAAACACAATTACAAAAAATGTTGAAATCGAAGATCCATAGAGCGACTGTCACGCAAG<br>CTGATTTACATTACGTTGGTTCTGTTACCGTTGATGAGGACTTGTTGGACGCTGCAGACTTATTGCCAGG<br>CGAACAAGTTGCAATTGTCGATATCACAAATGGTGCAAGGCTTGAGACTTATGTTATTCCAGGAGAAAGA<br>GGATCAGGCGTTATCGGTATTAATGGTGCTGCAGCTCACCTCGTTCACCCTTCGGATCTCGTCATTCTTA<br>TCTCCTACGGTGTGATGGACGAAACAGAAGCGAAGTCCTATATTCCTAAGGTGGTTCATGTGGATGCCGC<br>CAACAGAGTTGTCGAATTAGGTGGTGATCCATCTGCCCCAGTCCCAGGTGCAGACGATCAGCGTAGATCA<br>GATTTGGGTGTTCCTGTCAGATAGATCTAACTAATGCTTTTACTAAATATCTAG |
| Q4JXL3 | AAAACACACAAACAAACACAATTACAAAAAATGTTGAGAACCATGCTTAAATCTAAGATTCACAGAGCAA<br>CGGTCACTCAGGCCGATCTCCATTATGTTGGCTCATGTACTATTGATGCTGACTTGATGGAAGCAGCTGA<br>TATCTTAGAGGGTGACCAGATTGACATCGTCGATATCGATAATGGAAACAGATTGACCACTTACGCCATT<br>ACAGGTGATAGGGGTACAGGTGTGATTGGTATTAACGGTGCTGCTGCGAGATTGATTTCCCCAGGTGACT<br>TAGTGATCATCATCGGATATGCGCAATACTCGGAAGAAGATCTTAAGAACTACGCCTCTCGTGTCGTTTT<br>TGTTGATGGTAATAATAAACAAGTTGAGTTAGGTGGAGACCCTGCACAAGTTCCAGATGGTTCCGGCTTG<br>AAGAATCCTAGACATCCAGAAGCATAGATCTAACTAATGCTTTTACTAAATATCTAG |
| A0A8D8QWK3 | AAAACACACAAACAAACACAATTACAAAAAGCTGCATGGGGTGGTGGTGCTTTAGTCTCCAGAAAACATA<br>GGCATTTATTAAATGGTATTGAGAGAGCAGATTCTGTTACATGGAATCCTCACAAATTGCTCACAGCTCC<br>TCAACAATGTTCTGTTTTTCTTACTAGACACTCTACGATTTTGACTGAATGTCATTCGGCCTCAGCCACC<br>TACTTGTTCCAAAAAGATAAGTTTTACGACACCAAGTATGACTCCGGCGATAAGCACATTCAATGCGGAA<br>GAAAGCCAGATGTTTTGAAATTTTGGTTTATGTGGAAGGCTAAGGGTACAGAAGGACTTGAGGCACATAT<br>CGATAAATCGTTCGACAATGCCAAGTATTTCACAGATAAGATTAGAACTAGGGACGGCTTTAAATTGGTG<br>TTGGACGAGCCAGAGTGTACTAATATCTCGTTCTGGTATATGCCACCTTCTTTAAGAGGTAAGGAAAACC<br>AGGCGGATTTTAACGATCTCTTACATAAGGTCGCGCCAAAGATCAAGGAACGTATGATGAAGACCGGTTC<br>CATGATGATTACCTACCAGCCAATCCACGATCTCCCTAACTTCTTCCGTTTGGTTTTGCAAAACTCAGCA<br>CTTAATCACTCAGATATGGACTTCTTTATCGAAGAAATTGAAAGATTGGGTCATGATTTATAGATCTAAC<br>TAATGCTTTTACTAAATATCTAG |
| Q9X4N0 | AAAACACACAAACAAACACAATTACAAAAAATGTTGAGAACTATCTTAGGTTCGAAAATCCACAGAGCGA<br>CGGTTACACAAGCTGATTTAGACTACGTGGGTTCTGTCACTATTGATGCAGATTTAGTTCATGCCGCAGG<br>TTTGATTGAAGGCGAGAAGGTCGCCATTGTTGACATTACAAATGGTGCACGTTTGGAAACATACGTTATC |

| Uniprot ID | Sequence (5' to 3') |
| --- | --- |
| | GTCGGTGATGCTGGAACCGGTAACATTTGTATCAACGGAGCAGCGGCACATTTGATCAATCCTGGTGACC<br>TCGTCATCATTATGTCATATCTTCAGGCAACCGATGCTGAGGCGAAGGCATATGAACCAAAGATTGTTCA<br>CGTTGATGCTGACAACAGGATTGTGGCCTTGGGCAATGATTTGGCCGAAGCTTTACCAGGTTCCGGTCTC<br>CTTACTTCCAGATCTATCTAGATCTAACTAATGCTTTTACTAAATATCTAG |
| A0A2A3E1X6 | AAAACACACAAACAAACACAATTACAAAAAATGGAAAAACAACCTTTAAACCCAAAAGATGATAAGCCTC<br>CTTCTTATTCTGCCGCCACCGCCCCTAAGGTTAACTGTTCATGGCAACCACCACCAGGCTATAATTCTAC<br>TCAATGCCAAGAATCCACTTCTTGGCCTCCTCCACCAGGTTATTACCCTTCGGCTTCGGAAACCAATAAT<br>CATAACTACGTTCCATCTTATGGTTCGACACAGTCCACTACGATTATTATGCCAGAAATCATTCTCGTCG<br>GCGGTTGTCCTGCCTGTAGACCTAAGGTTGTTAATACACAGCAATCCTGTCCTTGGTTAGTTGACATCAA<br>ATACAAGGGTAACGATCATCATGGTTCCCGTTTGAAGAAGACCTCCATGCCTGCTAACGAAGAGACACTC<br>TCAATGACTTCAGACCAAATTAAGGATAACTTGTTTGGTAAGGTGTACGATGATTCAATTCAAAACAATT<br>TCGATTGCGAGTCTTCGTCAGATGACGAGGAAGATTACCAAAATGATTGTTCTAGATCGATTGTTAAGGA<br>AAACTTTAAGGAGATCTGTAACTACAAGTCACTTCCTGTCAGAGAAATCCACAAGAAGTTCATGAAGTCA<br>TTTGTTAATTTGCTCTTGGAAGAAGCTGTGTTCGAAGGTACATTAAGGAGAAATAGAGTCGTTGAATGGA<br>TTGAGCCAACGACATTACACTCTATCATTGACTTGAAGTTGTCCGATCAGGGTTGTTCATATGAAACCCT<br>TCTCACCTTAGCACACAATGTGATTAAGTACTCTGTTAAGACGGGCCATCCAAGATTCATCAATCAGTTA<br>TACTCCTCTGTTGACCCTTATGGTTTGTTAGGTCAATGGTTAACTGATGCATTGAATCCTTCCGTTTATA<br>CGTATGAAGTCTCTCCAGTTTTCTCGTTAATGGAAGAGGAAATTTTGAGAGAGATGAGAAAGATTATTGG<br>TTGGAGAGATGGTAGATCTGAAGGTATCTTCTGTCCAGGTGGATCCATCGCGAATGGCTACGCCATCAAT<br>TTGGCAAGATACTACAAATTCCCTAGATTGAAGGAATTGGGACTCTTTAATACTGGCAGATTAATCATTT<br>TCACATCCAGGGATGCACATTATTCTGTTAAAAAGTTGGGAGCCTTCCTTGGAATTGGTACTGAGAACGT<br>CTATGAGGTGAAAACAGATGACAAAGGAAAGATGTGTATTACCGATTTGAAGATCCAAATTAAAAAGGCG<br>TTACAAGAGAACGCTATCCCTCTTATGGTTTCAGCAACAGCAGGTACCACTGTCCTTGGTGCTTTTGATC<br>CATTGAAGAACATCGCAGCGATTTGTAAGAACTACAATCTCTGGTTCCATGTTGACGCTGCATGGGGAGG<br>CGGTGTCTTAATGTCCAAAAAGTACAAGTACCTTTTGGACGGTATCGAGCTTGCGGACTCGGTCACTTGG<br>AACCCACATAAGTTGTTGGCTGCTCCACAACAGTGTTCTACATTGTTATTGAGACATGAAGGTTTATTGC<br>AAGACGCACACGGTTCAAAGGCATCGTACTTATTTCAGCCAGATAAGTTTTACGATACCTCGTTTGACTC<br>CGGAGATAAACACATTCAATGTGGTCGTAGAGCAGACGTGTTGAAATTTTGGTTTATGTGGAAAGCCAAG<br>GGTACAAGAGGTCTCGAGAAGCATGTTGATAGGGTTTTTAAGTTGGCGCGTTATTTTACTAACTACATCA<br>GACACAGAGAGGGCTTTAAATTAATCTTGGAACCAGAATGCACCAATGTCTGTTTCTGGTATGTCCCACC<br>ATCTAAGAGACAGTTACAAAACGAAGAGTTGTTAAAAACCTTGCAAAAGATTGGTCCTGCAGTTAAAGAA<br>CGTATGATGAAAAAGGGTTCGATGTTAATTACTTATCAACCTCTCCGTGAATTGCCAAATTTCTTCAGGT<br>TAGTCTTGCAGAACTCTGGTCTTACAGAAACTGATATGCGTTTCTTTGCTGAAGAAATTGAAAGGTTAGC<br>TATTGATTTATAGATCTAACTAATGCTTTTACTAAATATCTAG |
| Q5E6F9 | AAAACACACAAACAAACACAATTACAAAAAATGGTTACTGATAACAAAACTGCAGACGCTTCATTTGAAT<br>CCCTTTTGAGGATCTTCACCGTCCCTGAAGCCCCTGACTCAACCTTAGGTATTATCGAGAAGGAACTCTC<br>TCAGAATTTAAATCAATTTCTTAGAGAACATATTGTTGCGGAAGAAAAACCATTGACGGAAATTGAGAAA<br>GACTTTACTGATTCTTCCATGCCAGAGTCTCCTACTTATGTTTCTGAACATACTGAACATTTGTTAGACA<br>CATTGGTTTCGCAATCCGTCCATACATCAGCCCCATCTTTTATTGGACATATGACATCAGCGTTGCCATA<br>CTTTCTTATGCCATTGTCCAAGATTATGATTGCCTTGAACCAGAACTTAGTTAAGATCGAGACCTCCAAG<br>GCATTCACCCCATTAGAGAGGCAAGTTTTAGGTATGCTTCACAGATTAATCTTCGGTCAAAAGGATTCCT<br>TCTATCAACATTGGATGCATTCTGCTGATCACTCACTTGGAGCATTTTGTTCCGGTGGTACTATTGCCAA<br>TATCACAGCATTGTGGGTCGCAAGAAATAGATTATTAAAACCTGAAGGTGATTTCGAAGGTATCGCGAAG<br>CAGGGCTTGTTTGCAGCATTAATGCACTACAAGTGTAACGGTCTTGCTATCTTCGTGTCGGAGAGAGGCC<br>ATTACTCGTTGAAGAAGGCCGCAGACGTCTTGGGTATCGGTCAAGATGGAGTTATTGCCGTTAAGACTGA<br>TAATAATAATCGTGTCTGTTTAGACGATTTGGAGCTCAAGATTGCCCAAGCTAAGGCTAAGAATATTAAG<br>CCTTTGGCTATCGTTGGTGTTGCTGGTACAACTGAAACAGGCTCAATTGATCCTTTGCGTGAACTCGCAA<br>ATGTTGCTCAAAGGGAAGGTTGTCATTTCCATGTCGACGCAGCTTGGGGCGGTGCAACTTTGATGTCGAA<br>TACATACCGTCACTTACTCGATGGTATCGATTTGGCGGATTCTGTCACCATTGATGCACATAAGCAATTA<br>TACGTCCCAATGGGTGCTGGTATGGTTATTTTTAAAGACCCTGAATTGATGTCCTCAATCCAACATCACG<br>CCGAATATATCTTAAGAAAGGGTTCTAAAGATTTGGGAAGACATACCCTCGAAGGTTCTAGATCTGGAAT<br>GGCTATGCTCCTTTACTCGTGCTTCAACGTGATTTCGCGTCCAGGTTATGAGTTATTAATTAACCAATCA<br>ATTGAAAAGGCGCACTACTTTGCCGACTTGATTCAGCAACAAGATGATTTCGAATTGATTACGGAACCAG<br>AGTTATGTCTCTTAACATATAGATACGTTCCATCTAACGTGAAGGCTGCGTTGGCAATTGCAACAGATGA<br>ACAGAAGATCGAGATCTATGAGCACTTGGATAATCTCACGAAATATATCCAAAAGACTCAAAGAGAGACA<br>GGCAAATCGTTTGTTTCTAGAACCAGACTTACTCCTGAAGCATACCAACACCAACCTACGATTGTGTTCA<br>GAGTTGTCTTGGCGAACCCATTGACCACCAAAGAAATCTTGCAGAACGTTTTAATCGAACAGAGAGAAAT<br>TGCCTCGTCTTCCGAGATTTCTTTGCCATTGTTAAACCAGATTGTCGGAAACATTTTGCACTAGATCTAA<br>CTAATGCTTTTACTAAATATCTAG |

| Uniprot ID | Sequence (5' to 3') |
|---|---|
| C1D1G6 | AAAACACACAAACAAACACAATTACAAAAAATGGAAAGAATTATGTTTCGTGCCAAGATTCATCGTGCTA<br>CGGTGACTCAGGCCGATTTGGACTATGTCGGTTCGGTTACAATCGATCAAGATCTTTTAGATGCAGCAGA<br>TATTCTCGTTAATGAAAAGGTCGATATCTGGAACATCACCAATGGTAATAGACTCCACACTTACGCCTTG<br>TCCGGTCCTAGGGGTTCTGGAGTTATTGGAATTAATGGTGCTGCGGCACATTTGATGAGACCTGGTGACA<br>TGGTTATTATCGCAGCGTTCGGCAACTTTACAGAGGAGGAAGCAAGAGTCTTGGAACCAAAAGTTGTCTT<br>GGTTGACGCTAGAAACAGATTATTAGAACTTGAGCCAGCTTAGATCTAACTAATGCTTTTACTAAATATC<br>TAG |
| A0A1D7VLP2 | AAAACACACAAACAAACACAATTACAAAAAATGTTGAGGACCATGTTTAAATCGAAGATTCACAGAGCGA<br>CAGTCACGGAAGCAGATTTGCATTATGTGGGCTCCGTCACTATTGACGCTGCATTAATGGAGGCAGCAGA<br>CCTTTTACCAGGTGAACTCGTTCATATTGTTGATATTGATAACGGCGCCAGATTGGAGACTTACGTCATC<br>GAAGGTGCGAGAGATTCAGGAGTTATCGGTATTAATGGTGCTGCAGCACATTTGGTCCATCCTGGTGATC<br>GTGTCATCTTGATTTCTTACGCGCAAGTTGAAGACGCAGAAGCTCGTGCTTTAAGACCAAAGGTTGTTCA<br>CGTTGATGCAGGAAATAGAATCGTGGCTCTTGGTGCCGACGCTTCGGAGCCTGTTCCAGGTTCCGATACA<br>TCTAGGTCACCAAGAGCCGCCGCCGTCTAGATCTAACTAATGCTTTTACTAAATATCTAG |
| Q118E6 | AAAACACACAAACAAACACAATTACAAAAAATGACTACCATTAAGTTGATGCACGCTAAATTACATAGAG<br>TTCGTGTCACCGATGCAAAGCCTGATTATGTCGGATCGGTCACAATTGATTCCGACTTGTTGGAAAAGGT<br>GGGTATTTTACCACTCGAAGAAGTTCAAATTGTGAATTTAAATAACGGTCAGAGGTGGCTCACATATGTT<br>TTGCCTGGTCAGGCCGGTTCCGGAATGGTTTGTCCAAATGGTGGTGCCGCATTGTTATGCAAAAAGGACG<br>ATATCTTAATCATCTGGGCTAACGAACAATGTGATAGAGCTGAAGTTTTAGAGAACGGTCACAAGGCGAA<br>AATTTTCGTCGCAGACGAGAACAATCATTGTAGAGAGTTTTTGTACCAGATTCTTAATCCAAACCAAGGC<br>TCTGTTGAATTTCATTCACTTCCAGTTATTCCTGAAGGCAAGACGTCTGAGTACTTGCAACAAATGATCG<br>AAACTAATAAGATCTAGATCTAACTAATGCTTTTACTAAATATCTAG |
| A0A8D8A8L5 | AAAACACACAAACAAACACAATTACAAAAAATGGGAATTGGTTCTGACAATGTTTATCCTATTAGGACAG<br>ACGCCGTCGGTAAGATCCAGCCAGATCATTTAGAAGCAGAAATTCTTAGAGCAAAGTCAGAGGGTGCTTT<br>ACCATTCATGGTGTCAGCCACTGCCGGTACAACTGTCATCGGTGCGTTTGACCCTCTCGAGCAAATCGCA<br>GACTTGTGTCAAAAGTACGATTTGTGGATGCATGTCGATGCGGCTTGGGGAGGTGGTGCATTGATGTCTA<br>AGAAGTACAGAACTTTGCTCAAGGGAGTCGAAAGAGCTGATTCCGTTACTTGGAATCCTCACAAGTTATT<br>GGCTGCACCTCAACAATGTTCAACCTTTCTTACAAGACACGAGGGCATCTTATCCGGTTGTCACTCTACC<br>AACGCGACTTACTTGTTTCAGAAGGATAAATTTTATGACACCCAGTACGACACCGGAGATAAGCATATTC<br>AATGTGGTAGAAGAGCCGATGTTTTAAAGTTTTGGTTCATGTGGAGAGCAAAAGGTACATCGGGTTTCGA<br>ACAGCACATCGATAAAGTCTTTGAGAACGCAGAGTACTTCACGAATTCCATTAAGGCCAGACCTGGTTTC<br>GAAATGGTGATTGAAAATCCAGAATGTACGAACGTTTGCTTTTGGTATGTTCCACCAGGTTTGAGGCAAG<br>TCCCACGTGACTCTGCAGAGTTCGGTGAACGTTTGCATAAGGTTGCTCCTAAGGTTAAAGAAAGAATGAT<br>GAGAGAAGGCTCTATGATGATTACATACCAACCAATTCATGATAAACCAAACTTCTTTCGTTTGGTTCTC<br>CAAAACTCGGGCTTGGATAAATCCGATATGAATTATATTATCGATGAAATCGAAAGGCTTGCTTCGGATT<br>TATAGATCTAACTAATGCTTTTACTAAATATCTAG |
| A0A0C5BI73 | AAAACACACAAACAAACACAATTACAAAAAATGTACAGAACCATGTTATCCGGTAAAATCCATAGATTAA<br>CCGTTACCCACGCGGATCTTCACTACGTTGGCTCTGTTACAATCGACAGAGATTTGATGGAAGCGGCCGA<br>CCTCTTACCAGGTGAGCAGGTGACTATTGTTGATATCACGAATGGTGCTAGGCTCGAAACTTATGTGTTG<br>GAGGGCGAGAGAGGATCGGGAGTCGTTGGTATTAATGGTGCAGCCGCTCATTTGGTTCATCCTGGTGATT<br>TGGTCATTGTTATTGCATATCAACAAGTTCCATCCGTTTCGGCAAAGGATTTTGTCCCACATGTCGTCCA<br>CGTTGATGAAAACAATCGTGTCATCTCTTTGGGTACTGATGCCGCTGAAGCAGTGACAAACGACGTCGAA<br>AGATCACCTTTCACTACAACACTTTTGTAGATCTAACTAATGCTTTTACTAAATATCTAG |
| A0A8K1X5N7 | AAAACACACAAACAAACACAATTACAAAAAATGGAAGAAACTGTTTTACGTGAAATGAGAAAGATTGTTG<br>GTTTTAAGGACGGTAAGGGTGATGGTATCTTCTGTCCTGGTGGATCTATCTCGAACGGTTACGGTTTGTC<br>TTGTGCAAGACATAAGAAGTTTCCAGACATCAAGACTAAGGGTTTGAACTCATTGCCAAGATTGGTTTTG<br>TTCACGTCCGAGGATGCACATTATTCGATTAAGAAGATGGCGTCCTTTCAAGGTGTTGGTTCGGATAATG<br>TGTACTGTATTAAGACCGACAGACGTGGCAAGATGGATATCTTAGATCTCGAGGCACAAATCATTAAGTC<br>TATCGAAGAAGGTGCCGTCCCATTTATGGTCTCTGCTACTTCTGGTACTACGGTTTTAGGAGCCTTTGAT<br>CCTTTGAATGAAATCGCAGATATTTGCGAGAAATATGGCTTGTGGATGCATGTGGATGCCGCTTGGGGTG<br>GTGGTGCGTTGATGTCACAGAAGTACAGAAAGTTGCTTACAGGCATTGAAAGAGCTGATTCCGTTACTTG<br>GAATCCACATAAACTCCTCACCGCCCCACAACAATGTTCGGCATTTCTTTTAAGGCACGAAGGTTTGTTA<br>GCGGAAGCACACGGATTAAACGCCGCTTACTTGTTCCAGAAAGACAAGTTCTATGATACCAAGTACGATA<br>CAGGAGACAAACATATTCAATGTGGTAGAAGGGCAGACGTCTTGAAGTTCTGGTACATGTGGAAGGCTAA<br>GGGCACAAACGGTCTCGTCAAACACATTGAAAGAGTTTTTGAGAACGCTGAGTACTTTACCGATTTAATC<br>CGTAACAGACCAGGTTTCGAAATGGTCCACCCTGAACCTGAGTGTACTAATATTTGTTTTTGGTACATTC<br>CTAAATCATTAAGAGATTGCAAAGACAACGATAATTATAAGGAAAGATTGCACAGAGTGGCTCCTGCGAT<br>GAAAGAGAAGATGATGAAGGAAGGAACAATGATGGTCACATATCAAGCACAGAAAGGACATCCAAATTTC |

| Uniprot ID | Sequence (5' to 3') |
|---|---|
| | TTTAGGATTGTTTTCCAGAATTCCGGCTTGAACAGATCCGACATGGAACATCTTGTTCAAGAGTTCGAAC<br>GTTTAGGTTCAAATCTTTAGATCTAACTAATGCTTTTACTAAATATCTAG |
| P52999 | AAAACACACAAACAAACACAATTACAAAAAATGTATAGGACAATGATGAATGGTAAATTGCATAGAGCGA<br>CAGTTACTGAAGCGAATTTGAACTACGTTGGATCTATTACGATCGATGAAGACCTTATTGATGCAGTTGG<br>TATGTTACCTAACGAGAAGGTGCAAATTGTTAATAACAATAACGGTGCCCGTCTCGAAACCTATATTATC<br>CCAGGCAATAGAGGCTCAGGTGTCATCTGTTTAAATGGTGCTGCTGCAAGATTGGTGCAAGAAGGTGACA<br>AGGTCATCATTATGTCTTACAAAATGATGTCCGATCAAGAGGCTGAATCGCACGAACCAAAGGTTGCAGT<br>CTTAAACGATCAGAATAGAATTGAGCAGATGTTGGGAAACGAACCTGCCAGAACTTAGATCTAACTAATG<br>CTTTTACTAAATATCTAG |
| Q9EYU1 | AAAACACACAAACAAACACAATTACAAAAAATGAGAAAACTCGTGGCGGGTAAGCTTCACGGTATTCATG<br>TTACAGAAGCTAACCTTAACTACCATGGCTCAATCACCCTCGATCCTGATCACTGCGAAGCTGCAGGTAT<br>TTTGCCAATGGAGTTCGTTGAGATTTGGAATAAGAACTCCGGCGCCAGAATCTCCACATATGTTATCTTA<br>GGAGAAAGAGGTTCTAGGTGTTGTATTTTGAATGGTGCTGCAGCACGTACTTGTCAACCAGATGATCCAA<br>TCATTGTCTGTAATTCGATTTACTTGGACGAGGCCCATATCACTTCATTAAAGCCTAGAATCGTCACCTT<br>TGACCAGGATAATCATATTTTAGATCGTTTGTCCTACTCTGTTGACATTGATACGGACGGAAGATATTCT<br>TTTGCTATTTTGGACGAAGCAGATGAACCATTGGTTATCCCTGCCTTAGTCTCGGGTGCGTAGATCTAAC<br>TAATGCTTTTACTAAATATCTAG |
| A0A1B6I626 | AAAACACACAAACAAACACAATTACAAAAAAGAACTTTGAAGGAAGGTGCCAAGCCTTTTATGGTTTCTG<br>CAACAGCTGGTACCACTGTTTTAGGTGCTTTCGACCCATTAGTTGCTTTGGCGGATGTGTGTGAAGCACA<br>CAAACTTTGGTTACATGTTGACGCGGCGTGGGGTGGTGGAGCTTTGATGTCTCGTAGGCACAGACATTTG<br>TTGAATGGTATCCAACGTGCAGATTCAGTTACCTGGAATCCACATAAGTTACTCACGGCTCCTCAGCAGT<br>GTTCAACATTCCTTACTAGACACACAGACTTACTTGGACCTTGCCATTCCGCAGGTGCAACCTACCTCTT<br>CCAAAAGGATAAGTTTTATGACACTAAGTACGATACGGGCGACAAGCATGTCCAATGTGGTAGAAGAGCA<br>GACGTTTTGAAATTTTGGTTCATGTGGAAGGCCAAGGGCTCCGATGGTTTCGAAGCACATATCGATAGAG<br>TCTTTGATAACGCTCAATATTTGACAGAGCAAATTAGACATAGGCCAGGATTTAAGTTGGTGTTGGATAA<br>ACCTGAGTGTACCAACGTCTCGTTTTGGTACATTCCACCATCGTTAAGAGGTCAAGATACTGCCCCAGAT<br>TATCACCAGAGACTCCACAAAGTCGCCCCTAAATAGATCTAACTAATGCTTTTACTAAATATCTAG |
| A0A146M9I1 | AAAACACACAAACAAACACAATTACAAAAAATGCCAGCTTCCGAGGCGTCCGCGGAAAGACCAAGAGCAG<br>ACGTGGTTATTGAAGAAATTGATGGTGCTCCATCAAAGGAGGCATTTCTTTCTGCACCAGACAAGGAGAT<br>CCATGGACAATTTTTACACAGAGTCCAACAGATGTTATTGGATCATGCATTTCAATCAACTCATAGAGAA<br>TCTAAAGTCACAAGATACGTCGATCCAGATAGGTTACGTGAAATCATTGATATGAATTTGAGAAAGGAAC<br>CTGCTTCTGAAGAAGCACTCTTATCGTATATCCGTGACACCATGAAGTACTCAGTCAAAACAGGCCATCC<br>TTTCTTTGTTAACCAATTGTTCTCATCGCTTGATCCATATGCCTTAGCGGGTGCTTGGGTTACCGACTCT<br>TTAAACCCATCTGTTTACACTTACGAAGTCTCTCCTGTGTTAACACTTATGGAAGAATCTGTTTTGAAGG<br>AAATGCGTTCGATTGTCGGTTGGTCTGATGGAGACGGTATCTTTGCCCCAGGAGGTTCCATTGCAAACGG<br>TTATGCTATCTCCTGTGCTAGGTTCCACAAGATCCCAGATATTAAGGCCAAAGGCCTCCATGGTGTGCCT<br>AGATTAGTTTTGTTCACCTCGGAGGATGCACACTATTCTATCAAGAAGTTGGCGGCGTTTGAAGGTTTAG<br>GTTCAGACAATGTTTACTCCGTTAGAACTGACCCTAGAGGTAAAATGGATGTCTTGCATTTGAGAGAGTT<br>GGTTCAAAGGGCCAAAGCAGAGGGTGCAGTTCCTTTCATGGTTTCTGCCACATCGGGTACTACGGTTTTA<br>GGTGCATACGATCCTCTCGAAAAGATTCACGAAGTGTGTGAAGAGGAGGGAATGTGGTTGCACGTTGATG<br>CCGCTTGGGGTGGTGGCGCACTTGTCTCCAAGAAGTACAGACACTTGCTCAAGGGAATTGAGAAGGCAGA<br>TTCCGTCACTTGGAACCCTCATAAGATGTTAGCCGCTCCACAACAGTGCTCAACATTGTTGACCAAGCAC<br>AAGACCATTTTGGCGGAATGTCATTCTGCTTCGGCACAGTACTTATTCCAGAAGGACAAATTCTATGACA<br>CGTCCTTCGATACAGGTGATAAGCATATTCAATGTGGCAGAAGAGCTGATGTCTTTAAATTTTGGTTGAT<br>GTGGAAAGCCAAAGGTACTGATGGTTTGGAAAAGCATGTTAATCGTGTTTAGATCTAACTAATGCTTTTA<br>CTAAATATCTAG |
| A0K1U2 | AAAACACACAAACAAACACAATTACAAAAAATGAATAGGACTATGTTTAAGTCCAAGATTCATAGAGCTA<br>CAGTCACTCACGCGGATTTGCATTATGTGGGCTCGGTCACTGTTGATTTGGATTTATTAGAAGCCGCAGA<br>TATCTTGCCAGGTGAACTCGTTGCAATTGTTGATGTTACCAACGGTGCCAGACTCGAAACCTACACGATT<br>GCAGGTGAGAGAGGATCAGGTGTCATCGGTATCAATGGTGCGGCTGCTCATCTTATGCACGAAAACGATA<br>TCGTTATTCTTATCACTTATGCCGAGATGACAACCGAAGAGGCAAAAGCTTACACACCTAAGGTGGTTCA<br>TGTCGACAAGGACAACAAAATTGTTCAAATTGGAAATGACCCAGCTGAAGGCCACACAAGAGGTTTAATG<br>CGTCCACCTTTCGCCTTGAACAATTCTGCATTGAATTAGATCTAACTAATGCTTTTACTAAATATCTAG |
| B8ZU50 | AAAACACACAAACAAACACAATTACAAAAAATGTTGAGAACTATGTTAAAGTCTAAGATTCACAGAGCGA<br>CTGTTACCCAGGCATATTTGCATTATGTTGGATCGGTGACAATCGACGCTGATCTTATGGGTGCCGCAGA<br>CCTTTTGGAGGGTGAACAAGTGACAATCGTCGACATTAACAATGGTGCAAGACTCGTTACTTACGCGATC<br>GCAGGCGAAAGAGGCACCGGTGTCATCGGTATTAATGGTGCTGCTGCCCATTTGGTTCACCCAGGTGATT<br>TAGTTATTCTCATTTCTTACGGCACGATGGAAGACGCAGAGGCACATGCCTACCAACCACGTATTGTCTT |

| Uniprot ID | Sequence (5' to 3') |
| --- | --- |
| | TGTCGATGCTGATAACAAACCAATTGATTTGGGACATGATCCTGGTTCAGTTCCTTTAGATATCTCCGTT<br>GCCGCGGAATTGTTCGATCCAAGAATCGGTGCTAGGTAGATCTAACTAATGCTTTTACTAAATATCTAG |
| A7IAB9 | AAAACACACAAACAAACACAATTACAAAAAATGTTATCGAACGGTATCTCAGAGGATGATTTGTTTTCCT<br>TTCTTTCTTTGCACAAGAAGGAAGATCTCGACCATAGATATATCCTCTCATCCATGTGTACTTTGCCACA<br>CCCTGTTGCAGTCAGAGCCCATTGTATGTTTATGGAAACCAATTTGGGTGACCCTGGCTTATTCCCTGGT<br>ACAGCTGCTCTCGAGAGGTTATTGGTGGAAAGATTGGGTACATTGTTCCATCATAAGAATGCGGGAGGCT<br>ACGCTACCTCTGGTGGTACTGAGTCTAATATTCAGGCCTTAAGGTTAGCCAAAGCGCTCCGTCCAGGTTC<br>TTCTCCAAACGTTGTTTTACCTGAATCGGTTCATTTCTCCTTCAAGAAGGCTTGCGATTTATTGTCGTTA<br>GAAATGAGGTCAGTGCCACTCGGTACCGATAGAAGAATCATGGCAGACAAGGCTGCGGAATTGATCGATA<br>AGAATACCATCTGTTTGGTGGGCGTTGCAGGTACCACTGAATACGGAATGGTCGATCCTATTGCGGATCT<br>CGCCAAAATTGCAGCCCAACAAGATATTTTCCTTCACGTCGACGCAGCATTTGGAGGTATGGTTATTCCA<br>TTCCTTCCTAAGCCTGTTCCATTTGATTTTGCTTTGCCTGGTGTCACTACTTTAGCAGTCGATCCACATA<br>AGATGGGTATGTCTACGATTCCAGCCGGAGTTTTATTGACAAGAGAACCTGACATGCTTGACGCCTTGAA<br>CATCGATACTCCATATTTGACAGTTAAGAAGGGATACACGTTGGGTGGTACTCGTCCTGGTGCTCCAATG<br>GCAGGTGCCTTGGCTGTCTTGGATTATTTAGGTATTTCCGGTATGAAAGCAGTCGTTGCGGGATGTATGA<br>AAAATACAGAAAGATTAATCGCTGGTATGGAGACTAGAGGTATTCAACCAGCTGCATCACCTGATGTTAA<br>CGTTGCAACCTTCGTTTGTGATAGAGTGCCAGAGCCTTGGAAAGTCTCCTGGACAAGAGCTGGCCACTTA<br>CGTATTGTTTGTATGCCACACGTCACAGCAGATAGAATTGAAGCATTTCTTTCGGACTTTGGCGACATGT<br>ACGCGTAGATCTAACTAATGCTTTTACTAAATATCTAG |
| A0A9W6G5Z9 | AAAACACACAAACAAACACAATTACAAAAAATGATTAGGACAATGTTCAAGTCGAAAATCCATAGAGCAA<br>CCGTTACTCAAGCCGATTTGCACTATGTCGGCTCTGTTACTGTTGATGCGGATTTATTAGATGCTGCAGA<br>TATCTTAGACGGCGAATTGGTTGCTATCGTCGACATTACCAATGGTGCACGTTTGGAAACATACACTATC<br>GCTGGTGAGAGAGGTACGGGTGTCATCGGAATTAATGGAGCCGCTGCTAGACTCGTCCAACCTGGTGACC<br>TTGTGATTTTGATTTCATACGCACAGATGGAGGATGCCGAAGCCCGTGCCTTTAAGCCAACCGTTGTTCA<br>CGTGAACGAAAGAAACGAAATTATTGAGGTCAACGCAGATAAGGCTCATCCAGCGCCAGGTACAGCAGGT<br>AATCCTGTTGCATCCCCACTTGCGGCAAGATAGATCTAACTAATGCTTTTACTAAATATCTAG |
| A0A6I4NYF5 | AAAACACACAAACAAACACAATTACAAAAAATGCCATCATTCCGTCGTACAATGTTGAAATCAAAGATTC<br>ACAGAGCGGCCGTGACTCATGCTGACTTACACTACGTTGGTTCCATCACTGTCGATGCAGACCTTTTGGA<br>TGCCGCAGATTTGTTGCCTGGCGAGTTAGTTACCGTCGTCAACATTAATACCGGTGCTAGGTTTGAAACT<br>TATACGATTGAGGGTGAAAGAGGTTCCGGTGTGATTGGTGTTAATGGCGCGGCCGCAAGACTTGTCGCTG<br>TTGGTGACTTGGTCATTCTCATCTCGTGGGCATCGATTGATTCTGAAGAAGCTAGAGAATTAGTTCCAGC<br>AATCGTTCATGTTGATGCAGCCAACAGAATCATCTCTTTGGGAGGAGATCCAGCAGAAGCTTTAGTTCCA<br>GGTTTGGAGACACCTCCTCTCGCCGTCAGAGCGTAGATCTAACTAATGCTTTTACTAAATATCTAG |
| Q0RB93 | AAAACACACAAACAAACACAATTACAAAAAATGCTTAGAACAATGTTGACAGCAAAGATTCACCGTGCGA<br>CCGTTACTCAGGCTGACCTCCATTACGTTGGTTCGGTTACTATTGATGCAGATTTGCTTGAGGCCGCTGA<br>TCTCTTACCAGGTGAGCAAGTTACTATCGTTGATATCAATAATGGTGCTAGATTAGAAACCTATGCGATC<br>GCTGGTCCAGCAGGATCCGGTGTGATTGGAATTAACGGTGCCGCCGCGAGGTTAGTCCAACCAGGCGACT<br>TGGTCATTATCATTGCCTACGGTATGATGGACGATGCAGAAGCTAGGCGTCACGTCCCAAGAGTTTTGTT<br>TGTGGATGCTGAAAATAGAATCATTGGTCGTGGACATGACCCTGCAGAAGCTTTGCCTGATGATCCTTCT<br>TCATTGAGAGGCGACTTGGCCGTCCCTGGTAACCCAGTCACTGCAGCGGCAAGAAGAGGTACCCCAACAC<br>ATCAAGCACCTGTCGCATTACCTGCCTCTAGAACGGTTGTTGCTCCAAGATAGATCTAACTAATGCTTTT<br>ACTAAATATCTAG |
| A0A132MNB2 | AAAACACACAAACAAACACAATTACAAAAAATGTTACGTACAATGCTCAAGTCAAAAATTCATAGAGCTA<br>CTGTTACTCAAGCAGATCTTCATTATGTCGGTTCGGTTACAATCGACGAAGACCTTATGGATGCCGCTGA<br>TTTGTTGCCAGGTGAGCAAGTCCAGATTGTCGATATTGATAACGGAGCGCGTTTAGAAACCTACGTTATT<br>CCAGGCGAAAGGGGTTCCGGTGTTATTGGTGTTAATGGTGCGGCTGCAAGATTGGTGCACCCTGGAGACT<br>TAGTTATCATCATTTCTTACATGACAGTCACTGATGCCGAGGCAAGAACGTTGAAGCCTAGAGTGGTCTT<br>TGTTGGAACCAGAGAAGGTAGGGCTAATAGAATCATCGGTACCGGTGGCGATCCAGCAGAAGCCGTTCCA<br>GGTACTGGCACAGTTAGAGGTGACGTCATTGTGTAGATCTAACTAATGCTTTTACTAAATATCTAG |
| A0ABT1JDK7 | AAAACACACAAACAAACACAATTACAAAAAATGGACCTTGAAATGCTTCAGGCGAAAATTCACAGAGCTA<br>CAGTTACTAGGGCCGACTTGCATTACGTTGGTTCATTGACGTTGGATGCAGATTTGATTGCAGCCGCTGG<br>TATCTTACCTGGTCAAAAGGTTGATGTTGTCGATATTGACAACGGTGCGCGTTTGTCTACATACGTGATT<br>GAGGGTCCAGCAGGATCCGGCACCGTCGGTATTAATGGAGCAGCAGCTAGATTAATCTCTCCAGGTGATC<br>TCGTTATCATTATCGCTTATGCGAGAATGCCTGCCGAAGAAGCAAGGAGATTTGTCCCATCGGTCGTTTT<br>CGTCGATGAACGTAATAGAGTGGTTAGACAAGGTGTCGATCCTGCTGAGCCTTTACCAGGTACTGTTCAT<br>GGAGCCACCCCAGACGGCGGTGCAGCTTAGATCTAACTAATGCTTTTACTAAATATCTAG |

| Uniprot ID | Sequence (5' to 3') |
|---|---|
| Q8U9Q9 | AAAACACACAAACAAACACAATTACAAAAAATGATTAGAATCGTTAGAGCAGAGTTGCATGGTATCACTG<br>TCACTGGTGCCGATTTAGATTACCATGGATCGATTACATTAGACCCTGAGCACTGTGACTTGGCTGGCAT<br>TAGACCACTCGAATTTGTCGATATTTGGAACAAGCAATCCGGAGCAAGAATCTCAACTTACGTCATTTTT<br>GGTGAAGCTGGTTCGAAGTGTTGTATCTTGAATGGCGCAGCCGCCAGGACTTGTCAAAGAGGTGATCAGG<br>TTATTATCTGCTCTTCTTCTTATGTTGAAGAGACAGCGCTTTACGACACCGCTCCTATGGTTTTGACATT<br>CACCCCAAAGAATGATGTCAAAGATGTGTTAAGATATCGTGTTAAAGAAACCGCTAGACGTCCATTTGAT<br>TTCTTCATTGAAACGCTTCCAAGAGCGGAAAACTCAGACGTTGATGATGGTTCCGCAGCAGACAGGGCCA<br>TCGCTTAGATCTAACTAATGCTTTTACTAAATATCTAG |
| Q7UQU4 | AAAACACACAAACAAACACAATTACAAAAAATGGTTGATACTCCATACAGAAAGATGTTGGCCGCTAAAA<br>TCCATCGTGCCACAGTCACGGGTGCCGACGTTAATTATGAGGGTTCCCTTACTGTCCCTCCTGAGCTCTT<br>AGTTGCAGCTAAGATTCATCCTTACGAATCGTTACATGTGTGGAACGTCACCAGAGGTACTAGACTTGAA<br>ACATATGCGATTGAAGGATTACCAAACTCCAATGATGTTTGTGCCAACGGTGCAGCGGCACACTTGATCA<br>GGCCAGGAGATCATGTCATTTTGGCTGCATACGCTATGGTTCCAGAAGCGGACGCAGCAACCCATAAGCC<br>AAGATTGATTTTTGTTGACGATAATAACCAATTGTCTCACGTGGGTCCTGAAATCGCTGGTCCAAATCTC<br>AGATCTGATTCAGATGACACCCACTTGGTTCGTTCAACTGAAATGACACCTGATGGCCAGCCATTAGCAG<br>AGGGCTGTTAGATCTAACTAATGCTTTTACTAAATATCTAG |
| Q2VZZ9 | AAAACACACAAACAAACACAATTACAAAAAATGATGAAGATTATCAGAGCTAAGTTGCATGGTATTCGTG<br>TTACTAACGCAGACCTTAATTATCACGGTTCCATTACATTAGATCCTGAACAATGTGAGATGGCCGGAAT<br>TTATCCAATGGAATTTGTCGATATCTGGAACAAAAACTCAGCAGCCAGGATTTCTACTTACGTTATCTTT<br>GGTGAGCCTGGTTCAAGATGTTGCGTTTTAAATGGTGCCGCAGCTAGAACCTGTCAGAAGGGAGATGAAT<br>TAATCATTGCTGCTTCCGCCGATATTTCGGGCCCAGAAAAGTTGTACGATATCAAGCCAAGAATCCTCAC<br>ATTCCTTCCTGACAATCATGTGGACCAAGTCCTCTACTACGACGTCTTCCAATCTGAGAAACGTCCATAT<br>GATTTTAGAATCGTTGATGCAGATAAACATACGATCGAATCATGTCATACCTGGCCAAATGTTGACATTA<br>CTAAGTTGAGATCTGATTTGGCGGCAAAGGGTTGGTCGGAAGCGGAAATTGATTCGTTTATTGCTTCCCA<br>CTTCTCTTTGTAGATCTAACTAATGCTTTTACTAAATATCTAG |
| S4XK29 | AAAACACACAAACAAACACAATTACAAAAAATGAACAGGAGAATGTTTAAATCTAAGATCCATCGTGCTA<br>CCGTTACACATGCTGATCTCCATTATGAGGGTTCAGTCACCATCGATGTGGATTTATTGGAAGCAGCGGA<br>CATTTTGCCATACGAAGCCGTTGACATTTGGAATGTTACTAGAGGCTCGAGACTTACTACTTACGCCTTA<br>GCAGGAGCAAGAGGATCCGGCGTTGTTTGTATCAATGGTGCAGCTGCACACCTCAATCAACCTGGTGACT<br>TAGTGATTATTGCGACGTTCGCTGATATGGATGAAGCCGAAGCAAGAAGGCACGCTCCAAGAGTCTTGCG<br>TGTCGATGGTCAGAACAGACCATTGGCGGATCAACCACCTGAAGCCCCAGGTCCATTGGCTCCTGCAGCC<br>CCTCTTGCGGGTCAACCAGGTCCATCTCCTGAGACACCTGCTCCAAGAGCACCTAACTAGATCTAACTAA<br>TGCTTTTACTAAATATCTAG |

### 8.2. Experimental and automation workflows for new PAND discovery

For the PAND enzyme-discovery workflow, iBioGenie first generated the detailed experimental plan and organized it into eight modules (**fig. S30A-B**). BiofoundryAI then mapped the experimental operations to the available biofoundry capabilities and generated the executable workflows for each stage (**fig. S30C**). 32 candidates were codon optimized and synthesized by TWIST with overlapping sequences with backbone plasmids. Before execution, the synthesized PAND variant inserts and gel-purified pZF-TDH3p-PAND-Eno2t backbone were arranged in 96-well format, together with competent *E. coli* DH10B cells, *P. kudriavzevii* SD108ΔURA3, selective media, yeast-transformation reagents, fermentation media, and materials required for LC-MS/MS analysis. Variant identities and plate positions were defined at the beginning of the workflow and maintained through construction, transformation, fermentation, and screening. Wild-type PAND, a positive control, and an empty-vector or parent-strain background control were included as experimental references.

Module 1 prepared the DNA inputs for assembly. Synthesized PAND variant fragments and the gel-purified linearized plasmid backbone were consolidated into a 96-well source layout while maintaining the identity of each variant. Module 2 assembled each PAND insert with the corresponding plasmid backbone by HiFi assembly. Reactions were prepared in 96-well format using approximately 2 µL of insert, ~30 ng of backbone, and 5 µL of NEBuilder HiFi Master Mix in a 10 µL reaction. The assembly plate was incubated at 50 °C for 30 min before being carried into bacterial transformation (**fig. S30D**).

Module 3 transformed the assembled plasmids into chemically competent DH10B cells. The cells were incubated with the assembly products on ice, heat shocked at 42 °C, recovered in SOC medium for 1 h at 37 °C, and plated on ampicillin-containing selective medium for overnight growth at 37 °C. Module 4 recovered individual bacterial clones and prepared plasmid DNA for yeast transformation. Colonies corresponding to each construct were inoculated into 96-deepwell plates and grown overnight at 37 °C and 900 rpm while preserving variant identity. The cultures were then processed using a 96-well plasmid miniprep workflow, and the resulting plasmid DNA was collected for transformation into the production host.

Module 5 introduced the PAND constructs into *P. kudriavzevii* SD108ΔURA3. Yeast cells were grown in YPD to an OD600 of approximately 0.8–1.0 and prepared for LiAc/PEG-mediated transformation. The purified PAND plasmids were transferred into the corresponding yeast transformation positions, and transformed cells were subsequently plated on SC-URA selective medium and incubated at 30 °C until individual colonies formed. Module 6 established the biological replicates and performed plate-format fermentation for 3-HP production. Three independent yeast colonies were selected for each construct and carried forward as biological replicates. Cultures were grown in uracil-dropout medium containing 20 g/L glucose in deep-well plates at 30 °C and 900 rpm for approximately 2 days. During fermentation, 3-HP accumulated in the culture medium, allowing the extracellular supernatant to be used directly for downstream screening.

Module 7 prepared the fermentation samples and quantified secreted 3-HP by acoustic droplet ejection–open port interface mass spectrometry (ADE-OPI-MS). Cultures were centrifuged to separate the yeast cells from the product-containing supernatant, and clarified supernatants were transferred while maintaining the correspondence between fermentation wells and analytical samples. Before analysis, samples were diluted 20-fold and analyzed using an Echo® MS+ system coupled to a SCIEX ZenoTOF™ 7600 mass spectrometer. The carrier solvent consisted of 80:20 (v/v) acetonitrile containing 4 mM ammonium carbonate and 0.01% (v/v) ammonium hydroxide and was delivered at 350 µL/min. For each sample, 12.5 nL was acoustically ejected into the open port interface at 20 Hz in Wide peak mode, with an inter-sample interval of 3000 ms. Measurements were performed in negative ion mode using Zeno trap-enabled high-resolution MRM (ZenoMRMHR). 3-Hydroxypropionate (3-HP; $C_3H_6O_3$) was monitored as the $[M-H]^-$ ion at *m/z* 89.024, with the fragment ion at *m/z* 59.011 used for quantification (**fig. S33**). Source parameters were GS1 at 90 psi, GS2 at 30 psi, curtain gas at 35 psi, CAD gas at 7, source temperature at 300 °C, and spray voltage at −4500 V. The declustering potential and collision energy were −70 V and −15 V, respectively. TOF MS accumulation time was 0.05 s with Zeno pulsing enabled, and the instrument was calibrated using X500 ESI Negative Calibration Solution delivered through the Calibrant Delivery System. Module 8 processed the screening data and connected the analytical measurements back to the original variant identities. Data were processed in SCIEX OS using the Summation algorithm with an S/N threshold of 3, a Gaussian smoothing width of 1.0 point, a 1.0-s summation window, and a noise baseline set to 10% of baseline. Peak area was used for quantification with 1/x-weighted linear regression against an eight-point external calibration series containing 0, 0.001, 0.0025, 0.005, 0.01, 0.025, 0.05, and 0.1 g/L 3-HP. All calibration curves achieved $R^2 \geq 0.99$. Back-calculated concentrations were corrected for the 20-fold sample dilution before downstream analysis. The resulting 3-HP measurements were then organized by construct and replicate, summarized across the triplicate fermentations, and compared with the corresponding experimental controls to generate the final screening results used for downstream analysis and subsequent design rounds.

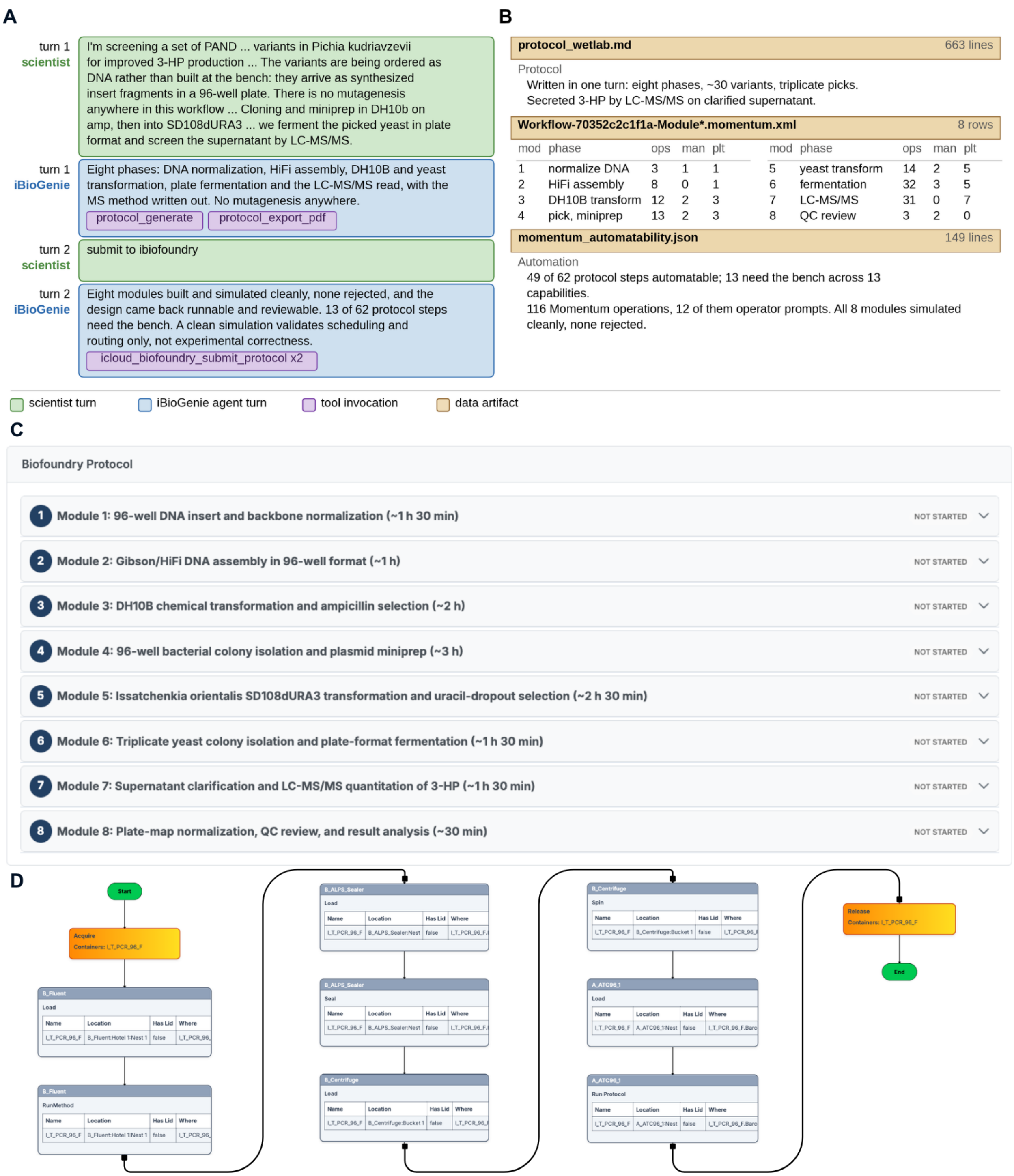


**Fig. S30. panD variants ordered as DNA, assembled and screened for 3-HP.** The variants arrive as synthesized insert fragments rather than being built at the bench, so there is no mutagenesis anywhere in this workflow: each insert is joined to a linearized backbone by HiFi assembly, carried through *E. coli* DH10B into *P. kudriavzevii* SD108dURA3, fermented in plate format, and read for secreted 3-HP by LC-MS/MS on clarified supernatant. A clean simulation validates scheduling, labware and routing, not experimental correctness. **(A)** Both exchanges, with the scientist and the agent condensed with an LLM; the submit tool shows twice because the first call errored and the second returned the build. **(B)** The delivered artifacts, all eight modules simulating cleanly with none rejected. **(C)** The same protocol in the web application as the eight modules built from it, from 96-well insert and backbone normalization through LC-MS/MS quantitation to plate-map normalization and QC review, with estimated durations. **(D)** The Momentum workflow graph for the Gibson and HiFi assembly module.

#### 8.3. Agentic AI pipeline for enzyme retrieval of aspartate decarboxylase

After identifying an improved PAND candidate enzyme through the retrieval workflow, we next asked whether the best retrieved enzyme could be further optimized by targeted protein engineering. For this second iteration, we selected *AF*panD, an aspartate 1-decarboxylase from *Aliivibrio fischeri* ES114, as the engineering target. This enzyme belongs to the group II PLP-dependent decarboxylase family, as expected. The objective of this step was to design a focused library of single amino-acid substitutions predicted to improve *AF*panD catalytic performance while maintaining protein stability and experimental feasibility for one round of SDM.

The design process began with target information extraction. The *AF*panD protein sequence and functional annotation were retrieved from UniProt, confirming its identity as a 547-aa aspartate 1-decarboxylase associated with pantothenate biosynthesis. Because no experimental structure was available for this enzyme, the corresponding AlphaFold DB model was used as the structural basis for design. The predicted model had high overall confidence, which supported its use for active-site analysis, substrate-pocket mapping, and mutation selection. Conserved PLP-dependent decarboxylase motifs were then extracted from the sequence to define the catalytic machinery.

To generate a structural context for rational design, the active-site pocket was first mapped using pocket detection and cofactor placement. fPocket analysis identified a pocket containing the key PLP-decarboxylase signature residues, establishing this region as the catalytic pocket. PLP was then placed into the AlphaFold model using a combination of docking and cofactor transplantation. Initial GNINA docking confirmed that the predicted active-site pocket could accommodate PLP with high confidence. Next, L-aspartate was docked into the PLP-bound *AF*panD structure to define substrate-contacting residues and possible activity-limiting interactions. GNINA docking was used to generate candidate L-aspartate binding poses, and the selected pose was chosen based on its compatibility with the expected Schiff-base-forming geometry between the substrate amino group and the PLP aldehyde. This analysis identified residues around the substrate amine, α-carboxylate, β-carboxylate, PLP ring, and phosphate-binding region. The final active-site context was therefore generated by integrating sequence motifs, fPocket pocket detection, GNINA cofactor and substrate docking, and AlphaFill PLP placement into a single catalytic-residue and substrate-contact map.

Using this active-site context, iBioGenie then performed mutation selection through a combination of its native rational design tool, evolutionary analysis through MSA, zero-shot sequence scoring using ESM2, and thermostability filtering. A MSA was generated from *AF*panD homologs to estimate the evolutionary tolerance of each position and identify consensus-supported substitutions. This conservation profile was used to remove highly conserved residues from consideration and to prioritize mutations where the proposed amino acid had natural precedent among homologs. In parallel, ESM2 zero-shot scoring was used to estimate whether each substitution was sequence-compatible with the broader protein context, while ThermoMPNN was used to predict the stability effect of each mutation. These filters helped remove substitutions that were strongly evolutionarily disfavored or predicted to destabilize the fold.

After candidate generation, a set of explicit selection rules was applied to produce a focused experimental panel. Candidate mutations were required to pass a stability filter based on ThermoMPNN, retaining mutations with predicted $\Delta\Delta G$ values below 1.5 kcal/mol. Highly conserved positions were removed using an MSA-based filter, excluding sites where the wild-type residue frequency exceeded 90%. ESM2 scores were used as an additional plausibility filter, retaining mutations with scores above −3. The remaining mutations were then ranked using an ensemble selection strategy that integrated mechanistic relevance, active-site shell proximity, evolutionary support, sequence-model compatibility, and predicted stability. This produced a final panel of 30 top-ranked mutations for experimental testing.

Finally, the selected mutations were converted into experimentally actionable designs by generating SDM primers. The AFpanD ORF was verified to translate exactly to the UniProt protein sequence with no positional offset, ensuring that mutation numbering in the protein sequence matched the DNA template. For each of the 30 selected substitutions, primer pairs were designed for site-directed mutagenesis, producing 30 SDM primer pairs ready for experimental construction. This completed the second iteration of the retrieval-and-engineering strategy: after retrieving a stronger PAND enzyme, iBioGenie used target annotation, structural modeling, cofactor and substrate context generation, MSA analysis, ESM2 scoring, ThermoMPNN stability prediction, rational design, ensemble ranking, and primer design to generate a focused 30-mutation *AF*panD engineering library for further improving 3HP production (**table S19 and fig. S31**). Of the 30 proposed variants, 29 were successfully constructed, tested, and yielded retrievable activity measurements.

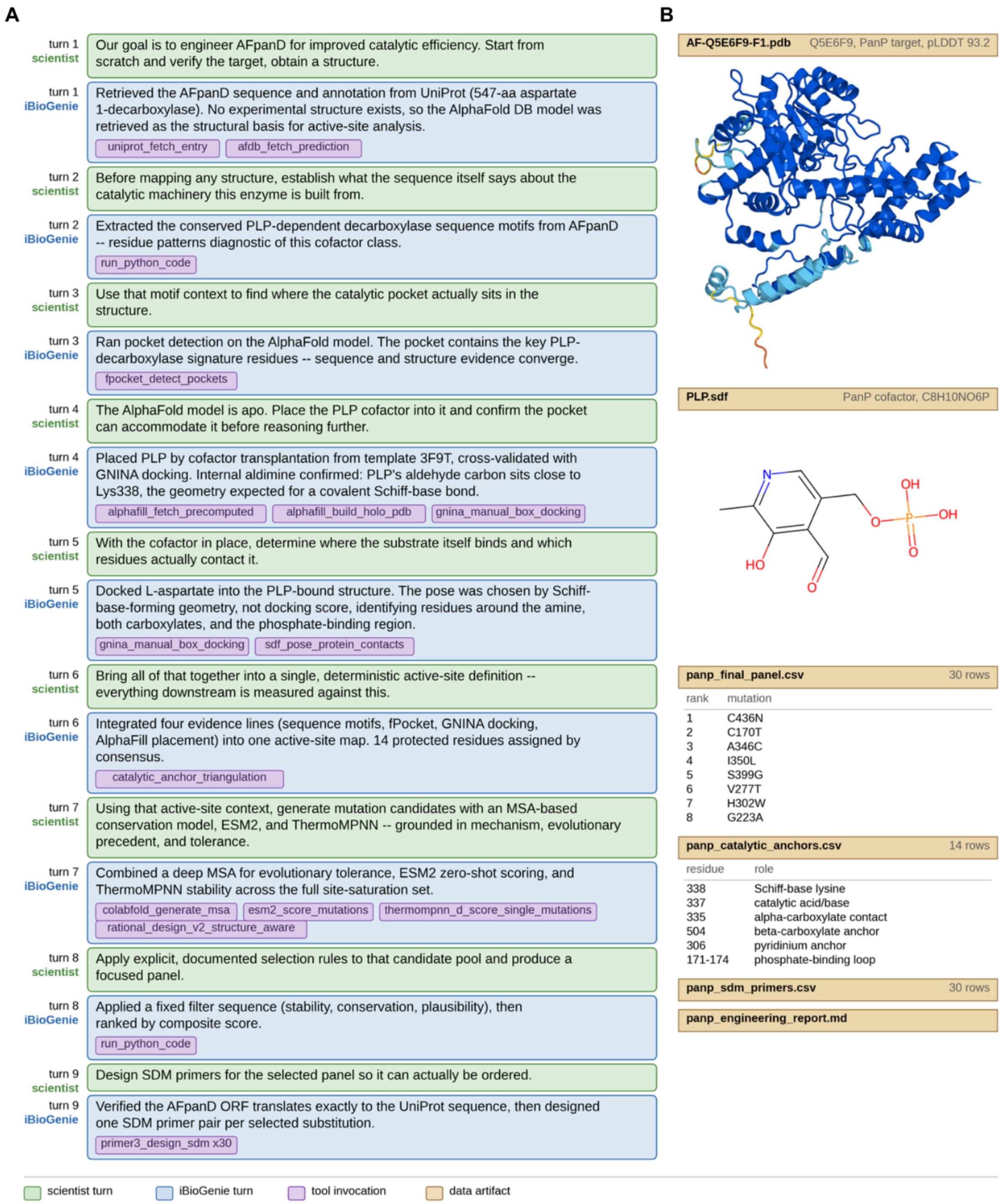


**Fig. S31. iBioGenie AfpanD engineering for improved catalytic efficiency. (A)** A summary of the scientist guided a structure- and sequence-informed engineering workflow, including conserved-motif analysis, pocket detection, PLP placement, L-aspartate docking, and integration of these signals into an active-site map with 14 protected residues. Mutation candidates were then generated using evolutionary conservation, ESM2, ThermoMPNN, and structure-aware design, filtered and ranked to a 30-variant panel, and converted into SDM primer designs. **(B)** Representative outputs include the AfPanD structure, PLP cofactor, the 29 ranked mutations, catalytic-anchor residues, SDM primers, and the final engineering report.

**Table S19. SDM design and primers for *AF*panD (Q5E6F9)**

| Variant | Forward (5' to 3') | Reverse (5' to 3') |
|---|---|---|
| C436N | CGGAACCAGAGTTAAACCTCTTAACATATAGA | TCTATATGTTAAGAGGTTTAACTCTGGTTCCG |
| C170T | TGGAGCATTTACCTCCGGTGGTACT | AGTACCACCGGAGGTAAATGCTCCA |
| A346C | TCCCAATGGGTTGCGGTATGGTTATTT | AAATAACCATACCGCAACCCATTGGGA |
| H302W | GGAAGGTTGTTGGTTCCATGTCGAC | GTCGACATGGAACCAACAACCTTCC |
| G223A | GTCGGAGAGAGCCCATTACTCGTTGA | TCAACGAGTAATGGGCTCTCTCCGAC |
| I350L | GCTGGTATGGTTCTGTTTAAAGACCCTGA | TCAGGGTCTTTAAACAGAACCATACCAGC |
| L437I | ACGGAACCAGAGTTATGTATTTTAACATATAG ATACGT | ACGTATCTATATGTTAAAATACATAACTCTGGT TCCGT |
| S399G | CAACGTGATTGGCCGTCCAGGTTAT | ATAACCTGGACGGCCAATCACGTTG |
| S285A | TGAAACAGGCGCGATTGATCCTTTG | CAAAGGATCAATCGCGCCTGTTTCA |
| V277T | TATCGTTGGTACCGCTGGTACAACTG | CAGTTGTACCAGCGGTACCAACGATA |
| T313A | GGGCGGTGCAGCGTTGATGTCGAATA | TATTCGACATCAACGCTGCACCGCCC |
| N250G | ACTGATAATAATGGTCGTGTCTGTTTAGACGA | TCGTCTAAACAGACACGACCATTATTATCAGT |
| I178L | CTATTGCCAATCTGACAGCATTGTGGG | CCCACAATGCTGTCAGATTGGCAATAG |
| D327E | CGATGGTATCGAATTGGCGGATTCTG | CAGAATCCGCCAATTCGATACCATCG |
| G276A | GGCTATCGTTGCGGTTGCTGGTACAA | TTGTACCAGCAACCGCAACGATAGCC |
| N268G | AGCTAAGGCTAAGGGTATTAAGCCTTTGG | CCAAAGGCTTAATACCCTTAGCCTTAGCT |
| M389L | TGGAATGGCTCTGCTCCTTTACTCG | CGAGTAAAGGAGCAGAGCCATTCCA |
| T78L | TTCTGAACATCTGGAACATTTGTTAGACACA | TGTGTCTAACAAATGTTCCAGATGTTCAGAA |
| V277I | GGCTATCGTTGGTATTGCTGGTACAACT | AGTTGTACCAGCAATACCAACGATAGCC |
| L437V | CGGAACCAGAGTTATGTGTGTTAACATATAGA | TCTATATGTTAACACACATAACTCTGGTTCCG |
| L438V | CCAGAGTTATGTCTCGTGACATATAGATACGT | ACGTATCTATATGTCACGAGACATAACTCTGG |
| Y392W | CTATGCTCCTTTGGTCGTGCTTCAAC | GTTGAAGCACGACCAAAGGAGCATAG |
| V342L | GCAATTATACCTGCCAATGGGTGCTG | CAGCACCCATTGGCAGGTATAATTGC |
| F395L | CTTTACTCGTGCCTGAACGTGATTTCG | CGAAATCACGTTCAGGCACGAGTAAAG |
| I175L | CGGTGGTACTCTGGCCAATATCACA | TGTGATATTGGCCAGAGTACCACCG |
| F503L | ACCTACGATTGTGCTGAGAGTTGTCTTG | CAAGACAACTCTCAGCACAATCGTAGGT |
| M344V | TATACGTCCCAGTGGGTGCTGGTATG | CATACCAGCACCCACTGGGACGTATA |
| V506I | GTTCAGAGTTATTTTGGCGAACCCATTGAC | GTCAATGGGTTCGCCAAAATAACTCTGAAC |
| M344L | TATACGTCCCACTGGGTGCTGGTATG | CATACCAGCACCCAGTGGGACGTATA |
| V342I | GCAATTATACATTCCAATGGGTGCTGGTATG | CATACCAGCACCCATTGGAATGTATAATTGC |

#### 8.4. Experimental and automation workflows for PAND engineering

For this PAND engineering workflow, iBioGenie's generated experimental plan organized the workflow into seven modules (**fig. S32A-B**). BiofoundryAI then mapped the experimental operations to the available biofoundry capabilities and generated the executable automation workflows for each stage. The workflow maintained variant identity in 96-well format from mutagenesis through fermentation and LC-MS/MS screening (**fig. S32C**). The parent pZF-TDH3p-PAND-Eno2t plasmid, mutation-specific primer plates, prepared vector backbone, *E. coli* DH10B cells, *P. kudriavzevii* SD108ΔURA3, selective media, yeast-transformation reagents, and LC-MS/MS materials were prepared before execution. Wild-type PAND, a positive-control variant, and an empty-vector control were carried alongside the engineered variants during the downstream stages.

Module 1 generated the mutation-containing PAND fragments by site-directed mutagenesis. Forward and reverse reactions were prepared separately in 96-well PCR plates using Q5 polymerase in 50 µL reactions. The PCR products were quality checked before being treated with DpnI at 37 °C to reduce carryover of the parental plasmid. Module 2 assembled the forward and reverse mutagenic fragments with the prepared vector backbone. The three DNA components were consolidated by variant and incubated at 50 °C to generate the reconstructed plasmids.

Module 3 recovered the assembled constructs in DH10B. Following transformation and recovery, cells were selected on ampicillin-containing agar and grown overnight at 37 °C. Individual colonies were then transferred into selective liquid medium while preserving the variant-to-well mapping established during mutagenesis. Module 4 purified the resulting plasmids using a 96-well miniprep workflow, including cell resuspension, lysis, neutralization, clarification, binding, washing, and elution. The purified plasmid plate then served as the DNA input for transformation of the yeast production host.

Module 5 transferred the PAND constructs into *P. kudriavzevii* SD108ΔURA3 using LiAc/PEG-mediated transformation with single-stranded carrier DNA. Purified plasmid DNA was added to the corresponding yeast transformation positions, and transformed cells were selected on SC-URA medium at 30 °C until isolated colonies formed. Module 6 established the biological replicates and performed the 3-HP production screen. Three independent yeast colonies were selected for each construct and inoculated directly into 2 mL deep-well fermentation cultures. Cultures were grown at 30 °C for approximately 2 days, allowing secreted 3-HP to accumulate in the fermentation medium.

Module 7 prepared the fermentation samples and quantified 3-HP exactly as described in Section 8.2. Cultures were centrifuged, and clarified supernatants were collected while maintaining their correspondence to the original fermentation wells. Samples were analyzed by ADE-OPI-MS using the Echo® MS+ system coupled to a SCIEX ZenoTOF™ 7600 mass spectrometer, with 3-HP measured in negative ion mode using ZenoMRMHR. The resulting measurements were linked to the corresponding PAND variant and biological replicate for downstream comparison of 3-HP production (**fig. S32D**).

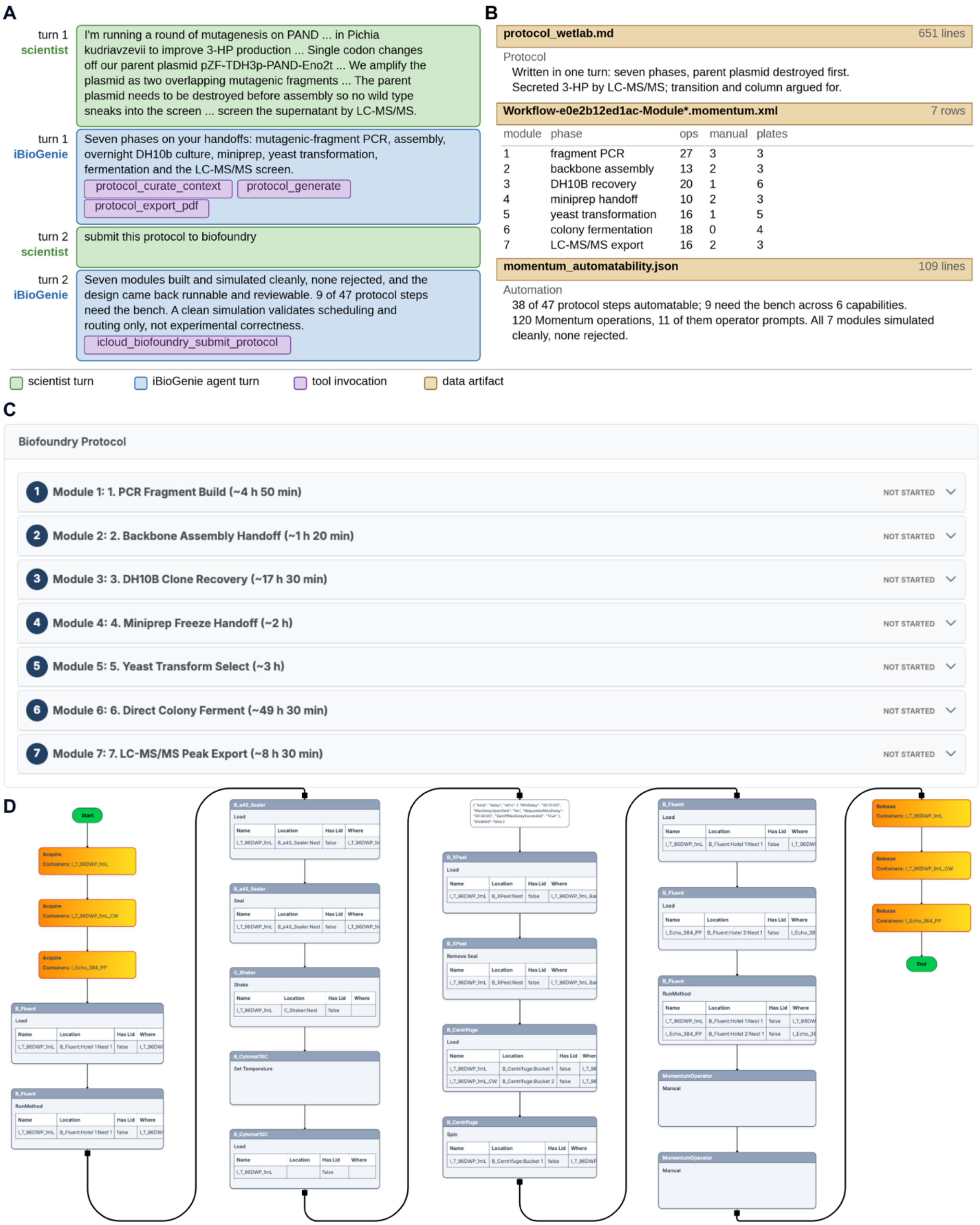


**Fig. S32. panD variants built by site-directed mutagenesis and screened for 3-HP.** The same host, product and analytical endpoint as fig. S30, reached the other way: single codon changes amplified as two overlapping mutagenic fragments off the parent plasmid, with the parent destroyed before assembly so no wild type reaches the screen. A clean simulation validates scheduling, labware and routing, not experimental correctness. **(A)** Both exchanges, with the scientist and the agent for metagenesis protocol design. **(B)** The delivered artifacts, all seven modules simulating cleanly with none rejected. **(C)** The same protocol in the web application as the seven modules built from it, from PCR fragment build through DH10B clone recovery and yeast transformation to direct colony fermentation and the LC-MS/MS peak export, with estimated durations. **(D)** The Momentum workflow graph for the LC-MS/MS peak export module, showing plate sealing and peeling, centrifugation, the liquid handler steps and the operator prompts that bracket the instrument run.

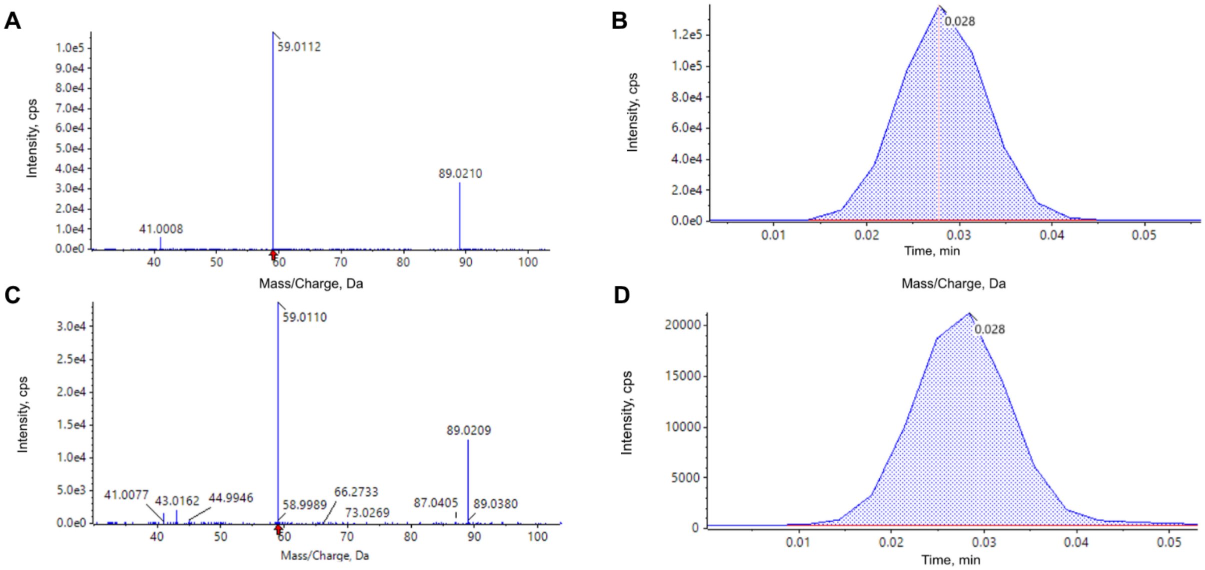


**Fig. S33. LC-MS/MS confirmation and quantitation of secreted 3-HP.** All four panels are negative-ion acquisitions of the same analyte. **(A)** Mass spectrum of standard of 0.1 g/L of 3-HP with the base peak at m/z 59.011 and the m/z 89.021 ion at roughly a third of its intensity, and a red marker at m/z 59.011. **(B)** Extracted-ion chromatogram for the standard, peaking at 0.028 min. **(C)** The same acquisition for an example sample. **(D)** Extracted-ion chromatogram for one sample.

### 9. Case study III: Metabolic engineering of *Saccharomyces cerevisiae* for β-carotene production through combinatorial gene knockout

#### 9.1. Agentic AI pipeline for gene knockout selection

Starting from base strain CEN.PK113-7D carrying the integrated β-carotene pathway and the additional ΔCIT2 deletion, iBioGenie was instructed to find further gene deletions for improving β-carotene production. It ran four classical strain-design methods, FSEOF, MCS, OptKnock and RobustKnock (*11*), against Yeast9 v9.0.2 + Crt, a machine-learning method, flux cone learning (*12*), and it searched a curated literature library together with SI data from previous β-carotene studies (*13*), with single additional deletions imposed as a constraint of the automated gene knockout protocol. MCS, OptKnock and RobustKnock returned zero feasible single-deletion interventions. The issue with these methods for our single additional gene deletion scenario is that they enforce growth coupling, and with ΔCIT2 already in place no single deletion on Yeast9 + Crt achieves it: the exhaustive single-deletion search ran to completion and returned nothing. Larger intervention sets were not searched, so what puts growth coupling out of reach is the one-deletion budget rather than the network alone. Flux cone learning collapsed to majority-class prediction on the Özaydın visually labeled β-carotene data, reaching 0.35 held-out balanced accuracy against a 0.33 random baseline and predicting the majority class for about 83% of test samples so it was discarded (*12*). Candidate deletions were then filtered through SGD gene essentiality and FVA feasibility at biomass ≥ 1.3. The final gene list contained 14 unique gene knockouts applied to the ΔCIT2 strain: 2 Özaydın visual screen picks, 2 FSEOF picks, 8 multi-evidence literature picks, and 2 non-GEM regulators (**table S20**). 12 of the 14 have literature evidence for carotenoid or terpenoid improvement in wild-type strains, indicating that multi-method candidate selection favors the literature route under this strain-construction scenario. One target, IFA38, was carried forward precisely because its evidence was contradictory. The model-based arms supported it, with the strongest flux-sampling signal of any candidate, while the visual screen scored it as decreased, so building it could directly test the model prediction against the screen. The strain could not be constructed after successful gRNA identification, so we could not evaluate the method disagreement. SaCas9 gRNAs for full ORF deletions were designed over two rounds. Round 1 took the top three guides per gene by score from the deletion-gRNA library of a previous publication (*14*); round 2 used CHOPCHOP for the eight genes that round 1 did not construct (*15*). Overlapping sense and antisense primers for homologous-recombination donors were designed with Primer3 (**fig. S35**).

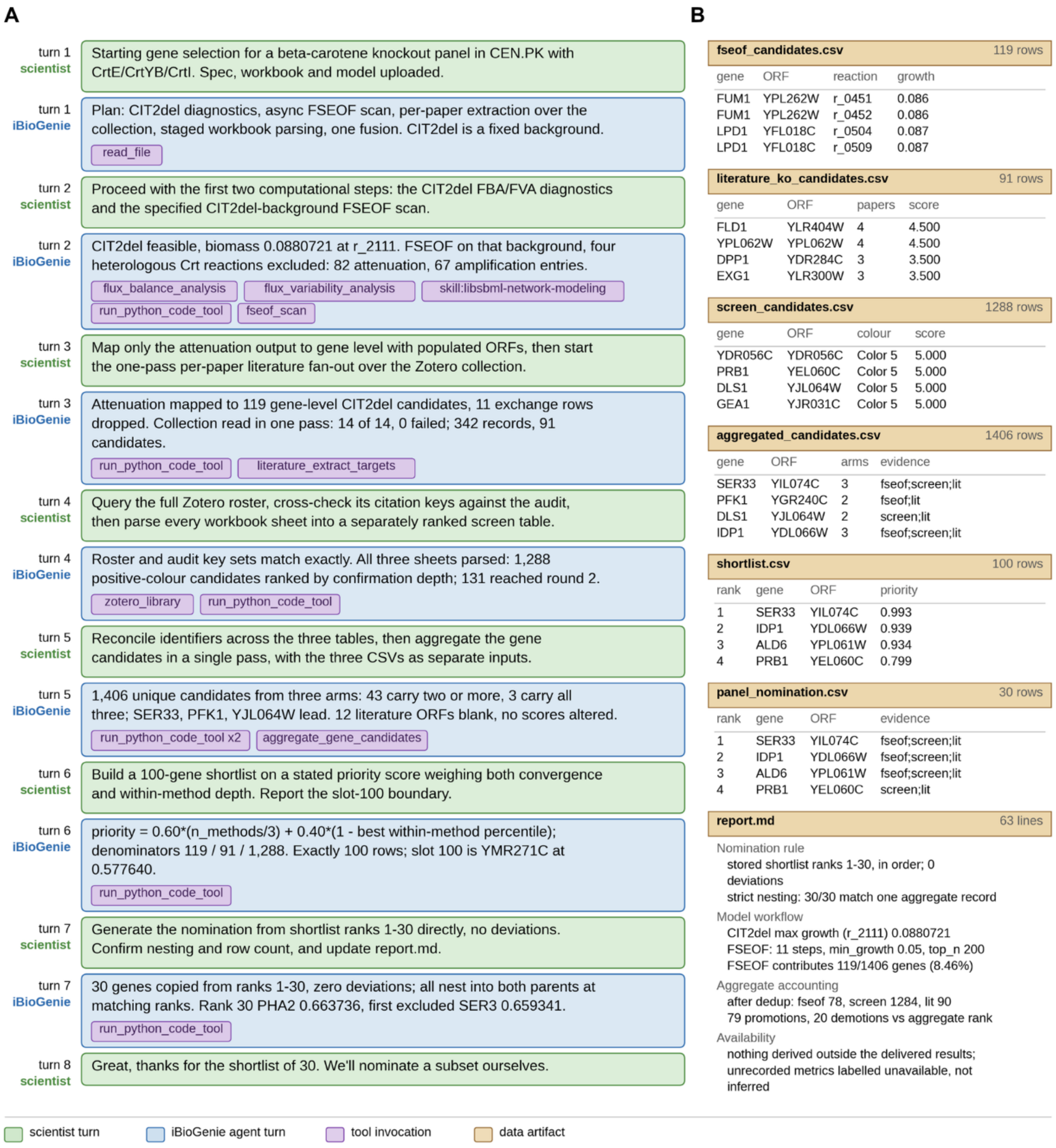

**Fig. S34. iBioGenie gene knockout selection for β-carotene production.** (**A**) A summary of the scientist-guided session that ranks knockout candidates for the 14-gene *S. cerevisiae* CEN.PK β-carotene panel. All seven exchanges and a closing turn are shown; the scientist gates each step, from an FSEOF scan on a fixed ΔCIT2 background through a literature fan-out and workbook parsing to aggregation of genes from different methods. (**B**) Representative outputs include the seven delivered files with their leading rows and true row counts: 119 FSEOF, 91 literature and 1,288 screen candidates aggregate to 1,406 unique genes, which a stated priority score cuts to a 100-gene shortlist and then to a 30-gene nomination. A similar session produced the 14 genes that were built.

Because on-target gRNA activity deviates from computational prediction, each gRNA was evaluated experimentally in a killing assay rather than accepted on its predicted score. iBioGenie carried out design and screening over two rounds: round 1 screened 42 guides, 3 per gene across all 14 genes, and round 2 screened 32 guides, 4 per gene for the 8 genes whose round-1 guides all failed. The six genes for which round 1 already yielded an efficient guide were not carried into round 2, and none of the round-2 gRNAs had been screened for those genes in round 1. Across the 74 guides screened, 21 were efficient and collectively covered all 14 genes (**figs. S34-S35, tables S21 to S24**).

Chaining a design workflow across model construction, flux analysis, literature extraction, screen parsing, scoring, and final human selection makes full list reproducibility difficult because each step produces stochastic model output, and small differences can compound (**fig. S36A**). To address reproducibility of the gene knockout list, we took the original sessions that generated the 14-gene candidate list (**table S20**) and converted the trajectory into a single starting prompt for the simulated researcher. We analyzed 21 completed replicate trajectories using the final tools from the original list generation and assessed where the 14 genes fall across the 21 ranked lists. Twelve of the 14 genes recover within the top 110 candidates, and seven recover within the top 14. The two that do not are exactly the two derived from flux analysis rather than from the literature or the screen, and they fail in different ways (**fig. S36B**). ALD6 appears in every replicate list but at a median rank of 318. IFA38, the contradictory pick built to test the model-based prediction against the Özaydın screen, is absent from every replicate list. Its absence traces to a change in the default solver in FSEOF. Even under the original solver, IFA38's flux correlation fell inside a 201-way tie, so its rank carried no ranking information. The FSEOF slot was filled by ranking candidates on an FVA ceiling shift that was numerically zero for every tied candidate, so the choice among them was not made on signal; IFA38 was then retained as a deliberate test case, adding one further FSEOF-derived gene whose prediction contradicted the ordinal visual screen of Özaydın et al. The reproducibility study suggests that designs arrived at through multiturn chats can be evaluated with replicate trajectories to ensure original designs are not hypersensitive to stochastic model trajectories (**fig. S36**).

**Table S20. List of genes for β-carotene production.** Fourteen single-gene knockout targets, each designed on the ΔCIT2 base as 14 strains. Twelve were selected from prior carotenoid and terpenoid engineering literature and two from an FSEOF scan. Yeast9 records whether the gene is in the Yeast9 genome-scale model, so that flux predictions were available for it. Confidence is HIGH for three or more independent supporting lines, MED for two, and LOW for one line plus a biological rationale; conflict marks opposing signals treated as a model-resolution experiment. DLS1 is the gene built at the locus the selection pipeline nominates as the overlapping dubious ORF YJL064W and is listed here under its own name and systematic ORF.

| # | Gene | ORF | Source | Yeast9 | Confidence |
|---|---|---|---|---|---|
| 1 | LAC1 | YKL008C | Lit | yes | HIGH |
| 2 | ROX1 | YPR065W | Lit | no | HIGH |
| 3 | PFK1 | YGR240C | Lit | yes | HIGH |
| 4 | EXG1 | YLR300W | Lit | yes | HIGH |
| 5 | PAH1 | YMR165C | Lit | yes | HIGH |
| 6 | DLS1 | YJL065C | Lit | no | MED |

| # | Gene | ORF | Source | Yeast9 | Confidence |
|---|---|---|---|---|---|
| 7 | SOL1 | YNR034W | Lit | no | MED |
| 8 | SER33 | YIL074C | Lit | yes | MED |
| 9 | PCT1 | YGR202C | Lit | yes | MED |
| 10 | HXK2 | YGL253W | Lit | yes | MED |
| 11 | GEA1 | YJR031C | Lit | no | LOW |
| 12 | VPS1 | YKR001C | Lit | no | LOW |
| 13 | IFA38 | YBR159W | FSEOF | yes | HIGH (conflict) |
| 14 | ALD6 | YPL061W | FSEOF | yes | HIGH (mech) |

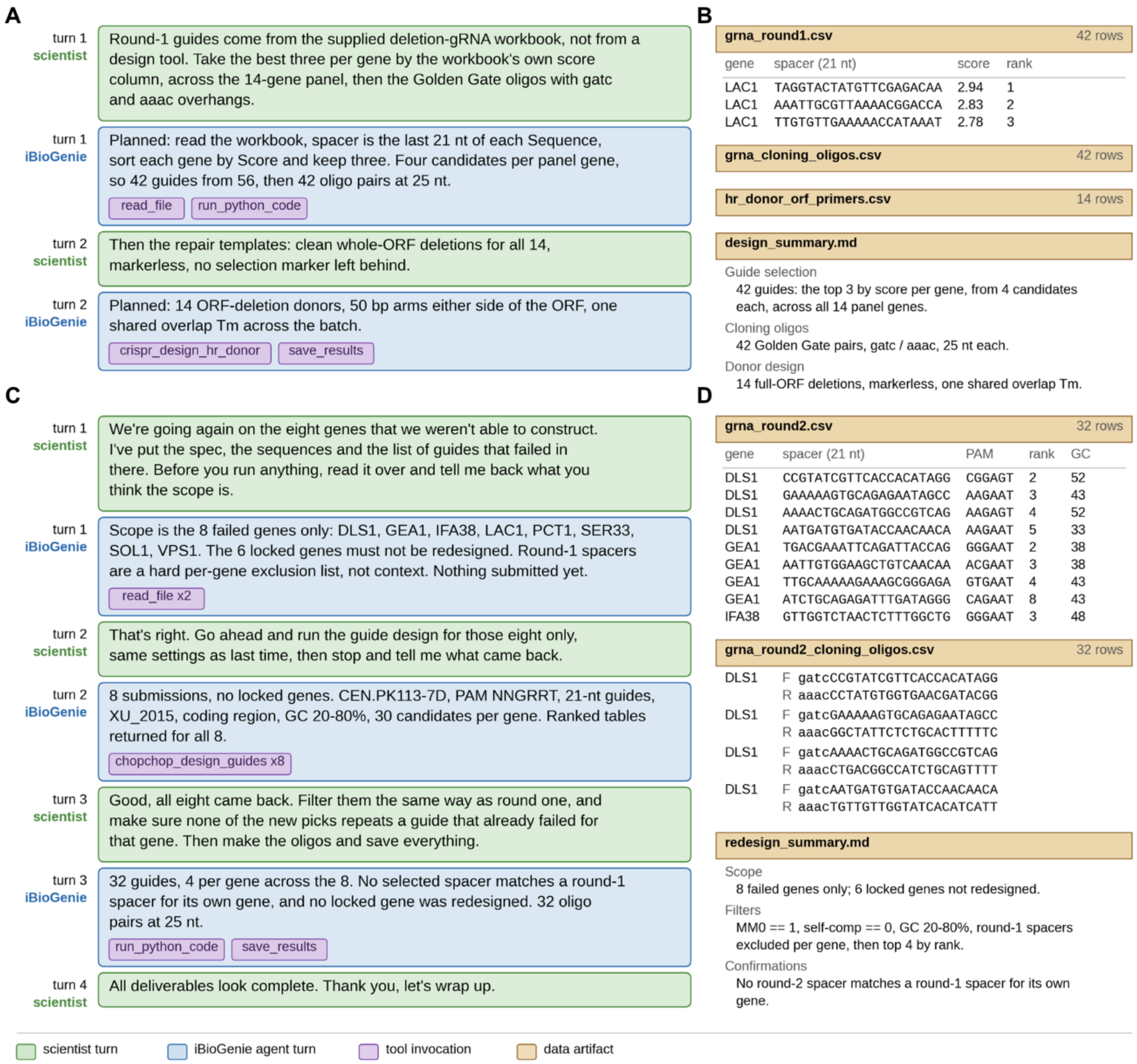


**Fig. S35. iBioGenie gRNA design for the β-carotene panel, rounds 1 and 2.** A summary for the 14-gene panel in one figure (**A**) A summary of round-1 guide selection. Guides are not designed de novo but taken from the supplied deletion-gRNA workbook of a previous publication, the best three per gene by its own score column. (**B**) The round-1 artifacts: 42 guides, 42 Golden Gate oligo pairs carrying gatc and aaac overhangs, and 14 full-ORF-deletion donors, the 42 forward oligos reproducing **table S21** row for row. (**C**) A summary of round-2 redesign session, all four exchanges, covering the eight genes the panel could not construct. (**D**) Its delivered files: 32 guides, four per gene across the eight, and their 32 cloning oligo pairs.

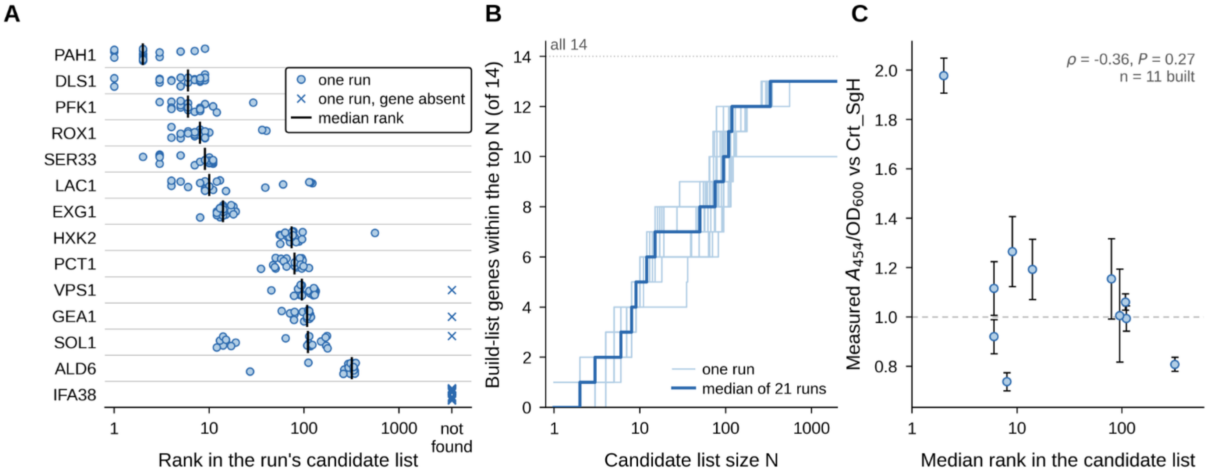


**Fig. S36. Reproducibility of gene selection for the β-carotene case study.** One prompt posed the original 14-gene *S. cerevisiae* knockout selection task to the simulated researcher, with added detail directing iBioGenie to the same tools. (**A**) Rank of each gene in each run on a log axis, the median rank as a black tick and a cross where a run's list lacks the gene, genes ordered by median rank. Seven recover at median rank 2-14 (PAH1, DLS1, PFK1, ROX1, SER33, LAC1, EXG1), five at 74-110 (HXK2, PCT1, VPS1, GEA1, SOL1), ALD6 at 318, and IFA38 is never recovered; IFA38 was chosen for its disagreement between FSEOF and the literature (12). DLS1 is the gene built at the locus nominated as the overlapping dubious ORF YJL064W and is counted under either identifier. (**B**) Build-list genes within the top N candidates, one curve per run and the median: 7 at N = 30, 10 at N = 100, 12 at N = 300, 13 at N = 1000. IFA38 is always absent. (**C**) Measured β-carotene against predicted rank for the eleven genes built, as A454/OD600 relative to the Crt_SgH reference (dashed line), error bars the SD across replicates and rank taken from the medians in (A). The trend runs in the expected direction but is not significant at this size (Spearman ρ = -0.36, P = 0.27, n = 11). LAC1, HXK2 and IFA38 failed construction at transformation and do not appear.

**Table S21. first round of gRNA design for 14 genes.**

| Well Position | Name | Forward (5' to 3') | Reverse (5' to 3') |
|---|---|---|---|
| A1 | LAC1_Sg1 | gatcTAGGTACTATGTTCGAGACAA | aaacTTGTCTCGAACATAGTACCTA |
| B1 | LAC1_Sg2 | gatcAAATTGCGTTAAAACGGACCA | aaacTGGTCCGTTTTAACGCAATTT |
| C1 | LAC1_Sg3 | gatcTTGTGTTGAAAAACCATAAAT | aaacATTTATGGTTTTTCAACACAA |
| D1 | ROX1_Sg1 | gatcAACGCAGTTAGACAATCAACA | aaacTGTTGATTGTCTAACTGCGTT |
| E1 | ROX1_Sg2 | gatcTCTGTTCAGACAGCACTACCA | aaacTGGTAGTGCTGTCTGAACAGA |
| F1 | ROX1_Sg3 | gatcATCTGTTGTTCTCCAGTACAG | aaacCTGTACTGGAGAACAACAGAT |
| G1 | PFK1_Sg1 | gatcTCTATGGAATTCAGAAAGCGT | aaacACGCTTTCTGAATTCCATAGA |
| H1 | PFK1_Sg2 | gatcCTTAACAGCATCGACACCTTG | aaacCAAGGTGTCGATGCTGTTAAG |
| A2 | PFK1_Sg3 | gatcGATTGGAGCAGACATCACTAA | aaacTTAGTGATGTCTGCTCCAATC |
| B2 | EXG1_Sg1 | gatcTTGGAAAGCCCAGTAACCGAT | aaacATCGGTTACTGGGCTTTCCAA |
| C2 | EXG1_Sg2 | gatcCACTTCTGTCAATATTTAGGT | aaacACCTAAATATTGACAGAAGTG |
| D2 | EXG1_Sg3 | gatcTACAAATGATGACAACGACGA | aaacTCGTCGTTGTCATCATTTGTA |
| E2 | PAH1_Sg1 | gatcGCTGGACAAGCTGATTCCACG | aaacCGTGGAATCAGCTTGTCCAGC |
| F2 | PAH1_Sg2 | gatcTTGATCGCAAATAACTCCTCG | aaacCGAGGAGTTATTTGCGATCAA |
| G2 | PAH1_Sg3 | gatcTGCAGTACGTAGGCAGAGCTC | aaacGAGCTCTGCCTACGTACTGCA |
| H2 | SOL1_Sg1 | gatcCGTTAATCCAAGTGTTGCACG | aaacCGTGCAACACTTGGATTAACG |

| Well Position | Name | Forward (5' to 3') | Reverse (5' to 3') |
|---|---|---|---|
| A3 | SOL1_Sg2 | gatcCACAGAATCCAAAACGATGGA | aaacTCCATCGTTTTGGATTCTGTG |
| B3 | SOL1_Sg3 | gatcTGCACCCAGCGGACCTTCGAA | aaacTTCGAAGGTCCGCTGGGTGCA |
| C3 | SER33_Sg1 | gatcTCACCAATTACCAATTCTGCT | aaacAGCAGAATTGGTAATTGGTGA |
| D3 | SER33_Sg2 | gatcCCAGAAAAGTTCATGGCACGT | aaacACGTGCCATGAACTTTTCTGG |
| E3 | SER33_Sg3 | gatcGCAATCAAAATCTTCAAGGAT | aaacATCCTTGAAGATTTTGATTGC |
| F3 | PCT1_Sg1 | gatcTGAAACTTTAACGCACTGCAG | aaacCTGCAGTGCGTTAAAGTTTCA |
| G3 | PCT1_Sg2 | gatcAGAAGGAGGCTCGTTACACCA | aaacTGGTGTAACGAGCCTCCTTCT |
| H3 | PCT1_Sg3 | gatcGATAAGGATGAGAGTAAGAAC | aaacGTTCTTACTCTCATCCTTATC |
| A4 | HXK2_Sg1 | gatcTACACTGACCCAGAAACTAAG | aaacCTTAGTTTCTGGGTCAGTGTA |
| B4 | HXK2_Sg2 | gatcCAATAAATTCCCACAATTCGT | aaacACGAATTGTGGGAATTTATTG |
| C4 | HXK2_Sg3 | gatcTGATTTGGGTGGTACCAACTT | aaacAAGTTGGTACCACCCAAATCA |
| D4 | GEA1_Sg1 | gatcCGAAAATGTCGAGCCAAATTA | aaacTAATTTGGCTCGACATTTTCG |
| E4 | GEA1_Sg2 | gatcAAAGTTGCAAAAAGAAAGCGG | aaacCCGCTTTCTTTTTGCAACTTT |
| F4 | GEA1_Sg3 | gatcACGACTACCTCTGATAACGAT | aaacATCGTTATCAGAGGTAGTCGT |
| G4 | VPS1_Sg1 | gatcACGTCAAAGTAAGAACATGCG | aaacCGCATGTTCTTACTTTGACGT |
| H4 | VPS1_Sg2 | gatcACGATGATGACAAGAAAAAGG | aaacCCTTTTTCTTGTCATCATCGT |
| A5 | VPS1_Sg3 | gatcTTCTTCTGTGCCCATTAACTT | aaacAAGTTAATGGGCACAGAAGAA |
| B5 | IFA38_Sg1 | gatcATTCTAGCCATTGACATTGCG | aaacCGCAATGTCAATGGCTAGAAT |
| C5 | IFA38_Sg2 | gatcGTGAAAGCTGAAAACAAGAAG | aaacCTTCTTGTTTTCAGCTTTCAC |
| D5 | IFA38_Sg3 | gatcAACTGCGCACACAATTCCTTA | aaacTAAGGAATTGTGTGCGCAGTT |
| E5 | ALD6_Sg1 | gatcGTCTACTAAGTAAGTTGGCTG | aaacCAGCCAACTTACTTAGTAGAC |
| F5 | ALD6_Sg2 | gatcAACGATGTTGACGACACCAGC | aaacGCTGGTGTCGTCAACATCGTT |
| G5 | ALD6_Sg3 | gatcGCATTTAAAGGTGTGACAGCA | aaacTGCTGTCACACCTTTAAATGC |
| H5 | DLS1_Sg1 | gatcCTGACGGCCATCTGCAGTTT | aaacAAACTGCAGATGGCCGTCAG |
| A6 | DLS1_Sg2 | gatcCTTCGTCCAGGTGCTGACAC | aaacGTGTCAGCACCTGGACGAAG |
| B6 | DLS1_Sg3 | gatcATCCGAAGTGTCCATATATT | aaacAATATATGGACACTTCGGAT |

**Table S22. Second round of gRNA design for 8 genes.**

| Well Position | Name | Forward (5' to 3') | Reverse (5' to 3') |
|---|---|---|---|
| A1 | LAC1_Sg1 | gatcACATCCATTAGGAATTCACGT | aaacACGTGAATTCCTAATGGATGT |
| B1 | LAC1_Sg2 | gatcGTTACACGTCACTTCTAAGCA | aaacTGCTTAGAAGTGACGTGTAAC |
| C1 | LAC1_Sg3 | gatcGCATGACGGTAACTTATCTCT | aaacAGAGATAAGTTACCGTCATGC |
| D1 | LAC1_Sg4 | gatcGGAAATCCAACATTTATATTG | aaacCAATATAAATGTTGGATTTCC |
| E1 | GEA1_Sg1 | gatcTGACGAAATTCAGATTACCAG | aaacCTGGTAATCTGAATTTCGTCA |
| F1 | GEA1_Sg2 | gatcAATTGTGGAAGCTGTCAACAA | aaacTTGTTGACAGCTTCCACAATT |
| G1 | GEA1_Sg3 | gatcCTTGCAACTGCCTTTCCACCG | aaacCGGTGGAAAGGCAGTTGCAAG |
| H1 | GEA1_Sg4 | gatcTTGCAAAAAGAAAGCGGGAGA | aaacTCTCCCGCTTTCTTTTTGCAA |
| A2 | PCT1_Sg1 | gatcTGTCATCCGGTGATTTAGCGG | aaacCCGCTAAATCACCGGATGACA |
| B2 | PCT1_Sg2 | gatcCTCCGGCTACAGAGTTTGCCA | aaacTGGCAAACTCTGTAGCCGGAG |
| C2 | PCT1_Sg3 | gatcTGGGAGGCGTTGAGTTAGTGT | aaacACACTAACTCAACGCCTCCCA |
| D2 | PCT1_Sg4 | gatcGCGTTGAGTTAGTGTTTGAAT | aaacATTCAAACACTAACTCAACGC |

| Well Position | Name | Forward (5' to 3') | Reverse (5' to 3') |
|---|---|---|---|
| E2 | SOL1_Sg1 | gatcCTTAAGGGCAGAGTTTTGTGC | aaacGCACAAAACTCTGCCCTTAAG |
| F2 | SOL1_Sg2 | gatcCAATAATGGAAAGGCCTGAAA | aaacTTTCAGGCCTTTCCATTATTG |
| G2 | SOL1_Sg3 | gatcGCCTTAACGGAAATGGTAACA | aaacTGTTACCATTTCCGTTAAGGC |
| H2 | SOL1_Sg4 | gatcTCGTTATTCCCCAATTTTCAA | aaacTTGAAAATTGGGGAATAACGA |
| A3 | DLS1_Sg1 | gatcCCGTATCGTTCACCACATAGG | aaacCCTATGTGGTGAACGATACGG |
| B3 | DLS1_Sg2 | gatcGAAAAAGTGCAGAGAATAGCC | aaacGGCTATTCTCTGCACTTTTTC |
| C3 | DLS1_Sg3 | gatcAAAACTGCAGATGGCCGTCAG | aaacCTGACGGCCATCTGCAGTTTT |
| D3 | DLS1_Sg4 | gatcAATGATGTGATACCAACAACA | aaacTGTTGTTGGTATCACATCATT |
| E3 | IFA38_Sg1 | gatcGTTGGTCTAACTCTTTGGCTG | aaacCAGCCAAAGAGTTAGACCAAC |
| F3 | IFA38_Sg2 | gatcCTTTAGATAATTCCCCAGCCA | aaacTGGCTGGGGAATTATCTAAAG |
| G3 | IFA38_Sg3 | gatcTAATTCTGATAGATTTATGGA | aaacTCCATAAATCTATCAGAATTA |
| H3 | IFA38_Sg4 | gatcACCACTGTATGTAGCCAAAAG | aaacCTTTTGGCTACATACAGTGGT |
| A4 | SER33_Sg1 | gatcAATCTTGTAAATTATCGGCAG | aaacCTGCCGATAATTTACAAGATT |
| B4 | SER33_Sg2 | gatcACAAGTCTTCTCTACCTGAGG | aaacCCTCAGGTAGAGAAGACTTGT |
| C4 | SER33_Sg3 | gatcTCAAGGATCAGGGTTACCAAG | aaacCTTGGTAACCCTGATCCTTGA |
| D4 | SER33_Sg4 | gatcTGCATTGACATTTTCCAACAG | aaacCTGTTGGAAAATGTCAATGCA |
| E4 | VPS1_Sg1 | gatcGACAATCTGAAGACAATAAAG | aaacCTTTATTGTCTTCAGATTGTC |
| F4 | VPS1_Sg2 | gatcTGCTCAAAACAACCGAACTAG | aaacCTAGTTCGGTTGTTTTGAGCA |
| G4 | VPS1_Sg3 | gatcTGTCATAGATATTTTGGCTGG | aaacCCAGCCAAAATATCTATGACA |
| H4 | VPS1_Sg4 | gatcAAGACAATAAAGAGGAATGGG | aaacCCCATTCCTCTTTATTGTCTT |

### Table S23. Overlap PCR primers for 14 genes.

| Forward primer (F) | Sequence | Reverse primer (R) | Sequence |
|---|---|---|---|
| ALD6_KO_donor_F | AAAACATCAAGAAACATCTTTAACATACACAAACACATACTATCAGAATAGTACCAACCT | ALD6_KO_donor_R | TTATATGAAAGTATTTTGTGTATATGACGGAAAGAAATGCAGGTTGGTACTATTCTGATA |
| EXG1_KO_donor_F | TGCCTTTGATTTTTTGTTTACTTTCTTTTTCTAGTTAATTACCAACTAAAGCTCATTTTA | EXG1_KO_donor_R | TTAAGTTTTGCTGATGAAAAATAACATTAGAAAATTCAGCTAAAATGAGCTTTAGTTGGT |
| GEA1_KO_donor_F | CGCAACTTTGGTAAGAAGCGCCCAGGTTTAACTAATACGGCTTCATATACACCAACGATA | GEA1_KO_donor_R | AAAAAATTCCATCATACTAAGTTTCTTGGTATAGTTACATTATCGTTGGTGTATATGAAG |
| HXK2_KO_donor_F | TTTGAAAAGGTTGTAGGAATATAATTCTCCACACATAATAAGTACGCTAACTTAATTTGT | HXK2_KO_donor_R | AGTAGAAAAAGGGCACCTTCTTGTTGTTCAAACTTAATTTACAAATTAAGTTAGCGTACT |
| IFA38_KO_donor_F | ATAGAAGAACAGTCTTCAAAGCAATCAATTCGTGTCGGTGCAGCTTAAGATTATCGTTCT | IFA38_KO_donor_R | ATATATATATATATATATGTATTGTATAAAAACTATCTCGAGAACGATAATCTTAAGCTG |
| LAC1_KO_donor_F | TTATTATTCATACCTCCGGTAAACATTTAGATAGACACAGTATCAATAAAGGAAAACGGT | LAC1_KO_donor_R | TGTAAAGAATTAATGTGTAATGGTTATACTACTTAAAAACACCGTTTTCCTTTATTGATA |
| PAH1_KO_donor_F | TTTTACCTTCTAAGAAACATACAGGGAAGAAATTACTGAAGATAGACACAAGATTCTTGT | PAH1_KO_donor_R | CTTAATATGCAGTATGGATCGTTATAAATAATATTCGGCTACAAGAATCTTGTGTCTATC |

| Forward primer (F) | Sequence | Reverse primer (R) | Sequence |
|---|---|---|---|
| PCT1_KO_donor_F | CTCTTGCTATTCTCTCTTGGTAGTACAAAGCACAGCAGACACAAAGAAAATCCAATCTCT | PCT1_KO_donor_R | ACAAAAAGGGTTGGTTGAAGGGAGAGAGAAAGAGACGCACAGAGATTGGATTTTCTTTGT |
| PFK1_KO_donor_F | TATTTGGGAAAGCTTTTATATAAAAAATCTGAAACAAAATCATATCAAAGGATTGCAATG | PFK1_KO_donor_R | GCATGCCATTTTTACCTCCTTTTGCTTAACTTAAACTTTTCATTGCAATCCTTTGATATG |
| ROX1_KO_donor_F | GAAAATACTAATACTTCTTCACACAAAAGAACGCAGTTAGACAATCAACATTTTTTTCCA | ROX1_KO_donor_R | AAGGGAATATAGTATAATATAATATAACGGAAAGAAGAAATGGAAAAAAATGTTGATTGT |
| SER33_KO_donor_F | TTGTAACATTAAAAAGTAACAAACACTGATTTCGGGTATTTCCTCCCTAATCAGATAAAT | SER33_KO_donor_R | TGAAGATAGGGGAGGATTTTAAATCTATACATATATTTTTATTTATCTGATTAGGGAGGA |
| SOL1_KO_donor_F | ATAAAAGAGGCCCGAGAAAATAGATTTTAACTGGAATAATTTTTTGCAAGGTGTTTAATT | SOL1_KO_donor_R | ATTAGTTGTCATTATCATCAGAAAGAAAAAGAACCCAAAAAATTAAACACCTTGCAAAAA |
| VPS1_KO_donor_F | ACCAAAATAAGGACCGTACGAAAACTGCACATTTTATATTATCAGATATCTTCCTCATCT | VPS1_KO_donor_R | AGGGAGAAATACTCAAAACCAAGCTTGAGTCGACCGGTATAGATGAGGAAGATATCTGAT |
| DLS1_KO_donor_F | TGGGCTTCGGAGCCGTATCGTTCACCACATAGGCGGAGTAAACTTCATTATCACATCGAG | DLS1_KO_donor_R | AAACAGAAGAAACAAGAGAGAATAGCGTCAGGATAGCTCGCTCGATGTGATAATGAAGTT |

**Table S24. Colony PCR primers used to genotype the 14 gene knockouts.**

| KO Gene | Forward (5' to 3') | Reverse (5' to 3') |
|---|---|---|
| ALD6 | CCATGTGGGCAAATTCGTAA | CTAAAAGCGGTCCACGTTAG |
| EXG1 | CGTCATATGAGCAGTTGCAG | TTTTATGCAAGACGACACGG |
| GEA1 | CTTGAATCATCGCGTTGGAT | CCGTCTCCCACACAATACTA |
| HXK2 | GACCACGAAACACCTTTGAT | AGAGGAAGTGTAGAGAGGGT |
| IFA38 | CTTGAAATTCCAGAAGGGCC | TCTGGGCTATTTTCCACTGT |
| LAC1 | TATTGTTTCCCTCCTTGGCT | ACGATAGCTCCACTGTTGAA |
| PAH1 | GGTTTCCCTCCCTTTAGACA | GCATTGTATTTCGGCATTGC |
| PCT1 | TACCCGTATTTGTCGTTTGC | GCTTCAAGGAACGCACTAAA |
| PFK1 | CGACTTGCATCAACCTTTGA | AATCAAGGCGCCATTATTCC |
| ROX1 | TCTTCGTTTTCACTTTGCGT | CTGCGCCTAACACCAATATC |
| SER33 | GTACCAGGAAGCGTGATAGA | CTGGGTGGTCTTTTACTGGA |
| SOL1 | ACAACAAGTTGGAGAGCAAC | AGCATGCTTTTCTCTTAGCG |
| VPS1 | AATGCTCGAGGGTAAACCA | ACGTTCGGAAATCATGAACA |
| DLS1 | TAAACGGTTTGTCGGTTTCC | GTCCCTTCCCTATGATGAGG |

#### 9.2. Experimental and automation workflows for gene knockout selection

The full protocol was run as an integrated liquid-handling workflow on a Tecan Fluent deck coupled to an Echo acoustic dispenser, an on-deck thermocycler, a Cytomat automated incubator, an automated centrifuge, and a PIXL colony picker, with plate sealing and transfers handled by the platform's robotic arm; all steps were carried out in 96-well/384-well format with well-position mapping preserved end to end.

The campaign was organized into two module sets, one for building and screening the guide plasmids and one for constructing and assaying the knockouts. Each module was sized to finish within one working day, and adjacent modules were linked through plate handoffs rather than run as a single continuous process, so a failed step could be repeated without restarting the campaign. BiofoundryAI generated each module as a simulation-tested workflow and scheduled execution on the biofoundry through Momentum and the main robotic arm. Durations given below are the hands-on runtime of each module and exclude unattended incubation.

**Guide-plasmid assembly and screening**

The full β-carotene knockout screen was executed as an integrated liquid-handling workflow on a Tecan Fluent deck coupled to an Echo acoustic dispenser, an automated thermocycler, a Cytomat incubator, an Agilent automated centrifuge, a PIXL colony picker and a Tecan Infinite plate reader, with plate sealing, unsealing and inter-device transfers handled by the platform's robotic arm. All steps were carried out in 96-well and 384-well format, and well-position mapping, and reformatting was preserved end-to-end, so that a given guide, plasmid, and occupy the same address. The host was an *S. cerevisiae* CEN.PK strain carrying the crtE/crtYB/crtI pathway, together with a constitutively expressed SaCas9 and a KanMX (G418) marker integrated into the genome. The guide-RNA plasmid carried URA3 for yeast selection and ampicillin resistance for E. coli; paired BsaI recognition sites sit immediately upstream of the SaSgRNA scaffold, between the SNR52 promoter and the SUP4 terminator. Round 1 screened 42 guides against a 14-gene panel, and round 2 screened 32 redesigned guides against the eight panel genes that had not been edited, using the same protocol document resubmitted without edits (**figs. S37 and S38**).

Module 1 annealed the guide oligonucleotide pairs. Complementary gRNA oligo pairs bearing BsaI-compatible overhangs were ordered in 96-well plates and 5′-phosphorylated in 50 μL reactions containing up to 300 pmol of 5′ termini, 1× T4 PNK reaction buffer, 1 mM ATP and 10 U of T4 polynucleotide kinase, incubated at 37 °C for 30 min and heat-inactivated at 65 °C for 20 min. Equimolar forward and reverse oligos were annealed on the thermocycler by holding at 95 °C for 2 min and cooling from 95 °C to 25 °C over 45 min, and the resulting duplexes were diluted 1:10 to 1:200 in nuclease-free water.

Module 2 assembled the guide plasmids by Golden Gate cloning. The Fluent then combined each diluted duplex with a Golden Gate master mix in 12 μL reactions containing 5 μL of NEBridge Ligase Master Mix, 1 μL of BsaI-HF v2 and approximately 100 ng of the BsaI acceptor backbone, and the reactions were cycled on the automated thermocycler for 30 cycles of 37 °C for 1 min and 16 °C for 1 min, followed by a BsaI heat-kill at 60 °C for 5 min, so that each spacer was inserted into the backbone in a single one-pot Golden Gate reaction.

Module 3 transformed 8 μL of each assembly into homemade 96-well chemically competent *E. coli* DH5α cells. Reactions were held on a cold block for 30 min, heat-shocked at 42 °C for 60 s on the thermal block, returned to the cold block for 2 min, recovered in 100 μL of SOC medium at 37 °C for 1 h with shaking, and the outgrowth was dispensed by the Fluent onto 8-well Omnitray LB agar plates containing 100 μg/mL ampicillin. Plates were incubated in the Cytomat at 37 °C for 16 h until single colonies appeared. In Module 4, PIXL picked one colony per guide while preserving the well mapping and inoculated it into 1 mL of LB with 100 μg/mL ampicillin in 96-deepwell plates for overnight growth at 37 °C and 900 rpm. Plasmids were extracted from the cultures on the Fluent using the PureLink Pro Quick96 kit, eluting in 50 μL and targeting at least 500 ng of plasmid per guide so that each downstream yeast reaction could receive 500–1000 ng.

Module 5 screened the verified guide plasmids for on-target cutting activity before the knockout construction. Each guide plasmid was transformed alone, without a repair donor, into the CEN.PK strain with SaCas9 integration by automated LiAc/PEG transformation. Yeast cells were grown overnight in YPAD at 30 °C, sub-cultured to an $OD_{600}$ of approximately 0.2, grown for 4 h to an $OD_{600}$ of 0.8–1.0, washed twice with sterile water and concentrated to 30 µL per reaction; each reaction combined 240 µL of 50% (w/v) PEG 3350, 36 µL of 1 M lithium acetate, 10 µL of boiled and snap-cooled carrier ssDNA, the concentrated cells and 500–1000 ng of guide plasmid, and was heat-shocked in the Cytomat at 42 °C for 3 h with shaking at 900 rpm. Transformants were washed on the automated centrifuge into SD-URA with 20 g/L glucose, Echo-spotted onto SD-URA agar containing 200 µg/mL G418 to maintain the Cas9 plasmid and incubated at 30 °C for 2 days. Because double-strand breaks that cannot be repaired are lethal in yeast, an active guide depletes colonies relative to a non-targeting control. Module 6 scored guide activity from these colony counts. The resulting gRNA efficiencies were then used as feedback for the design of the next round or the construction of knockout strains.

**β-carotene knockout strain construction**
Module 1 reconciled the target panel and assigned the donors. A target-tracking worksheet was built with one row per target, recording the target identifier and the intended edit class. Module 2 built the repair templates. Each homologous-recombination donor was amplified by overlap-extension PCR, fusing the 5′ and 3′ flanks of the target ORF so that repair at the Cas9 cut site deletes the coding sequence; 50-µL reactions contained 1× Q5 reaction buffer, 200 µM dNTPs, 0.5 µM of each primer, 3% DMSO and 0.02 U/µL Q5 High-Fidelity DNA polymerase, and were cycled at 98 °C for 30 s, 17 cycles of 98 °C for 10 s, 57 °C for 10 s and 72 °C for 10 s, and a final extension at 72 °C for 2 min.

Module 3 delivered each guide and its corresponding donor into the production strain. Each guide plasmid and its homology donor were then co-transformed into the β-carotene strain using the same automated LiAc/PEG workflow as the killing screen, with the donor added to the transformation mix. Transformants were washed on the automated centrifuge, Echo-spotted onto SD-URA plus G418 Omnitray plates and grown for 2 days at 30 °C. A positive-control knockout was carried through this and the following module as the baseline pigment reference.

Module 4 picked three colonies per transformant for inoculation and performed colony PCR for genotyping. Modules 5 and 6 then pre-cultured the selected strains and ran a four-day fermentation. Module 7 quantified β-carotene production for each knockout strain. Harvested cells were pelleted, resuspended in 300 µL of 3 N HCl, lysed by boiling for 5 min followed by 5 min on a cooled thermal block, washed with dd$H_2O$, and extracted into 300 µL of DMSO; debris was removed by centrifugation and the pigmented supernatant was transferred for reading. Absorbance at 454 nm was measured on the Tecan Infinite plate reader, and each well was normalized as a β-carotene index of $A_{454}$ divided by $OD_{600}$ and reported as fold-change relative to the positive control. A screening plate carried 11 target-gene knockouts alongside triplicate non-targeting and positive controls.

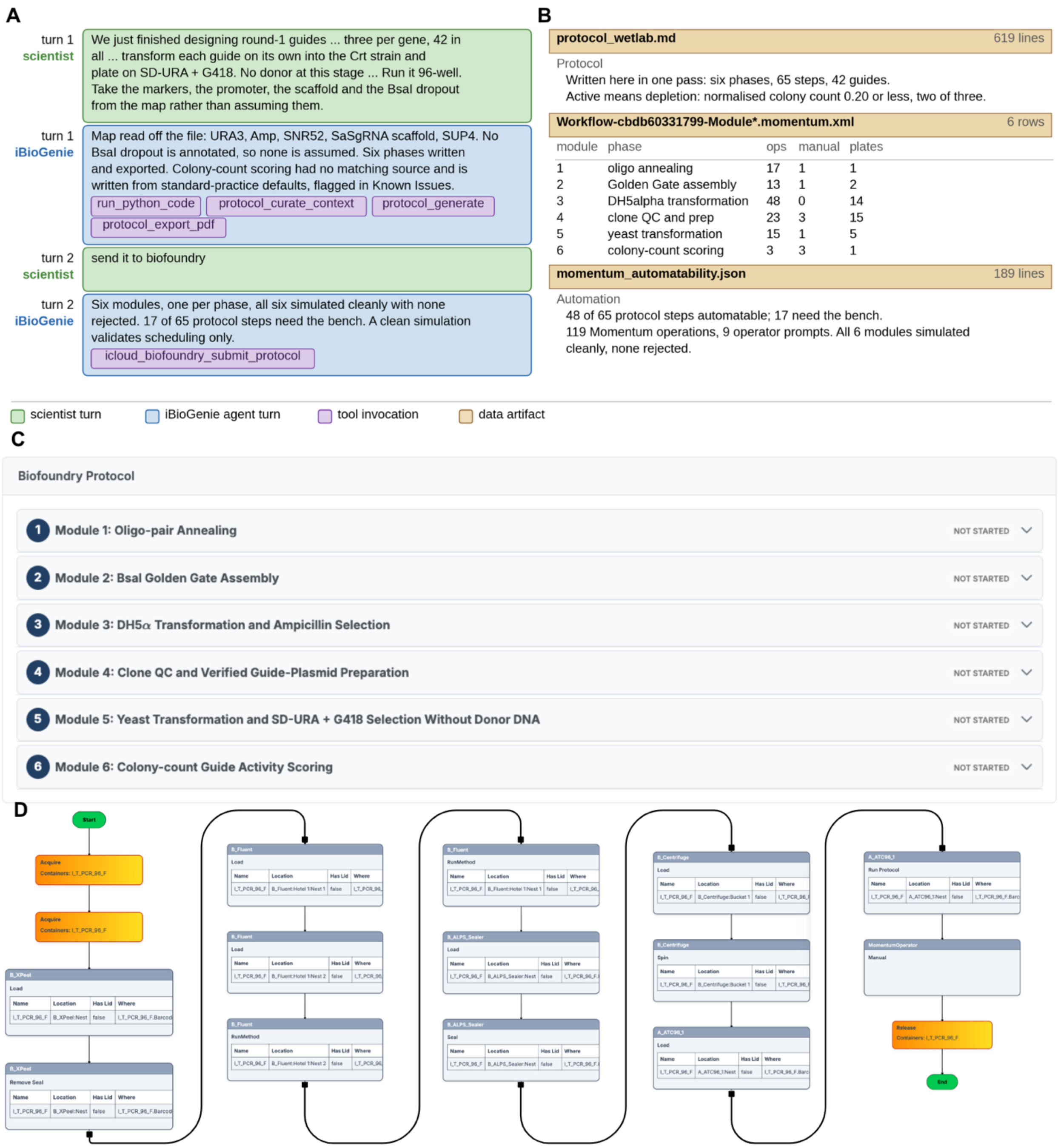


**Fig. S37. Round-1 guides screened by *S. cerevisiae* killing screen.** iBioGenie session writes and submits the bench protocol that scores the 42 round-1 guides by colony depletion on the 14-gene *S. cerevisiae* CEN.PK β-carotene panel. A clean simulation validates scheduling, labware and routing, not experimental correctness. (**A**) A summary of the chat session for designing wetlab protocol for killing assay with submission to BiofoundryAI. The opening ask supplies the plasmid map and directs the agent to take the markers, promoter, scaffold and BsaI site from it. (**B**) Delivered artifacts: the 619-line protocol and its guide-call rule, where active denotes a normalized colony count at or below 0.20 in two of three replicates; the six Momentum modules with their operations, operator prompts and plates, all six modules passing simulation with none rejected. (**C**) The same protocol in the web application as the six modules built from it. (**D**) Momentum workflow graph for module 2, BsaI Golden Gate assembly.

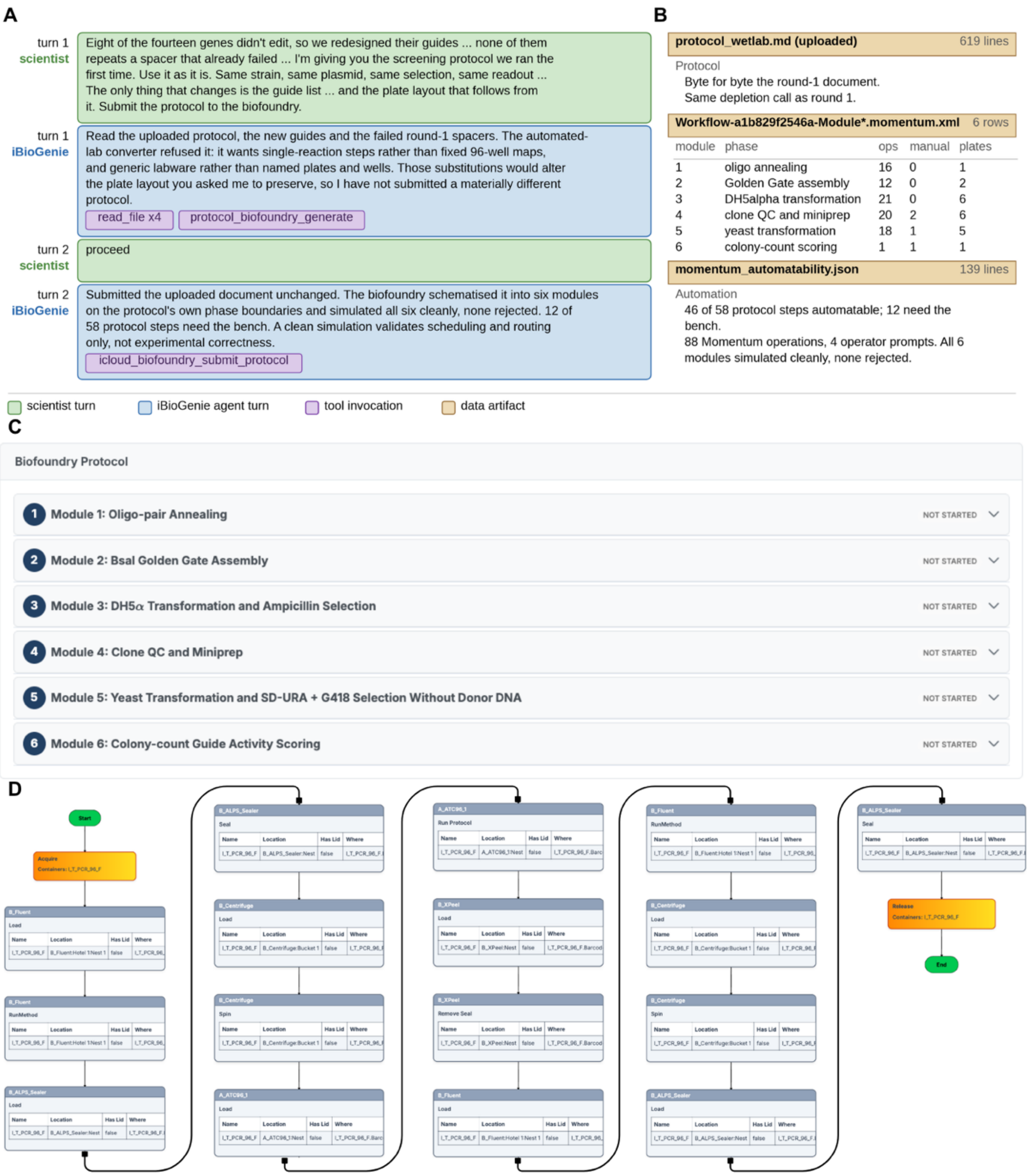


**Fig. S38. Round-2 guides screened by *S. cerevisiae* killing screen.** The eight panel genes that did not edit received four new guides each, and iBioGenie session resubmitted the round-1 screening document for them without edits. A clean simulation validates scheduling, labware and routing, not experimental correctness. (**A**) In the intervening turn, the agent refuses the document because it requires single-reaction steps and generic labware in place of the fixed 96-well maps the protocol specifies, and the agent submits unchanged only after the scientist confirms. (**B**) Delivered artifacts: six Momentum modules and 88 operations built from a document byte-identical to the round-1 submission, with 46 of 58 steps automatable and all six modules simulating cleanly. (**C**) The same protocol in the web application as the six modules built from it. (**D**) Momentum workflow graph for module 1, oligo-pair annealing.

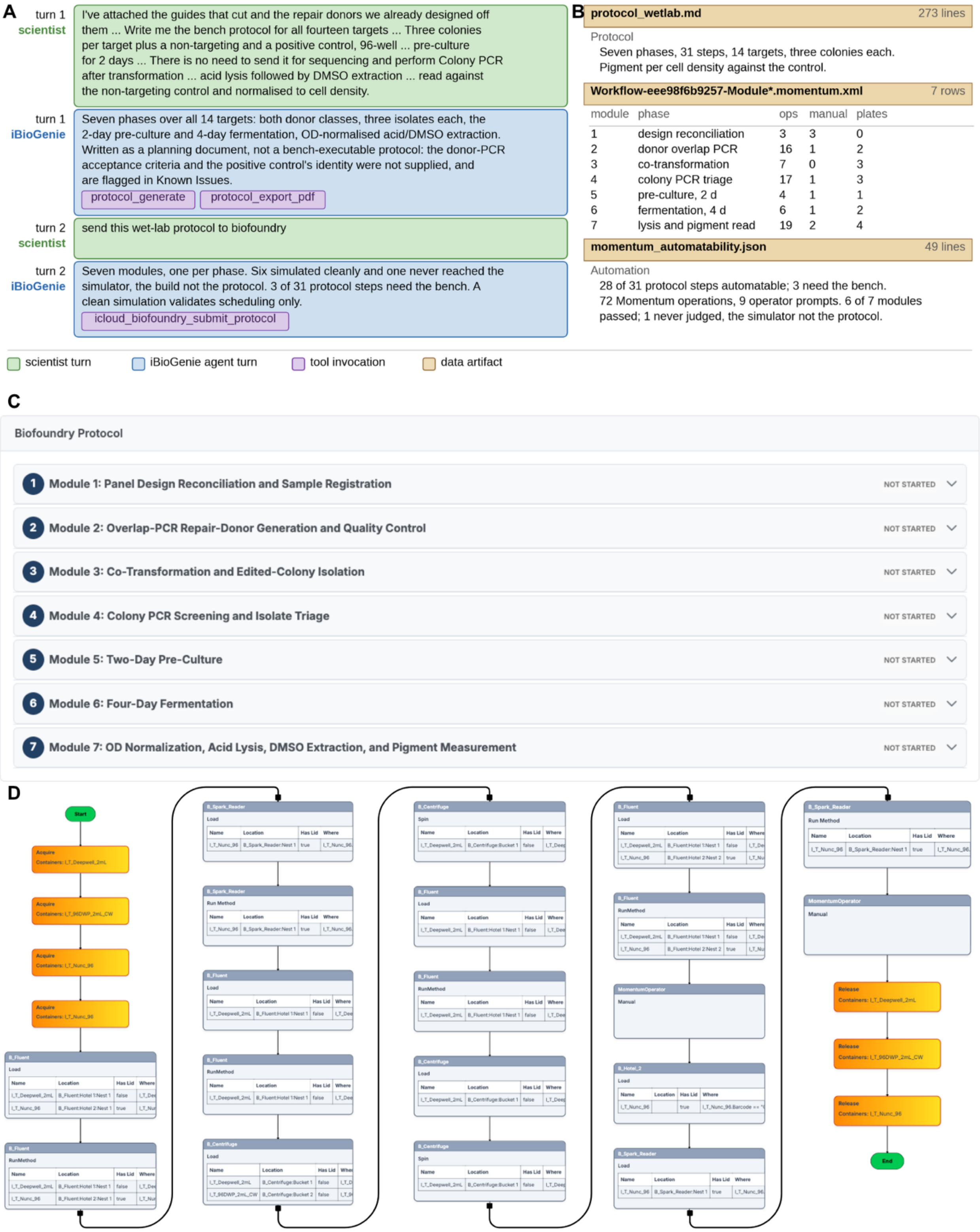


**Fig. S39. Fourteen-target knockout panel construction and β-carotene detection.** An iCloudBiofoundry session drafts and submits the protocol that builds the fourteen knockouts and reads their pigment. (**A**) A summary of the session where iBioGenie helps design the seven-phase wet lab protocol and flags potential issues including donor-PCR acceptance criteria and positive control were not supplied. (**B**) Delivered artifacts: the 273-line protocol, seven phases over 14 targets at three colonies each, its readout pigment normalized to cell density against the non-targeting control. Six modules simulated cleanly and one never reached the simulator, reported as not simulated rather than as failing. (**C**) The same protocol in the web application as the seven modules built from it, from panel reconciliation and donor generation through co-transformation and colony-PCR triage to fermentation and the pigment read. (**D**) Momentum workflow graph for module 7, OD normalization, acid lysis, DMSO extraction and pigment measurement.

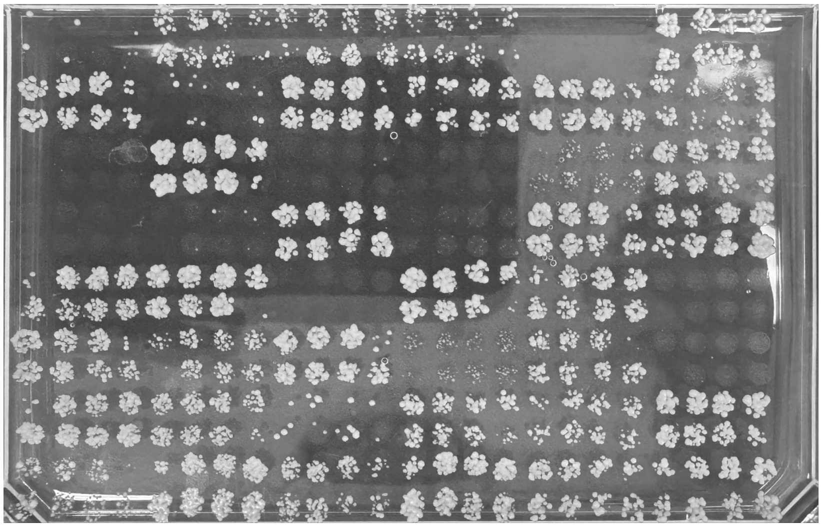

**Fig. S40. Round 1 of the guide RNA killing assay plated by Echo dispenser.** Each position is one yeast transformation from the 42 guides round-1 panel, dispensed onto SD-URA agar containing 200 µg/mL G418 on an Omnitray layout. Plates were incubated at 30 °C for 48 h. The spot-to-spot difference visible here is the assay: with no repair donor supplied, a position carrying an inactive guide grows to confluence, whereas an active guide cuts the genome and depletes its position to a sparse or empty spot.

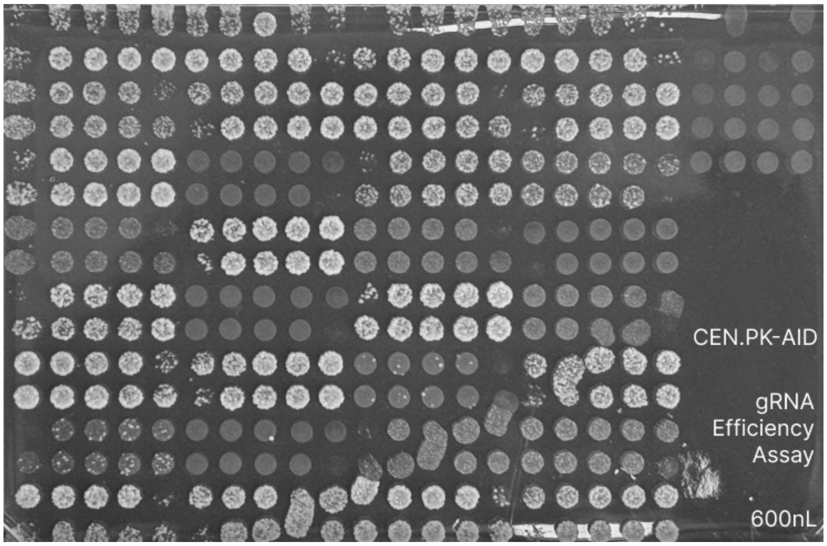


**Fig. S41. Round 2 of the same screening, run on the 32 redesigned guides built for the eight panel genes that did not edit in round 1.** Medium, selection, and incubation are identical to **fig. S40**; only the guide set and the plate layout differ. The on-plate annotation records the assay and the 600 nL acoustic droplet volume dispensed per position.

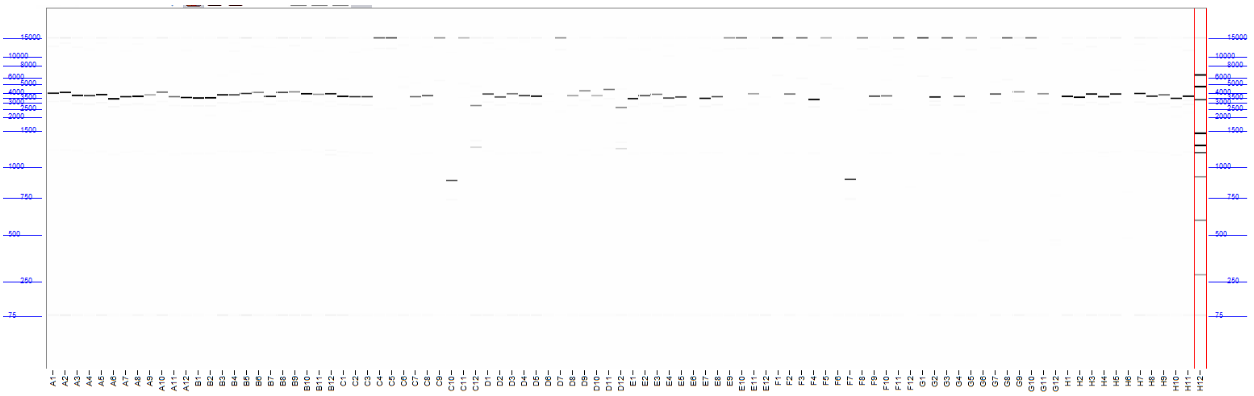

**Fig. S42. Quality check and verification of first round of gRNA plasmid by capillary electrophoresis (Agilent Fragment Analyzer, kit #DNF-920).** Observed fragment sizes were compared to expected product sizes to confirm gRNA plasmid by enzyme digestion.

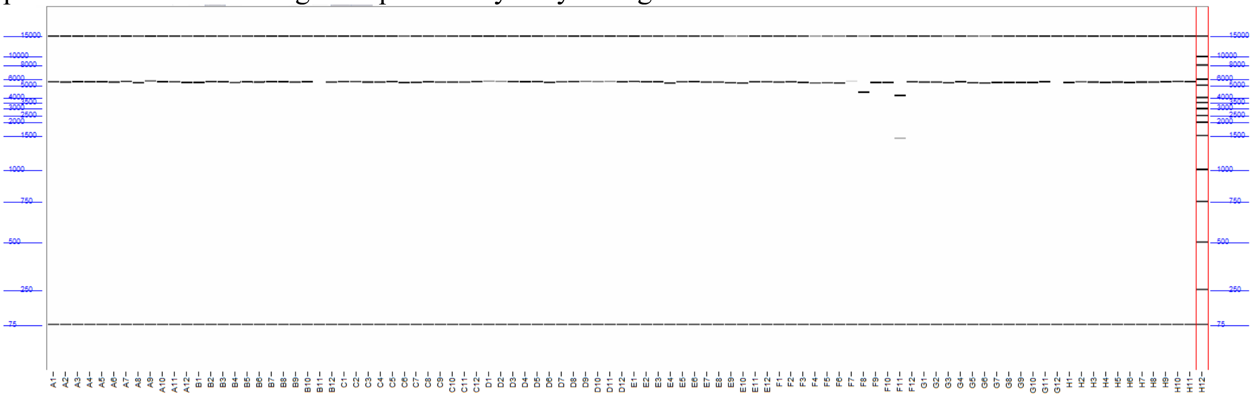

**Fig. S43. Round 2 guide plasmids verification (Agilent Fragment Analyzer, kit #DNF-920), lanes again labelled by source well.** Digestion conditions, acceptance criterion and marker positions are as in fig. S42, applied unchanged to the redesigned guide set.

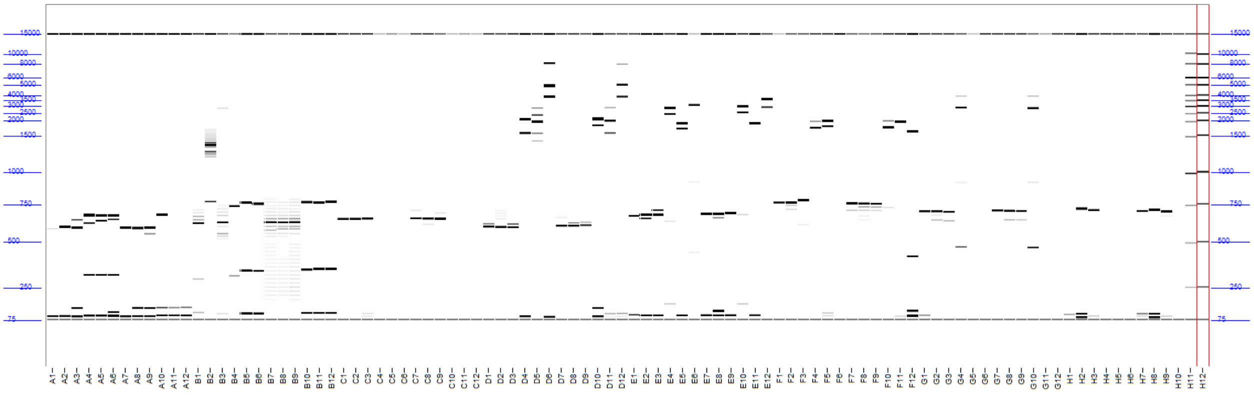

**Fig. S44. Genotype verification of the knockout isolates by capillary electrophoresis (Agilent Fragment Analyzer, kit #DNF-920).** Each lane is a colony PCR product spanning the targeted locus of one isolate, labelled by its source well. Because each of the 14 targets carries its own flank design, the expected amplicon size differs from target to target, and an edit is confirmed when the product matches the size predicted for the donor-repaired deletion allele rather than the unedited locus. Consistent with the ranking design of the screen, genotyping was performed retrospectively rather than as a gate before culture.

### 10. Conclusion on case studies

Across three distinct case studies, iCloudBiofoundry was evaluated for both its scientific design capabilities and its ability to translate those designs into executable experimental workflows. The studies covered different types of synthetic biology problems and required substantially different computational reasoning, design strategies, and experimental procedures. In all three cases, the platform was able to successfully coordinate the required design and execution steps, demonstrating its potential as a versatile system for a broad range of synthetic biology applications. All three case studies involved complex, multi-stage computational and experimental workflows requiring the coordinated use of specialized tools, instruments, and intermediate outputs that would otherwise be difficult to assemble and manage manually (**table S25**). iCloudBiofoundry was able to manage these complex workflows and show the practical potential of the platform to make AI-guided, automated experimentation more accessible and to help democratize a scientific workflow in which complex biological design and physical experimentation can be coordinated through a unified interface.

**Table S25. Number of computational steps, automation modules, devices, and experimental operations required across the iCloudBiofoundry case studies.**

| Experiment | Computational steps | Experimental modules | Unique devices | Experimental Steps |
|---|---|---|---|---|
| Designing improved phytase enzyme variants | 6 | 7 | 13 | 212 |
| Enzyme discovery for improved 3HP production | 7 | 8 | 14 | 176 |
| Enzyme engineering for improved 3HP production | 11 | 7 | 16 | 189 |
| engineering of *S. cerevisiae* for β-carotene production | 7 | 6+6+7 | 15 | 488 |
| Total | 31 | 41 | 19 | 1065 |
